# Comprehensive profiling of L1 retrotransposons in mouse

**DOI:** 10.1101/2023.11.13.566638

**Authors:** Xuanming Zhang, Ivana Celic, Hannah Mitchell, Sam Stuckert, Lalitha Vedula, Jeffrey S. Han

## Abstract

L1 elements are retrotransposons currently active in mammals. Although L1s are typically silenced in most normal tissues, elevated L1 expression is associated with a variety of conditions, including cancer, aging, infertility, and neurological disease. These associations have raised interest in the mapping of human endogenous *de novo* L1 insertions, and a variety of methods have been developed for this purpose. Adapting these methods to mouse genomes would allow us to monitor endogenous *in vivo* L1 activity in controlled, experimental conditions using mouse disease models. Here we use a modified version of transposon insertion profiling, called nanoTIPseq, to selectively enrich young mouse L1s. By linking this amplification step with nanopore sequencing, we identified >95% annotated L1s from C57BL/6 genomic DNA using only 200,000 sequencing reads. In the process, we discovered 82 unannotated L1 insertions from a single C57BL/6 genome. Most of these unannotated L1s were near repetitive sequence and were not found with short-read TIPseq. We used nanoTIPseq on individual mouse breast cancer cells and were able to identify the annotated and unannotated L1s, as well as new insertions specific to individual cells, providing proof of principle for using nanoTIPseq to interrogate retrotransposition activity at the single cell level *in vivo*.

## INTRODUCTION

Transposable elements (TEs) are often referred as “genetic parasites” that have thrived and mobilized in mammalian genomes for millions of years (1, 2). TEs have played a critical role in shaping human genomes, making up at least 45% of the human genome, with <u>L</u>ong <u>In</u>terspersed <u>E</u>lements (LINE-1, L1) comprising 17% (1). L1 has been extensively investigated not only due to its abundance in the genome but also its activity in humans, which can drive mutations and endanger the integrity of the genome (3–7). L1 mobilizes in the genome via a “copy and paste” mechanism, utilizing two proteins encoded by the L1 ORF1 and ORF2 genes (8, 9). L1 ORF1p possesses nucleic acid binding activity (10), presumably facilitating the binding of L1 mRNA in *cis* (11) to form the ribonucleoprotein (RNP) complex in the cytoplasm (12–15). The L1 RNP complex is then transported to the nucleus through an uncertain mechanism which may vary depending on cell type (16, 17). ORF2p contains an endonuclease domain (EN) that nicks the genome at the 5’-TTTT/AA-3’ preferred insertion site (8, 18), followed by converting L1 mRNA to cDNA using ORF2p reverse transcriptase (RT) (19). This process is called target-primed reverse transcription (TPRT), and the mobilization of L1 is called retrotransposition (20, 21). L1 RT is susceptible to early termination often resulting in 5’ truncated L1 insertions, which are unable to further retrotranspose in the genome. It has been reported that the majority of new L1 insertions are caused by a few “hot” transposable full-length L1s (3). The human genome has ∼100 L1s capable of retrotransposition (3, 5) and L1 dysregulation has been shown to be associated with many health-related issues (6, 7, 22–28). Mouse models exist for many of these conditions; for example, L1 overexpression is observed in piRNA mutant male mice germ cells, where massive DNA damage and meiotic arrest occur (29–31). The presumption is that L1 overexpression leads to these genotoxic effects due to endonuclease-mediated cutting of DNA and possibly new L1 insertions. However, to date there is no clear-cut, direct evidence for meiotic *de novo* retrotransposition of endogenous L1s in these mice. Tracking endogenous L1 insertion activity in various mouse disease models would be beneficial for understanding the role and impact of L1 on these conditions. In addition, L1 has been found to insert at CRISPR-cas9 editing sites (32), demonstrating the potential impact of L1 on CRISPR related gene therapy and the importance to be able to monitor these events. A variety of methods have been used to profile human L1 insertions, including bioinformatic analysis of high coverage whole-genome sequencing (WGS) and targeted L1 enrichment followed by sequencing (33–36). Early forays into profiling mouse L1s have begun but can be costly due to the amount of sequencing data required (37, 38).

TIPseq (transposon insertion profiling by sequencing) is a promising method for comprehensive profiling of fixed, polymorphic, and *de novo* transposon insertions (35, 39). In TIPseq, a transposon specific primer is used in a ligation-mediated PCR to enrich for transposon sequences. The resulting amplicon mixture is subjected to paired-end short read deep sequencing. This method has been used to successfully profile the currently active family of human L1 retrotransposons, L1Hs, in a variety of samples (35). The success of this protocol relies on the specificity of the L1Hs-specific primer. Adapting TIP-seq to mouse genomes is desirable because it would allow us to monitor endogenous *in vivo* L1 activity in controlled, experimental conditions using mouse disease models. However, adapting L1 TIPseq to mouse genomes presents additional challenges. In mouse (as compared to humans), not only are there more active transposons (40), there are multiple active L1 subfamilies (Tf, Gf, and A) (2, 41). Within the active mouse L1 subfamilies, there are more young, potentially active family members. There is an estimated 30x more active L1s in mouse as compared to the active L1 population in humans (41). Thus, we would expect that comprehensive profiling of this larger and more complex population of young L1s in mice would require greater sequencing depth and specificity.

Here, we describe a modified TIPseq protocol to capture young, actively retrotransposing L1 elements in mouse. We focus on the L1MdTf subfamily, as this family is responsible for the vast majority (>80%) of *de novo* L1 insertions in mouse (38, 42). This modified protocol utilizes long read Nanopore sequencing and allows the comprehensive, unambiguous identification of fixed, polymorphic, and *de novo* L1MdTf elements from pooled or single cells. As little as 200,000 reads (115 million bases (MB)) of sequencing data can be used to identify 98% of L1MdTf elements in a genome, making this a cost-effective method of transposon profiling that can be routinely implemented even by smaller labs.

## MATERIAL AND METHODS

### Genomic DNA isolation

Mouse tail (1cm) was used to extract the genomic DNA (gDNA) using NEB Monarch® Genomic DNA Purification Kit (NEB T3010L). DNA concentration was measured on Qubit dsDNA BR assay kit (cat: Q32856) and visualized on 1% agarose gel.

### Library preparation and sequencing

Purified gDNA was sheared to an average size of 1500 bp using ultrasonication (Covaris M220). Library input was examined on an Agilent bioanalyzer 2100 using the high sensitivity DNA Kit (cat: 5067-4626). The sheared DNA was end polished (NEBNext® Ultra^™^ II End Repair/dA-Tailing Module E7546S), followed by ligation to T-overhang vectorette adapters (39, 43) (NEBNext® Quick Ligation Module E6056S; see Table S1 for primer sequences). We recommend cleaning up the ligation product before PCR amplification to remove excessive adapters which minimize potential undesired concatemers and other non-specific amplification. We used bead purification (AMPure XP Reagent A63880, 1x ratio) to remove small DNA fragments and excessive adapters. Touchdown PCR (Table S2) with Platinum^™^ II Taq Hot-Start DNA Polymerase (cat: 14966001) was performed to balance yield and specificity, using an L1 specific primer and vectorette primers. The PCR products were cleaned up with beads (0.7x ratio) and quality controlled using Qubit assays and the Agilent bioanalyzer 2100. For short read sequencing, PCR amplicons were sheared to a size of ∼400 bp and sequenced by BGI (150 bp paired-end reads, 30 million reads ∼10Gb data).

To sequence using Oxford Nanopore (ONT), we used a MinION sequencer with R10.4.1 flow cells. Purified TIPseq PCR amplicons were used to construct the ONT library using Native Barcoding Kit 24 V14 (SQK-NBD114.24). Sequencing was carried out in house to our desired sequence depth. For whole genome sequencing, high molecular weight gDNA was used as the direct input for the Native Barcoding Kit 24 V14.

### L1 annotation curation

TIPseqHunter utilizes a small set of 200 high-confidence “fixed present” L1HS insertions in humans, providing essential alignment characteristics for building an accurate machine learning model (35). We downloaded all L1MdTf1-3 elements from the UCSC table browser using the filter option (mm39, filtered for L1MdTf*) for mouse L1 annotation. We manually curated the annotation file to better represent the active L1 population of interest to us. We removed L1s that did not contain the JH922 primer sequence, as they were presumably older elements not relevant for our purpose. We also separated individual elements that were bookended together erroneously as a single L1 element (e.g. Table S3). This step effectively removed most L1MdTf3 elements (older elements) and left us with 3266 elements, which better represented a younger and active L1s in the mouse genome.

### TIPseqHunter data analysis

For short-read TIPseq data, we ran the TIPseqHunter pipeline as previously described (35, 39). Briefly, pair-end reads were quality controlled and trimmed to remove Illumina adapter sequences, vectorette adapter sequences, and reads with poor quality scores. The reads were subsequently mapped to the mm39 reference genome where L1MdTf1-3 elements were masked,using bowtie2/2.3.3 with the following settings: -X 1000 --local --phred33 --sensitive. l1hskey was changed to G(6833) to map the 5’ most position of TuJH922 primer in the mouse L1MdTf_1 consensus sequence. We replaced the human L1 reference with our customized mouse L1 annotation file.

For downsampling, seqtk sample was used to scaledown to 0.01x, 0.1x, 0.2x, 0.5x of the original fastq files in triplicates using different seeding keys. The scaledown fastq files were used to run TIPseqHunter as described above.

### Long-read TIPseq data analysis

The raw sequencing data was processed during sequencing runs by the MinKNOW application (fast basecalling model, trim barcodes on). Reads were aligned to the mm39 reference genome using the integrated aligner, minimap2. Default settings were applied, producing alignments in the bam format. To extract the clipped sequences from each alignment, the SamExtractClip tool (Jvarkit) was utilized. These clipped sequences were then remapped to the L1MdTf_1 consensus sequence (44) using bowtie2. A customized scoring system (--sensitive -N 1

–mp 1,1 –rdg 5,2 –rfg 5,2) was employed to achieve a balance between sensitivity and accuracy. Reads that mapped to the L1MdTf_1 consensus sequences were filtered based on the presence of a poly-A tail at the end of the alignment. By default, the filtering required at least 5 consecutive As at the end. The filtered reads were subsequently used to retrieve the original mapping location in mm39 using the read name where the mapped intervals from were added to the potential insertions list, which was formatted as a bed file. The bedtools cluster function (45) was employed to identify regions with individual reads. Regions with more than 3 supporting reads were merged using the bedtools merge function, resulting in the final set of insertion candidates. The precise insertion site was determined using an in-house script that based on the clipped position obtained in the header region from the SamExtractClip. The full pipeline is available at https://github.com/JHanLab/NanoTipSeq.

### Single cell whole genome amplification

Cultured 4226 cells (46) were washed with PBS and stained with propidium iodide. Single live cells were sorted into individual wells of a 96 well lo-bind plate by the Louisiana State University CIMC core, based on PI staining, forward scatter, and side scatter. Freshly sorted single cells were used to perform single-cell whole genome amplification (WGA) by either MDA (multiple displacement amplification) or PTA (primary template directed amplification) following manufacturer recommendations. Qiagen REPLI-g single cell kit (cat 150343) was used for MDA, and BioSkryb Genomics ResolveDNA®kit was used for PTA. The amplified genomes were quality controlled using Qubit assays and the Agilent bioanalyzer 2100. Amplified DNA was used in our TIP-seq protocols, library construction, and sequencing as described above.

## RESULTS AND DISCUSSION

### L1 primer design

The L1MdTf lineage contains 3 families, namely L1MdTf1, L1MdTf2 and L1MdTf3. These families are responsible for all of the reported spontaneous *de novo* germ line insertions in the literature (42). Because we were unable to identify primers that would specifically amplify all 3 L1MdTf families, we chose to focus on targeting L1MdTf1 and L1MdTf2, as these two families account for 80% of the reported spontaneous *de novo* germ line insertions and are more closely related to each other than to L1MdTf3. L1MdTf1 and L1MdTf2 are also younger than L1MdTf3, with average age of 0.25 million years and 0.27 million years, respectively (2). Thus, we reasoned that monitoring L1MdTf1 and L1MdTf2 insertions would be an acceptable proxy for L1 activity. Our strategy to target L1MdTf1/2 elements involved designing L1 primers in the L1 3’ UTR region, where the Tf subfamilies exhibited substantial differences from other subfamilies. One of the Tf1/2-specific primers is shown in figure 1A. We inspected the multi-sequence alignment of consensus sequences from each family. We found a conserved 17bp region where an ATA->GTG change appears to be diagnostic for L1MdTf1 and L1MdTf2. We then designed the TuJH922 primer spanning this region with 3’ end terminating at this diagnostic GTG trinucleotide. We performed an *in silico* search for TuJH922 sequence in all C57BL/6 L1 elements (total of 410549 elements). A perfect match for TuJH922 was found in 56.6% of L1MdTf1 (1479/2613) and 51.3% of L1MdTf2 (1950/3801) elements but rarely in other subfamilies (24 total instances from all other subfamilies combined). The phylogenetic tree shown was built using the L1 ORF2 sequence (2) (Fig. 1B). Moreover, all previously identified “hot” L1MdTf1/2 elements or L1MdTf1/2 *de novo* germ line insertions contain an exact match for TuJH922 in their 3’ UTR (when sequence spanning that region is available to examine), (42, 47) indicating the suitability of TuJH922 to target young, potentially transposable L1 elements in the mouse genome.

**Fig. 1.**
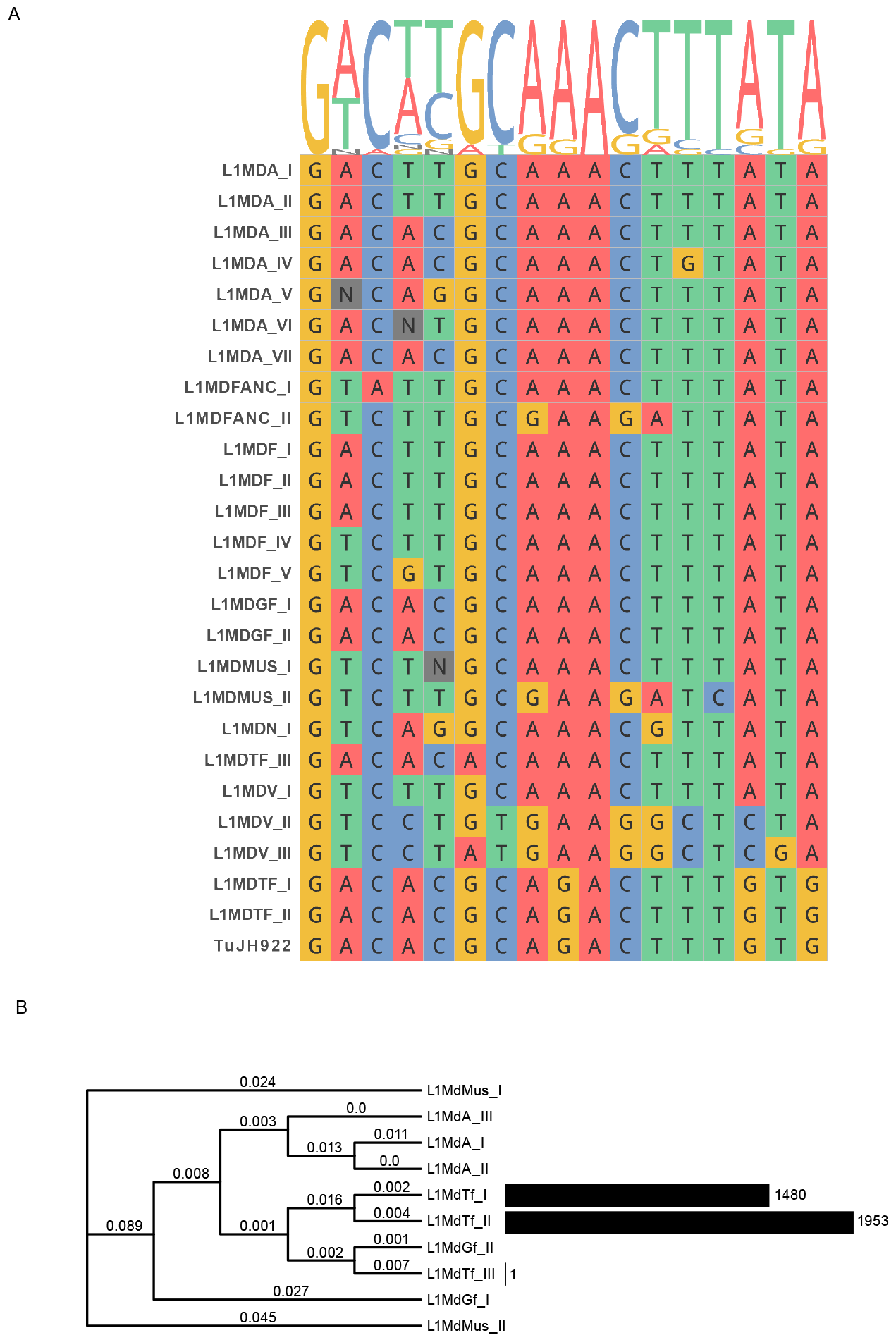
Design of L1MdTf specific primer. (**A**) Mouse L1-specific primer design based on alignment of consensus sequences from major L1 subfamilies. We selected TuJH922 for its diagnostic trinucleotides “GTG” in the 3’ end, which distinguish young active L1s from inactive old L1s. (**B**) Number of elements containing an exact match of TuJH922 primer sequence for each L1 subfamily.

### Mouse L1 TIPseq using short-read sequencing

The original TIPseq protocol relies on ligation-mediated PCR to selectively amplify L1Hs from human DNA, involving five main steps: 1) Genomic DNA (gDNA) extraction and digestion. Six 5 or 6 base cutter restriction enzymes are used individually to digest the genome to achieve even fragmentation of gDNA. 2) Vectorette adapter ligation. The vectorette adapters consist of two short oligonucleotides that are complementary on both ends but mismatched in the center. This structure allows amplifying a region of interest with knowing only one end of the target. 3) TIPseq PCR by using an L1 specific primer and vectorette primer. The vectorette primer has identity to the bottom mismatched strand of the vectorette adapter, which has no complementary sequence to anneal to. The complement of the bottom strand of the vectorette adapter is only present after synthesis from the L1 primer, allowing specific amplification of L1 adjacent sequence. 4) Amplicon shearing and 150 bp paired-end sequencing. 5) Machine-learning algorithm to identify annotated L1s and new L1 insertions.

We used C57BL/6 genomic DNA to evaluate the sensitivity of the TIPseq pipeline for identifying L1 elements in the mouse genome through short-read sequencing (Fig. 2A). TIPseq PCR amplicons were sequenced to a depth of 30 million reads (150 bp x2, 10Gb of data) for each sample. We employed alignment evidence such as genomic-genomic read pairs and L1-genomic junction reads (Fig. 2B, C) to detect L1 elements in the short-read data. TipseqHunter (35, 39) was used to analyze the short-read data, and we successfully identified over 90% (∼3000) of pre-existing L1MdTf1 and L1MdTf2 (annotated L1s) using different L1 specific primers (Fig. 2D). The primers performed similarly with respect to L1 identification and 3’ UTR coverage (Fig. 2D, E). To assess primer specificity, we aligned reads to the L1MdTf_1 consensus sequence and found that 99% of reads contained the diagnostic “GTG” nucleotides (Fig. S1). In contrast, TIPseq amplicons generated with a less specific primer (TuJH801) resulted in only 25% of reads containing the GTG nucleotides and were largely comprised of older mouse L1s. To test how much data we needed to achieve good recovery of annotated L1s, we employed multiple rounds of randomly selecting 50%, 20%, 10%, and 1% of the original data and analyzing with the TipseqHunter pipeline. With as little as 10% of the original data, we were able to detect over 79% of annotated L1MdTf1 and L1MdTf2 L1s from sample using the TuJH922 primer (Fig. 2D, Fig. S2). For samples made from TuJH922, the average annotated L1 recovery rate was 90%, 88%, 84%, 79% and 44% for 100%, 50%, 20%, 10% and 1% of the original data, respectively. These results demonstrate that mouse TIPseq coupled with short-read sequencing can detect up to 80% of pre-existing L1 elements using as few as 2 million reads (1Gb).

**Fig. 2.**
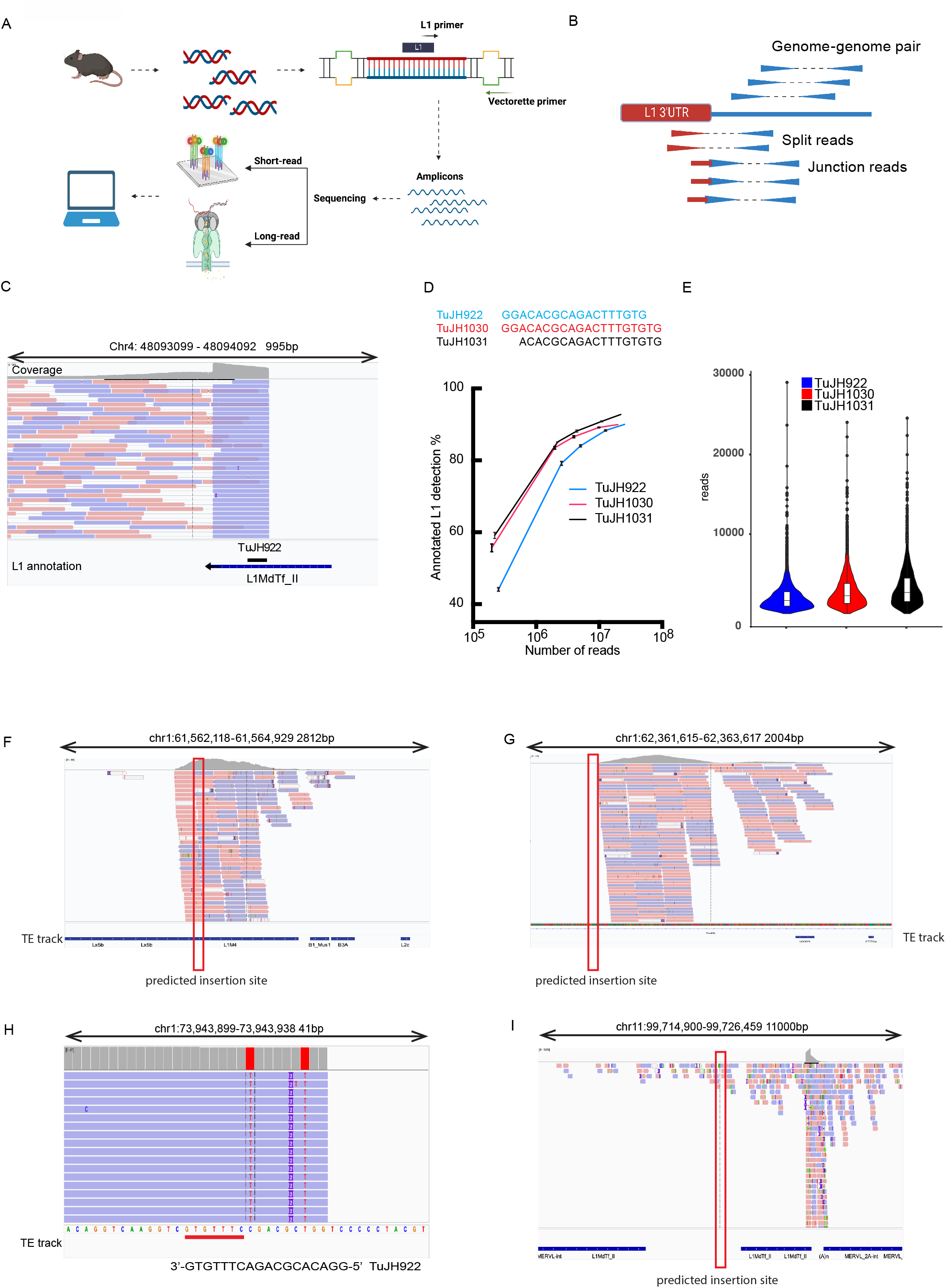
Short-read TIPseq identifies annotated L1s in the mouse genome. (**A**) TIPseq workflow consists of five main steps: 1) genomic DNA extraction and fragmentation 2) vectorette adapter ligation 3) TIPseq PCR using mouse L1 specific primers 4) sequencing the amplicons 5) customized bioinformatic pipeline analysis. (**B**) Alignment evidence supporting the detection of annotated L1 elements, including genome-genome pairs, junction reads, and split-reads (**C**) Example of typical mapping evidence supporting the identification of an existing L1MdTf element. (**D**) Scaling-down experiments. Original sequencing files were scaled down to 50%, 20%, 10%, 1% in triplicates and analyzed using TIPseqhunter. Percentage of annotated L1s identified are shown on Y-axis, and number of reads scaled to are shown on X-axis. (**E**) Coverage of L1 3’ 300bp flank region. The box plot displays 25% to 75% percentile with median shown as the middle line in the box. (**F**) Example of a TIPseqhunter predicted non-reference insertion that falls inside another transposable element. Because we have low confidence in short reads mapped to a repetitive region, we discarded this prediction. (**G**) Potential non-reference L1 insertion site predicted by TIPseqhunter is highlighted in red box. (**H**) False-positive predictions resulting from mis-priming of the TuJH922 primer into a similar region. (**I**) False-positive prediction with low coverage and were between two annotated L1s, possibly caused by mismapping or non-specific mapping.

Our goal in using TIPseq is to be able to detect new, unannotated mouse L1 insertions. In our original TuJH922 dataset, TIPseqHunter predicted a total of 869 candidate L1 insertions absent from the reference genome. Many of these candidate insertions were predicted to be within an existing repetitive sequence (e.g. Fig. 2F). We filtered out these candidates within repetitive sequence because we had low confidence in the accuracy and uniqueness of the alignments. This left 113 candidates for further validation (Table S4). We visually inspected alignments of these remaining candidates in IGV and checked for partial primer sequence that may have caused mis-priming incidents. After examining all candidates, we identified only 17 high-confidence potential insertions (e.g. Fig. 2G). In contrast, we found 37 mis-priming cases (e.g. Fig. 2H), 26 candidates located near an annotated L1 with very low read coverage (e.g. Fig. 2I), and 36 candidates with low mapping quality (Table S4, Fig. S3). Thus, although TIPseqHunter had the sensitivity to identify most existing, annotated young mouse L1s, the high false positive rate for unannotated insertions led us to search for improvements to identify unannotated mouse L1s.

### Mouse TIPseq with long-read sequencing (nanoTIPseq)

Short-read sequencing data has several limitations to identify L1 insertions: 1) Low mappability to highly repetitive regions, 2) Sophisticated algorithms needed to reconstruct and identify potential non-reference (polymorphic or *de novo*) insertions, and 3) greater chance of artifacts and need to infer insertion site structure due to short reads requires additional validation. To overcome these limitations, we tested TIPseq with long-read sequencing. Oxford nanopore technologies (ONT) sequencing is a 3^rd^ generation sequencing method that produces long continuous readout of DNA strands ranging from a couple hundred to millions of bases (48, 49). The advantages of ONT include portability, affordability and instantaneity which are ideal for real-time applications (49–51). Although ONT is prone to high error rates as compared to short-read sequencing methods, the long reads still usually allow unambiguous alignment, even within repetitive regions. High coverage ONT whole genome sequencing has been used to detect L1 insertions (34, 37, 38); however, whole genome sequencing may not be cost effective when examining a large number of samples, such as studies involving single cell analysis. Sequencing an L1-enriched TIPseq reaction is expected to decrease costs considerably. We used the same TIPseq amplicon mixtures previously sequenced with short-reads (Fig 2D) to prepare a library for ONT. We needed only 200,000 reads (129Mb) of the long-read data to identify, with at least 5 support reads, 89.3% of our annotated L1MdTf1 and L1MdTf2 L1s in the C57BL/6 genome (Table S5). Thus, a similar number of elements can be detected with 81x less data using long-read sequencing, when comparing short-read TIPseq. In most cases, individual reads encompass an entire amplicon, starting from the TuJH922 primer followed by the L1 3’UTR, poly(A) tail, flanking genomic sequence, and the vectorette linker (Fig 3A). The original TIPseq protocol entails digestion of genomic DNA with various restriction enzymes before vectorette adapter ligation, and long reads can be grouped into multiple populations with discrete endpoints based on restriction site positions (Fig. 3B). To directly compare TIPseq short-read and long-read data, we normalized the number of reads mapping to annotated L1 3’ UTR (starting from TuJH922 sequence to end of the element) by counts per million mapped reads (CPM) (Fig. 3C). Long reads cover annotated L1 3’ UTR 2.44x better than short reads, despite having less sequencing depth compared to short reads. In addition, long read data allowed us to detect 59% (192/326) of those annotated L1s that were not found by TIPseqHunter. To distinguish TIPseq with short reads and long reads, we call TIPseq with long reads “nanoTIPseq”. NanoTIPseq allowed us to detect L1s in highly repetitive regions, a major shortcoming of short-read sequencing. We found 3.26x better coverage in the 300 bp downstream region of the annotated L1s using long-read sequencing, demonstrating the capability of nanoTIPseq to detect L1s in a complex region (Fig. 3D, Fig. S4).

**Fig. 3.**
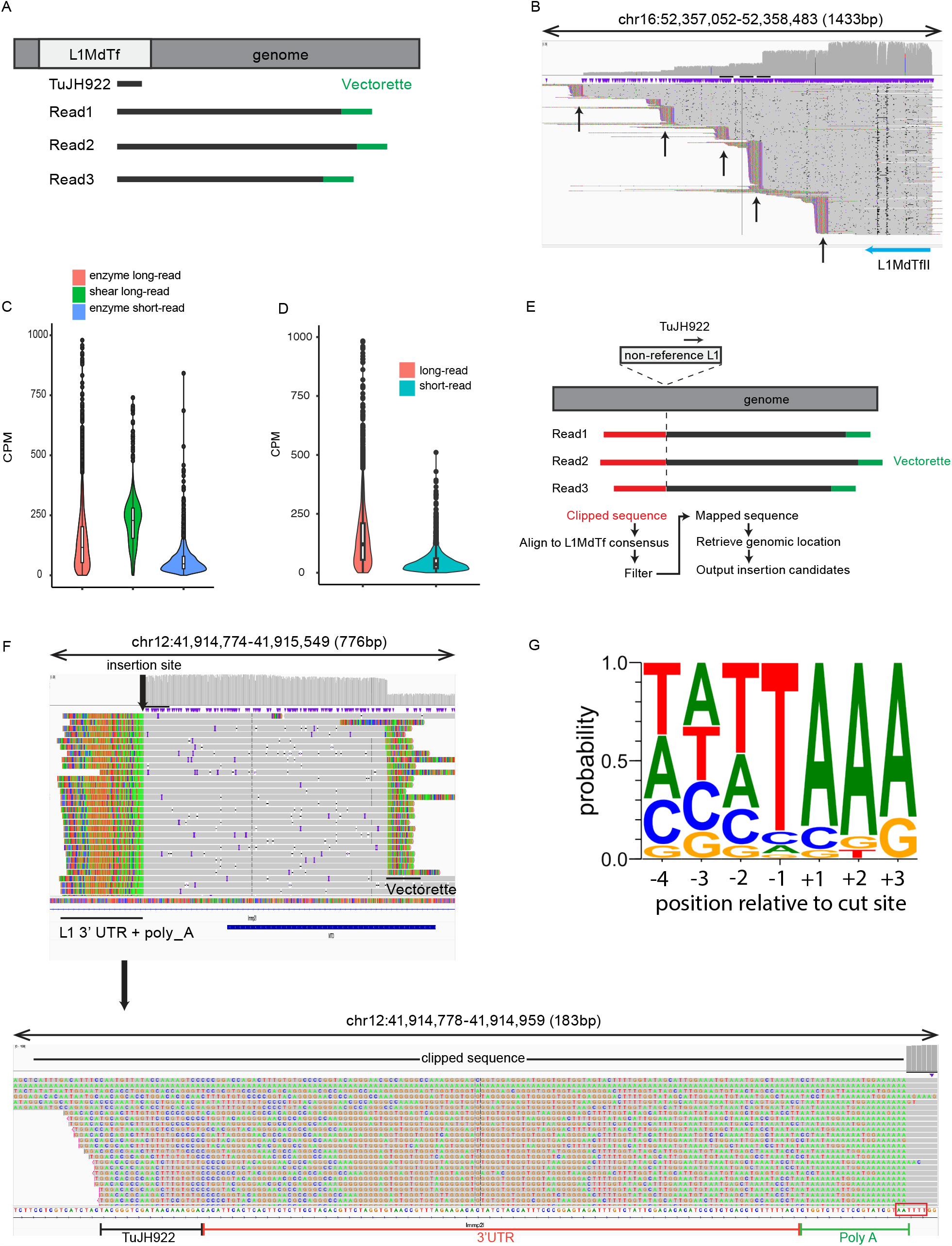
nanoTIPseq detection of non-annotated L1s in the mouse genome. (**A**) Typical alignment evidence for annotated L1 elements. One single alignment should contain L1 primer (TuJH922), L1 3’ UTR sequence, adjacent genomic sequence and vectorette adapter sequence. (**B**) Representative image showing a L1MdTfII element on chr16. Blue arrow shows the direction and location of the L1 element. Black arrow indicates the location of restriction enzyme sites used to digest gDNA. (**C**) Annotated L1 3’ UTR coverage among C57BL/6 samples processed with enzyme-digested long-read (pink), enzymatic-digested short-read (blue) and sonicated long-read (green). (**D**) Annotated L1 3’ adjacent genomic region coverage among C57BL/6 sample processed with enzymatic digestion and sequenced either with short-read (cyan) or long-read (pink). Long-read method demonstrated 3.26x better coverage than short-reads. (**E**) Scheme showing the basic idea behind nanoTIPseq. The non-reference L1 which is absent from the reference is showing in over-hang box and TuJH922 mapping location is shown. The clipped sequence showed in red and should contain TuJH922, L1 3’ UTR and a poly-A tract. The mapped sequence is shown in black, and vectorette sequence is shown in green. (**F**) A typical non-reference insertion site should consist of a clipped sequence region (non-aligned bases shown in color, A:green, T:red, G:orange, C: blue) at the 3’ integration site (zoomed in image is shown below) that contains TuJH922 sequence, L1 3’ UTR sequence and a poly-A tract. Insertion site site (5’-AAAA/TT-3’) is highlighted by the red box. There is no annotated L1 at or nearby this insertion site. This alignment evidence suggests that this is a non-reference L1 in the C57BL/6 mouse we tested. (**G**) Sequence logo of non-reference L1 insertions site recapitulates L1 ORF2p preference site TT/AAAA motif.

To discover L1 insertions not present in the reference genome, we developed a customized bioinformatics pipeline for nanoTIPseq (Fig. 3E). Briefly, at a new L1 insertion site we anticipate the inserted L1 sequence will be missing from the reference genome, thus resulting in L1 sequence being clipped from the reads during alignment. The reads will still map to the insertion site owing to the genomic sequence in the same read. We extracted the clipped sequences and aligned them to the L1MdTf_I consensus using Minimap2. Subsequently, we selected the clipped reads that mapped to the L1 3’ UTR and possessed a poly(A) tail. By utilizing alignment of flanking genomic sequence from the corresponding original reads, we identified putative new L1 insertion sites (Fig. 3E, F). Employing a minimum of 5 support reads as a cut-off for potential insertions, our pipeline predicted a total of 54 insertions that were present in our C57BL/6 mouse but absent from the C57BL/6 reference genome (Table S6). We manually curated the potential insertion sites in the Integrative Genomics Viewer (IGV) and called 43 out of 54 insertions as likely true insertions based on the poly(A) tail, number and quality of supporting reads, alignments, and putative endonuclease cleavage site. We observed a bias towards the endonuclease cleavage preference (TT/AAAA) at the putative insertion sites (Fig. 3G) (52). The majority of the false positive cases were caused by poor sequencing quality. Of the 43 non-reference L1s identified by nanotip-seq, only 6 were found when TIPseq was coupled with short reads (Fig. 4).

**Fig. 4.**
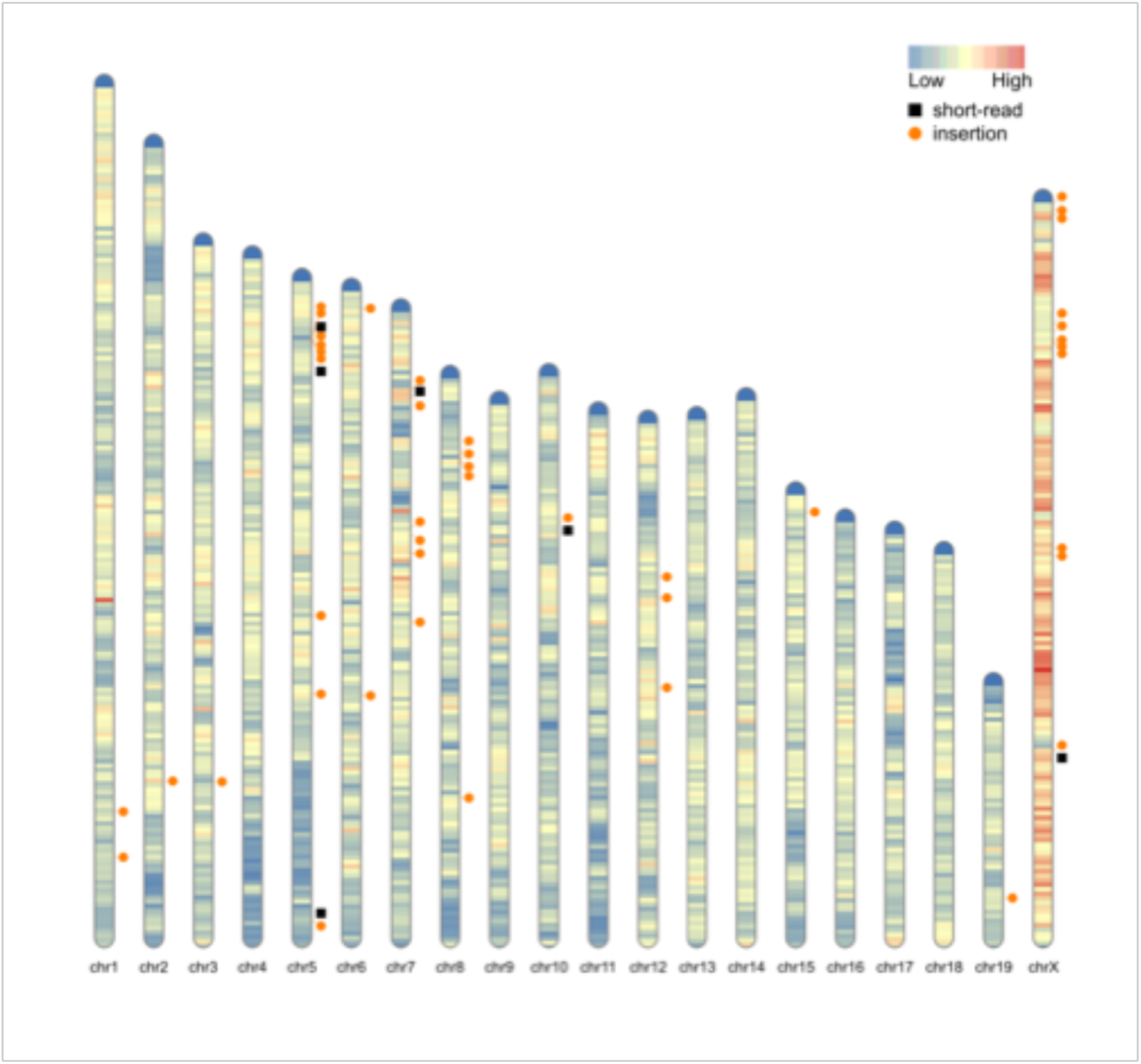
Genomic distribution of non-reference L1 insertions identified by long and short reads. Chromosome idiogram showing the identified non-reference L1s in our C57BL/6 mouse (orange circles). Black squares indicate insertions also found by short-read TIPseqhunter. Each chromosome is color-coded to show the density of repetitive elements.

In our initial experiments, we digested genomic DNA with restriction enzymes to adhere closely to the original TIPseq protocols published for human L1 (35, 39). To eliminate the need for restriction enzymes, we replaced enzymatic digestion with shearing, A-tailing, and ligation to a universal vectorette with a 3’ T-overhang. These steps drastically reduced sample processing time, and with 200,000 reads (115Mb), using sheared DNA method identified over 98.2% annotated L1 (“Sheared_B6_230217” from Table S5) and had 4x more coverage than enzymatic digested TIPseq (Fig. 3C) and identified 88% (288/326) of annotated L1s not found by short reads. Our pipeline also predicted 93 non-reference insertions (Table S7). We used manual examination of alignments at the locations of these predicted insertions on IGV to exclude 11 as false positives, leaving a total of 82 non-reference insertions identified. We picked 18 of these non-reference insertions and all were confirmed by genomic PCR or low coverage whole genome sequencing with ONT (Table S8). Notably, shearing also resulted in a continuous distribution of read lengths, as opposed to the discrete read lengths obtained when digesting the genomic DNA with restriction enzymes (Fig. S5). This lifted the concern that many of the reads were PCR duplicates. These results suggest that shearing has similar, if not higher, effectiveness compared to enzymatic digestion for our purpose.

### nanoTIPseq is capable of detecting lower frequency insertion events

Somatic mutations can lead to catastrophic consequences such as cancer and neurodevelopmental diseases (53, 54). While L1 insertions have traditionally been considered to occur primarily in the germline, it has been increasingly recognized that somatic L1 insertions can be detected in human and mouse genomes (6, 24–26, 55). To simulate the heterogeneity of somatic insertion events, we mixed FVB mouse gDNA with C57BL/6 gDNA at a 1:4 ratio. We used this mixture to prepare a library for nanoTIPseq. Additionally, we prepared a separate nanoTIPseq library using 100% FVB gDNA to serve as a reference for existing FVB L1 insertions. In the 100% FVB nanoTIPseq library, we discovered 1937 possible FVB strain related insertions that are absent from the C57BL/6 genome. In the 1:4 FVB:C57BL/6 nanoTIPseq library, we identified 1465 of the FVB specific L1s (76%) (Fig. S6A). Upon manual examination of our sequencing data for the 472 non-detected FVB specific L1s, we found that only 62/472 had no supporting reads in the 1:4 mixture data. Most had some alignment evidence supporting the presence of the non-detected elements in the library (Fig. S6B), suggesting the sensitivity of our procedure can be further improved, possibly by enhancing the bioinformatic pipeline or by increasing the sequencing depth. Nevertheless, these data demonstrated the ability for nanoTIPseq to detect insertion events present at less than one copy per genome.

### Single cell nanoTIPseq in mouse

Next, we asked how well mouse TIPseq performs with single cells. Short-read TIPseq has been reported to work with human single-cell samples, with limited detection sensitivity and a high false-positive rate (56). In this case, multi-displacement genome amplification (MDA) was used to amplify single cell genomes. Although MDA has advantages such as having high yield and low error rate, MDA also has disadvantages such as large drop-out rate, chimeric reads, and large variation between read length (57, 58). Primary template directed amplification (PTA) is a modified version of MDA, with incorporation of a reaction terminator (57). PTA decreases the formation of the “branches” seen in MDA, and ultimately leads to more uniform amplification of the target genome (57). We have tried both MDA and PTA whole genome amplification on single cells from 4226 cells, a breast cancer line derived from the MMTV-Wnt mouse model (46). As expected, MDA samples showed larger molecular weight amplification products, while PTA samples produced precise products around 1.5kb (Fig. S7). As a rough quality control measure, we designed primers to amplify an L1 adjacent region on each mouse chromosome. All PTA and MDA samples were positive for each chromosome (Table S9). We performed nanoTIPseq on 5 PTA amplified 4226 cells and 1 MDA amplified 4226 cell. From PTA amplified DNA, we identified on average 95.4% (∼3116/3266) of annotated C57BL/6 L1 insertions (Table S5), and 64/1937 (∼3%) L1 insertions from our reference of FVB L1s. Because the MMTV-Wnt mice were originally made in the FVB background, but subsequently backcrossed and maintained in the C57BL/6 background, these results are consistent with the backgrounds of the mice from which the 4226 cell line was derived. It is worth noting that the MDA amplified samples have a slightly higher mean CPM (215) compared to PTA amplified samples (192,196,199,187,220 for samples 1-5 respectively), however the overall number of annotated L1s recovered from MDA amplified genomes is less than the PTA group (88% vs 95.4%) at 200,000 reads. Furthermore, MDA amplified samples have higher variance than PTA samples where the L1 3’UTR coverage is consistent across PTA amplified single cell samples and resembles nanoTIPseq libraries made from unamplified bulk genomic DNA (Figure 5A), with an average coverage of ∼192 CPM. In addition, in all 5 single cell nanoTIPseq reactions, we found total of 130 non-reference L1s and 58 of them were not previously observed in either FVB or C57BL/6 background, indicating those insertion events occurred in a common progenitor cell or were polymorphic insertions in the original mice where the cancer originated (Figure 5B-D, Table S10) The consistency among the data from 5 individual single cells suggest that PTA coupled nanoTIPseq can be used for reliable identification of new L1 insertions from single cells.

**Fig. 5.**
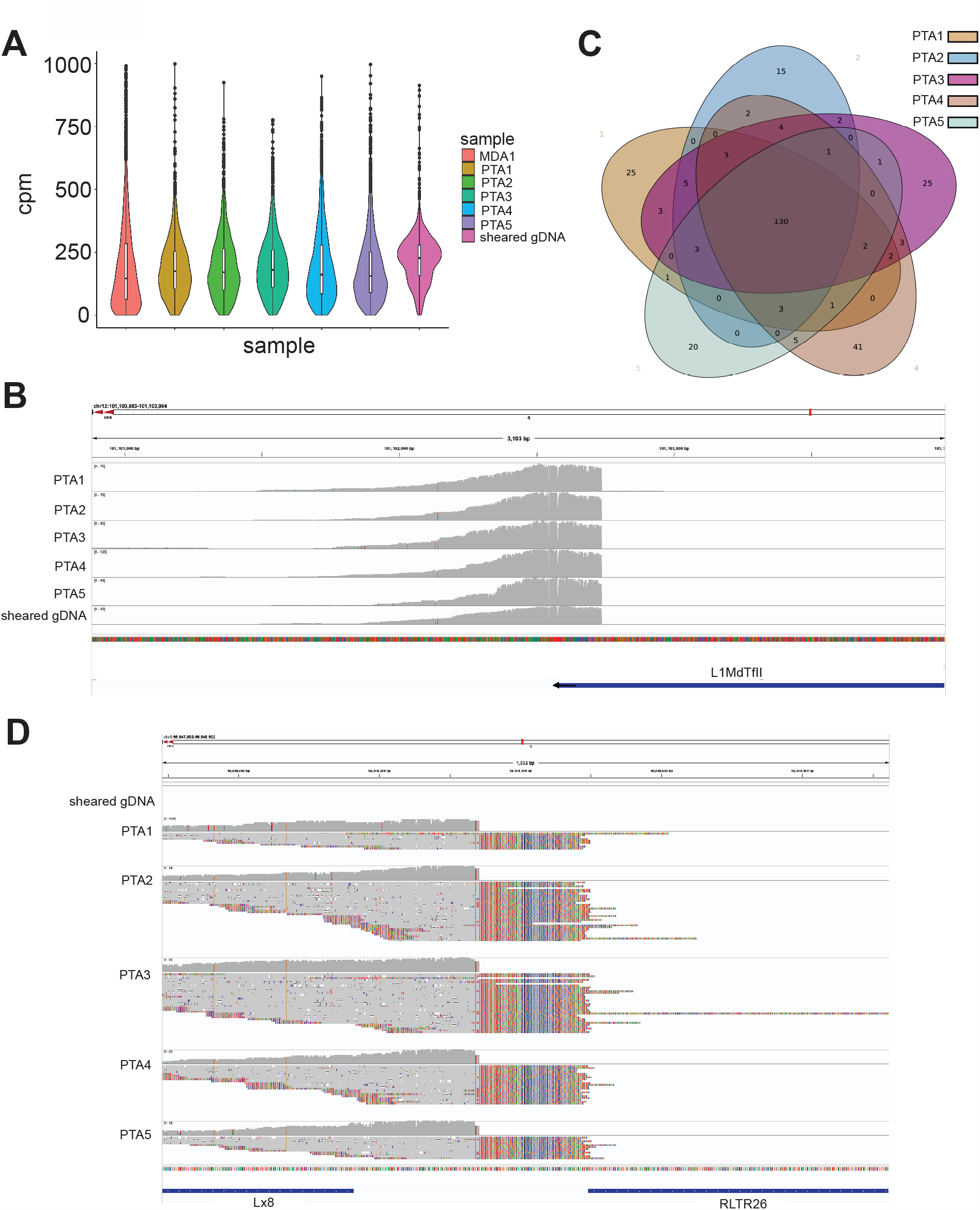
nanoTIPseq in 4226 single cell L1 detection. (**A**) PTA single-cell amplification showed consistent annotated L1 coverage among 5 different single cells. (**B**) Annotated L1 were readily detected in all 5 tested single cells nanoTIPseq samples. (**C**) Non-reference insertions from each single cell were intersected, and 130 common non-reference L1 insertions were identified in all 5 single cell samples, demonstrating good consistency and specificity in nanoTIPseq. (**D**) A representative example of a common non-reference L1 insertion found only in 4226 cells, but not C57BL/6 gDNA (“sheared gDNA”).

Aberrant expression of L1 has been correlated with many abnormal states in mammals (6, 7, 22–28). However, in most of these cases it is unclear whether L1 expression contributes to the underlying pathophysiology of disease. Mouse models are critical tools for studying human health and disease and will be valuable for evaluating the role of L1 in disease. One key aspect to characterizing the role of L1 is monitoring new L1 insertions. Whole genome sequencing and targeted L1 enrichment followed by sequencing have previously been used to identify human L1 insertions (33–36), but the sequencing depth required (ranging from 5 GB-120 GB) can create a cost limitation for smaller labs when a large number of samples or single cells need to be processed. WGS has also been used to identify mouse L1 insertions, but again the sequencing depth required poses an obstacle for many labs. nanoTIPseq allows us to identify >95% of mouse L1s from single cells or bulk tissue while requiring less than 150 MB of sequencing data per sample, a 30-fold reduction in sequencing data required per sample as compared to other methods. This opens the possibility of designing large studies with mouse disease models, then isolating tissues and/or single cells at various times during disease progression to pinpoint the cell type, timing, and frequency of L1 integration. Single-cell nanoTIPseq should also enable the unambiguous determination of the timing of L1 retrotransposition during normal development. Finally, because the human genome has far less active elements than the mouse genome (∼100 active humans L1s vs ∼3000 active mouse L1s) (3, 47), we would expect that using nanoTIPseq to profile active human L1s could require as little as 5 MB of sequencing data per sample, allowing low cost screening for retrotransposition events in the clinic (e.g. during prenatal testing).

