## Supplemental material for "Comprehensive profiling of L1 retrotransposons in mouse"

### SUPPLEMENTARY MATERIAL

### SUPPLEMENTARY FIGURES

**Fig. S1.** Base composition of L1 3′ UTR sequence in TIPseq amplicons.

TIPseq amplicons generated using TuJH922 or TuJH801 primers were mapped to the L1MdTf_1 consensus sequence (consensus sequence shown at the bottom). Grey indicates a match with the reference sequence. (**A**) The 4 TuJH922 samples had over 95% GTG at positions 6848-6850, while the 4 TuJH801 samples had predominantly ATA at position 6848-6850 (**B**) Detailed base composition at position 6848-6850 for TuJH801 samples 1-4 and TuJH922 samples 1-2. Coordinates 6848-6850 are shown on the x-axis, and the percentage of particular bases is shown on the y-axis.

**Fig. S2.** Representative examples of alignment evidence for detection of annotated L1s using short-read TIPseq data.

Shown are 4 examples viewed in IGV. L1 location and direction are shown on the bed file track. TuJH922 matched sequence is shown as the black box.

**Fig.S3.** Representative examples of TIPseq short-read false positives.

(**A**) An example of poor mapping of a 70bp region, where both end of the reads are being soft clipped. (**B**) An example of low-quality mapping of a non-repetitive region.

**Fig. S4.** Long-reads map better in repetitive regions compared to short-reads.

This IGV screenshot shows an example where an annotated L1 is immediately upstream of other repetitive elements (L1x_IV and L1MDA_VII). The short reads are not able to map effectively to the downstream region, causing the TIPseqhunter algorithm to not detect this L1 element. Nanopore reads map to this repetitive region, providing solid evidence for detection of this L1 loci. The black boxes highlight the coverage for long vs short reads. The reads are colored by read strand.

**Fig. S5.** Alignment characteristics of shearing vs enzymatic digestion methods.

A nanoTIPseq sample using sheard DNA is shown on the top of the IGV screenshot, with continuous coverage with various read lengths. In contrast, the bottom section of the IGV screenshot showcases samples prepared via enzymatic digestion where only two lengths of reads were observed.

**Fig. S6.** nanoTIPseq detection of low frequency insertion events. (**A**) Example of an FVB specific insertion identified in both the FVB only sample and FBV spike-in sample. Coverage of FVB only, spike-in, and C57BL/6 are shown on the top section. Read alignments are shown on the bottom section. Colored bases are being soft clipped. (**B**) Example of an FVB insertion not detected in spike-in samples by our algorithm, but present in the sequencing data. The alignment data demonstrates evidence of the insertion in the spike-in samples, but the low read coverage is under the detection limit of the nanoTIPseq pipeline.

**Fig. S7.** Distribution profile of whole genome amplification samples.

1 ng of each sample was loaded on an Agilent High Sensitivity DNA Assay chip.

**SUPPLEMENTARY TABLES**

**Table S1.** Primers used in this study

**Table S2.** PCR cycling parameters used for TIPseq in this study.

**Table S3.** Bookended elements that were separated in our L1 annotation file.

**Table S4.** Manually curated TIPseqHunter candidates from C57BL/6 gDNA.

**Table S5.** Coverage of annotated L1s identified by nanoTIPseq. Shown are number of annotated L1s (top) or percentage of annotated L1s (bottom) identified using 200,000 reads.

**Table S6.** Manually curated nanoTIPseq candidates from enzymatically digested C57BL/6 gDNA.

**Table S7.** Manually curated nanoTIPseq candidates from sheared C57BL/6 gDNA.

**Table S8.** Confirmation of 18 non-reference nanoTIPseq L1 insertions by genomic PCR or whole genome sequencing.

**Table S9.** Quality control of whole genome amplification by chromosome PCRs.

**Table S10.** Unique non-reference L1s found in 4226 cells.

Figure S1

**A**

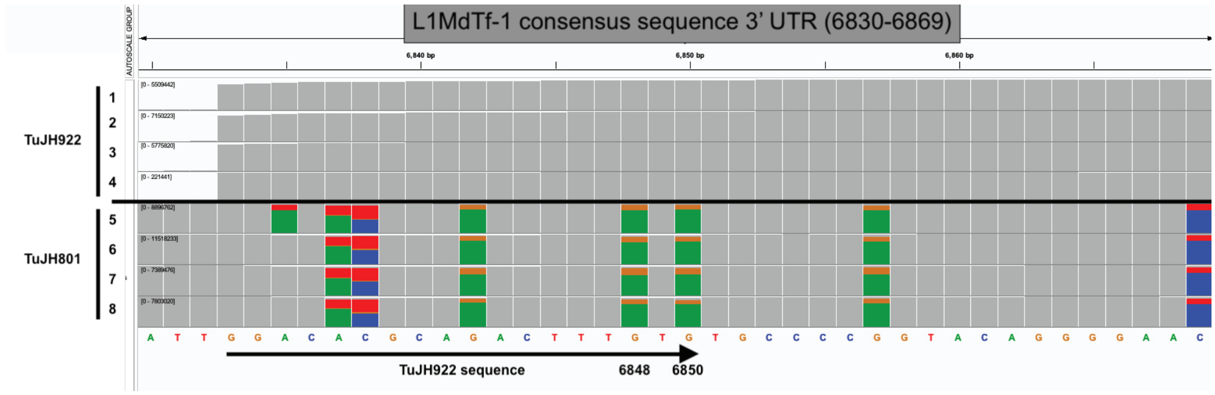
L1MdTf-1 consensus sequence 3' UTR (6830-6869)
AUTOSCALE GROUP
6,840 bp
6,850 bp
6,860 bp
TuJH922
TuJH801
1
2
3
4
5
6
7
8
A T T G G A C A C G C A G A C T T T G T G T G C C C C G G T A C A G G G G A A C
TuJH922 sequence
6848
6850

**B**

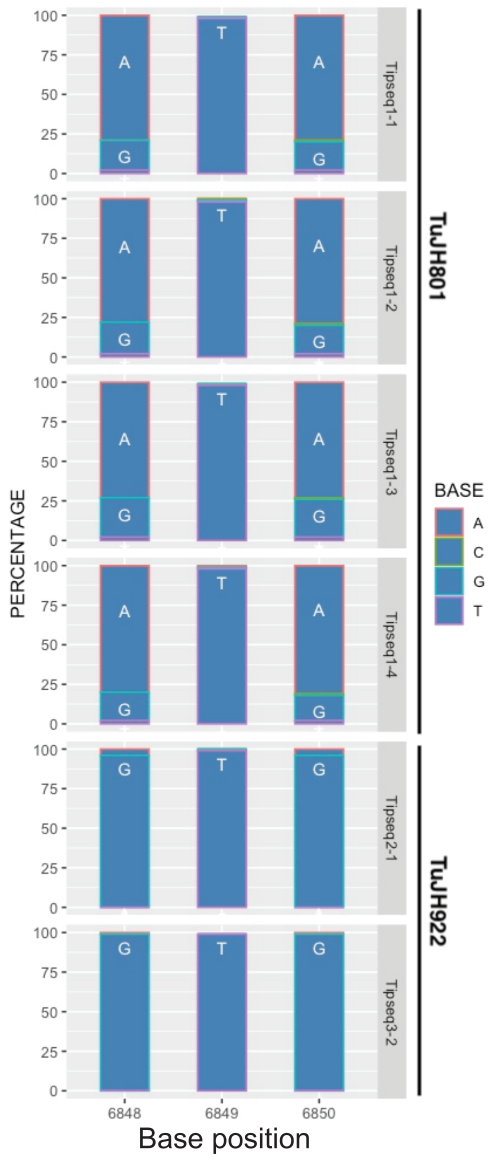
PERCENTAGE
100
75
50
25
0
A
G
T
Tipseq1-1
Tipseq1-2
Tipseq1-3
Tipseq1-4
Tipseq2-1
Tipseq3-2
TuJH801
TuJH922
BASE
A
C
G
T
6848
6849
6850
Base position

Figure S2

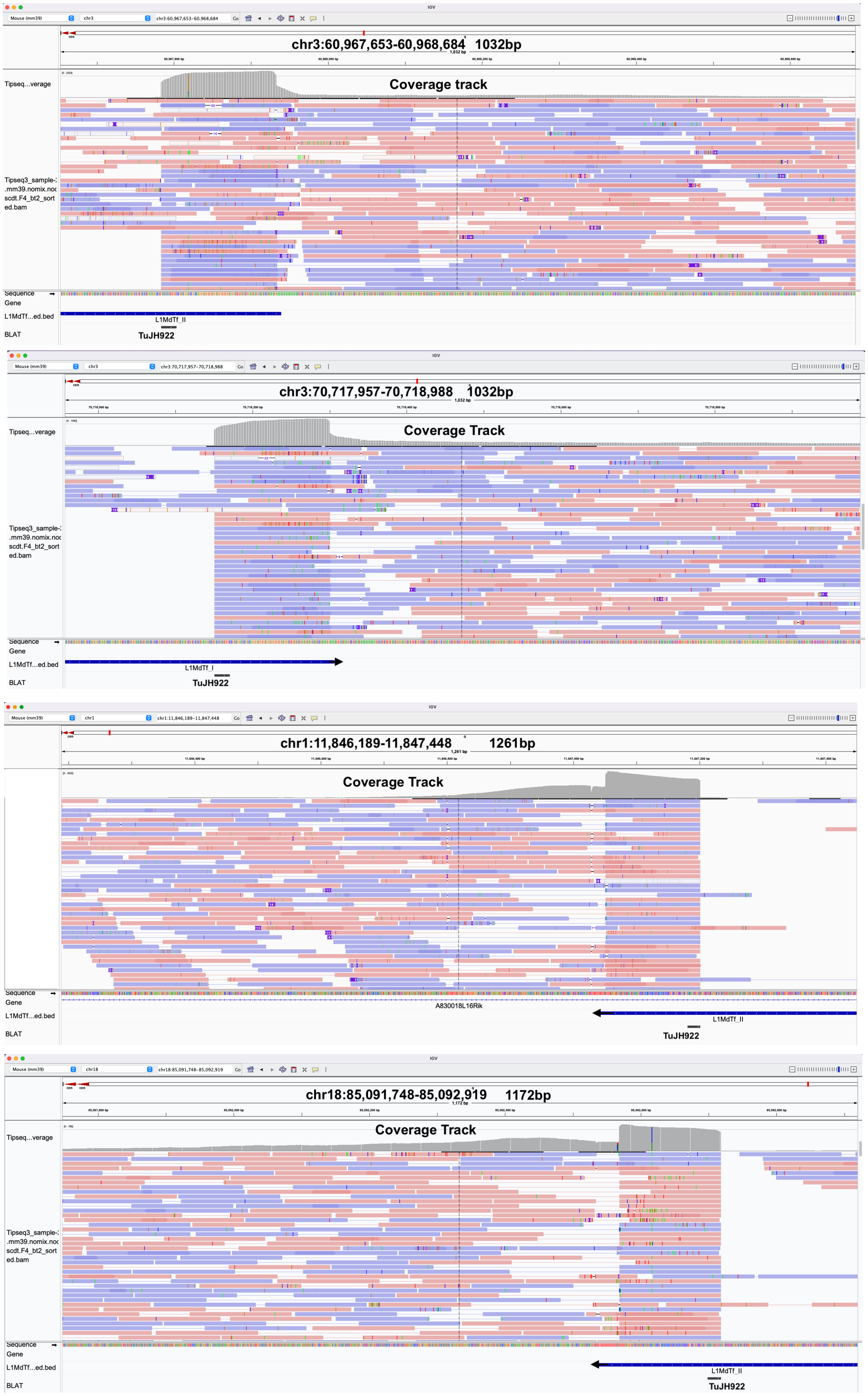
chr3:60,967,653-60,968,684
1032bp
Coverage track
Tipseq...verage
Tipseq3_sample-
.mm39.nomix.noc
scdt.F4_bt2_sort
ed.bam
Sequence
Gene
L1MdTf...ed.bed
L1MdTf_II
BLAT
TuJH922
chr3:70,717,957-70,718,988
1032bp
Coverage Track
Tipseq...verage
Tipseq3_sample-
.mm39.nomix.noc
scdt.F4_bt2_sort
ed.bam
Sequence
Gene
L1MdTf...ed.bed
L1MdTf_I
BLAT
TuJH922
chr1:11,846,189-11,847,448
1261bp
Coverage Track
Sequence
Gene
A830018L16Rik
L1MdTf...ed.bed
L1MdTf_II
BLAT
TuJH922
chr18:85,091,748-85,092,919
1172bp
Coverage Track
Tipseq...verage
Tipseq3_sample-
.mm39.nomix.noc
scdt.F4_bt2_sort
ed.bam
Sequence
Gene
L1MdTf...ed.bed
L1MdTf_II
BLAT
TuJH922

Figure S3

A

B

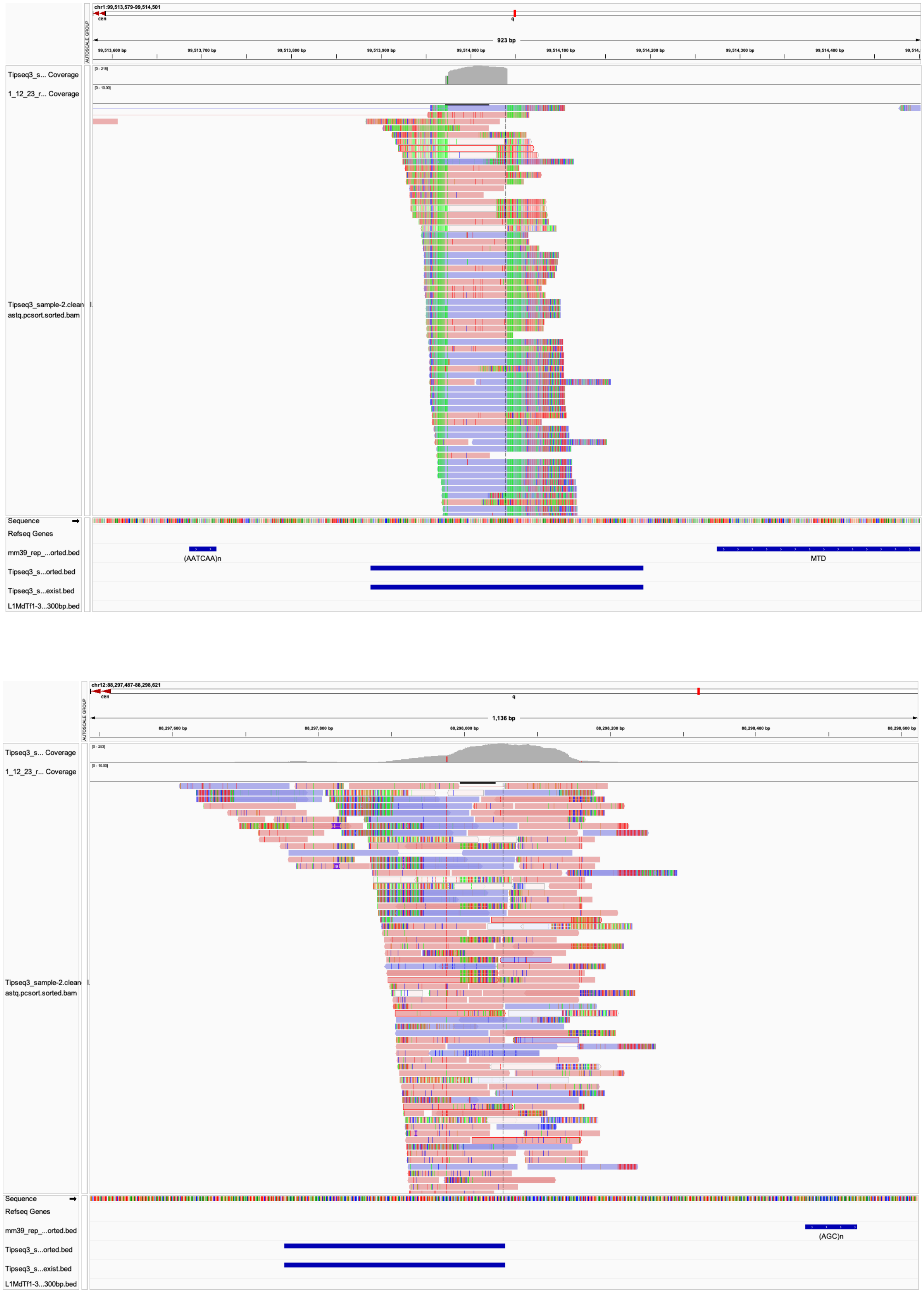
chr1:99,513,579-99,514,501
923 bp
Tipseq3_s... Coverage
1_12_23_r... Coverage
Tipseq3_sample-2.clean astq.pcsort.sorted.bam
Sequence
Refseq Genes
mm39_rep_...orted.bed
(AATCAA)n
MTD
Tipseq3_s...orted.bed
Tipseq3_s...exist.bed
L1MdTf1-3...300bp.bed
chr12:88,297,487-88,298,621
1,136 bp
Tipseq3_s... Coverage
1_12_23_r... Coverage
Tipseq3_sample-2.clean astq.pcsort.sorted.bam
Sequence
Refseq Genes
mm39_rep_...orted.bed
(AGC)n
Tipseq3_s...orted.bed
Tipseq3_s...exist.bed
L1MdTf1-3...300bp.bed

Figure S4

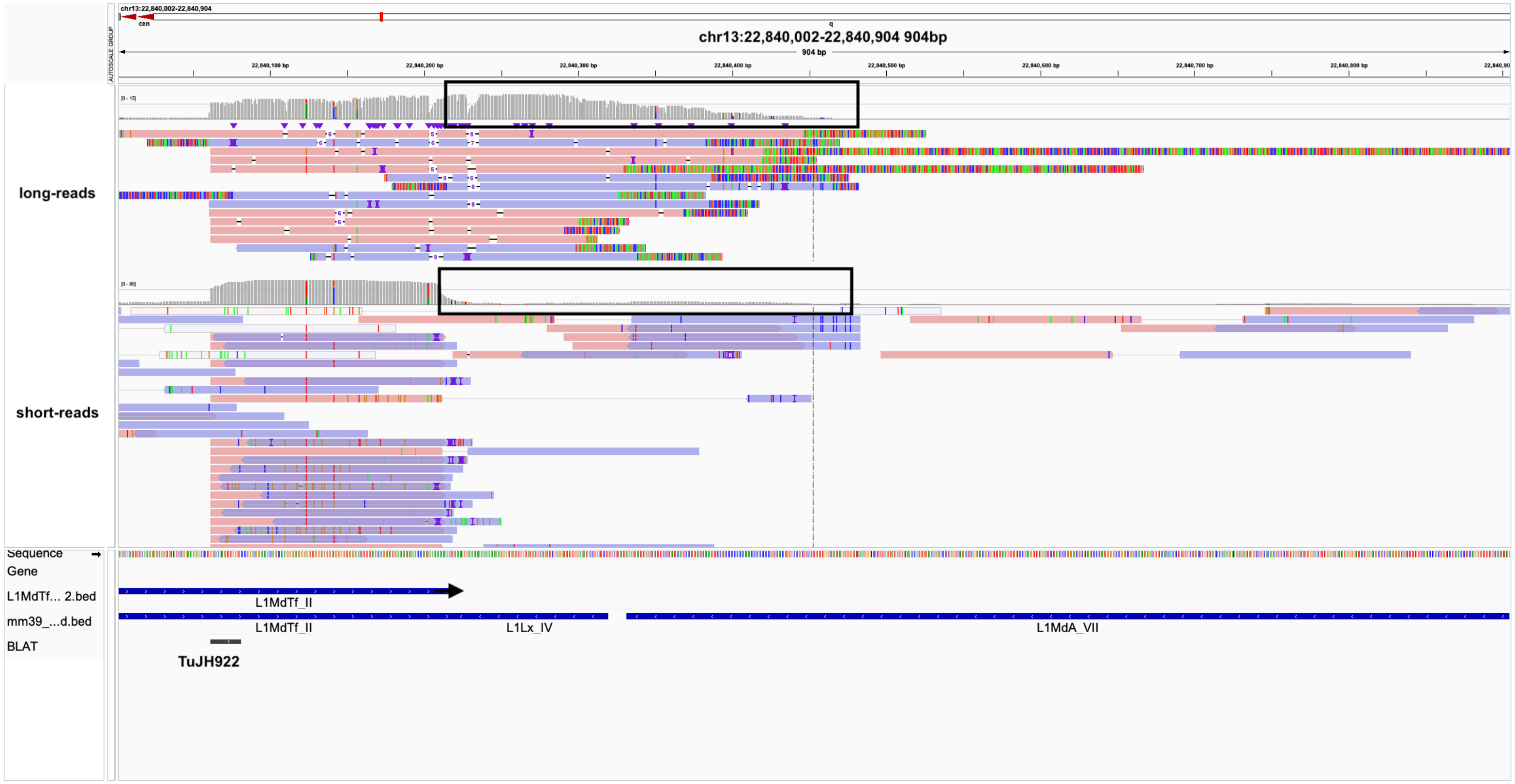

chr13:22,840,002-22,840,904 904bp
904 bp
22,840,100 bp
22,840,200 bp
22,840,300 bp
22,840,400 bp
22,840,500 bp
22,840,600 bp
22,840,700 bp
22,840,800 bp
long-reads
short-reads
Sequence
Gene
L1MdTf... 2.bed
mm39_...d.bed
BLAT
L1MdTf_II
L1MdTf_II
L1Lx_IV
L1MdA_VII
TuJH922

Figure S5

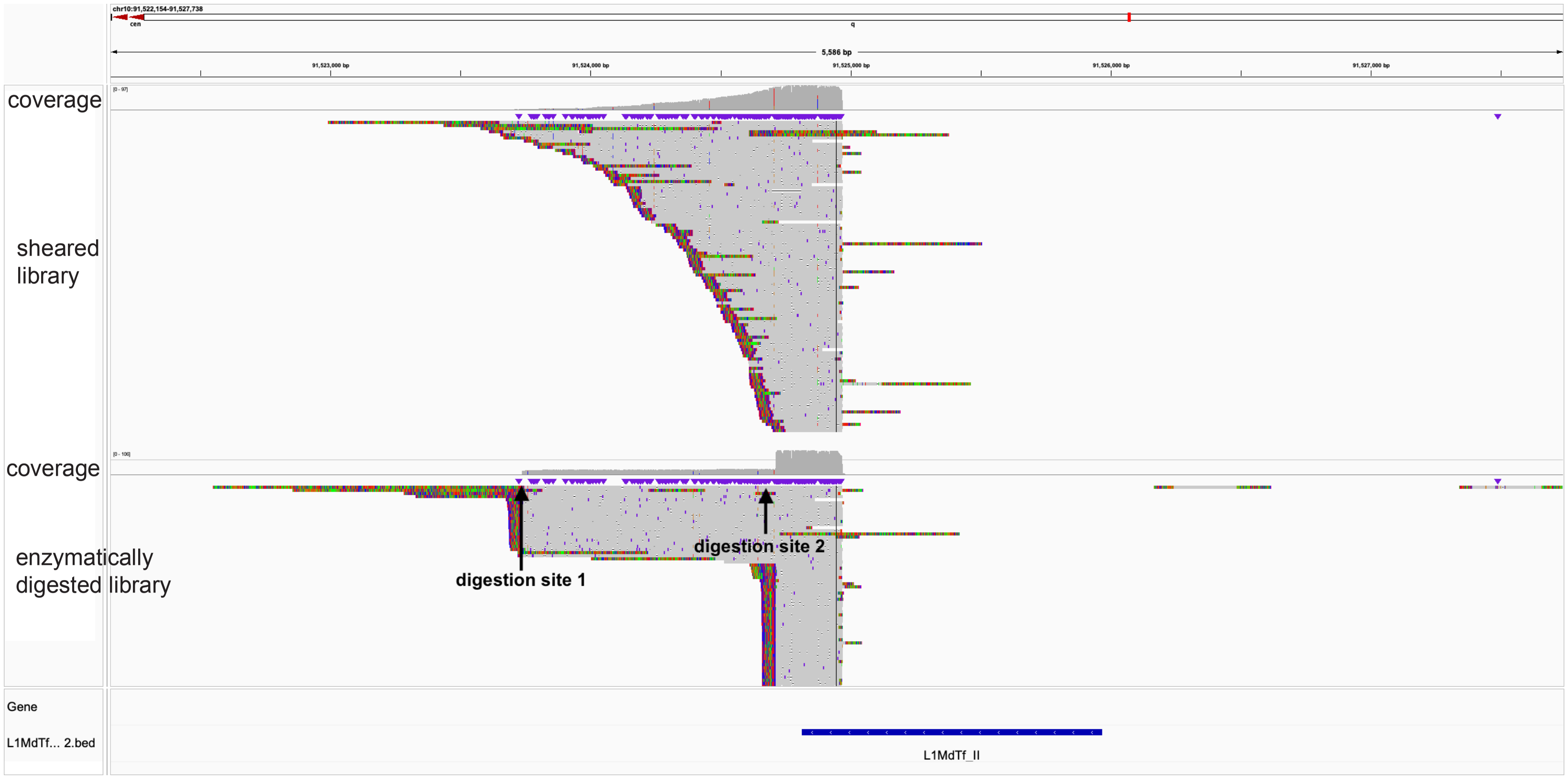
chr10:91,522,154-91,527,738
cen
q
5,586 bp
91,523,000 bp
91,524,000 bp
91,525,000 bp
91,526,000 bp
91,527,000 bp
coverage
sheared library
coverage
enzymatically digested library
digestion site 1
digestion site 2
Gene
L1MdTf... 2.bed
L1MdTf_II

Figure S6

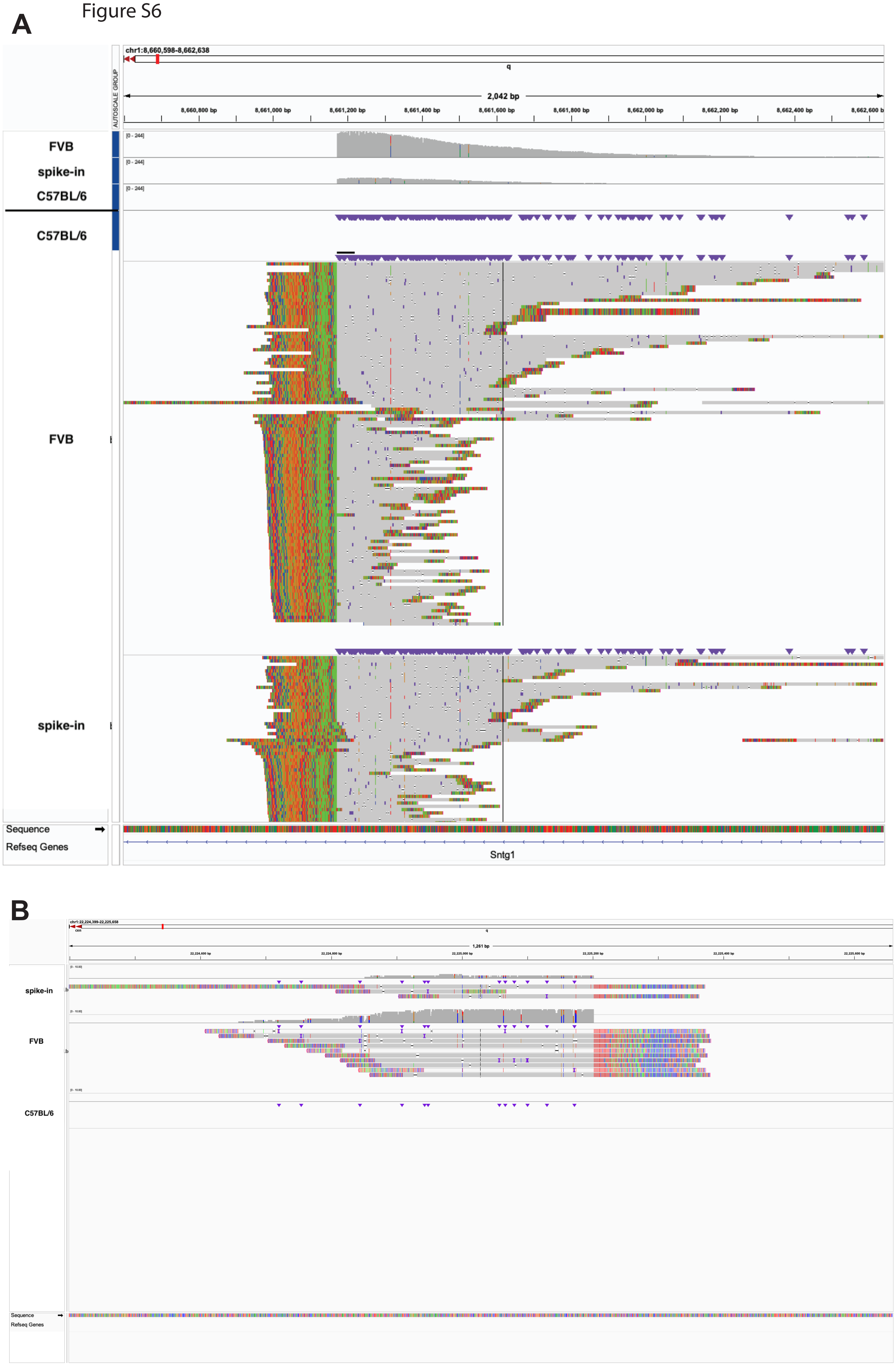
A
chr1:8,660,598-8,662,638
q
2,042 bp
8,660,800 bp
8,661,000 bp
8,661,200 bp
8,661,400 bp
8,661,600 bp
8,661,800 bp
8,662,000 bp
8,662,200 bp
8,662,400 bp
8,662,600 bp
AUTOSCALE GROUP
FVB
spike-in
C57BL/6
C57BL/6
FVB
spike-in
Sequence
Refseq Genes
Sntg1
B
spike-in
FVB
C57BL/6
Sequence
Refseq Genes

Figure S7

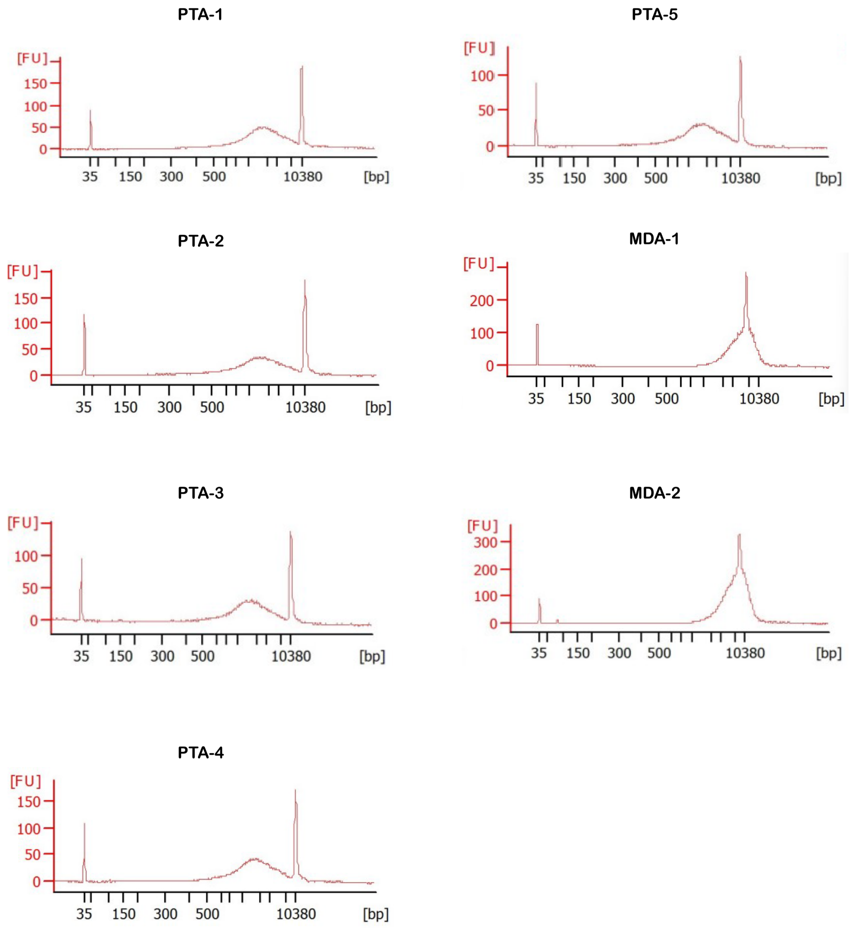

**Table S1. Primer list**

| Primer name | Sequence (5' to 3') | Description |
|---|---|---|
| TuJH800 | GAAGGAGAGGACGCTGTCTGTCGAAGGTAAGGAACGGACGAGAGAAGGGAGAG | common plus vectorette linker |
| TuJH801 | CTCTCCCTTCTCGGATCTTAA | Vectorette primer |
| TuJH802 | AACATTATGAACTAACCAGTACCCCT | L1 primer |
| TuJH922 | GGACACGCAGACTTTGTG | L1 primer |
| TuJH1030 | GGACACGCAGACTTTGTGTG | L1 primer |
| TuJH1031 | ACACGCAGACTTTGTGTG | L1 primer |
| TuJH1054 | CTCTCCCTTCTCGGATCTTAACCGTTCGTACGAGAATCGCTGTCCTCTCCTTCT | 3' T overhang common minus vectorette linker |
| TuJH1083 | AAGGGATAGTAGTGGTGGCC | validation primer for non-reference L1 chr 1:164797576 |
| TuJH1067 | GGGGAAGGGATAAAGCTGGA | validation primer for non-reference L1 chr 5:145538591 |
| TuJH1071 | ACCATGAGTCAGAAAGCAATGG | validation primer for non-reference L1 chr 7:51248689 |
| TuJH1076 | GTGCCTTATCAGAATCAGAGAGC | validation primer for non-reference L1 chr 7:56935951 |
| TuJH1143 | TGGGTGTTGGATCTTGTCAAA | validation primer for non-reference L1 chr 8:20487110 |
| TuJH1055 | GTCTGTGTGCTGCGTTTACA | validation primer for non-reference L1 chr 10:35898425 |
| TuJH1132 | AGAGAAAGTGGGGAAAAGCC | validation primer for non-reference L1 chr 12:41914804 |
| TuJH1060 | ATAAACCCGTTCCTCCCGAA | validation primer for non-reference L1 chr 12:37250143 |
| TuJH1087 | CTCTCGCCCATCCCAGAAAT | validation primer for non-reference L1 chr 12:62017979 |
| TuJH1063 | TGGCAGGGTGTAGAATCTCA | validation primer for non-reference L1 chr 19:50374143 |
| TuJH1095 | CGAAGCTTACTATCCAATGCCA | validation primer for non-reference L1 chr X:3312993 |
| TuJH1108 | CTTCCAATCACATGCCCACA | validation primer for non-reference L1 chr X:4442610 |
| TuJH1112 | CACTAGTTGCTGCACTCCAA | validation primer for non-reference L1 chr X:5091199 |
| TuJH1099 | CCATGCCCTTGTTCCAGTATC | validation primer for non-reference L1 chr X:31975242 |
| TuJH1103 | TGGTGTAATTCTGATAGGCTTGC | validation primer for non-reference L1 chr X:33311863 |
| TuJH1120 | TGCACCTCAGTACCTTGTTCT | validation primer for non-reference L1 chr X:33900422 |
| TuJH1079 | CTGCGGAACCATAATTGCTGA | validation primer for non-reference L1 chr X:81997828 |
| TuJH1091 | GGTCATCATTGTAAGATAGGGGA | validation primer for non-reference L1 chr X:125685683 |
| TuJH722 | GCCATTTAAGCAAGGCAACT | WGS coverage test chr 1 |
| TuJH1005 | AAGAGAAGGCCAACTGAAGGG | WGS coverage test chr 1 |
| TuJH927 | GGAATGCCCCACAACTCTGA | WGS coverage test chr 2 |
| TuJH928 | CACCTCTGAGGCCCTTGATC | WGS coverage test chr 2 |
| TuJH969 | GCAGTCAGGCTAGGGTCAAG | WGS coverage test chr 3 |
| TuJH970 | TTCAAAACCAGGGGCAGTGT | WGS coverage test chr 3 |
| TuJH971 | GTCGGCATGCATCCCAGATA | WGS coverage test chr 4 |
| TuJH972 | CTCCCAAGTCAGCACCTCTG | WGS coverage test chr 4 |
| TuJH973 | CATTGCAAGGCGTTCACACA | WGS coverage test chr 5 |
| TuJH974 | GGGCAGAAGTATGGGTGTCC | WGS coverage test chr 5 |
| TuJH935 | TGATCCACCTACTGGCCTCA | WGS coverage test chr 6 |
| TuJH936 | CAGGTGCCATCTTGAGAGCA | WGS coverage test chr 6 |
| TuJH937 | GTCACAGGACAAAGAGAGGCT | WGS coverage test chr 7 |
| TuJH938 | ATGGGGTGCCATACAGGGTA | WGS coverage test chr 7 |
| TuJH939 | TTGCCTTTGTGGTTGCCAAC | WGS coverage test chr 8 |
| TuJH940 | TGCCTGGTGTTAAAGCCCAA | WGS coverage test chr 8 |
| TuJH738 | CACTTCCCCTTCATTCCAGA | WGS coverage test chr 9 |
| TuJH1011 | TGGTGTCGGGAGTGTTCAAG | WGS coverage test chr 9 |
| TuJH983 | AAGTTCTGGCCCTTCCTGTC | WGS coverage test chr 10 |
| TuJH984 | TGCCATTGCCTTCATACCCT | WGS coverage test chr 10 |
| TuJH742 | CCTCCAAGCTCAGTCCAAAG | WGS coverage test chr 11 |
| TuJH1013 | TCCTCTCCTCAGGGCATTGA | WGS coverage test chr 11 |
| TuJH947 | ACTTCTTTGAACGTGAGCTGGA | WGS coverage test chr 12 |
| TuJH948 | ACTGTGGCATGGTCCCTTAC | WGS coverage test chr 12 |
| TuJH746 | TGCCTCAGTCTTCCGATCTT | WGS coverage test chr 13 |
| TuJH1015 | GGTACACTGGCCCTTGGAAC | WGS coverage test chr 13 |
| TuJH991 | CACTGCTGTAAATTCCGCGG | WGS coverage test chr 14 |
| TuJH992 | AAATGCAGAACACGCTGACC | WGS coverage test chr 14 |
| TuJH993 | TTGCCCCACAAGTTGTTCCT | WGS coverage test chr 15 |
| TuJH994 | GCTTTATAGCCAGGCCCACA | WGS coverage test chr 15 |
| TuJH995 | GGATTTGTGCTTGCTGGCAG | WGS coverage test chr 16 |
| TuJH996 | CTTCCCTGGAACCAGCATGT | WGS coverage test chr 16 |
| TuJH997 | CGGCAATTGCAGCTTGTCAT | WGS coverage test chr 17 |
| TuJH998 | TGTGCTGAGAACCAGAGCAG | WGS coverage test chr 17 |
| TuJH959 | ATGACTCCCACTGAAGCTGC | WGS coverage test chr 18 |
| TuJH960 | GAACTGTCTCGGACACTGCA | WGS coverage test chr 18 |
| TuJH758 | AGCCAATCTGGGTCCCTAGT | WGS coverage test chr 19 |
| TuJH1021 | GTCCCTGACCTGTGACCTTG | WGS coverage test chr 19 |
| TuJH963 | CCTGGCTTGCTGTCCTGTTT | WGS coverage test chr X |
| TuJH964 | TGGGTTTGCACACCAAGTCT | WGS coverage test chr X |

**Table S2. TIPseq PCR program**

| Temperature (˚C) | Duration (min) | Cycles |
|---|---|---|
| 95 | 3 | 1 |
| 95 | 1 | |
| 72 | 1 | 5 |
| 72 | 2 | |
| 95 | 1 | |
| 68 | 1 | 5 |
| 72 | 2 | |
| 95 | 45s | |
| 64 | 1 | 15 |
| 72 | 2 | |
| 95 | 45s | |
| 60 | 1 | 15 |
| 72 | 2 | |
| 72 | 5 | 1 |
| 4 | hold | |

**Table S3. Bookended L1s in reference genome**

| Chromosome | Start | End | Type | Strand | Length |
|---|---|---|---|---|---|
| chr1 | 4629820 | 4636224 | L1MdTf_I | + | 6404 |
| chr1 | 4636224 | 4637217 | L1MdTf_I | + | 993 |
| chr1 | 5259367 | 5264947 | L1MdTf_III | + | 5580 |
| chr1 | 5264947 | 5265940 | L1MdTf_III | + | 993 |
| chr1 | 5420616 | 5426477 | L1MdTf_I | + | 5861 |
| chr1 | 5426477 | 5427492 | L1MdTf_I | + | 1015 |
| chr1 | 6063319 | 6068585 | L1MdTf_III | + | 5266 |
| chr1 | 6068585 | 6069577 | L1MdTf_III | + | 992 |
| chr1 | 7260691 | 7266040 | L1MdTf_III | + | 5349 |
| chr1 | 7266040 | 7267033 | L1MdTf_III | + | 993 |
| chr1 | 7913663 | 7919046 | L1MdTf_III | + | 5383 |
| chr1 | 7919046 | 7920062 | L1MdTf_III | + | 1016 |
| chr1 | 9128380 | 9133778 | L1MdTf_III | + | 5398 |
| chr1 | 9133778 | 9134773 | L1MdTf_III | + | 995 |
| chr1 | 9310739 | 9316370 | L1MdTf_III | + | 5631 |
| chr1 | 9316370 | 9317388 | L1MdTf_III | + | 1018 |
| chr1 | 12246967 | 12247337 | L1MdTf_III | + | 370 |
| chr1 | 12247337 | 12248330 | L1MdTf_III | + | 993 |
| chr1 | 14183463 | 14189308 | L1MdTf_II | + | 5845 |
| chr1 | 14189308 | 14190301 | L1MdTf_II | + | 993 |
| chr1 | 15153731 | 15159604 | L1MdTf_III | + | 5873 |
| chr1 | 15159604 | 15160596 | L1MdTf_III | + | 992 |
| chr1 | 17537078 | 17542717 | L1MdTf_II | + | 5639 |
| chr1 | 17542718 | 17543710 | L1MdTf_II | + | 992 |
| chr1 | 18155466 | 18156777 | L1MdTf_III | + | 1311 |
| chr1 | 18156777 | 18157769 | L1MdTf_III | + | 992 |
| chr1 | 18519273 | 18521733 | L1MdTf_I | + | 2460 |
| chr1 | 18521733 | 18522746 | L1MdTf_I | + | 1013 |
| chr1 | 18542908 | 18548443 | L1MdTf_III | + | 5535 |
| chr1 | 18548443 | 18549435 | L1MdTf_III | + | 992 |
| chr1 | 18942648 | 18947882 | L1MdTf_II | + | 5234 |
| chr1 | 18947882 | 18948875 | L1MdTf_II | + | 993 |
| chr1 | 19442514 | 19447940 | L1MdTf_II | + | 5426 |
| chr1 | 19447940 | 19448933 | L1MdTf_II | + | 993 |
| chr1 | 19469014 | 19474254 | L1MdTf_II | + | 5240 |
| chr1 | 19474254 | 19475273 | L1MdTf_II | + | 1019 |
| chr1 | 20319346 | 20320399 | L1MdTf_I | + | 1053 |
| chr1 | 20320399 | 20321392 | L1MdTf_I | + | 993 |
| chr1 | 20502437 | 20508723 | L1MdTf_I | + | 6286 |
| chr1 | 20508723 | 20509716 | L1MdTf_I | + | 993 |
| chr1 | 22230488 | 22232692 | L1MdTf_I | + | 2204 |

| | | | | | |
|---|---|---|---|---|---|
| chr1 | 22232692 | 22233685 | L1MdTf_I | + | 993 |
| chr1 | 25158855 | 25164533 | L1MdTf_II | + | 5678 |
| chr1 | 25164533 | 25165526 | L1MdTf_II | + | 993 |
| chr1 | 25629914 | 25635341 | L1MdTf_I | + | 5427 |
| chr1 | 25635341 | 25636334 | L1MdTf_I | + | 993 |
| chr1 | 25701136 | 25707376 | L1MdTf_I | + | 6240 |
| chr1 | 25707376 | 25708369 | L1MdTf_I | + | 993 |
| chr1 | 26840422 | 26842084 | L1MdTf_I | + | 1662 |
| chr1 | 26842084 | 26843077 | L1MdTf_I | + | 993 |
| chr1 | 27178814 | 27183736 | L1MdTf_III | + | 4922 |
| chr1 | 27183736 | 27184758 | L1MdTf_III | + | 1022 |
| chr1 | 27198486 | 27204277 | L1MdTf_I | + | 5791 |
| chr1 | 27204277 | 27205289 | L1MdTf_I | + | 1012 |
| chr1 | 27794175 | 27799400 | L1MdTf_I | + | 5225 |
| chr1 | 27799400 | 27800393 | L1MdTf_I | + | 993 |
| chr1 | 27871124 | 27871482 | L1MdTf_III | - | 358 |
| chr1 | 27871482 | 27872036 | L1MdTf_III | + | 554 |
| chr1 | 28032373 | 28032418 | L1MdTf_III | + | 45 |
| chr1 | 28032418 | 28033411 | L1MdTf_III | + | 993 |
| chr1 | 28326665 | 28332930 | L1MdTf_II | + | 6265 |
| chr1 | 28332930 | 28333923 | L1MdTf_II | + | 993 |
| chr1 | 28706844 | 28707395 | L1MdTf_II | - | 551 |
| chr1 | 28707395 | 28708042 | L1MdTf_II | + | 647 |
| chr1 | 29645510 | 29646443 | L1MdTf_II | + | 933 |
| chr1 | 29646443 | 29647436 | L1MdTf_II | + | 993 |
| chr1 | 29742506 | 29747933 | L1MdTf_I | + | 5427 |
| chr1 | 29747933 | 29748926 | L1MdTf_I | + | 993 |
| chr1 | 30769424 | 30774945 | L1MdTf_III | + | 5521 |
| chr1 | 30774945 | 30775938 | L1MdTf_III | + | 993 |
| chr1 | 30892762 | 30898013 | L1MdTf_I | + | 5251 |
| chr1 | 30898013 | 30899012 | L1MdTf_I | + | 999 |
| chr1 | 31114201 | 31120049 | L1MdTf_I | + | 5848 |
| chr1 | 31120049 | 31121063 | L1MdTf_I | + | 1014 |
| chr1 | 32704745 | 32710146 | L1MdTf_II | + | 5401 |
| chr1 | 32710146 | 32711166 | L1MdTf_II | + | 1020 |
| chr1 | 32818410 | 32819619 | L1MdTf_I | + | 1209 |
| chr1 | 32819619 | 32820632 | L1MdTf_I | + | 1013 |
| chr1 | 33201616 | 33206966 | L1MdTf_III | + | 5350 |
| chr1 | 33206966 | 33207964 | L1MdTf_III | + | 998 |
| chr1 | 33446494 | 33452955 | L1MdTf_II | + | 6461 |
| chr1 | 33452955 | 33453948 | L1MdTf_II | + | 993 |
| chr1 | 41115989 | 41116266 | L1MdTf_III | + | 277 |
| chr1 | 41116267 | 41117258 | L1MdTf_III | + | 991 |
| chr1 | 42373829 | 42379193 | L1MdTf_I | + | 5364 |
| chr1 | 42379193 | 42380185 | L1MdTf_I | + | 992 |

| | | | | | |
|---|---|---|---|---|---|
| chr1 | 44515984 | 44519615 | L1MdTf_I | - | 3631 |
| chr1 | 44519615 | 44525908 | L1MdTf_I | + | 6293 |
| chr1 | 44942468 | 44947493 | L1MdTf_II | + | 5025 |
| chr1 | 44947493 | 44948511 | L1MdTf_II | + | 1018 |
| chr1 | 44998420 | 45000000 | L1MdTf_II | + | 1580 |
| chr1 | 45000000 | 45004695 | L1MdTf_I | + | 4695 |
| chr1 | 45693388 | 45698636 | L1MdTf_III | + | 5248 |
| chr1 | 45698636 | 45699629 | L1MdTf_III | + | 993 |
| chr1 | 46622521 | 46627753 | L1MdTf_II | + | 5232 |
| chr1 | 46627753 | 46628746 | L1MdTf_II | + | 993 |
| chr1 | 47144987 | 47150432 | L1MdTf_II | + | 5445 |
| chr1 | 47150432 | 47151416 | L1MdTf_II | + | 984 |
| chr1 | 47626850 | 47632084 | L1MdTf_I | + | 5234 |
| chr1 | 47632084 | 47633077 | L1MdTf_I | + | 993 |
| chr1 | 47654565 | 47654771 | L1MdTf_III | - | 206 |
| chr1 | 47654771 | 47655751 | L1MdTf_III | + | 980 |
| chr1 | 47724768 | 47728590 | L1MdTf_III | + | 3822 |
| chr1 | 47728591 | 47729583 | L1MdTf_III | + | 992 |
| chr1 | 48584707 | 48591403 | L1MdTf_I | + | 6696 |
| chr1 | 48591403 | 48592396 | L1MdTf_I | + | 993 |
| chr1 | 48724488 | 48725044 | L1MdTf_III | + | 556 |
| chr1 | 48725044 | 48726030 | L1MdTf_III | + | 986 |
| chr1 | 48854140 | 48859588 | L1MdTf_III | + | 5448 |
| chr1 | 48859588 | 48860580 | L1MdTf_III | + | 992 |
| chr1 | 50369130 | 50374557 | L1MdTf_II | + | 5427 |
| chr1 | 50374557 | 50375550 | L1MdTf_II | + | 993 |
| chr1 | 50499700 | 50500000 | L1MdTf_I | + | 300 |
| chr1 | 50500000 | 50500090 | L1MdTf_II | + | 90 |
| chr1 | 52439079 | 52445036 | L1MdTf_II | + | 5957 |
| chr1 | 52445036 | 52446032 | L1MdTf_II | + | 996 |
| chr1 | 52978668 | 52984555 | L1MdTf_I | + | 5887 |
| chr1 | 52984555 | 52985548 | L1MdTf_I | + | 993 |
| chr1 | 54065363 | 54070578 | L1MdTf_II | + | 5215 |
| chr1 | 54070578 | 54071571 | L1MdTf_II | + | 993 |
| chr1 | 54426337 | 54431866 | L1MdTf_I | + | 5529 |
| chr1 | 54431866 | 54432859 | L1MdTf_I | + | 993 |
| chr1 | 56419491 | 56424919 | L1MdTf_I | + | 5428 |
| chr1 | 56424919 | 56425912 | L1MdTf_I | + | 993 |
| chr1 | 56497793 | 56500000 | L1MdTf_II | - | 2207 |
| chr1 | 56500000 | 56504004 | L1MdTf_II | - | 4004 |
| chr1 | 61553400 | 61559515 | L1MdTf_I | + | 6115 |
| chr1 | 61559515 | 61560508 | L1MdTf_I | + | 993 |
| chr1 | 61931930 | 61933638 | L1MdTf_III | + | 1708 |
| chr1 | 61933638 | 61934631 | L1MdTf_III | + | 993 |
| chr1 | 62334554 | 62340471 | L1MdTf_I | + | 5917 |

| | | | | | |
|---|---|---|---|---|---|
| chr1 | 62340471 | 62341464 | L1MdTf_I | + | 993 |
| chr1 | 65546710 | 65546981 | L1MdTf_III | - | 271 |
| chr1 | 65546981 | 65547205 | L1MdTf_III | + | 224 |
| chr1 | 67636580 | 67637093 | L1MdTf_III | - | 513 |
| chr1 | 67637093 | 67637750 | L1MdTf_III | + | 657 |
| chr1 | 68637415 | 68643025 | L1MdTf_III | + | 5610 |
| chr1 | 68643025 | 68644018 | L1MdTf_III | + | 993 |
| chr1 | 68664071 | 68669669 | L1MdTf_III | + | 5598 |
| chr1 | 68669669 | 68670662 | L1MdTf_III | + | 993 |
| chr1 | 69270590 | 69276681 | L1MdTf_I | + | 6091 |
| chr1 | 69276681 | 69277674 | L1MdTf_I | + | 993 |
| chr1 | 70247873 | 70253279 | L1MdTf_II | + | 5406 |
| chr1 | 70253279 | 70254272 | L1MdTf_II | + | 993 |
| chr1 | 70726300 | 70728876 | L1MdTf_I | + | 2576 |
| chr1 | 70728876 | 70729869 | L1MdTf_I | + | 993 |
| chr1 | 71289718 | 71294928 | L1MdTf_II | + | 5210 |
| chr1 | 71294928 | 71295921 | L1MdTf_II | + | 993 |
| chr1 | 71571212 | 71574231 | L1MdTf_II | + | 3019 |
| chr1 | 71574231 | 71575227 | L1MdTf_II | + | 996 |
| chr1 | 72717480 | 72718120 | L1MdTf_III | - | 640 |
| chr1 | 72718120 | 72718469 | L1MdTf_III | + | 349 |
| chr1 | 73506052 | 73508386 | L1MdTf_I | + | 2334 |
| chr1 | 73508386 | 73509398 | L1MdTf_I | + | 1012 |
| chr1 | 76331998 | 76333348 | L1MdTf_III | + | 1350 |
| chr1 | 76333348 | 76334341 | L1MdTf_III | + | 993 |
| chr1 | 76436631 | 76442062 | L1MdTf_I | + | 5431 |
| chr1 | 76442062 | 76443055 | L1MdTf_I | + | 993 |
| chr1 | 76716259 | 76718426 | L1MdTf_I | - | 2167 |
| chr1 | 76718426 | 76719408 | L1MdTf_I | + | 982 |
| chr1 | 79358552 | 79364031 | L1MdTf_I | + | 5479 |
| chr1 | 79364031 | 79365024 | L1MdTf_I | + | 993 |
| chr1 | 80113289 | 80117117 | L1MdTf_I | + | 3828 |
| chr1 | 80117117 | 80118109 | L1MdTf_I | + | 992 |
| chr1 | 80961496 | 80967105 | L1MdTf_II | + | 5609 |
| chr1 | 80967105 | 80968098 | L1MdTf_II | + | 993 |
| chr1 | 81397383 | 81403386 | L1MdTf_II | + | 6003 |
| chr1 | 81403386 | 81404377 | L1MdTf_II | + | 991 |
| chr1 | 81476190 | 81481612 | L1MdTf_I | + | 5422 |
| chr1 | 81481612 | 81482605 | L1MdTf_I | + | 993 |
| chr1 | 83116749 | 83117418 | L1MdTf_II | - | 669 |
| chr1 | 83117418 | 83117723 | L1MdTf_II | + | 305 |
| chr1 | 83797333 | 83802548 | L1MdTf_II | + | 5215 |
| chr1 | 83802548 | 83803567 | L1MdTf_II | + | 1019 |
| chr1 | 84549463 | 84553194 | L1MdTf_II | + | 3731 |
| chr1 | 84553194 | 84554187 | L1MdTf_II | + | 993 |

| | | | | | |
|---|---|---|---|---|---|
| chr1 | 85035692 | 85037866 | L1MdTf_III | + | 2174 |
| chr1 | 85037866 | 85038859 | L1MdTf_III | + | 993 |
| chr1 | 86914407 | 86915454 | L1MdTf_III | - | 1047 |
| chr1 | 86915454 | 86916054 | L1MdTf_III | + | 600 |
| chr1 | 89686360 | 89686674 | L1MdTf_II | - | 314 |
| chr1 | 89686674 | 89687397 | L1MdTf_II | + | 723 |
| chr1 | 94028654 | 94034102 | L1MdTf_II | + | 5448 |
| chr1 | 94034102 | 94035098 | L1MdTf_II | + | 996 |
| chr1 | 94078569 | 94084226 | L1MdTf_II | + | 5657 |
| chr1 | 94084226 | 94085221 | L1MdTf_II | + | 995 |
| chr1 | 95153203 | 95156390 | L1MdTf_III | + | 3187 |
| chr1 | 95156390 | 95157381 | L1MdTf_III | + | 991 |
| chr1 | 95827105 | 95827322 | L1MdTf_I | - | 217 |
| chr1 | 95827322 | 95828018 | L1MdTf_II | + | 696 |
| chr1 | 96168541 | 96173856 | L1MdTf_III | + | 5315 |
| chr1 | 96173856 | 96174849 | L1MdTf_III | + | 993 |
| chr1 | 96668730 | 96672244 | L1MdTf_III | + | 3514 |
| chr1 | 96672244 | 96673237 | L1MdTf_III | + | 993 |
| chr1 | 97793420 | 97798800 | L1MdTf_III | + | 5380 |
| chr1 | 97798800 | 97799793 | L1MdTf_III | + | 993 |
| chr1 | 98128859 | 98134250 | L1MdTf_III | + | 5391 |
| chr1 | 98134250 | 98135241 | L1MdTf_III | + | 991 |
| chr1 | 98294164 | 98299803 | L1MdTf_I | + | 5639 |
| chr1 | 98299803 | 98300796 | L1MdTf_I | + | 993 |
| chr1 | 98312883 | 98318615 | L1MdTf_III | + | 5732 |
| chr1 | 98318615 | 98319608 | L1MdTf_III | + | 993 |
| chr1 | 98801052 | 98802056 | L1MdTf_III | + | 1004 |
| chr1 | 98802056 | 98803049 | L1MdTf_III | + | 993 |
| chr1 | 100063123 | 100063466 | L1MdTf_III | - | 343 |
| chr1 | 100063466 | 100063699 | L1MdTf_III | + | 233 |
| chr1 | 101172822 | 101174904 | L1MdTf_I | + | 2082 |
| chr1 | 101174904 | 101175897 | L1MdTf_I | + | 993 |
| chr1 | 101230143 | 101235585 | L1MdTf_II | + | 5442 |
| chr1 | 101235585 | 101236581 | L1MdTf_II | + | 996 |
| chr1 | 101332631 | 101333186 | L1MdTf_III | + | 555 |
| chr1 | 101333186 | 101334177 | L1MdTf_III | + | 991 |
| chr1 | 102928344 | 102933826 | L1MdTf_III | + | 5482 |
| chr1 | 102933826 | 102934820 | L1MdTf_III | + | 994 |
| chr1 | 104145389 | 104150816 | L1MdTf_I | + | 5427 |
| chr1 | 104150816 | 104151809 | L1MdTf_I | + | 993 |
| chr1 | 105495702 | 105500000 | L1MdTf_III | + | 4298 |
| chr1 | 105500000 | 105501131 | L1MdTf_III | + | 1131 |
| chr1 | 107555990 | 107559332 | L1MdTf_II | + | 3342 |
| chr1 | 107559333 | 107560326 | L1MdTf_II | + | 993 |
| chr1 | 107750516 | 107756647 | L1MdTf_I | + | 6131 |

| | | | | | |
|---|---|---|---|---|---|
| chr1 | 107756647 | 107757640 | L1MdTf_I | + | 993 |
| chr1 | 108332786 | 108333100 | L1MdTf_II | - | 314 |
| chr1 | 108333100 | 108336584 | L1MdTf_II | - | 3484 |
| chr1 | 108962378 | 108968236 | L1MdTf_I | + | 5858 |
| chr1 | 108968236 | 108969229 | L1MdTf_I | + | 993 |
| chr1 | 109799269 | 109804889 | L1MdTf_II | + | 5620 |
| chr1 | 109804889 | 109805885 | L1MdTf_II | + | 996 |
| chr1 | 112765840 | 112771758 | L1MdTf_II | + | 5918 |
| chr1 | 112771758 | 112772753 | L1MdTf_II | + | 995 |
| chr1 | 113823245 | 113824779 | L1MdTf_II | + | 1534 |
| chr1 | 113824779 | 113825772 | L1MdTf_II | + | 993 |
| chr1 | 114410422 | 114410838 | L1MdTf_II | - | 416 |
| chr1 | 114410838 | 114410936 | L1MdTf_I | + | 98 |
| chr1 | 114485034 | 114491146 | L1MdTf_I | + | 6112 |
| chr1 | 114491146 | 114492158 | L1MdTf_I | + | 1012 |
| chr1 | 115374968 | 115380819 | L1MdTf_II | + | 5851 |
| chr1 | 115380819 | 115381812 | L1MdTf_II | + | 993 |
| chr1 | 115727163 | 115732300 | L1MdTf_II | + | 5137 |
| chr1 | 115732300 | 115733037 | L1MdTf_II | + | 737 |
| chr1 | 115733037 | 115733335 | L1MdTf_II | + | 298 |
| chr1 | 115733336 | 115734328 | L1MdTf_II | + | 992 |
| chr1 | 117024205 | 117024397 | L1MdTf_III | + | 192 |
| chr1 | 117024397 | 117025390 | L1MdTf_III | + | 993 |
| chr1 | 117785441 | 117790398 | L1MdTf_III | + | 4957 |
| chr1 | 117790398 | 117791390 | L1MdTf_III | + | 992 |
| chr1 | 119672049 | 119677536 | L1MdTf_II | + | 5487 |
| chr1 | 119677536 | 119678529 | L1MdTf_II | + | 993 |
| chr1 | 120921718 | 120927173 | L1MdTf_I | + | 5455 |
| chr1 | 120927173 | 120928158 | L1MdTf_I | + | 985 |
| chr1 | 120963383 | 120968653 | L1MdTf_II | + | 5270 |
| chr1 | 120968653 | 120969646 | L1MdTf_II | + | 993 |
| chr1 | 121665612 | 121670960 | L1MdTf_III | + | 5348 |
| chr1 | 121670960 | 121671953 | L1MdTf_III | + | 993 |
| chr1 | 122014148 | 122019793 | L1MdTf_II | + | 5645 |
| chr1 | 122019793 | 122020786 | L1MdTf_II | + | 993 |
| chr1 | 122801337 | 122801949 | L1MdTf_I | + | 612 |
| chr1 | 122801949 | 122802942 | L1MdTf_I | + | 993 |
| chr1 | 123442128 | 123443414 | L1MdTf_III | + | 1286 |
| chr1 | 123443414 | 123444407 | L1MdTf_III | + | 993 |
| chr1 | 123516889 | 123517439 | L1MdTf_II | + | 550 |
| chr1 | 123517439 | 123518429 | L1MdTf_II | + | 990 |
| chr1 | 123784770 | 123790206 | L1MdTf_I | + | 5436 |
| chr1 | 123790206 | 123791199 | L1MdTf_I | + | 993 |
| chr1 | 124069080 | 124074482 | L1MdTf_I | + | 5402 |
| chr1 | 124074482 | 124075475 | L1MdTf_I | + | 993 |

| | | | | | |
|---|---|---|---|---|---|
| chr1 | 124404275 | 124404459 | L1MdTf_III | + | 184 |
| chr1 | 124404460 | 124405449 | L1MdTf_III | + | 989 |
| chr1 | 125062446 | 125067906 | L1MdTf_II | + | 5460 |
| chr1 | 125067906 | 125068890 | L1MdTf_II | + | 984 |
| chr1 | 125068935 | 125071572 | L1MdTf_I | + | 2637 |
| chr1 | 125071572 | 125072558 | L1MdTf_I | + | 986 |
| chr1 | 125438572 | 125440331 | L1MdTf_II | + | 1759 |
| chr1 | 125440331 | 125441324 | L1MdTf_II | + | 993 |
| chr1 | 126719942 | 126725075 | L1MdTf_I | + | 5133 |
| chr1 | 126725075 | 126726065 | L1MdTf_I | + | 990 |
| chr1 | 127772518 | 127772928 | L1MdTf_II | - | 410 |
| chr1 | 127772928 | 127773040 | L1MdTf_II | + | 112 |
| chr1 | 128700937 | 128706425 | L1MdTf_III | + | 5488 |
| chr1 | 128706425 | 128707437 | L1MdTf_III | + | 1012 |
| chr1 | 130418829 | 130424456 | L1MdTf_II | + | 5627 |
| chr1 | 130424456 | 130425449 | L1MdTf_II | + | 993 |
| chr1 | 131256516 | 131256741 | L1MdTf_II | - | 225 |
| chr1 | 131256741 | 131256952 | L1MdTf_II | + | 211 |
| chr1 | 131608280 | 131613732 | L1MdTf_II | + | 5452 |
| chr1 | 131613732 | 131614725 | L1MdTf_II | + | 993 |
| chr1 | 137265162 | 137266578 | L1MdTf_III | - | 1416 |
| chr1 | 137266578 | 137273241 | L1MdTf_I | - | 6663 |
| chr1 | 138169321 | 138170613 | L1MdTf_II | + | 1292 |
| chr1 | 138170613 | 138171606 | L1MdTf_II | + | 993 |
| chr1 | 138721332 | 138722397 | L1MdTf_II | + | 1065 |
| chr1 | 138722397 | 138723389 | L1MdTf_II | + | 992 |
| chr1 | 139457460 | 139457549 | L1MdTf_I | - | 89 |
| chr1 | 139457549 | 139457757 | L1MdTf_II | + | 208 |
| chr1 | 139545717 | 139547600 | L1MdTf_II | + | 1883 |
| chr1 | 139547601 | 139548595 | L1MdTf_II | + | 994 |
| chr1 | 139644069 | 139649938 | L1MdTf_II | + | 5869 |
| chr1 | 139649938 | 139650951 | L1MdTf_II | + | 1013 |
| chr1 | 139835072 | 139840742 | L1MdTf_III | + | 5670 |
| chr1 | 139840742 | 139841735 | L1MdTf_III | + | 993 |
| chr1 | 140037013 | 140038462 | L1MdTf_I | + | 1449 |
| chr1 | 140038462 | 140039455 | L1MdTf_I | + | 993 |
| chr1 | 140442377 | 140445058 | L1MdTf_II | + | 2681 |
| chr1 | 140445058 | 140446054 | L1MdTf_II | + | 996 |
| chr1 | 140851631 | 140857148 | L1MdTf_I | + | 5517 |
| chr1 | 140857148 | 140858141 | L1MdTf_I | + | 993 |
| chr1 | 141117432 | 141123485 | L1MdTf_II | + | 6053 |
| chr1 | 141123485 | 141124479 | L1MdTf_II | + | 994 |
| chr1 | 141175832 | 141176645 | L1MdTf_III | + | 813 |
| chr1 | 141176646 | 141177641 | L1MdTf_III | + | 995 |
| chr1 | 141371854 | 141372621 | L1MdTf_I | - | 767 |

| | | | | | |
|---|---|---|---|---|---|
| chr1 | 141372621 | 141372811 | L1MdTf_I | + | 190 |
| chr1 | 141625788 | 141631102 | L1MdTf_II | - | 5314 |
| chr1 | 141631102 | 141631983 | L1MdTf_II | + | 881 |
| chr1 | 142599494 | 142600148 | L1MdTf_III | + | 654 |
| chr1 | 142600148 | 142601107 | L1MdTf_III | + | 959 |
| chr1 | 142964861 | 142965635 | L1MdTf_I | - | 774 |
| chr1 | 142965635 | 142966122 | L1MdTf_I | + | 487 |
| chr1 | 143175222 | 143175286 | L1MdTf_III | + | 64 |
| chr1 | 143175286 | 143176306 | L1MdTf_III | + | 1020 |
| chr1 | 143737833 | 143743481 | L1MdTf_II | + | 5648 |
| chr1 | 143743481 | 143744474 | L1MdTf_II | + | 993 |
| chr1 | 143966854 | 143967590 | L1MdTf_III | + | 736 |
| chr1 | 143967590 | 143968583 | L1MdTf_III | + | 993 |
| chr1 | 144196646 | 144202130 | L1MdTf_III | + | 5484 |
| chr1 | 144202130 | 144203098 | L1MdTf_III | + | 968 |
| chr1 | 144757570 | 144757674 | L1MdTf_III | + | 104 |
| chr1 | 144757674 | 144758665 | L1MdTf_III | + | 991 |
| chr1 | 145968914 | 145969147 | L1MdTf_III | + | 233 |
| chr1 | 145969147 | 145970141 | L1MdTf_III | + | 994 |
| chr1 | 146067720 | 146068402 | L1MdTf_I | - | 682 |
| chr1 | 146068402 | 146069529 | L1MdTf_I | + | 1127 |
| chr1 | 147372693 | 147378098 | L1MdTf_II | + | 5405 |
| chr1 | 147378098 | 147379091 | L1MdTf_II | + | 993 |
| chr1 | 147547003 | 147552424 | L1MdTf_I | + | 5421 |
| chr1 | 147552424 | 147553417 | L1MdTf_I | + | 993 |
| chr1 | 147647766 | 147647851 | L1MdTf_III | + | 85 |
| chr1 | 147647851 | 147648843 | L1MdTf_III | + | 992 |
| chr1 | 148429018 | 148434224 | L1MdTf_III | + | 5206 |
| chr1 | 148434224 | 148435216 | L1MdTf_III | + | 992 |
| chr1 | 149059335 | 149060414 | L1MdTf_II | - | 1079 |
| chr1 | 149060414 | 149060797 | L1MdTf_II | + | 383 |
| chr1 | 150108185 | 150114188 | L1MdTf_I | + | 6003 |
| chr1 | 150114188 | 150115204 | L1MdTf_I | + | 1016 |
| chr1 | 150148751 | 150154184 | L1MdTf_II | + | 5433 |
| chr1 | 150154184 | 150155203 | L1MdTf_II | + | 1019 |
| chr1 | 150727293 | 150727802 | L1MdTf_II | + | 509 |
| chr1 | 150727802 | 150728823 | L1MdTf_II | + | 1021 |
| chr1 | 157087253 | 157087694 | L1MdTf_III | + | 441 |
| chr1 | 157087694 | 157088678 | L1MdTf_III | + | 984 |
| chr1 | 158159089 | 158165154 | L1MdTf_I | + | 6065 |
| chr1 | 158165154 | 158166172 | L1MdTf_I | + | 1018 |
| chr1 | 158707140 | 158712840 | L1MdTf_II | + | 5700 |
| chr1 | 158712840 | 158713833 | L1MdTf_II | + | 993 |
| chr1 | 158854602 | 158855256 | L1MdTf_III | + | 654 |
| chr1 | 158855256 | 158856248 | L1MdTf_III | + | 992 |

| | | | | | |
|---|---|---|---|---|---|
| chr1 | 160946432 | 160946491 | L1MdTf_III | - | 59 |
| chr1 | 160946491 | 160947460 | L1MdTf_III | + | 969 |
| chr1 | 161760396 | 161765736 | L1MdTf_I | + | 5340 |
| chr1 | 161765736 | 161766729 | L1MdTf_I | + | 993 |
| chr1 | 161891153 | 161896736 | L1MdTf_I | + | 5583 |
| chr1 | 161896736 | 161897729 | L1MdTf_I | + | 993 |
| chr1 | 162317109 | 162322604 | L1MdTf_III | + | 5495 |
| chr1 | 162322604 | 162323597 | L1MdTf_III | + | 993 |
| chr1 | 162627992 | 162629129 | L1MdTf_III | + | 1137 |
| chr1 | 162629129 | 162630143 | L1MdTf_III | + | 1014 |
| chr1 | 162647458 | 162652885 | L1MdTf_II | + | 5427 |
| chr1 | 162652885 | 162653908 | L1MdTf_II | + | 1023 |
| chr1 | 163106005 | 163106226 | L1MdTf_II | + | 221 |
| chr1 | 163106226 | 163107219 | L1MdTf_II | + | 993 |
| chr1 | 163287319 | 163287414 | L1MdTf_III | + | 95 |
| chr1 | 163287414 | 163288407 | L1MdTf_III | + | 993 |
| chr1 | 163611403 | 163617042 | L1MdTf_I | + | 5639 |
| chr1 | 163617042 | 163618035 | L1MdTf_I | + | 993 |
| chr1 | 165237260 | 165237914 | L1MdTf_II | + | 654 |
| chr1 | 165237914 | 165238907 | L1MdTf_II | + | 993 |
| chr1 | 165354183 | 165360456 | L1MdTf_II | + | 6273 |
| chr1 | 165360456 | 165361449 | L1MdTf_II | + | 993 |
| chr1 | 166699894 | 166702505 | L1MdTf_II | + | 2611 |
| chr1 | 166702505 | 166703500 | L1MdTf_II | + | 995 |
| chr1 | 166740030 | 166745461 | L1MdTf_II | + | 5431 |
| chr1 | 166745461 | 166746454 | L1MdTf_II | + | 993 |
| chr1 | 167024516 | 167030104 | L1MdTf_III | + | 5588 |
| chr1 | 167030104 | 167031097 | L1MdTf_III | + | 993 |
| chr1 | 168547917 | 168553389 | L1MdTf_III | + | 5472 |
| chr1 | 168553389 | 168554374 | L1MdTf_III | + | 985 |
| chr1 | 168981215 | 168981377 | L1MdTf_III | - | 162 |
| chr1 | 168981377 | 168981929 | L1MdTf_III | + | 552 |
| chr1 | 169253466 | 169258719 | L1MdTf_III | + | 5253 |
| chr1 | 169258719 | 169259712 | L1MdTf_III | + | 993 |
| chr1 | 170515585 | 170516358 | L1MdTf_III | - | 773 |
| chr1 | 170516358 | 170517407 | L1MdTf_III | + | 1049 |
| chr1 | 172829718 | 172830271 | L1MdTf_III | + | 553 |
| chr1 | 172830271 | 172831264 | L1MdTf_III | + | 993 |
| chr1 | 173660508 | 173665995 | L1MdTf_III | + | 5487 |
| chr1 | 173665995 | 173666988 | L1MdTf_III | + | 993 |
| chr1 | 174770551 | 174776011 | L1MdTf_III | + | 5460 |
| chr1 | 174776011 | 174777004 | L1MdTf_III | + | 993 |
| chr1 | 175623399 | 175626036 | L1MdTf_I | - | 2637 |
| chr1 | 175626036 | 175626478 | L1MdTf_I | + | 442 |
| chr1 | 175644428 | 175649645 | L1MdTf_III | + | 5217 |

| | | | | | |
|---|---|---|---|---|---|
| chr1 | 175649646 | 175650639 | L1MdTf_III | + | 993 |
| chr1 | 176477387 | 176478211 | L1MdTf_II | + | 824 |
| chr1 | 176478211 | 176479204 | L1MdTf_II | + | 993 |
| chr1 | 179098307 | 179098405 | L1MdTf_I | + | 98 |
| chr1 | 179098405 | 179099424 | L1MdTf_I | + | 1019 |
| chr1 | 184887745 | 184893214 | L1MdTf_II | + | 5469 |
| chr1 | 184893214 | 184894207 | L1MdTf_II | + | 993 |
| chr1 | 186794848 | 186800510 | L1MdTf_I | + | 5662 |
| chr1 | 186800510 | 186801503 | L1MdTf_I | + | 993 |
| chr1 | 187124933 | 187131165 | L1MdTf_II | + | 6232 |
| chr1 | 187131165 | 187132158 | L1MdTf_II | + | 993 |
| chr1 | 190346462 | 190347879 | L1MdTf_I | - | 1417 |
| chr1 | 190347879 | 190349408 | L1MdTf_I | + | 1529 |
| chr1 | 192752931 | 192758241 | L1MdTf_I | + | 5310 |
| chr1 | 192758241 | 192759234 | L1MdTf_I | + | 993 |
| chr1 | 193152237 | 193152799 | L1MdTf_II | - | 562 |
| chr1 | 193152799 | 193154132 | L1MdTf_II | + | 1333 |
| chr2 | 3843302 | 3848734 | L1MdTf_III | + | 5432 |
| chr2 | 3848734 | 3849727 | L1MdTf_III | + | 993 |
| chr2 | 4276682 | 4280396 | L1MdTf_III | + | 3714 |
| chr2 | 4280396 | 4281410 | L1MdTf_III | + | 1014 |
| chr2 | 6844851 | 6850471 | L1MdTf_II | + | 5620 |
| chr2 | 6850471 | 6851468 | L1MdTf_II | + | 997 |
| chr2 | 7419904 | 7425302 | L1MdTf_II | + | 5398 |
| chr2 | 7425302 | 7426295 | L1MdTf_II | + | 993 |
| chr2 | 8042230 | 8047670 | L1MdTf_II | + | 5440 |
| chr2 | 8047670 | 8048663 | L1MdTf_II | + | 993 |
| chr2 | 8106963 | 8107225 | L1MdTf_II | - | 262 |
| chr2 | 8107225 | 8107304 | L1MdTf_II | + | 79 |
| chr2 | 8843275 | 8847937 | L1MdTf_II | + | 4662 |
| chr2 | 8847937 | 8848930 | L1MdTf_II | + | 993 |
| chr2 | 9221332 | 9223050 | L1MdTf_I | + | 1718 |
| chr2 | 9223050 | 9224043 | L1MdTf_I | + | 993 |
| chr2 | 10540516 | 10541134 | L1MdTf_III | - | 618 |
| chr2 | 10541134 | 10541436 | L1MdTf_III | + | 302 |
| chr2 | 10648351 | 10648858 | L1MdTf_III | + | 507 |
| chr2 | 10648858 | 10649876 | L1MdTf_III | + | 1018 |
| chr2 | 11126032 | 11132307 | L1MdTf_I | + | 6275 |
| chr2 | 11132307 | 11133300 | L1MdTf_I | + | 993 |
| chr2 | 12955359 | 12958268 | L1MdTf_I | + | 2909 |
| chr2 | 12958268 | 12959288 | L1MdTf_I | + | 1020 |
| chr2 | 13185236 | 13185658 | L1MdTf_II | + | 422 |
| chr2 | 13185658 | 13186651 | L1MdTf_II | + | 993 |
| chr2 | 13662828 | 13668679 | L1MdTf_I | + | 5851 |
| chr2 | 13668679 | 13669672 | L1MdTf_I | + | 993 |

| | | | | | |
|---|---|---|---|---|---|
| chr2 | 14300581 | 14306247 | L1MdTf_III | + | 5666 |
| chr2 | 14306247 | 14307240 | L1MdTf_III | + | 993 |
| chr2 | 14731779 | 14737287 | L1MdTf_III | + | 5508 |
| chr2 | 14737287 | 14738280 | L1MdTf_III | + | 993 |
| chr2 | 15805846 | 15805967 | L1MdTf_II | + | 121 |
| chr2 | 15805967 | 15806950 | L1MdTf_II | + | 983 |
| chr2 | 15919858 | 15925414 | L1MdTf_II | + | 5556 |
| chr2 | 15925414 | 15926410 | L1MdTf_II | + | 996 |
| chr2 | 16950269 | 16954473 | L1MdTf_II | + | 4204 |
| chr2 | 16954473 | 16955466 | L1MdTf_II | + | 993 |
| chr2 | 17491136 | 17496294 | L1MdTf_III | + | 5158 |
| chr2 | 17496294 | 17497345 | L1MdTf_III | + | 1051 |
| chr2 | 18481759 | 18487238 | L1MdTf_II | + | 5479 |
| chr2 | 18487238 | 18488230 | L1MdTf_II | + | 992 |
| chr2 | 19047066 | 19049805 | L1MdTf_II | + | 2739 |
| chr2 | 19049805 | 19050795 | L1MdTf_II | + | 990 |
| chr2 | 19586198 | 19592062 | L1MdTf_II | + | 5864 |
| chr2 | 19592062 | 19593055 | L1MdTf_II | + | 993 |
| chr2 | 23829963 | 23835390 | L1MdTf_II | + | 5427 |
| chr2 | 23835390 | 23836383 | L1MdTf_II | + | 993 |
| chr2 | 23836386 | 23840883 | L1MdTf_II | + | 4497 |
| chr2 | 23840883 | 23841876 | L1MdTf_II | + | 993 |
| chr2 | 23841880 | 23846461 | L1MdTf_II | + | 4581 |
| chr2 | 23846461 | 23846769 | L1MdTf_II | + | 308 |
| chr2 | 25535737 | 25536313 | L1MdTf_I | - | 576 |
| chr2 | 25536313 | 25537168 | L1MdTf_I | + | 855 |
| chr2 | 36998175 | 37000000 | L1MdTf_III | + | 1825 |
| chr2 | 37000000 | 37004013 | L1MdTf_III | + | 4013 |
| chr2 | 37943735 | 37949709 | L1MdTf_I | + | 5974 |
| chr2 | 37949709 | 37950702 | L1MdTf_I | + | 993 |
| chr2 | 40677811 | 40683040 | L1MdTf_III | + | 5229 |
| chr2 | 40683040 | 40684033 | L1MdTf_III | + | 993 |
| chr2 | 41963521 | 41968633 | L1MdTf_II | + | 5112 |
| chr2 | 41968633 | 41969626 | L1MdTf_II | + | 993 |
| chr2 | 42155883 | 42157025 | L1MdTf_I | + | 1142 |
| chr2 | 42157025 | 42158039 | L1MdTf_I | + | 1014 |
| chr2 | 42459717 | 42464985 | L1MdTf_III | + | 5268 |
| chr2 | 42464985 | 42465979 | L1MdTf_III | + | 994 |
| chr2 | 43298615 | 43304078 | L1MdTf_II | + | 5463 |
| chr2 | 43304078 | 43305071 | L1MdTf_II | + | 993 |
| chr2 | 44249063 | 44254467 | L1MdTf_I | + | 5404 |
| chr2 | 44254467 | 44255460 | L1MdTf_I | + | 993 |
| chr2 | 44501461 | 44506900 | L1MdTf_III | + | 5439 |
| chr2 | 44506901 | 44507897 | L1MdTf_III | + | 996 |
| chr2 | 44836432 | 44836585 | L1MdTf_III | - | 153 |

| | | | | | |
|---|---|---|---|---|---|
| chr2 | 44836585 | 44837495 | L1MdTf_III | + | 910 |
| chr2 | 45571719 | 45573134 | L1MdTf_II | + | 1415 |
| chr2 | 45573134 | 45574118 | L1MdTf_II | + | 984 |
| chr2 | 46865404 | 46868008 | L1MdTf_II | + | 2604 |
| chr2 | 46868008 | 46869001 | L1MdTf_II | + | 993 |
| chr2 | 46907305 | 46912526 | L1MdTf_III | + | 5221 |
| chr2 | 46912527 | 46913520 | L1MdTf_III | + | 993 |
| chr2 | 48468908 | 48474636 | L1MdTf_II | + | 5728 |
| chr2 | 48474636 | 48475629 | L1MdTf_II | + | 993 |
| chr2 | 50490319 | 50490421 | L1MdTf_III | + | 102 |
| chr2 | 50490421 | 50495887 | L1MdTf_III | + | 5466 |
| chr2 | 50497163 | 50497499 | L1MdTf_III | + | 336 |
| chr2 | 50497499 | 50498490 | L1MdTf_III | + | 991 |
| chr2 | 50916084 | 50921533 | L1MdTf_III | + | 5449 |
| chr2 | 50921533 | 50922526 | L1MdTf_III | + | 993 |
| chr2 | 50940818 | 50942011 | L1MdTf_III | - | 1193 |
| chr2 | 50942011 | 50942472 | L1MdTf_III | + | 461 |
| chr2 | 51373847 | 51379111 | L1MdTf_III | + | 5264 |
| chr2 | 51379111 | 51380104 | L1MdTf_III | + | 993 |
| chr2 | 53954914 | 53960355 | L1MdTf_III | + | 5441 |
| chr2 | 53960355 | 53961348 | L1MdTf_III | + | 993 |
| chr2 | 54275041 | 54280956 | L1MdTf_I | + | 5915 |
| chr2 | 54280956 | 54281949 | L1MdTf_I | + | 993 |
| chr2 | 54536327 | 54536861 | L1MdTf_II | - | 534 |
| chr2 | 54536861 | 54537526 | L1MdTf_II | + | 665 |
| chr2 | 54607755 | 54613337 | L1MdTf_II | + | 5582 |
| chr2 | 54613337 | 54614321 | L1MdTf_II | + | 984 |
| chr2 | 54631534 | 54636958 | L1MdTf_II | + | 5424 |
| chr2 | 54636958 | 54637951 | L1MdTf_II | + | 993 |
| chr2 | 55103660 | 55105495 | L1MdTf_III | + | 1835 |
| chr2 | 55105495 | 55106516 | L1MdTf_III | + | 1021 |
| chr2 | 55382259 | 55387346 | L1MdTf_II | + | 5087 |
| chr2 | 55387346 | 55388330 | L1MdTf_II | + | 984 |
| chr2 | 55561405 | 55566558 | L1MdTf_II | + | 5153 |
| chr2 | 55566558 | 55567551 | L1MdTf_II | + | 993 |
| chr2 | 56188703 | 56193983 | L1MdTf_II | + | 5280 |
| chr2 | 56193983 | 56194976 | L1MdTf_II | + | 993 |
| chr2 | 56964765 | 56964999 | L1MdTf_III | + | 234 |
| chr2 | 56964999 | 56965992 | L1MdTf_III | + | 993 |
| chr2 | 57900211 | 57901981 | L1MdTf_II | - | 1770 |
| chr2 | 57901981 | 57902482 | L1MdTf_II | + | 501 |
| chr2 | 62904915 | 62908638 | L1MdTf_II | + | 3723 |
| chr2 | 62908638 | 62909631 | L1MdTf_II | + | 993 |
| chr2 | 63227613 | 63233243 | L1MdTf_II | + | 5630 |
| chr2 | 63233243 | 63234258 | L1MdTf_II | + | 1015 |

| | | | | | |
|---|---|---|---|---|---|
| chr2 | 64142961 | 64148941 | L1MdTf_I | + | 5980 |
| chr2 | 64148941 | 64149954 | L1MdTf_I | + | 1013 |
| chr2 | 64438717 | 64438891 | L1MdTf_I | + | 174 |
| chr2 | 64438891 | 64439884 | L1MdTf_I | + | 993 |
| chr2 | 64619255 | 64624522 | L1MdTf_III | + | 5267 |
| chr2 | 64624522 | 64625515 | L1MdTf_III | + | 993 |
| chr2 | 65605432 | 65610860 | L1MdTf_I | + | 5428 |
| chr2 | 65610860 | 65611853 | L1MdTf_I | + | 993 |
| chr2 | 66574548 | 66579740 | L1MdTf_III | + | 5192 |
| chr2 | 66579740 | 66580733 | L1MdTf_III | + | 993 |
| chr2 | 67769467 | 67775352 | L1MdTf_III | + | 5885 |
| chr2 | 67775352 | 67776345 | L1MdTf_III | + | 993 |
| chr2 | 68600704 | 68606121 | L1MdTf_II | + | 5417 |
| chr2 | 68606121 | 68607111 | L1MdTf_II | + | 990 |
| chr2 | 76785646 | 76790862 | L1MdTf_II | + | 5216 |
| chr2 | 76790862 | 76791856 | L1MdTf_II | + | 994 |
| chr2 | 76951449 | 76951801 | L1MdTf_I | - | 352 |
| chr2 | 76951801 | 76952195 | L1MdTf_II | + | 394 |
| chr2 | 77831998 | 77833254 | L1MdTf_III | + | 1256 |
| chr2 | 77833254 | 77834247 | L1MdTf_III | + | 993 |
| chr2 | 78335232 | 78340379 | L1MdTf_I | + | 5147 |
| chr2 | 78340379 | 78341372 | L1MdTf_I | + | 993 |
| chr2 | 78447193 | 78452793 | L1MdTf_I | + | 5600 |
| chr2 | 78452793 | 78453776 | L1MdTf_I | + | 983 |
| chr2 | 79767000 | 79773297 | L1MdTf_II | + | 6297 |
| chr2 | 79773297 | 79774288 | L1MdTf_II | + | 991 |
| chr2 | 82746583 | 82752228 | L1MdTf_II | + | 5645 |
| chr2 | 82752228 | 82753247 | L1MdTf_II | + | 1019 |
| chr2 | 82878614 | 82880335 | L1MdTf_II | + | 1721 |
| chr2 | 82880335 | 82881328 | L1MdTf_II | + | 993 |
| chr2 | 82902782 | 82907992 | L1MdTf_III | + | 5210 |
| chr2 | 82907992 | 82908979 | L1MdTf_III | + | 987 |
| chr2 | 83243845 | 83249874 | L1MdTf_II | + | 6029 |
| chr2 | 83249874 | 83250867 | L1MdTf_II | + | 993 |
| chr2 | 83260284 | 83266139 | L1MdTf_I | + | 5855 |
| chr2 | 83266139 | 83267132 | L1MdTf_I | + | 993 |
| chr2 | 83386901 | 83392956 | L1MdTf_I | + | 6055 |
| chr2 | 83392956 | 83393949 | L1MdTf_I | + | 993 |
| chr2 | 85568899 | 85573899 | L1MdTf_III | + | 5000 |
| chr2 | 85573899 | 85574889 | L1MdTf_III | + | 990 |
| chr2 | 87736730 | 87742175 | L1MdTf_II | + | 5445 |
| chr2 | 87742175 | 87743167 | L1MdTf_II | + | 992 |
| chr2 | 88071556 | 88076983 | L1MdTf_II | + | 5427 |
| chr2 | 88076983 | 88077979 | L1MdTf_II | + | 996 |
| chr2 | 88494521 | 88500000 | L1MdTf_I | - | 5479 |

| | | | | | |
|---|---|---|---|---|---|
| chr2 | 88500000 | 88500977 | L1MdTf_II | - | 977 |
| chr2 | 88978832 | 88979369 | L1MdTf_I | - | 537 |
| chr2 | 88979369 | 88980777 | L1MdTf_I | + | 1408 |
| chr2 | 89917749 | 89923729 | L1MdTf_I | + | 5980 |
| chr2 | 89923729 | 89924722 | L1MdTf_I | + | 993 |
| chr2 | 89997117 | 90000000 | L1MdTf_I | - | 2883 |
| chr2 | 90000000 | 90003770 | L1MdTf_II | - | 3770 |
| chr2 | 90203610 | 90209270 | L1MdTf_I | + | 5660 |
| chr2 | 90209270 | 90210263 | L1MdTf_I | + | 993 |
| chr2 | 95244427 | 95250069 | L1MdTf_I | + | 5642 |
| chr2 | 95250069 | 95251062 | L1MdTf_I | + | 993 |
| chr2 | 95945989 | 95946181 | L1MdTf_II | + | 192 |
| chr2 | 95946181 | 95947174 | L1MdTf_II | + | 993 |
| chr2 | 96264065 | 96266657 | L1MdTf_III | + | 2592 |
| chr2 | 96266657 | 96267647 | L1MdTf_III | + | 990 |
| chr2 | 97231412 | 97231907 | L1MdTf_III | + | 495 |
| chr2 | 97231907 | 97232900 | L1MdTf_III | + | 993 |
| chr2 | 97515330 | 97515746 | L1MdTf_III | + | 416 |
| chr2 | 97515746 | 97516739 | L1MdTf_III | + | 993 |
| chr2 | 97590958 | 97595141 | L1MdTf_I | + | 4183 |
| chr2 | 97595141 | 97596134 | L1MdTf_I | + | 993 |
| chr2 | 97786594 | 97787698 | L1MdTf_II | + | 1104 |
| chr2 | 97787698 | 97788694 | L1MdTf_II | + | 996 |
| chr2 | 98137752 | 98143399 | L1MdTf_I | + | 5647 |
| chr2 | 98143399 | 98144392 | L1MdTf_I | + | 993 |
| chr2 | 98239771 | 98245020 | L1MdTf_III | + | 5249 |
| chr2 | 98245020 | 98246013 | L1MdTf_III | + | 993 |
| chr2 | 98259235 | 98265145 | L1MdTf_II | + | 5910 |
| chr2 | 98265145 | 98266137 | L1MdTf_II | + | 992 |
| chr2 | 98374596 | 98375883 | L1MdTf_III | - | 1287 |
| chr2 | 98375883 | 98376126 | L1MdTf_III | + | 243 |
| chr2 | 98949817 | 98950150 | L1MdTf_III | + | 333 |
| chr2 | 98950150 | 98951142 | L1MdTf_III | + | 992 |
| chr2 | 99082583 | 99082804 | L1MdTf_I | + | 221 |
| chr2 | 99082804 | 99083797 | L1MdTf_I | + | 993 |
| chr2 | 99291091 | 99295631 | L1MdTf_II | - | 4540 |
| chr2 | 99295631 | 99296216 | L1MdTf_II | + | 585 |
| chr2 | 99359812 | 99360210 | L1MdTf_III | + | 398 |
| chr2 | 99360210 | 99361194 | L1MdTf_III | + | 984 |
| chr2 | 99865612 | 99869401 | L1MdTf_III | + | 3789 |
| chr2 | 99869402 | 99870395 | L1MdTf_III | + | 993 |
| chr2 | 101174585 | 101180012 | L1MdTf_I | + | 5427 |
| chr2 | 101180012 | 101181005 | L1MdTf_I | + | 993 |
| chr2 | 101311897 | 101317302 | L1MdTf_II | + | 5405 |
| chr2 | 101317302 | 101318295 | L1MdTf_II | + | 993 |

| | | | | | |
|---|---|---|---|---|---|
| chr2 | 106183614 | 106183856 | L1MdTf_III | + | 242 |
| chr2 | 106183856 | 106184849 | L1MdTf_III | + | 993 |
| chr2 | 106995897 | 107000000 | L1MdTf_III | - | 4103 |
| chr2 | 107000000 | 107002095 | L1MdTf_III | - | 2095 |
| chr2 | 107567583 | 107567801 | L1MdTf_III | - | 218 |
| chr2 | 107567801 | 107567905 | L1MdTf_III | + | 104 |
| chr2 | 107807168 | 107812612 | L1MdTf_II | + | 5444 |
| chr2 | 107812612 | 107813596 | L1MdTf_II | + | 984 |
| chr2 | 108233399 | 108233578 | L1MdTf_I | + | 179 |
| chr2 | 108233578 | 108234571 | L1MdTf_I | + | 993 |
| chr2 | 109654754 | 109654806 | L1MdTf_III | + | 52 |
| chr2 | 109654806 | 109655799 | L1MdTf_III | + | 993 |
| chr2 | 110067558 | 110072965 | L1MdTf_I | + | 5407 |
| chr2 | 110072965 | 110073958 | L1MdTf_I | + | 993 |
| chr2 | 110375999 | 110381212 | L1MdTf_II | + | 5213 |
| chr2 | 110381212 | 110382205 | L1MdTf_II | + | 993 |
| chr2 | 110582401 | 110583257 | L1MdTf_III | - | 856 |
| chr2 | 110583257 | 110585193 | L1MdTf_III | + | 1936 |
| chr2 | 114310484 | 114315910 | L1MdTf_III | + | 5426 |
| chr2 | 114315910 | 114316904 | L1MdTf_III | + | 994 |
| chr2 | 114331998 | 114333466 | L1MdTf_I | + | 1468 |
| chr2 | 114333466 | 114334459 | L1MdTf_I | + | 993 |
| chr2 | 114986321 | 114992684 | L1MdTf_I | - | 6363 |
| chr2 | 114992685 | 114993637 | L1MdTf_III | - | 952 |
| chr2 | 115622202 | 115623656 | L1MdTf_III | + | 1454 |
| chr2 | 115623656 | 115624647 | L1MdTf_III | + | 991 |
| chr2 | 116284786 | 116290001 | L1MdTf_II | + | 5215 |
| chr2 | 116290001 | 116290994 | L1MdTf_II | + | 993 |
| chr2 | 116324565 | 116330434 | L1MdTf_II | + | 5869 |
| chr2 | 116330434 | 116331427 | L1MdTf_II | + | 993 |
| chr2 | 117723947 | 117725161 | L1MdTf_III | - | 1214 |
| chr2 | 117725162 | 117726049 | L1MdTf_III | + | 887 |
| chr2 | 122485272 | 122491118 | L1MdTf_III | + | 5846 |
| chr2 | 122491118 | 122492111 | L1MdTf_III | + | 993 |
| chr2 | 123444578 | 123447161 | L1MdTf_III | + | 2583 |
| chr2 | 123447162 | 123448178 | L1MdTf_III | + | 1016 |
| chr2 | 123722102 | 123727547 | L1MdTf_II | + | 5445 |
| chr2 | 123727547 | 123728539 | L1MdTf_II | + | 992 |
| chr2 | 124723060 | 124728287 | L1MdTf_III | + | 5227 |
| chr2 | 124728287 | 124729280 | L1MdTf_III | + | 993 |
| chr2 | 126080809 | 126086137 | L1MdTf_III | + | 5328 |
| chr2 | 126086137 | 126087130 | L1MdTf_III | + | 993 |
| chr2 | 127635077 | 127640941 | L1MdTf_II | + | 5864 |
| chr2 | 127640941 | 127641942 | L1MdTf_II | + | 1001 |
| chr2 | 128976214 | 128978403 | L1MdTf_III | + | 2189 |

| | | | | | |
|---|---|---|---|---|---|
| chr2 | 128978403 | 128979396 | L1MdTf_III | + | 993 |
| chr2 | 130162557 | 130163861 | L1MdTf_I | - | 1304 |
| chr2 | 130163861 | 130164485 | L1MdTf_II | - | 624 |
| chr2 | 133081177 | 133086181 | L1MdTf_III | + | 5004 |
| chr2 | 133086181 | 133087174 | L1MdTf_III | + | 993 |
| chr2 | 134317503 | 134317989 | L1MdTf_II | - | 486 |
| chr2 | 134317989 | 134318413 | L1MdTf_II | + | 424 |
| chr2 | 134417031 | 134418800 | L1MdTf_I | + | 1769 |
| chr2 | 134418800 | 134419793 | L1MdTf_I | + | 993 |
| chr2 | 134988719 | 134989997 | L1MdTf_II | - | 1278 |
| chr2 | 134989997 | 134990996 | L1MdTf_II | + | 999 |
| chr2 | 135354100 | 135359519 | L1MdTf_II | + | 5419 |
| chr2 | 135359519 | 135360512 | L1MdTf_II | + | 993 |
| chr2 | 136126610 | 136127885 | L1MdTf_III | + | 1275 |
| chr2 | 136127885 | 136128878 | L1MdTf_III | + | 993 |
| chr2 | 137611435 | 137611586 | L1MdTf_I | + | 151 |
| chr2 | 137611586 | 137612579 | L1MdTf_I | + | 993 |
| chr2 | 140382063 | 140387534 | L1MdTf_III | + | 5471 |
| chr2 | 140387534 | 140388527 | L1MdTf_III | + | 993 |
| chr2 | 140945100 | 140950694 | L1MdTf_II | + | 5594 |
| chr2 | 140950694 | 140951691 | L1MdTf_II | + | 997 |
| chr2 | 142793404 | 142794148 | L1MdTf_II | + | 744 |
| chr2 | 142794148 | 142795140 | L1MdTf_II | + | 992 |
| chr2 | 143567810 | 143573423 | L1MdTf_II | + | 5613 |
| chr2 | 143573423 | 143574443 | L1MdTf_II | + | 1020 |
| chr2 | 144733509 | 144734846 | L1MdTf_I | + | 1337 |
| chr2 | 144734846 | 144735839 | L1MdTf_I | + | 993 |
| chr2 | 145999405 | 146000000 | L1MdTf_III | - | 595 |
| chr2 | 146000000 | 146005628 | L1MdTf_III | - | 5628 |
| chr2 | 147189960 | 147191381 | L1MdTf_III | - | 1421 |
| chr2 | 147191381 | 147191766 | L1MdTf_III | + | 385 |
| chr2 | 148302267 | 148307979 | L1MdTf_I | + | 5712 |
| chr2 | 148307979 | 148308972 | L1MdTf_I | + | 993 |
| chr2 | 148599029 | 148604496 | L1MdTf_III | + | 5467 |
| chr2 | 148604496 | 148605490 | L1MdTf_III | + | 994 |
| chr2 | 149132991 | 149133645 | L1MdTf_III | + | 654 |
| chr2 | 149133646 | 149134638 | L1MdTf_III | + | 992 |
| chr2 | 150282464 | 150288281 | L1MdTf_I | + | 5817 |
| chr2 | 150288281 | 150289274 | L1MdTf_I | + | 993 |
| chr2 | 153934631 | 153935285 | L1MdTf_II | + | 654 |
| chr2 | 153935285 | 153936277 | L1MdTf_II | + | 992 |
| chr2 | 159188039 | 159193509 | L1MdTf_II | + | 5470 |
| chr2 | 159193509 | 159194505 | L1MdTf_II | + | 996 |
| chr2 | 159498169 | 159500000 | L1MdTf_II | + | 1831 |
| chr2 | 159500000 | 159503594 | L1MdTf_I | + | 3594 |

| | | | | | |
|---|---|---|---|---|---|
| chr2 | 161490165 | 161495369 | L1MdTf_II | + | 5204 |
| chr2 | 161495369 | 161496358 | L1MdTf_II | + | 989 |
| chr2 | 161496359 | 161496677 | L1MdTf_I | + | 318 |
| chr2 | 161496677 | 161497670 | L1MdTf_I | + | 993 |
| chr2 | 161984838 | 161985299 | L1MdTf_III | - | 461 |
| chr2 | 161985299 | 161985508 | L1MdTf_III | + | 209 |
| chr2 | 162216935 | 162217686 | L1MdTf_I | + | 751 |
| chr2 | 162217686 | 162218679 | L1MdTf_I | + | 993 |
| chr2 | 170925582 | 170930592 | L1MdTf_II | + | 5010 |
| chr2 | 170930592 | 170931607 | L1MdTf_II | + | 1015 |
| chr2 | 171418994 | 171424874 | L1MdTf_III | + | 5880 |
| chr2 | 171424874 | 171425888 | L1MdTf_III | + | 1014 |
| chr2 | 171848973 | 171855255 | L1MdTf_I | + | 6282 |
| chr2 | 171855255 | 171856268 | L1MdTf_I | + | 1013 |
| chr2 | 175104399 | 175105306 | L1MdTf_I | + | 907 |
| chr2 | 175105306 | 175106299 | L1MdTf_I | + | 993 |
| chr2 | 176674472 | 176679896 | L1MdTf_II | + | 5424 |
| chr2 | 176679896 | 176680888 | L1MdTf_II | + | 992 |
| chr2 | 178416009 | 178421423 | L1MdTf_III | + | 5414 |
| chr2 | 178421423 | 178422420 | L1MdTf_III | + | 997 |
| chr3 | 3994028 | 4000000 | L1MdTf_III | - | 5972 |
| chr3 | 4000000 | 4000709 | L1MdTf_III | - | 709 |
| chr3 | 4649575 | 4655426 | L1MdTf_I | + | 5851 |
| chr3 | 4655426 | 4656448 | L1MdTf_I | + | 1022 |
| chr3 | 6915027 | 6920495 | L1MdTf_II | + | 5468 |
| chr3 | 6920495 | 6921490 | L1MdTf_II | + | 995 |
| chr3 | 7147949 | 7153399 | L1MdTf_I | + | 5450 |
| chr3 | 7153399 | 7154392 | L1MdTf_I | + | 993 |
| chr3 | 7156667 | 7161906 | L1MdTf_III | + | 5239 |
| chr3 | 7161906 | 7162899 | L1MdTf_III | + | 993 |
| chr3 | 7282378 | 7288026 | L1MdTf_II | + | 5648 |
| chr3 | 7288026 | 7289019 | L1MdTf_II | + | 993 |
| chr3 | 7376204 | 7381631 | L1MdTf_II | + | 5427 |
| chr3 | 7381631 | 7382646 | L1MdTf_II | + | 1015 |
| chr3 | 7975406 | 7980645 | L1MdTf_II | + | 5239 |
| chr3 | 7980645 | 7981638 | L1MdTf_II | + | 993 |
| chr3 | 10741716 | 10742655 | L1MdTf_III | + | 939 |
| chr3 | 10742655 | 10743648 | L1MdTf_III | + | 993 |
| chr3 | 11075488 | 11075831 | L1MdTf_III | + | 343 |
| chr3 | 11075832 | 11076825 | L1MdTf_III | + | 993 |
| chr3 | 11655507 | 11660967 | L1MdTf_I | + | 5460 |
| chr3 | 11660967 | 11661960 | L1MdTf_I | + | 993 |
| chr3 | 11856346 | 11861787 | L1MdTf_III | + | 5441 |
| chr3 | 11861788 | 11862779 | L1MdTf_III | + | 991 |
| chr3 | 12295430 | 12300917 | L1MdTf_III | + | 5487 |

| | | | | | |
|---|---|---|---|---|---|
| chr3 | 12300917 | 12301909 | L1MdTf_III | + | 992 |
| chr3 | 12881436 | 12883655 | L1MdTf_III | + | 2219 |
| chr3 | 12883655 | 12884648 | L1MdTf_III | + | 993 |
| chr3 | 15093515 | 15098985 | L1MdTf_II | + | 5470 |
| chr3 | 15098985 | 15099978 | L1MdTf_II | + | 993 |
| chr3 | 15144540 | 15149752 | L1MdTf_II | + | 5212 |
| chr3 | 15149752 | 15150764 | L1MdTf_II | + | 1012 |
| chr3 | 16142753 | 16147968 | L1MdTf_III | + | 5215 |
| chr3 | 16147968 | 16148968 | L1MdTf_III | + | 1000 |
| chr3 | 17052380 | 17054033 | L1MdTf_III | + | 1653 |
| chr3 | 17054033 | 17055026 | L1MdTf_III | + | 993 |
| chr3 | 17274578 | 17279920 | L1MdTf_II | + | 5342 |
| chr3 | 17279921 | 17280913 | L1MdTf_II | + | 992 |
| chr3 | 17437741 | 17442998 | L1MdTf_II | + | 5257 |
| chr3 | 17442998 | 17444008 | L1MdTf_II | + | 1010 |
| chr3 | 18485430 | 18490676 | L1MdTf_III | + | 5246 |
| chr3 | 18490676 | 18491669 | L1MdTf_III | + | 993 |
| chr3 | 18715761 | 18715818 | L1MdTf_II | - | 57 |
| chr3 | 18715818 | 18716351 | L1MdTf_II | + | 533 |
| chr3 | 19031406 | 19036935 | L1MdTf_I | + | 5529 |
| chr3 | 19036936 | 19037929 | L1MdTf_I | + | 993 |
| chr3 | 19904172 | 19907599 | L1MdTf_II | + | 3427 |
| chr3 | 19907599 | 19908592 | L1MdTf_II | + | 993 |
| chr3 | 20887697 | 20893182 | L1MdTf_III | + | 5485 |
| chr3 | 20893182 | 20894175 | L1MdTf_III | + | 993 |
| chr3 | 23407714 | 23413136 | L1MdTf_II | + | 5422 |
| chr3 | 23413136 | 23414134 | L1MdTf_II | + | 998 |
| chr3 | 23824884 | 23825971 | L1MdTf_II | + | 1087 |
| chr3 | 23825972 | 23826965 | L1MdTf_II | + | 993 |
| chr3 | 25038836 | 25044074 | L1MdTf_I | + | 5238 |
| chr3 | 25044074 | 25045061 | L1MdTf_I | + | 987 |
| chr3 | 25088311 | 25093744 | L1MdTf_III | + | 5433 |
| chr3 | 25093745 | 25094737 | L1MdTf_III | + | 992 |
| chr3 | 25305400 | 25310036 | L1MdTf_III | + | 4636 |
| chr3 | 25310036 | 25311055 | L1MdTf_III | + | 1019 |
| chr3 | 26838142 | 26843407 | L1MdTf_III | + | 5265 |
| chr3 | 26843407 | 26844401 | L1MdTf_III | + | 994 |
| chr3 | 28920957 | 28926342 | L1MdTf_I | + | 5385 |
| chr3 | 28926342 | 28927335 | L1MdTf_I | + | 993 |
| chr3 | 29573153 | 29578368 | L1MdTf_II | + | 5215 |
| chr3 | 29578368 | 29579361 | L1MdTf_II | + | 993 |
| chr3 | 29790811 | 29791204 | L1MdTf_II | + | 393 |
| chr3 | 29791204 | 29792197 | L1MdTf_II | + | 993 |
| chr3 | 33563585 | 33564148 | L1MdTf_II | - | 563 |
| chr3 | 33564148 | 33564990 | L1MdTf_II | + | 842 |

| | | | | | |
|---|---|---|---|---|---|
| chr3 | 33653812 | 33659206 | L1MdTf_III | + | 5394 |
| chr3 | 33659206 | 33660199 | L1MdTf_III | + | 993 |
| chr3 | 33822276 | 33828126 | L1MdTf_I | + | 5850 |
| chr3 | 33828126 | 33829139 | L1MdTf_I | + | 1013 |
| chr3 | 39199797 | 39199900 | L1MdTf_I | + | 103 |
| chr3 | 39199900 | 39200893 | L1MdTf_I | + | 993 |
| chr3 | 39272407 | 39272780 | L1MdTf_I | + | 373 |
| chr3 | 39272780 | 39273773 | L1MdTf_I | + | 993 |
| chr3 | 39656129 | 39661768 | L1MdTf_I | + | 5639 |
| chr3 | 39661768 | 39662761 | L1MdTf_I | + | 993 |
| chr3 | 39911874 | 39917217 | L1MdTf_III | + | 5343 |
| chr3 | 39917217 | 39918229 | L1MdTf_III | + | 1012 |
| chr3 | 40205424 | 40210791 | L1MdTf_II | + | 5367 |
| chr3 | 40210791 | 40211784 | L1MdTf_II | + | 993 |
| chr3 | 42036329 | 42036736 | L1MdTf_II | + | 407 |
| chr3 | 42036736 | 42037729 | L1MdTf_II | + | 993 |
| chr3 | 42201446 | 42201972 | L1MdTf_III | - | 526 |
| chr3 | 42201972 | 42202921 | L1MdTf_III | + | 949 |
| chr3 | 43062441 | 43064546 | L1MdTf_I | + | 2105 |
| chr3 | 43064546 | 43065539 | L1MdTf_I | + | 993 |
| chr3 | 43141075 | 43146195 | L1MdTf_II | + | 5120 |
| chr3 | 43146195 | 43147191 | L1MdTf_II | + | 996 |
| chr3 | 44053691 | 44059760 | L1MdTf_I | + | 6069 |
| chr3 | 44059760 | 44060780 | L1MdTf_I | + | 1020 |
| chr3 | 44491429 | 44492265 | L1MdTf_III | - | 836 |
| chr3 | 44492265 | 44492816 | L1MdTf_III | + | 551 |
| chr3 | 44576982 | 44582410 | L1MdTf_I | + | 5428 |
| chr3 | 44582410 | 44583402 | L1MdTf_I | + | 992 |
| chr3 | 44620186 | 44625851 | L1MdTf_II | + | 5665 |
| chr3 | 44625851 | 44626844 | L1MdTf_II | + | 993 |
| chr3 | 44811954 | 44817323 | L1MdTf_II | + | 5369 |
| chr3 | 44817323 | 44818316 | L1MdTf_II | + | 993 |
| chr3 | 44838869 | 44844193 | L1MdTf_III | + | 5324 |
| chr3 | 44844193 | 44845177 | L1MdTf_III | + | 984 |
| chr3 | 45302621 | 45307855 | L1MdTf_II | + | 5234 |
| chr3 | 45307855 | 45308852 | L1MdTf_II | + | 997 |
| chr3 | 45532652 | 45538956 | L1MdTf_II | + | 6304 |
| chr3 | 45538956 | 45539949 | L1MdTf_II | + | 993 |
| chr3 | 46153012 | 46158651 | L1MdTf_I | + | 5639 |
| chr3 | 46158651 | 46159644 | L1MdTf_I | + | 993 |
| chr3 | 46689417 | 46695347 | L1MdTf_I | + | 5930 |
| chr3 | 46695347 | 46696360 | L1MdTf_I | + | 1013 |
| chr3 | 47237391 | 47242778 | L1MdTf_III | + | 5387 |
| chr3 | 47242779 | 47243772 | L1MdTf_III | + | 993 |
| chr3 | 49077949 | 49081596 | L1MdTf_III | + | 3647 |

| | | | | | |
|---|---|---|---|---|---|
| chr3 | 49081596 | 49082589 | L1MdTf_III | + | 993 |
| chr3 | 49398020 | 49403259 | L1MdTf_III | + | 5239 |
| chr3 | 49403259 | 49404252 | L1MdTf_III | + | 993 |
| chr3 | 50008140 | 50013465 | L1MdTf_III | + | 5325 |
| chr3 | 50013465 | 50014484 | L1MdTf_III | + | 1019 |
| chr3 | 50045070 | 50045712 | L1MdTf_III | + | 642 |
| chr3 | 50045712 | 50046686 | L1MdTf_III | + | 974 |
| chr3 | 52790481 | 52791161 | L1MdTf_I | - | 680 |
| chr3 | 52791161 | 52791970 | L1MdTf_I | + | 809 |
| chr3 | 53093810 | 53099023 | L1MdTf_I | + | 5213 |
| chr3 | 53099023 | 53100012 | L1MdTf_I | + | 989 |
| chr3 | 56043670 | 56049069 | L1MdTf_I | + | 5399 |
| chr3 | 56049069 | 56050091 | L1MdTf_I | + | 1022 |
| chr3 | 56116696 | 56122236 | L1MdTf_III | + | 5540 |
| chr3 | 56122236 | 56123229 | L1MdTf_III | + | 993 |
| chr3 | 56421024 | 56426673 | L1MdTf_II | + | 5649 |
| chr3 | 56426673 | 56427669 | L1MdTf_II | + | 996 |
| chr3 | 56811312 | 56816734 | L1MdTf_II | + | 5422 |
| chr3 | 56816734 | 56817756 | L1MdTf_II | + | 1022 |
| chr3 | 58670120 | 58675482 | L1MdTf_III | + | 5362 |
| chr3 | 58675482 | 58676475 | L1MdTf_III | + | 993 |
| chr3 | 59209143 | 59210207 | L1MdTf_I | + | 1064 |
| chr3 | 59210207 | 59211200 | L1MdTf_I | + | 993 |
| chr3 | 59532822 | 59539161 | L1MdTf_III | + | 6339 |
| chr3 | 59539161 | 59540156 | L1MdTf_III | + | 995 |
| chr3 | 59699185 | 59699707 | L1MdTf_III | + | 522 |
| chr3 | 59699707 | 59700702 | L1MdTf_III | + | 995 |
| chr3 | 60604096 | 60604310 | L1MdTf_II | + | 214 |
| chr3 | 60604310 | 60605302 | L1MdTf_II | + | 992 |
| chr3 | 60966451 | 60966946 | L1MdTf_II | + | 495 |
| chr3 | 60966946 | 60967939 | L1MdTf_II | + | 993 |
| chr3 | 61627519 | 61632959 | L1MdTf_II | + | 5440 |
| chr3 | 61632959 | 61633952 | L1MdTf_II | + | 993 |
| chr3 | 62029654 | 62034887 | L1MdTf_II | + | 5233 |
| chr3 | 62034887 | 62035880 | L1MdTf_II | + | 993 |
| chr3 | 62113334 | 62119175 | L1MdTf_I | + | 5841 |
| chr3 | 62119175 | 62120168 | L1MdTf_I | + | 993 |
| chr3 | 62480818 | 62486239 | L1MdTf_III | + | 5421 |
| chr3 | 62486239 | 62487232 | L1MdTf_III | + | 993 |
| chr3 | 66341150 | 66342078 | L1MdTf_I | - | 928 |
| chr3 | 66342078 | 66342201 | L1MdTf_I | + | 123 |
| chr3 | 66615509 | 66618253 | L1MdTf_II | + | 2744 |
| chr3 | 66618253 | 66619246 | L1MdTf_II | + | 993 |
| chr3 | 66809746 | 66814952 | L1MdTf_II | + | 5206 |
| chr3 | 66814952 | 66815945 | L1MdTf_I | + | 993 |

| | | | | | |
|---|---|---|---|---|---|
| chr3 | 66815945 | 66816085 | L1MdTf_II | + | 140 |
| chr3 | 66816085 | 66817078 | L1MdTf_II | + | 993 |
| chr3 | 66817880 | 66826774 | L1MdTf_II | + | 8894 |
| chr3 | 66826774 | 66827767 | L1MdTf_II | + | 993 |
| chr3 | 67200656 | 67203838 | L1MdTf_I | + | 3182 |
| chr3 | 67203838 | 67204858 | L1MdTf_I | + | 1020 |
| chr3 | 67694029 | 67699877 | L1MdTf_II | + | 5848 |
| chr3 | 67699877 | 67700874 | L1MdTf_II | + | 997 |
| chr3 | 69803186 | 69803908 | L1MdTf_II | - | 722 |
| chr3 | 69803908 | 69805390 | L1MdTf_I | + | 1482 |
| chr3 | 70625945 | 70631392 | L1MdTf_II | + | 5447 |
| chr3 | 70631392 | 70632388 | L1MdTf_II | + | 996 |
| chr3 | 70682024 | 70682503 | L1MdTf_II | + | 479 |
| chr3 | 70682503 | 70683495 | L1MdTf_II | + | 992 |
| chr3 | 70716448 | 70717658 | L1MdTf_I | - | 1210 |
| chr3 | 70717658 | 70718306 | L1MdTf_I | + | 648 |
| chr3 | 70995395 | 70996860 | L1MdTf_I | - | 1465 |
| chr3 | 70996860 | 70997289 | L1MdTf_I | + | 429 |
| chr3 | 71173580 | 71173870 | L1MdTf_II | - | 290 |
| chr3 | 71173870 | 71173979 | L1MdTf_II | + | 109 |
| chr3 | 71317703 | 71318324 | L1MdTf_II | + | 621 |
| chr3 | 71318324 | 71319317 | L1MdTf_II | + | 993 |
| chr3 | 71678565 | 71680212 | L1MdTf_I | + | 1647 |
| chr3 | 71680212 | 71681206 | L1MdTf_I | + | 994 |
| chr3 | 72605464 | 72611147 | L1MdTf_III | + | 5683 |
| chr3 | 72611147 | 72612140 | L1MdTf_III | + | 993 |
| chr3 | 73771382 | 73777038 | L1MdTf_I | + | 5656 |
| chr3 | 73777038 | 73778031 | L1MdTf_I | + | 993 |
| chr3 | 74377923 | 74383610 | L1MdTf_III | + | 5687 |
| chr3 | 74383610 | 74384603 | L1MdTf_III | + | 993 |
| chr3 | 75279649 | 75285083 | L1MdTf_II | + | 5434 |
| chr3 | 75285083 | 75286076 | L1MdTf_II | + | 993 |
| chr3 | 76740771 | 76741151 | L1MdTf_III | - | 380 |
| chr3 | 76741151 | 76741453 | L1MdTf_III | + | 302 |
| chr3 | 77140951 | 77146386 | L1MdTf_III | + | 5435 |
| chr3 | 77146386 | 77147378 | L1MdTf_III | + | 992 |
| chr3 | 77612475 | 77617957 | L1MdTf_III | + | 5482 |
| chr3 | 77617958 | 77618951 | L1MdTf_III | + | 993 |
| chr3 | 78403687 | 78407493 | L1MdTf_I | + | 3806 |
| chr3 | 78407493 | 78408486 | L1MdTf_I | + | 993 |
| chr3 | 78653037 | 78657799 | L1MdTf_II | + | 4762 |
| chr3 | 78657799 | 78658792 | L1MdTf_II | + | 993 |
| chr3 | 78724592 | 78730600 | L1MdTf_I | + | 6008 |
| chr3 | 78730600 | 78731593 | L1MdTf_I | + | 993 |
| chr3 | 79546448 | 79547102 | L1MdTf_I | + | 654 |

| | | | | | |
|---|---|---|---|---|---|
| chr3 | 79547102 | 79548120 | L1MdTf_I | + | 1018 |
| chr3 | 80372699 | 80378356 | L1MdTf_II | + | 5657 |
| chr3 | 80378356 | 80379349 | L1MdTf_II | + | 993 |
| chr3 | 80445843 | 80446164 | L1MdTf_III | + | 321 |
| chr3 | 80446164 | 80447157 | L1MdTf_III | + | 993 |
| chr3 | 80649720 | 80655399 | L1MdTf_I | + | 5679 |
| chr3 | 80655399 | 80656392 | L1MdTf_I | + | 993 |
| chr3 | 81596865 | 81602113 | L1MdTf_II | + | 5248 |
| chr3 | 81602113 | 81603130 | L1MdTf_II | + | 1017 |
| chr3 | 82414704 | 82415160 | L1MdTf_III | + | 456 |
| chr3 | 82415160 | 82416152 | L1MdTf_III | + | 992 |
| chr3 | 84905771 | 84906325 | L1MdTf_II | - | 554 |
| chr3 | 84906326 | 84906759 | L1MdTf_I | + | 433 |
| chr3 | 86388068 | 86389124 | L1MdTf_I | + | 1056 |
| chr3 | 86389124 | 86390117 | L1MdTf_I | + | 993 |
| chr3 | 90766996 | 90768385 | L1MdTf_II | - | 1389 |
| chr3 | 90768385 | 90769286 | L1MdTf_II | + | 901 |
| chr3 | 91033932 | 91040419 | L1MdTf_II | + | 6487 |
| chr3 | 91040419 | 91041412 | L1MdTf_I | + | 993 |
| chr3 | 91609278 | 91614562 | L1MdTf_III | + | 5284 |
| chr3 | 91614562 | 91615555 | L1MdTf_III | + | 993 |
| chr3 | 91803166 | 91804703 | L1MdTf_III | + | 1537 |
| chr3 | 91804704 | 91805697 | L1MdTf_III | + | 993 |
| chr3 | 92520870 | 92521652 | L1MdTf_II | - | 782 |
| chr3 | 92521652 | 92521808 | L1MdTf_II | + | 156 |
| chr3 | 94175691 | 94181364 | L1MdTf_I | + | 5673 |
| chr3 | 94181364 | 94182357 | L1MdTf_I | + | 993 |
| chr3 | 94456883 | 94462675 | L1MdTf_I | + | 5792 |
| chr3 | 94462675 | 94463668 | L1MdTf_I | + | 993 |
| chr3 | 102300678 | 102306050 | L1MdTf_III | + | 5372 |
| chr3 | 102306050 | 102307042 | L1MdTf_III | + | 992 |
| chr3 | 102739982 | 102745654 | L1MdTf_I | + | 5672 |
| chr3 | 102745654 | 102746647 | L1MdTf_I | + | 993 |
| chr3 | 102773497 | 102778938 | L1MdTf_III | + | 5441 |
| chr3 | 102778938 | 102779930 | L1MdTf_III | + | 992 |
| chr3 | 103334859 | 103335622 | L1MdTf_II | + | 763 |
| chr3 | 103335622 | 103336643 | L1MdTf_II | + | 1021 |
| chr3 | 105252608 | 105253691 | L1MdTf_II | + | 1083 |
| chr3 | 105253691 | 105254687 | L1MdTf_II | + | 996 |
| chr3 | 106120690 | 106120813 | L1MdTf_III | - | 123 |
| chr3 | 106120813 | 106121251 | L1MdTf_III | + | 438 |
| chr3 | 106192476 | 106197953 | L1MdTf_III | + | 5477 |
| chr3 | 106197953 | 106198946 | L1MdTf_III | + | 993 |
| chr3 | 106321772 | 106327244 | L1MdTf_III | + | 5472 |
| chr3 | 106327244 | 106328236 | L1MdTf_III | + | 992 |

| | | | | | |
|---|---|---|---|---|---|
| chr3 | 110198823 | 110201508 | L1MdTf_I | + | 2685 |
| chr3 | 110201508 | 110202518 | L1MdTf_I | + | 1010 |
| chr3 | 111971966 | 111975678 | L1MdTf_I | + | 3712 |
| chr3 | 111975678 | 111976671 | L1MdTf_I | + | 993 |
| chr3 | 112188420 | 112193660 | L1MdTf_II | + | 5240 |
| chr3 | 112193660 | 112194653 | L1MdTf_II | + | 993 |
| chr3 | 113627829 | 113633039 | L1MdTf_I | + | 5210 |
| chr3 | 113633039 | 113634032 | L1MdTf_I | + | 993 |
| chr3 | 114463295 | 114468505 | L1MdTf_III | + | 5210 |
| chr3 | 114468505 | 114469498 | L1MdTf_III | + | 993 |
| chr3 | 115701752 | 115701908 | L1MdTf_III | - | 156 |
| chr3 | 115701908 | 115702683 | L1MdTf_III | + | 775 |
| chr3 | 115709559 | 115715601 | L1MdTf_II | + | 6042 |
| chr3 | 115715601 | 115716616 | L1MdTf_II | + | 1015 |
| chr3 | 116811794 | 116812907 | L1MdTf_III | + | 1113 |
| chr3 | 116812907 | 116813899 | L1MdTf_III | + | 992 |
| chr3 | 116823616 | 116829122 | L1MdTf_II | + | 5506 |
| chr3 | 116829122 | 116830115 | L1MdTf_II | + | 993 |
| chr3 | 117848702 | 117851209 | L1MdTf_II | - | 2507 |
| chr3 | 117851209 | 117852697 | L1MdTf_II | + | 1488 |
| chr3 | 118507418 | 118512834 | L1MdTf_II | + | 5416 |
| chr3 | 118512834 | 118513827 | L1MdTf_II | + | 993 |
| chr3 | 118718466 | 118719512 | L1MdTf_III | - | 1046 |
| chr3 | 118719512 | 118720394 | L1MdTf_III | + | 882 |
| chr3 | 119898324 | 119904219 | L1MdTf_II | + | 5895 |
| chr3 | 119904219 | 119905214 | L1MdTf_II | + | 995 |
| chr3 | 120114688 | 120115916 | L1MdTf_I | - | 1228 |
| chr3 | 120115916 | 120116024 | L1MdTf_I | + | 108 |
| chr3 | 120618860 | 120625347 | L1MdTf_II | + | 6487 |
| chr3 | 120625347 | 120626340 | L1MdTf_II | + | 993 |
| chr3 | 120640462 | 120644206 | L1MdTf_III | + | 3744 |
| chr3 | 120644206 | 120645199 | L1MdTf_III | + | 993 |
| chr3 | 120732211 | 120737824 | L1MdTf_I | + | 5613 |
| chr3 | 120737824 | 120738817 | L1MdTf_I | + | 993 |
| chr3 | 120856501 | 120858524 | L1MdTf_I | + | 2023 |
| chr3 | 120858524 | 120859517 | L1MdTf_I | + | 993 |
| chr3 | 122988449 | 122993705 | L1MdTf_III | + | 5256 |
| chr3 | 122993705 | 122994698 | L1MdTf_III | + | 993 |
| chr3 | 123619650 | 123619845 | L1MdTf_II | - | 195 |
| chr3 | 123619845 | 123619921 | L1MdTf_II | + | 76 |
| chr3 | 124075843 | 124081053 | L1MdTf_III | + | 5210 |
| chr3 | 124081053 | 124082046 | L1MdTf_III | + | 993 |
| chr3 | 126068428 | 126068510 | L1MdTf_II | - | 82 |
| chr3 | 126068510 | 126068670 | L1MdTf_II | + | 160 |
| chr3 | 127931641 | 127937278 | L1MdTf_II | + | 5637 |

| | | | | | |
|---|---|---|---|---|---|
| chr3 | 127937278 | 127938271 | L1MdTf_II | + | 993 |
| chr3 | 128336762 | 128337011 | L1MdTf_I | - | 249 |
| chr3 | 128337011 | 128337585 | L1MdTf_II | + | 574 |
| chr3 | 128818691 | 128824778 | L1MdTf_III | + | 6087 |
| chr3 | 128824778 | 128825771 | L1MdTf_III | + | 993 |
| chr3 | 129907966 | 129908134 | L1MdTf_III | - | 168 |
| chr3 | 129908134 | 129910075 | L1MdTf_III | + | 1941 |
| chr3 | 130567959 | 130573386 | L1MdTf_I | + | 5427 |
| chr3 | 130573386 | 130574379 | L1MdTf_I | + | 993 |
| chr3 | 131412417 | 131417702 | L1MdTf_III | + | 5285 |
| chr3 | 131417702 | 131418715 | L1MdTf_III | + | 1013 |
| chr3 | 134521104 | 134526543 | L1MdTf_II | + | 5439 |
| chr3 | 134526543 | 134527536 | L1MdTf_II | + | 993 |
| chr3 | 134586698 | 134593188 | L1MdTf_I | + | 6490 |
| chr3 | 134593188 | 134594181 | L1MdTf_I | + | 993 |
| chr3 | 135725043 | 135726189 | L1MdTf_III | + | 1146 |
| chr3 | 135726189 | 135727182 | L1MdTf_III | + | 993 |
| chr3 | 135991276 | 135991567 | L1MdTf_III | + | 291 |
| chr3 | 135991567 | 135992578 | L1MdTf_III | + | 1011 |
| chr3 | 137306184 | 137311646 | L1MdTf_I | + | 5462 |
| chr3 | 137311646 | 137312639 | L1MdTf_I | + | 993 |
| chr3 | 137364444 | 137369797 | L1MdTf_III | + | 5353 |
| chr3 | 137369797 | 137370790 | L1MdTf_III | + | 993 |
| chr3 | 139073103 | 139078318 | L1MdTf_I | + | 5215 |
| chr3 | 139078318 | 139079335 | L1MdTf_I | + | 1017 |
| chr3 | 139270491 | 139276543 | L1MdTf_I | + | 6052 |
| chr3 | 139276543 | 139277536 | L1MdTf_I | + | 993 |
| chr3 | 139862380 | 139868271 | L1MdTf_I | + | 5891 |
| chr3 | 139868271 | 139869284 | L1MdTf_I | + | 1013 |
| chr3 | 140218214 | 140223698 | L1MdTf_III | + | 5484 |
| chr3 | 140223698 | 140224692 | L1MdTf_III | + | 994 |
| chr3 | 144360183 | 144365568 | L1MdTf_III | + | 5385 |
| chr3 | 144365568 | 144366561 | L1MdTf_III | + | 993 |
| chr3 | 144534174 | 144540581 | L1MdTf_II | + | 6407 |
| chr3 | 144540581 | 144541574 | L1MdTf_II | + | 993 |
| chr3 | 145999493 | 146000000 | L1MdTf_III | + | 507 |
| chr3 | 146000000 | 146003931 | L1MdTf_III | + | 3931 |
| chr3 | 148023594 | 148025426 | L1MdTf_II | + | 1832 |
| chr3 | 148025426 | 148026419 | L1MdTf_II | + | 993 |
| chr3 | 149850729 | 149850811 | L1MdTf_III | + | 82 |
| chr3 | 149850811 | 149851804 | L1MdTf_III | + | 993 |
| chr3 | 152796296 | 152798125 | L1MdTf_I | - | 1829 |
| chr3 | 152798125 | 152799376 | L1MdTf_I | + | 1251 |
| chr3 | 155398614 | 155399643 | L1MdTf_I | - | 1029 |
| chr3 | 155399643 | 155399876 | L1MdTf_II | + | 233 |

| | | | | | |
|---|---|---|---|---|---|
| chr3 | 158402925 | 158408149 | L1MdTf_III | + | 5224 |
| chr3 | 158408149 | 158409142 | L1MdTf_III | + | 993 |
| chr3 | 158426171 | 158426633 | L1MdTf_III | + | 462 |
| chr3 | 158426633 | 158427626 | L1MdTf_III | + | 993 |
| chr3 | 158693507 | 158698956 | L1MdTf_II | + | 5449 |
| chr3 | 158698956 | 158699949 | L1MdTf_II | + | 993 |
| chr3 | 159090904 | 159096174 | L1MdTf_III | + | 5270 |
| chr3 | 159096174 | 159097167 | L1MdTf_III | + | 993 |
| chr3 | 159359875 | 159365317 | L1MdTf_III | + | 5442 |
| chr3 | 159365317 | 159366310 | L1MdTf_III | + | 993 |
| chr4 | 4268759 | 4274212 | L1MdTf_III | + | 5453 |
| chr4 | 4274212 | 4275227 | L1MdTf_III | + | 1015 |
| chr4 | 4608604 | 4608898 | L1MdTf_I | + | 294 |
| chr4 | 4608898 | 4609890 | L1MdTf_I | + | 992 |
| chr4 | 5927553 | 5932765 | L1MdTf_III | + | 5212 |
| chr4 | 5932765 | 5933182 | L1MdTf_III | + | 417 |
| chr4 | 7733688 | 7739145 | L1MdTf_III | + | 5457 |
| chr4 | 7739145 | 7740136 | L1MdTf_III | + | 991 |
| chr4 | 8191747 | 8196972 | L1MdTf_II | + | 5225 |
| chr4 | 8196972 | 8197965 | L1MdTf_II | + | 993 |
| chr4 | 8443045 | 8448571 | L1MdTf_III | + | 5526 |
| chr4 | 8448571 | 8449564 | L1MdTf_III | + | 993 |
| chr4 | 10109291 | 10109395 | L1MdTf_I | + | 104 |
| chr4 | 10109395 | 10110388 | L1MdTf_I | + | 993 |
| chr4 | 10491990 | 10497641 | L1MdTf_III | + | 5651 |
| chr4 | 10497641 | 10498628 | L1MdTf_III | + | 987 |
| chr4 | 12431153 | 12436765 | L1MdTf_II | + | 5612 |
| chr4 | 12436765 | 12437758 | L1MdTf_II | + | 993 |
| chr4 | 12467407 | 12473091 | L1MdTf_III | + | 5684 |
| chr4 | 12473091 | 12474086 | L1MdTf_III | + | 995 |
| chr4 | 14660982 | 14661686 | L1MdTf_III | + | 704 |
| chr4 | 14661686 | 14662679 | L1MdTf_III | + | 993 |
| chr4 | 15054443 | 15060106 | L1MdTf_II | + | 5663 |
| chr4 | 15060106 | 15061099 | L1MdTf_II | + | 993 |
| chr4 | 16058712 | 16063965 | L1MdTf_I | + | 5253 |
| chr4 | 16063965 | 16064958 | L1MdTf_I | + | 993 |
| chr4 | 16545274 | 16551098 | L1MdTf_II | + | 5824 |
| chr4 | 16551098 | 16552091 | L1MdTf_II | + | 993 |
| chr4 | 16706336 | 16707444 | L1MdTf_II | - | 1108 |
| chr4 | 16707444 | 16708283 | L1MdTf_II | + | 839 |
| chr4 | 17221332 | 17222927 | L1MdTf_II | + | 1595 |
| chr4 | 17222927 | 17223924 | L1MdTf_II | + | 997 |
| chr4 | 17408775 | 17414610 | L1MdTf_II | + | 5835 |
| chr4 | 17414610 | 17415620 | L1MdTf_II | + | 1010 |
| chr4 | 17571119 | 17576772 | L1MdTf_I | + | 5653 |

| | | | | | |
|---|---|---|---|---|---|
| chr4 | 17576772 | 17577765 | L1MdTf_I | + | 993 |
| chr4 | 17631088 | 17632017 | L1MdTf_I | + | 929 |
| chr4 | 17632017 | 17633010 | L1MdTf_I | + | 993 |
| chr4 | 17900873 | 17905070 | L1MdTf_III | + | 4197 |
| chr4 | 17905070 | 17906062 | L1MdTf_III | + | 992 |
| chr4 | 18021591 | 18027041 | L1MdTf_III | + | 5450 |
| chr4 | 18027042 | 18028034 | L1MdTf_III | + | 992 |
| chr4 | 19190076 | 19195632 | L1MdTf_I | + | 5556 |
| chr4 | 19195632 | 19196645 | L1MdTf_I | + | 1013 |
| chr4 | 19774576 | 19776488 | L1MdTf_II | - | 1912 |
| chr4 | 19776489 | 19776709 | L1MdTf_II | + | 220 |
| chr4 | 20420635 | 20426310 | L1MdTf_II | + | 5675 |
| chr4 | 20426310 | 20427327 | L1MdTf_II | + | 1017 |
| chr4 | 20868812 | 20870013 | L1MdTf_I | - | 1201 |
| chr4 | 20870013 | 20870253 | L1MdTf_II | + | 240 |
| chr4 | 21222954 | 21223139 | L1MdTf_II | - | 185 |
| chr4 | 21223139 | 21223496 | L1MdTf_II | + | 357 |
| chr4 | 21312410 | 21317640 | L1MdTf_III | + | 5230 |
| chr4 | 21317640 | 21318633 | L1MdTf_III | + | 993 |
| chr4 | 21374009 | 21379452 | L1MdTf_III | + | 5443 |
| chr4 | 21379452 | 21380445 | L1MdTf_III | + | 993 |
| chr4 | 21517395 | 21519168 | L1MdTf_III | + | 1773 |
| chr4 | 21519169 | 21520161 | L1MdTf_III | + | 992 |
| chr4 | 21650297 | 21656544 | L1MdTf_I | + | 6247 |
| chr4 | 21656544 | 21657537 | L1MdTf_I | + | 993 |
| chr4 | 22209814 | 22210314 | L1MdTf_III | - | 500 |
| chr4 | 22210314 | 22210594 | L1MdTf_III | + | 280 |
| chr4 | 22851465 | 22853987 | L1MdTf_II | + | 2522 |
| chr4 | 22853987 | 22854980 | L1MdTf_II | + | 993 |
| chr4 | 23946975 | 23947630 | L1MdTf_III | + | 655 |
| chr4 | 23947631 | 23948622 | L1MdTf_III | + | 991 |
| chr4 | 24117391 | 24119380 | L1MdTf_II | + | 1989 |
| chr4 | 24119380 | 24120374 | L1MdTf_II | + | 994 |
| chr4 | 24193703 | 24199986 | L1MdTf_II | + | 6283 |
| chr4 | 24199986 | 24200976 | L1MdTf_II | + | 990 |
| chr4 | 24295337 | 24300553 | L1MdTf_II | + | 5216 |
| chr4 | 24300553 | 24301545 | L1MdTf_II | + | 992 |
| chr4 | 25387912 | 25387955 | L1MdTf_III | - | 43 |
| chr4 | 25387955 | 25388231 | L1MdTf_III | + | 276 |
| chr4 | 25700552 | 25701664 | L1MdTf_III | + | 1112 |
| chr4 | 25701664 | 25702653 | L1MdTf_III | + | 989 |
| chr4 | 26109498 | 26114765 | L1MdTf_III | + | 5267 |
| chr4 | 26114765 | 26115758 | L1MdTf_III | + | 993 |
| chr4 | 27587754 | 27588384 | L1MdTf_I | - | 630 |
| chr4 | 27588384 | 27589373 | L1MdTf_I | + | 989 |

| | | | | | |
|---|---|---|---|---|---|
| chr4 | 28461503 | 28466877 | L1MdTf_I | + | 5374 |
| chr4 | 28466877 | 28467870 | L1MdTf_I | + | 993 |
| chr4 | 28746644 | 28751940 | L1MdTf_III | + | 5296 |
| chr4 | 28751941 | 28752932 | L1MdTf_III | + | 991 |
| chr4 | 30575882 | 30576853 | L1MdTf_I | - | 971 |
| chr4 | 30576853 | 30578981 | L1MdTf_I | + | 2128 |
| chr4 | 30630519 | 30632570 | L1MdTf_III | + | 2051 |
| chr4 | 30632570 | 30633563 | L1MdTf_III | + | 993 |
| chr4 | 30737964 | 30738851 | L1MdTf_II | + | 887 |
| chr4 | 30738851 | 30739844 | L1MdTf_II | + | 993 |
| chr4 | 31983876 | 31984092 | L1MdTf_III | + | 216 |
| chr4 | 31984092 | 31985084 | L1MdTf_III | + | 992 |
| chr4 | 35042621 | 35047836 | L1MdTf_I | + | 5215 |
| chr4 | 35047836 | 35048829 | L1MdTf_I | + | 993 |
| chr4 | 35181566 | 35186197 | L1MdTf_III | + | 4631 |
| chr4 | 35186197 | 35187190 | L1MdTf_III | + | 993 |
| chr4 | 35598192 | 35603578 | L1MdTf_III | + | 5386 |
| chr4 | 35603578 | 35604571 | L1MdTf_III | + | 993 |
| chr4 | 35627005 | 35632259 | L1MdTf_III | + | 5254 |
| chr4 | 35632259 | 35633252 | L1MdTf_III | + | 993 |
| chr4 | 37201621 | 37202285 | L1MdTf_II | + | 664 |
| chr4 | 37202285 | 37203296 | L1MdTf_II | + | 1011 |
| chr4 | 37343178 | 37348406 | L1MdTf_II | + | 5228 |
| chr4 | 37348406 | 37349399 | L1MdTf_II | + | 993 |
| chr4 | 37381741 | 37382421 | L1MdTf_III | - | 680 |
| chr4 | 37382421 | 37382544 | L1MdTf_III | + | 123 |
| chr4 | 37757432 | 37762899 | L1MdTf_III | + | 5467 |
| chr4 | 37762899 | 37763881 | L1MdTf_III | + | 982 |
| chr4 | 38549419 | 38554866 | L1MdTf_III | + | 5447 |
| chr4 | 38554866 | 38555859 | L1MdTf_III | + | 993 |
| chr4 | 39266359 | 39271648 | L1MdTf_III | + | 5289 |
| chr4 | 39271648 | 39272641 | L1MdTf_III | + | 993 |
| chr4 | 39430663 | 39431544 | L1MdTf_I | + | 881 |
| chr4 | 39431544 | 39432537 | L1MdTf_I | + | 993 |
| chr4 | 40083105 | 40088343 | L1MdTf_III | + | 5238 |
| chr4 | 40088343 | 40089336 | L1MdTf_III | + | 993 |
| chr4 | 40596337 | 40602843 | L1MdTf_I | + | 6506 |
| chr4 | 40602843 | 40603865 | L1MdTf_I | + | 1022 |
| chr4 | 43123380 | 43128631 | L1MdTf_I | + | 5251 |
| chr4 | 43128631 | 43129624 | L1MdTf_I | + | 993 |
| chr4 | 48246135 | 48248640 | L1MdTf_III | + | 2505 |
| chr4 | 48248640 | 48249658 | L1MdTf_III | + | 1018 |
| chr4 | 49265714 | 49271323 | L1MdTf_III | + | 5609 |
| chr4 | 49271323 | 49272314 | L1MdTf_III | + | 991 |
| chr4 | 50380331 | 50385939 | L1MdTf_II | + | 5608 |

| | | | | | |
|---|---|---|---|---|---|
| chr4 | 50385939 | 50386935 | L1MdTf_II | + | 996 |
| chr4 | 50534451 | 50534747 | L1MdTf_I | + | 296 |
| chr4 | 50534747 | 50535740 | L1MdTf_I | + | 993 |
| chr4 | 52063635 | 52068902 | L1MdTf_III | + | 5267 |
| chr4 | 52068902 | 52069895 | L1MdTf_III | + | 993 |
| chr4 | 52142154 | 52143534 | L1MdTf_I | + | 1380 |
| chr4 | 52143534 | 52144554 | L1MdTf_I | + | 1020 |
| chr4 | 53289647 | 53290949 | L1MdTf_III | + | 1302 |
| chr4 | 53290949 | 53291967 | L1MdTf_III | + | 1018 |
| chr4 | 53601819 | 53602153 | L1MdTf_II | + | 334 |
| chr4 | 53602153 | 53603150 | L1MdTf_II | + | 997 |
| chr4 | 53668403 | 53674903 | L1MdTf_I | + | 6500 |
| chr4 | 53674903 | 53675921 | L1MdTf_I | + | 1018 |
| chr4 | 58224805 | 58230036 | L1MdTf_II | + | 5231 |
| chr4 | 58230036 | 58231028 | L1MdTf_II | + | 992 |
| chr4 | 61394236 | 61395229 | L1MdTf_II | - | 993 |
| chr4 | 61395229 | 61395653 | L1MdTf_II | + | 424 |
| chr4 | 64372811 | 64379111 | L1MdTf_II | + | 6300 |
| chr4 | 64379111 | 64380104 | L1MdTf_II | + | 993 |
| chr4 | 64615336 | 64615812 | L1MdTf_II | - | 476 |
| chr4 | 64615812 | 64615916 | L1MdTf_II | + | 104 |
| chr4 | 65741534 | 65746767 | L1MdTf_I | + | 5233 |
| chr4 | 65746767 | 65747760 | L1MdTf_I | + | 993 |
| chr4 | 65996493 | 66000000 | L1MdTf_II | + | 3507 |
| chr4 | 66000000 | 66001952 | L1MdTf_II | + | 1952 |
| chr4 | 66273261 | 66279342 | L1MdTf_I | + | 6081 |
| chr4 | 66279342 | 66280361 | L1MdTf_I | + | 1019 |
| chr4 | 66636542 | 66641908 | L1MdTf_II | + | 5366 |
| chr4 | 66641908 | 66642927 | L1MdTf_II | + | 1019 |
| chr4 | 66693322 | 66698988 | L1MdTf_III | + | 5666 |
| chr4 | 66698989 | 66699982 | L1MdTf_III | + | 993 |
| chr4 | 67150771 | 67157145 | L1MdTf_III | + | 6374 |
| chr4 | 67157145 | 67158138 | L1MdTf_III | + | 993 |
| chr4 | 67738478 | 67738704 | L1MdTf_II | - | 226 |
| chr4 | 67738704 | 67738893 | L1MdTf_II | + | 189 |
| chr4 | 67834246 | 67840138 | L1MdTf_II | + | 5892 |
| chr4 | 67840138 | 67841122 | L1MdTf_II | + | 984 |
| chr4 | 67862538 | 67862999 | L1MdTf_II | + | 461 |
| chr4 | 67862999 | 67863992 | L1MdTf_II | + | 993 |
| chr4 | 69347028 | 69347129 | L1MdTf_III | - | 101 |
| chr4 | 69347129 | 69347771 | L1MdTf_III | - | 642 |
| chr4 | 69564637 | 69566953 | L1MdTf_I | + | 2316 |
| chr4 | 69566953 | 69567946 | L1MdTf_I | + | 993 |
| chr4 | 69625867 | 69631095 | L1MdTf_II | + | 5228 |
| chr4 | 69631095 | 69632090 | L1MdTf_II | + | 995 |

| | | | | | |
|---|---|---|---|---|---|
| chr4 | 69705124 | 69710510 | L1MdTf_III | + | 5386 |
| chr4 | 69710510 | 69711527 | L1MdTf_III | + | 1017 |
| chr4 | 69831720 | 69836816 | L1MdTf_III | + | 5096 |
| chr4 | 69836816 | 69837812 | L1MdTf_III | + | 996 |
| chr4 | 70765255 | 70766790 | L1MdTf_II | + | 1535 |
| chr4 | 70766790 | 70767786 | L1MdTf_II | + | 996 |
| chr4 | 71352488 | 71357964 | L1MdTf_III | + | 5476 |
| chr4 | 71357964 | 71358953 | L1MdTf_III | + | 989 |
| chr4 | 71745046 | 71746221 | L1MdTf_III | - | 1175 |
| chr4 | 71746221 | 71746538 | L1MdTf_III | + | 317 |
| chr4 | 72321563 | 72326847 | L1MdTf_III | + | 5284 |
| chr4 | 72326848 | 72327841 | L1MdTf_III | + | 993 |
| chr4 | 72597201 | 72603407 | L1MdTf_I | + | 6206 |
| chr4 | 72603407 | 72604400 | L1MdTf_I | + | 993 |
| chr4 | 73972201 | 73977654 | L1MdTf_I | + | 5453 |
| chr4 | 73977654 | 73978647 | L1MdTf_I | + | 993 |
| chr4 | 74782778 | 74788005 | L1MdTf_III | + | 5227 |
| chr4 | 74788005 | 74788998 | L1MdTf_III | + | 993 |
| chr4 | 75017161 | 75021858 | L1MdTf_I | + | 4697 |
| chr4 | 75021858 | 75022847 | L1MdTf_I | + | 989 |
| chr4 | 76646927 | 76652548 | L1MdTf_I | + | 5621 |
| chr4 | 76652548 | 76653541 | L1MdTf_I | + | 993 |
| chr4 | 76933545 | 76933614 | L1MdTf_III | + | 69 |
| chr4 | 76933614 | 76934607 | L1MdTf_III | + | 993 |
| chr4 | 76980956 | 76986585 | L1MdTf_II | + | 5629 |
| chr4 | 76986585 | 76987578 | L1MdTf_II | + | 993 |
| chr4 | 78598729 | 78599315 | L1MdTf_II | - | 586 |
| chr4 | 78599315 | 78599808 | L1MdTf_I | + | 493 |
| chr4 | 78798211 | 78798975 | L1MdTf_III | - | 764 |
| chr4 | 78798975 | 78800130 | L1MdTf_III | + | 1155 |
| chr4 | 79061023 | 79066691 | L1MdTf_II | + | 5668 |
| chr4 | 79066691 | 79067675 | L1MdTf_II | + | 984 |
| chr4 | 79175485 | 79180731 | L1MdTf_III | + | 5246 |
| chr4 | 79180731 | 79181722 | L1MdTf_III | + | 991 |
| chr4 | 79491486 | 79494976 | L1MdTf_III | + | 3490 |
| chr4 | 79494976 | 79495969 | L1MdTf_III | + | 993 |
| chr4 | 80320913 | 80320983 | L1MdTf_III | + | 70 |
| chr4 | 80320983 | 80321487 | L1MdTf_III | + | 504 |
| chr4 | 80617045 | 80618027 | L1MdTf_III | + | 982 |
| chr4 | 80618027 | 80619020 | L1MdTf_III | + | 993 |
| chr4 | 80731254 | 80736467 | L1MdTf_I | + | 5213 |
| chr4 | 80736467 | 80737488 | L1MdTf_I | + | 1021 |
| chr4 | 81323654 | 81328971 | L1MdTf_III | + | 5317 |
| chr4 | 81328971 | 81329976 | L1MdTf_III | + | 1005 |
| chr4 | 81441190 | 81441424 | L1MdTf_II | - | 234 |

| | | | | | |
|---|---|---|---|---|---|
| chr4 | 81441424 | 81441528 | L1MdTf_II | + | 104 |
| chr4 | 81959411 | 81965262 | L1MdTf_I | + | 5851 |
| chr4 | 81965262 | 81966285 | L1MdTf_I | + | 1023 |
| chr4 | 83876997 | 83882888 | L1MdTf_II | + | 5891 |
| chr4 | 83882888 | 83883902 | L1MdTf_II | + | 1014 |
| chr4 | 85071603 | 85071726 | L1MdTf_III | - | 123 |
| chr4 | 85071726 | 85071861 | L1MdTf_III | + | 135 |
| chr4 | 85809876 | 85815325 | L1MdTf_III | + | 5449 |
| chr4 | 85815325 | 85816318 | L1MdTf_III | + | 993 |
| chr4 | 86387361 | 86392615 | L1MdTf_II | + | 5254 |
| chr4 | 86392615 | 86393637 | L1MdTf_II | + | 1022 |
| chr4 | 86393637 | 86393777 | L1MdTf_III | + | 140 |
| chr4 | 86393777 | 86394770 | L1MdTf_III | + | 993 |
| chr4 | 88279089 | 88284655 | L1MdTf_I | + | 5566 |
| chr4 | 88284655 | 88285673 | L1MdTf_I | + | 1018 |
| chr4 | 88294439 | 88300419 | L1MdTf_II | + | 5980 |
| chr4 | 88300419 | 88301412 | L1MdTf_II | + | 993 |
| chr4 | 88586535 | 88591985 | L1MdTf_II | + | 5450 |
| chr4 | 88591985 | 88592978 | L1MdTf_II | + | 993 |
| chr4 | 89680882 | 89686546 | L1MdTf_II | + | 5664 |
| chr4 | 89686546 | 89687542 | L1MdTf_II | + | 996 |
| chr4 | 89920184 | 89925590 | L1MdTf_III | + | 5406 |
| chr4 | 89925590 | 89926580 | L1MdTf_III | + | 990 |
| chr4 | 90546980 | 90552227 | L1MdTf_III | + | 5247 |
| chr4 | 90552227 | 90553220 | L1MdTf_III | + | 993 |
| chr4 | 90867427 | 90873960 | L1MdTf_I | + | 6533 |
| chr4 | 90873960 | 90874981 | L1MdTf_I | + | 1021 |
| chr4 | 91058909 | 91062681 | L1MdTf_III | + | 3772 |
| chr4 | 91062682 | 91063675 | L1MdTf_III | + | 993 |
| chr4 | 91542854 | 91548958 | L1MdTf_I | + | 6104 |
| chr4 | 91548958 | 91549951 | L1MdTf_I | + | 993 |
| chr4 | 92699121 | 92704575 | L1MdTf_III | + | 5454 |
| chr4 | 92704575 | 92705566 | L1MdTf_III | + | 991 |
| chr4 | 92873871 | 92880007 | L1MdTf_II | + | 6136 |
| chr4 | 92880007 | 92880997 | L1MdTf_II | + | 990 |
| chr4 | 93591967 | 93597393 | L1MdTf_II | + | 5426 |
| chr4 | 93597393 | 93598386 | L1MdTf_II | + | 993 |
| chr4 | 93777422 | 93782662 | L1MdTf_III | + | 5240 |
| chr4 | 93782662 | 93783655 | L1MdTf_III | + | 993 |
| chr4 | 94095247 | 94095580 | L1MdTf_II | + | 333 |
| chr4 | 94095580 | 94096564 | L1MdTf_II | + | 984 |
| chr4 | 94779406 | 94785408 | L1MdTf_I | + | 6002 |
| chr4 | 94785408 | 94786401 | L1MdTf_I | + | 993 |
| chr4 | 94800855 | 94806832 | L1MdTf_I | + | 5977 |
| chr4 | 94806832 | 94807825 | L1MdTf_I | + | 993 |

| | | | | | |
|---|---|---|---|---|---|
| chr4 | 99619403 | 99625042 | L1MdTf_I | + | 5639 |
| chr4 | 99625042 | 99626035 | L1MdTf_I | + | 993 |
| chr4 | 102552871 | 102553620 | L1MdTf_III | - | 749 |
| chr4 | 102553620 | 102554269 | L1MdTf_III | + | 649 |
| chr4 | 103130513 | 103135935 | L1MdTf_I | + | 5422 |
| chr4 | 103135935 | 103136928 | L1MdTf_I | + | 993 |
| chr4 | 103946914 | 103952554 | L1MdTf_I | + | 5640 |
| chr4 | 103952554 | 103953547 | L1MdTf_I | + | 993 |
| chr4 | 105451173 | 105456521 | L1MdTf_III | + | 5348 |
| chr4 | 105456521 | 105457514 | L1MdTf_III | + | 993 |
| chr4 | 105492379 | 105493282 | L1MdTf_I | + | 903 |
| chr4 | 105493282 | 105494275 | L1MdTf_I | + | 993 |
| chr4 | 105592590 | 105598229 | L1MdTf_I | + | 5639 |
| chr4 | 105598229 | 105599222 | L1MdTf_I | + | 993 |
| chr4 | 108557758 | 108563286 | L1MdTf_II | + | 5528 |
| chr4 | 108563286 | 108564291 | L1MdTf_II | + | 1005 |
| chr4 | 110043902 | 110049591 | L1MdTf_III | + | 5689 |
| chr4 | 110049592 | 110050585 | L1MdTf_III | + | 993 |
| chr4 | 110603482 | 110604136 | L1MdTf_III | - | 654 |
| chr4 | 110604136 | 110604793 | L1MdTf_III | + | 657 |
| chr4 | 111039124 | 111042093 | L1MdTf_I | + | 2969 |
| chr4 | 111042093 | 111043086 | L1MdTf_I | + | 993 |
| chr4 | 111063663 | 111064917 | L1MdTf_I | + | 1254 |
| chr4 | 111064917 | 111065932 | L1MdTf_I | + | 1015 |
| chr4 | 111835744 | 111836466 | L1MdTf_II | - | 722 |
| chr4 | 111836466 | 111837133 | L1MdTf_II | + | 667 |
| chr4 | 112839022 | 112844275 | L1MdTf_II | + | 5253 |
| chr4 | 112844275 | 112845269 | L1MdTf_II | + | 994 |
| chr4 | 113293448 | 113293745 | L1MdTf_I | + | 297 |
| chr4 | 113293745 | 113294738 | L1MdTf_I | + | 993 |
| chr4 | 115194445 | 115199829 | L1MdTf_II | + | 5384 |
| chr4 | 115199829 | 115200826 | L1MdTf_II | + | 997 |
| chr4 | 115527854 | 115529444 | L1MdTf_II | + | 1590 |
| chr4 | 115529444 | 115530437 | L1MdTf_II | + | 993 |
| chr4 | 118703791 | 118709062 | L1MdTf_III | + | 5271 |
| chr4 | 118709062 | 118709493 | L1MdTf_III | + | 431 |
| chr4 | 122583926 | 122584113 | L1MdTf_III | + | 187 |
| chr4 | 122584113 | 122585106 | L1MdTf_III | + | 993 |
| chr4 | 143955891 | 143961219 | L1MdTf_III | + | 5328 |
| chr4 | 143961219 | 143962234 | L1MdTf_III | + | 1015 |
| chr4 | 144543195 | 144548791 | L1MdTf_III | + | 5596 |
| chr4 | 144548791 | 144549784 | L1MdTf_III | + | 993 |
| chr5 | 4651278 | 4656695 | L1MdTf_II | + | 5417 |
| chr5 | 4656695 | 4657706 | L1MdTf_II | + | 1011 |
| chr5 | 5453403 | 5458589 | L1MdTf_III | + | 5186 |

| | | | | | |
|---|---|---|---|---|---|
| chr5 | 5458590 | 5459582 | L1MdTf_III | + | 992 |
| chr5 | 6281988 | 6282126 | L1MdTf_III | + | 138 |
| chr5 | 6282126 | 6283120 | L1MdTf_III | + | 994 |
| chr5 | 6924640 | 6924774 | L1MdTf_II | + | 134 |
| chr5 | 6924774 | 6925786 | L1MdTf_II | + | 1012 |
| chr5 | 7135469 | 7141049 | L1MdTf_II | + | 5580 |
| chr5 | 7141049 | 7142042 | L1MdTf_II | + | 993 |
| chr5 | 8219166 | 8224491 | L1MdTf_III | + | 5325 |
| chr5 | 8224491 | 8225484 | L1MdTf_III | + | 993 |
| chr5 | 8316588 | 8321799 | L1MdTf_I | + | 5211 |
| chr5 | 8321799 | 8322811 | L1MdTf_I | + | 1012 |
| chr5 | 8601606 | 8606818 | L1MdTf_II | + | 5212 |
| chr5 | 8606818 | 8607811 | L1MdTf_II | + | 993 |
| chr5 | 9099288 | 9104929 | L1MdTf_II | + | 5641 |
| chr5 | 9104929 | 9105922 | L1MdTf_II | + | 993 |
| chr5 | 9380402 | 9385777 | L1MdTf_III | + | 5375 |
| chr5 | 9385777 | 9386770 | L1MdTf_III | + | 993 |
| chr5 | 9864578 | 9869807 | L1MdTf_III | + | 5229 |
| chr5 | 9869807 | 9870796 | L1MdTf_III | + | 989 |
| chr5 | 10339256 | 10343115 | L1MdTf_II | + | 3859 |
| chr5 | 10343115 | 10344104 | L1MdTf_II | + | 989 |
| chr5 | 10502107 | 10507337 | L1MdTf_III | + | 5230 |
| chr5 | 10507337 | 10508330 | L1MdTf_III | + | 993 |
| chr5 | 11267524 | 11268177 | L1MdTf_III | + | 653 |
| chr5 | 11268177 | 11269170 | L1MdTf_III | + | 993 |
| chr5 | 11419168 | 11419850 | L1MdTf_I | + | 682 |
| chr5 | 11419850 | 11420843 | L1MdTf_I | + | 993 |
| chr5 | 11848196 | 11848878 | L1MdTf_I | + | 682 |
| chr5 | 11848878 | 11849866 | L1MdTf_I | + | 988 |
| chr5 | 12864049 | 12865852 | L1MdTf_II | - | 1803 |
| chr5 | 12865852 | 12867473 | L1MdTf_II | + | 1621 |
| chr5 | 13934038 | 13935680 | L1MdTf_II | + | 1642 |
| chr5 | 13935680 | 13936677 | L1MdTf_II | + | 997 |
| chr5 | 15014127 | 15019476 | L1MdTf_III | + | 5349 |
| chr5 | 15019476 | 15020469 | L1MdTf_III | + | 993 |
| chr5 | 15930089 | 15930579 | L1MdTf_III | + | 490 |
| chr5 | 15930579 | 15931591 | L1MdTf_III | + | 1012 |
| chr5 | 16018221 | 16018564 | L1MdTf_III | + | 343 |
| chr5 | 16018564 | 16019557 | L1MdTf_III | + | 993 |
| chr5 | 17108413 | 17114483 | L1MdTf_II | + | 6070 |
| chr5 | 17114483 | 17115476 | L1MdTf_II | + | 993 |
| chr5 | 17145866 | 17151348 | L1MdTf_I | + | 5482 |
| chr5 | 17151348 | 17152341 | L1MdTf_I | + | 993 |
| chr5 | 17813445 | 17819718 | L1MdTf_I | + | 6273 |
| chr5 | 17819718 | 17820711 | L1MdTf_I | + | 993 |

| | | | | | |
|---|---|---|---|---|---|
| chr5 | 18004089 | 18010007 | L1MdTf_II | + | 5918 |
| chr5 | 18010007 | 18010982 | L1MdTf_II | + | 975 |
| chr5 | 19057355 | 19063049 | L1MdTf_III | + | 5694 |
| chr5 | 19063050 | 19064041 | L1MdTf_III | + | 991 |
| chr5 | 19681581 | 19686808 | L1MdTf_III | + | 5227 |
| chr5 | 19686808 | 19687801 | L1MdTf_III | + | 993 |
| chr5 | 22569470 | 22573402 | L1MdTf_III | + | 3932 |
| chr5 | 22573403 | 22574395 | L1MdTf_III | + | 992 |
| chr5 | 22661297 | 22666871 | L1MdTf_III | + | 5574 |
| chr5 | 22666871 | 22667866 | L1MdTf_III | + | 995 |
| chr5 | 22934806 | 22935253 | L1MdTf_III | + | 447 |
| chr5 | 22935253 | 22936248 | L1MdTf_III | + | 995 |
| chr5 | 23230002 | 23235619 | L1MdTf_III | + | 5617 |
| chr5 | 23235619 | 23236612 | L1MdTf_III | + | 993 |
| chr5 | 26584419 | 26590695 | L1MdTf_I | + | 6276 |
| chr5 | 26590695 | 26591688 | L1MdTf_I | + | 993 |
| chr5 | 26597506 | 26598580 | L1MdTf_III | + | 1074 |
| chr5 | 26598580 | 26599573 | L1MdTf_III | + | 993 |
| chr5 | 27501138 | 27506565 | L1MdTf_I | + | 5427 |
| chr5 | 27506565 | 27507558 | L1MdTf_I | + | 993 |
| chr5 | 28319858 | 28320200 | L1MdTf_I | + | 342 |
| chr5 | 28320200 | 28321193 | L1MdTf_I | + | 993 |
| chr5 | 28690017 | 28695735 | L1MdTf_II | + | 5718 |
| chr5 | 28695735 | 28696731 | L1MdTf_II | + | 996 |
| chr5 | 28809078 | 28814844 | L1MdTf_II | + | 5766 |
| chr5 | 28814844 | 28815837 | L1MdTf_II | + | 993 |
| chr5 | 29381735 | 29387374 | L1MdTf_I | + | 5639 |
| chr5 | 29387374 | 29388367 | L1MdTf_I | + | 993 |
| chr5 | 29600933 | 29601368 | L1MdTf_III | + | 435 |
| chr5 | 29601368 | 29602358 | L1MdTf_III | + | 990 |
| chr5 | 32145964 | 32151135 | L1MdTf_I | + | 5171 |
| chr5 | 32151135 | 32152152 | L1MdTf_I | + | 1017 |
| chr5 | 32593248 | 32593788 | L1MdTf_II | + | 540 |
| chr5 | 32593788 | 32594778 | L1MdTf_II | + | 990 |
| chr5 | 34419503 | 34421188 | L1MdTf_II | - | 1685 |
| chr5 | 34421189 | 34422883 | L1MdTf_II | + | 1694 |
| chr5 | 36395121 | 36402267 | L1MdTf_I | + | 7146 |
| chr5 | 36402267 | 36403257 | L1MdTf_I | + | 990 |
| chr5 | 37656626 | 37662119 | L1MdTf_III | + | 5493 |
| chr5 | 37662119 | 37663112 | L1MdTf_III | + | 993 |
| chr5 | 39733979 | 39739324 | L1MdTf_III | + | 5345 |
| chr5 | 39739324 | 39740304 | L1MdTf_III | + | 980 |
| chr5 | 41076070 | 41076200 | L1MdTf_II | - | 130 |
| chr5 | 41076200 | 41076428 | L1MdTf_II | + | 228 |
| chr5 | 41817099 | 41817441 | L1MdTf_III | + | 342 |

| | | | | | |
|---|---|---|---|---|---|
| chr5 | 41817441 | 41818434 | L1MdTf_III | + | 993 |
| chr5 | 42293495 | 42299405 | L1MdTf_II | + | 5910 |
| chr5 | 42299405 | 42300415 | L1MdTf_II | + | 1010 |
| chr5 | 42345633 | 42351660 | L1MdTf_II | + | 6027 |
| chr5 | 42351660 | 42352653 | L1MdTf_II | + | 993 |
| chr5 | 43255107 | 43258143 | L1MdTf_I | + | 3036 |
| chr5 | 43258143 | 43259136 | L1MdTf_I | + | 993 |
| chr5 | 45760989 | 45764922 | L1MdTf_III | - | 3933 |
| chr5 | 45764922 | 45765891 | L1MdTf_III | + | 969 |
| chr5 | 46142409 | 46142900 | L1MdTf_III | + | 491 |
| chr5 | 46142900 | 46143895 | L1MdTf_III | + | 995 |
| chr5 | 47863557 | 47864311 | L1MdTf_III | - | 754 |
| chr5 | 47864311 | 47864667 | L1MdTf_III | + | 356 |
| chr5 | 48584210 | 48584571 | L1MdTf_III | + | 361 |
| chr5 | 48584571 | 48585563 | L1MdTf_III | + | 992 |
| chr5 | 48681511 | 48685445 | L1MdTf_III | + | 3934 |
| chr5 | 48685445 | 48686438 | L1MdTf_III | + | 993 |
| chr5 | 48812567 | 48813157 | L1MdTf_III | + | 590 |
| chr5 | 48813157 | 48814148 | L1MdTf_III | + | 991 |
| chr5 | 54293509 | 54299007 | L1MdTf_III | + | 5498 |
| chr5 | 54299007 | 54300001 | L1MdTf_III | + | 994 |
| chr5 | 55336497 | 55337369 | L1MdTf_III | + | 872 |
| chr5 | 55337369 | 55338361 | L1MdTf_III | + | 992 |
| chr5 | 56346374 | 56352182 | L1MdTf_II | + | 5808 |
| chr5 | 56352182 | 56353198 | L1MdTf_II | + | 1016 |
| chr5 | 56559107 | 56560585 | L1MdTf_I | + | 1478 |
| chr5 | 56560585 | 56561577 | L1MdTf_I | + | 992 |
| chr5 | 56628847 | 56634714 | L1MdTf_I | + | 5867 |
| chr5 | 56634714 | 56635707 | L1MdTf_I | + | 993 |
| chr5 | 56650510 | 56655942 | L1MdTf_I | + | 5432 |
| chr5 | 56655942 | 56656953 | L1MdTf_I | + | 1011 |
| chr5 | 57810526 | 57814188 | L1MdTf_II | + | 3662 |
| chr5 | 57814188 | 57815181 | L1MdTf_II | + | 993 |
| chr5 | 58402704 | 58407953 | L1MdTf_III | + | 5249 |
| chr5 | 58407953 | 58408947 | L1MdTf_III | + | 994 |
| chr5 | 58647098 | 58647149 | L1MdTf_III | - | 51 |
| chr5 | 58647149 | 58647529 | L1MdTf_III | + | 380 |
| chr5 | 58887331 | 58888921 | L1MdTf_I | + | 1590 |
| chr5 | 58888921 | 58889941 | L1MdTf_I | + | 1020 |
| chr5 | 58935039 | 58940681 | L1MdTf_II | + | 5642 |
| chr5 | 58940681 | 58941674 | L1MdTf_II | + | 993 |
| chr5 | 60226973 | 60227805 | L1MdTf_II | - | 832 |
| chr5 | 60227805 | 60228720 | L1MdTf_II | + | 915 |
| chr5 | 60503321 | 60508641 | L1MdTf_III | + | 5320 |
| chr5 | 60508641 | 60509634 | L1MdTf_III | + | 993 |

| | | | | | |
|---|---|---|---|---|---|
| chr5 | 60627841 | 60630127 | L1MdTf_II | + | 2286 |
| chr5 | 60630127 | 60631120 | L1MdTf_II | + | 993 |
| chr5 | 60809023 | 60814579 | L1MdTf_II | + | 5556 |
| chr5 | 60814579 | 60815592 | L1MdTf_II | + | 1013 |
| chr5 | 61140994 | 61142494 | L1MdTf_III | + | 1500 |
| chr5 | 61142494 | 61143487 | L1MdTf_III | + | 993 |
| chr5 | 61370288 | 61375713 | L1MdTf_I | + | 5425 |
| chr5 | 61375713 | 61376706 | L1MdTf_I | + | 993 |
| chr5 | 61389639 | 61395108 | L1MdTf_III | + | 5469 |
| chr5 | 61395108 | 61396093 | L1MdTf_III | + | 985 |
| chr5 | 61450143 | 61456183 | L1MdTf_II | + | 6040 |
| chr5 | 61456183 | 61457180 | L1MdTf_II | + | 997 |
| chr5 | 61935637 | 61941504 | L1MdTf_II | + | 5867 |
| chr5 | 61941504 | 61942497 | L1MdTf_II | + | 993 |
| chr5 | 62366639 | 62372046 | L1MdTf_I | + | 5407 |
| chr5 | 62372046 | 62373039 | L1MdTf_I | + | 993 |
| chr5 | 62584856 | 62590321 | L1MdTf_I | + | 5465 |
| chr5 | 62590321 | 62591314 | L1MdTf_I | + | 993 |
| chr5 | 63525850 | 63531046 | L1MdTf_III | + | 5196 |
| chr5 | 63531046 | 63532028 | L1MdTf_III | + | 982 |
| chr5 | 63814720 | 63820143 | L1MdTf_II | + | 5423 |
| chr5 | 63820143 | 63821136 | L1MdTf_II | + | 993 |
| chr5 | 64275070 | 64280709 | L1MdTf_I | + | 5639 |
| chr5 | 64280709 | 64281702 | L1MdTf_I | + | 993 |
| chr5 | 68922578 | 68923645 | L1MdTf_I | + | 1067 |
| chr5 | 68923645 | 68924659 | L1MdTf_I | + | 1014 |
| chr5 | 69688821 | 69694723 | L1MdTf_III | + | 5902 |
| chr5 | 69694723 | 69695716 | L1MdTf_III | + | 993 |
| chr5 | 69886467 | 69891114 | L1MdTf_II | - | 4647 |
| chr5 | 69891114 | 69891367 | L1MdTf_II | + | 253 |
| chr5 | 69993156 | 69993447 | L1MdTf_I | - | 291 |
| chr5 | 69993447 | 69993505 | L1MdTf_II | + | 58 |
| chr5 | 70040146 | 70040928 | L1MdTf_I | + | 782 |
| chr5 | 70040928 | 70041922 | L1MdTf_I | + | 994 |
| chr5 | 70228399 | 70234236 | L1MdTf_I | + | 5837 |
| chr5 | 70234236 | 70235224 | L1MdTf_I | + | 988 |
| chr5 | 71713302 | 71719619 | L1MdTf_II | + | 6317 |
| chr5 | 71719619 | 71720612 | L1MdTf_II | + | 993 |
| chr5 | 71989180 | 71994835 | L1MdTf_I | + | 5655 |
| chr5 | 71994835 | 71995850 | L1MdTf_I | + | 1015 |
| chr5 | 72699695 | 72704804 | L1MdTf_II | + | 5109 |
| chr5 | 72704804 | 72705797 | L1MdTf_II | + | 993 |
| chr5 | 73091497 | 73093473 | L1MdTf_II | + | 1976 |
| chr5 | 73093473 | 73094462 | L1MdTf_II | + | 989 |
| chr5 | 73315454 | 73315536 | L1MdTf_I | - | 82 |

| | | | | | |
|---|---|---|---|---|---|
| chr5 | 73315536 | 73315732 | L1MdTf_II | + | 196 |
| chr5 | 74124103 | 74125181 | L1MdTf_III | + | 1078 |
| chr5 | 74125181 | 74126176 | L1MdTf_III | + | 995 |
| chr5 | 75301671 | 75301789 | L1MdTf_III | + | 118 |
| chr5 | 75301789 | 75302001 | L1MdTf_III | + | 212 |
| chr5 | 76651966 | 76657230 | L1MdTf_III | + | 5264 |
| chr5 | 76657230 | 76658217 | L1MdTf_III | + | 987 |
| chr5 | 77557486 | 77562891 | L1MdTf_I | + | 5405 |
| chr5 | 77562891 | 77563884 | L1MdTf_I | + | 993 |
| chr5 | 77778453 | 77779972 | L1MdTf_I | + | 1519 |
| chr5 | 77779972 | 77780955 | L1MdTf_I | + | 983 |
| chr5 | 77843246 | 77848611 | L1MdTf_III | + | 5365 |
| chr5 | 77848611 | 77849604 | L1MdTf_III | + | 993 |
| chr5 | 78560488 | 78565730 | L1MdTf_III | + | 5242 |
| chr5 | 78565730 | 78566723 | L1MdTf_III | + | 993 |
| chr5 | 78771090 | 78771743 | L1MdTf_III | - | 653 |
| chr5 | 78771743 | 78772123 | L1MdTf_III | + | 380 |
| chr5 | 79180718 | 79186146 | L1MdTf_II | + | 5428 |
| chr5 | 79186146 | 79187139 | L1MdTf_II | + | 993 |
| chr5 | 79387854 | 79388228 | L1MdTf_III | + | 374 |
| chr5 | 79388228 | 79389221 | L1MdTf_III | + | 993 |
| chr5 | 79995936 | 80000000 | L1MdTf_II | + | 4064 |
| chr5 | 80000000 | 80001803 | L1MdTf_II | + | 1803 |
| chr5 | 80257310 | 80259490 | L1MdTf_III | + | 2180 |
| chr5 | 80259490 | 80260483 | L1MdTf_III | + | 993 |
| chr5 | 81352973 | 81358437 | L1MdTf_I | + | 5464 |
| chr5 | 81358437 | 81359430 | L1MdTf_I | + | 993 |
| chr5 | 83054710 | 83060410 | L1MdTf_I | + | 5700 |
| chr5 | 83060410 | 83061403 | L1MdTf_I | + | 993 |
| chr5 | 83533165 | 83537573 | L1MdTf_II | - | 4408 |
| chr5 | 83537573 | 83537798 | L1MdTf_II | + | 225 |
| chr5 | 83931371 | 83937035 | L1MdTf_II | + | 5664 |
| chr5 | 83937035 | 83938028 | L1MdTf_II | + | 993 |
| chr5 | 84901651 | 84907140 | L1MdTf_III | + | 5489 |
| chr5 | 84907140 | 84908133 | L1MdTf_III | + | 993 |
| chr5 | 85411731 | 85417115 | L1MdTf_I | + | 5384 |
| chr5 | 85417115 | 85418131 | L1MdTf_I | + | 1016 |
| chr5 | 85665117 | 85668831 | L1MdTf_III | + | 3714 |
| chr5 | 85668831 | 85669820 | L1MdTf_III | + | 989 |
| chr5 | 86598187 | 86603438 | L1MdTf_III | + | 5251 |
| chr5 | 86603438 | 86604429 | L1MdTf_III | + | 991 |
| chr5 | 86741993 | 86747655 | L1MdTf_II | + | 5662 |
| chr5 | 86747655 | 86748648 | L1MdTf_II | + | 993 |
| chr5 | 86833591 | 86838761 | L1MdTf_III | + | 5170 |
| chr5 | 86838761 | 86839754 | L1MdTf_III | + | 993 |

| | | | | | |
|---|---|---|---|---|---|
| chr5 | 87245740 | 87246426 | L1MdTf_I | - | 686 |
| chr5 | 87246426 | 87248128 | L1MdTf_I | - | 1702 |
| chr5 | 88407144 | 88412398 | L1MdTf_III | + | 5254 |
| chr5 | 88412398 | 88413420 | L1MdTf_III | + | 1022 |
| chr5 | 88693741 | 88700036 | L1MdTf_I | + | 6295 |
| chr5 | 88700036 | 88701029 | L1MdTf_I | + | 993 |
| chr5 | 89475757 | 89481621 | L1MdTf_III | + | 5864 |
| chr5 | 89481622 | 89482615 | L1MdTf_III | + | 993 |
| chr5 | 89482615 | 89482913 | L1MdTf_III | + | 298 |
| chr5 | 89482914 | 89483907 | L1MdTf_III | + | 993 |
| chr5 | 89950282 | 89950949 | L1MdTf_I | - | 667 |
| chr5 | 89950949 | 89951815 | L1MdTf_I | + | 866 |
| chr5 | 91535653 | 91536554 | L1MdTf_II | + | 901 |
| chr5 | 91536554 | 91537573 | L1MdTf_II | + | 1019 |
| chr5 | 91979958 | 91985211 | L1MdTf_III | + | 5253 |
| chr5 | 91985211 | 91986204 | L1MdTf_III | + | 993 |
| chr5 | 93952154 | 93957591 | L1MdTf_II | + | 5437 |
| chr5 | 93957591 | 93958602 | L1MdTf_II | + | 1011 |
| chr5 | 95633559 | 95634449 | L1MdTf_III | - | 890 |
| chr5 | 95634449 | 95634565 | L1MdTf_III | + | 116 |
| chr5 | 98136584 | 98142688 | L1MdTf_III | + | 6104 |
| chr5 | 98142688 | 98143681 | L1MdTf_III | + | 993 |
| chr5 | 98156124 | 98159059 | L1MdTf_II | + | 2935 |
| chr5 | 98159059 | 98160052 | L1MdTf_II | + | 993 |
| chr5 | 99260367 | 99262028 | L1MdTf_II | - | 1661 |
| chr5 | 99262028 | 99265660 | L1MdTf_II | + | 3632 |
| chr5 | 101460210 | 101465683 | L1MdTf_III | + | 5473 |
| chr5 | 101465683 | 101466676 | L1MdTf_III | + | 993 |
| chr5 | 102556669 | 102562945 | L1MdTf_II | + | 6276 |
| chr5 | 102562945 | 102563957 | L1MdTf_II | + | 1012 |
| chr5 | 105169959 | 105175381 | L1MdTf_I | + | 5422 |
| chr5 | 105175382 | 105176374 | L1MdTf_I | + | 992 |
| chr5 | 105594817 | 105595247 | L1MdTf_III | - | 430 |
| chr5 | 105595247 | 105596038 | L1MdTf_III | + | 791 |
| chr5 | 105988509 | 105990335 | L1MdTf_I | + | 1826 |
| chr5 | 105990335 | 105991350 | L1MdTf_I | + | 1015 |
| chr5 | 109173446 | 109178672 | L1MdTf_III | + | 5226 |
| chr5 | 109178672 | 109179667 | L1MdTf_III | + | 995 |
| chr5 | 109270502 | 109276361 | L1MdTf_II | + | 5859 |
| chr5 | 109276361 | 109277382 | L1MdTf_II | + | 1021 |
| chr5 | 126453718 | 126453847 | L1MdTf_I | + | 129 |
| chr5 | 126453847 | 126454840 | L1MdTf_II | + | 993 |
| chr5 | 128095211 | 128095359 | L1MdTf_I | - | 148 |
| chr5 | 128095359 | 128095923 | L1MdTf_I | + | 564 |
| chr5 | 129715752 | 129716141 | L1MdTf_III | - | 389 |

| | | | | | |
|---|---|---|---|---|---|
| chr5 | 129716141 | 129716990 | L1MdTf_III | + | 849 |
| chr5 | 132758269 | 132763493 | L1MdTf_II | + | 5224 |
| chr5 | 132763493 | 132764486 | L1MdTf_II | + | 993 |
| chr5 | 133588685 | 133594341 | L1MdTf_II | + | 5656 |
| chr5 | 133594341 | 133595338 | L1MdTf_II | + | 997 |
| chr5 | 138565975 | 138566565 | L1MdTf_II | - | 590 |
| chr5 | 138566565 | 138567286 | L1MdTf_II | + | 721 |
| chr5 | 145973993 | 145979635 | L1MdTf_II | + | 5642 |
| chr5 | 145979635 | 145980625 | L1MdTf_II | + | 990 |
| chr5 | 146065811 | 146066007 | L1MdTf_I | + | 196 |
| chr5 | 146066007 | 146067000 | L1MdTf_I | + | 993 |
| chr5 | 151277101 | 151282540 | L1MdTf_III | + | 5439 |
| chr5 | 151282540 | 151283534 | L1MdTf_III | + | 994 |
| chr5 | 151578579 | 151584673 | L1MdTf_III | + | 6094 |
| chr5 | 151584673 | 151585697 | L1MdTf_III | + | 1024 |
| chr6 | 4092092 | 4097757 | L1MdTf_III | + | 5665 |
| chr6 | 4097758 | 4098755 | L1MdTf_III | + | 997 |
| chr6 | 4216953 | 4217285 | L1MdTf_II | - | 332 |
| chr6 | 4217285 | 4217362 | L1MdTf_II | + | 77 |
| chr6 | 4282293 | 4287534 | L1MdTf_II | + | 5241 |
| chr6 | 4287534 | 4288527 | L1MdTf_II | + | 993 |
| chr6 | 4329354 | 4329744 | L1MdTf_III | + | 390 |
| chr6 | 4329744 | 4330729 | L1MdTf_III | + | 985 |
| chr6 | 6748332 | 6748547 | L1MdTf_II | + | 215 |
| chr6 | 6748547 | 6749540 | L1MdTf_II | + | 993 |
| chr6 | 7225485 | 7226097 | L1MdTf_II | - | 612 |
| chr6 | 7226097 | 7226991 | L1MdTf_II | + | 894 |
| chr6 | 7663056 | 7664924 | L1MdTf_II | + | 1868 |
| chr6 | 7664924 | 7665917 | L1MdTf_II | + | 993 |
| chr6 | 9603654 | 9609189 | L1MdTf_I | + | 5535 |
| chr6 | 9609190 | 9610182 | L1MdTf_I | + | 992 |
| chr6 | 10147001 | 10152439 | L1MdTf_II | + | 5438 |
| chr6 | 10152439 | 10153432 | L1MdTf_II | + | 993 |
| chr6 | 11299837 | 11300262 | L1MdTf_I | + | 425 |
| chr6 | 11300262 | 11301278 | L1MdTf_I | + | 1016 |
| chr6 | 12522579 | 12527865 | L1MdTf_II | + | 5286 |
| chr6 | 12527865 | 12528859 | L1MdTf_II | + | 994 |
| chr6 | 13337939 | 13342397 | L1MdTf_I | + | 4458 |
| chr6 | 13342398 | 13343391 | L1MdTf_I | + | 993 |
| chr6 | 13790372 | 13791352 | L1MdTf_I | - | 980 |
| chr6 | 13791352 | 13791536 | L1MdTf_II | + | 184 |
| chr6 | 14635653 | 14635726 | L1MdTf_III | - | 73 |
| chr6 | 14635726 | 14636442 | L1MdTf_III | + | 716 |
| chr6 | 14794422 | 14799866 | L1MdTf_I | + | 5444 |
| chr6 | 14799866 | 14800859 | L1MdTf_I | + | 993 |

| | | | | | |
|---|---|---|---|---|---|
| chr6 | 14801445 | 14806904 | L1MdTf_III | + | 5459 |
| chr6 | 14806904 | 14807898 | L1MdTf_III | + | 994 |
| chr6 | 15055001 | 15055371 | L1MdTf_I | + | 370 |
| chr6 | 15055371 | 15056364 | L1MdTf_I | + | 993 |
| chr6 | 15684672 | 15690002 | L1MdTf_III | + | 5330 |
| chr6 | 15690003 | 15690996 | L1MdTf_III | + | 993 |
| chr6 | 15997470 | 16000000 | L1MdTf_II | + | 2530 |
| chr6 | 16000000 | 16003316 | L1MdTf_II | + | 3316 |
| chr6 | 16299670 | 16304800 | L1MdTf_II | + | 5130 |
| chr6 | 16304800 | 16305796 | L1MdTf_II | + | 996 |
| chr6 | 16391069 | 16396727 | L1MdTf_I | + | 5658 |
| chr6 | 16396727 | 16397738 | L1MdTf_I | + | 1011 |
| chr6 | 18899907 | 18905477 | L1MdTf_III | + | 5570 |
| chr6 | 18905478 | 18906471 | L1MdTf_III | + | 993 |
| chr6 | 19145459 | 19151800 | L1MdTf_I | + | 6341 |
| chr6 | 19151800 | 19152793 | L1MdTf_I | + | 993 |
| chr6 | 19839459 | 19841590 | L1MdTf_III | + | 2131 |
| chr6 | 19841590 | 19842583 | L1MdTf_III | + | 993 |
| chr6 | 20385228 | 20385971 | L1MdTf_II | - | 743 |
| chr6 | 20385971 | 20386223 | L1MdTf_II | + | 252 |
| chr6 | 20793426 | 20798848 | L1MdTf_II | + | 5422 |
| chr6 | 20798848 | 20799841 | L1MdTf_II | + | 993 |
| chr6 | 21014228 | 21019651 | L1MdTf_I | + | 5423 |
| chr6 | 21019651 | 21020644 | L1MdTf_I | + | 993 |
| chr6 | 21236313 | 21236361 | L1MdTf_II | - | 48 |
| chr6 | 21236361 | 21236664 | L1MdTf_II | + | 303 |
| chr6 | 21393955 | 21399848 | L1MdTf_I | + | 5893 |
| chr6 | 21399848 | 21400841 | L1MdTf_I | + | 993 |
| chr6 | 21451901 | 21457106 | L1MdTf_III | + | 5205 |
| chr6 | 21457106 | 21458100 | L1MdTf_III | + | 994 |
| chr6 | 21873283 | 21875802 | L1MdTf_II | + | 2519 |
| chr6 | 21875802 | 21876795 | L1MdTf_II | + | 993 |
| chr6 | 22125160 | 22130808 | L1MdTf_I | + | 5648 |
| chr6 | 22130808 | 22131826 | L1MdTf_I | + | 1018 |
| chr6 | 22624673 | 22625310 | L1MdTf_II | + | 637 |
| chr6 | 22625310 | 22626303 | L1MdTf_II | + | 993 |
| chr6 | 23364888 | 23370304 | L1MdTf_II | + | 5416 |
| chr6 | 23370304 | 23371297 | L1MdTf_II | + | 993 |
| chr6 | 23511278 | 23516909 | L1MdTf_II | + | 5631 |
| chr6 | 23516909 | 23517924 | L1MdTf_II | + | 1015 |
| chr6 | 23557676 | 23562949 | L1MdTf_III | + | 5273 |
| chr6 | 23562949 | 23563942 | L1MdTf_III | + | 993 |
| chr6 | 24429226 | 24434388 | L1MdTf_III | + | 5162 |
| chr6 | 24434388 | 24435381 | L1MdTf_III | + | 993 |
| chr6 | 24491745 | 24497382 | L1MdTf_II | + | 5637 |

| | | | | | |
|---|---|---|---|---|---|
| chr6 | 24497382 | 24498375 | L1MdTf_II | + | 993 |
| chr6 | 24673350 | 24678784 | L1MdTf_II | + | 5434 |
| chr6 | 24678784 | 24679774 | L1MdTf_II | + | 990 |
| chr6 | 25392941 | 25393408 | L1MdTf_II | + | 467 |
| chr6 | 25393408 | 25394392 | L1MdTf_II | + | 984 |
| chr6 | 27554232 | 27559906 | L1MdTf_III | + | 5674 |
| chr6 | 27559906 | 27560922 | L1MdTf_III | + | 1016 |
| chr6 | 32070527 | 32076590 | L1MdTf_I | + | 6063 |
| chr6 | 32076590 | 32077604 | L1MdTf_I | + | 1014 |
| chr6 | 35423196 | 35428827 | L1MdTf_II | + | 5631 |
| chr6 | 35428827 | 35429820 | L1MdTf_II | + | 993 |
| chr6 | 35769882 | 35770406 | L1MdTf_III | - | 524 |
| chr6 | 35770406 | 35770907 | L1MdTf_III | + | 501 |
| chr6 | 36473805 | 36479464 | L1MdTf_I | + | 5659 |
| chr6 | 36479464 | 36480457 | L1MdTf_I | + | 993 |
| chr6 | 41759845 | 41760026 | L1MdTf_I | - | 181 |
| chr6 | 41760026 | 41760616 | L1MdTf_I | + | 590 |
| chr6 | 42052181 | 42053525 | L1MdTf_III | + | 1344 |
| chr6 | 42053525 | 42054518 | L1MdTf_III | + | 993 |
| chr6 | 42147134 | 42147840 | L1MdTf_III | + | 706 |
| chr6 | 42147840 | 42148832 | L1MdTf_III | + | 992 |
| chr6 | 42427686 | 42432857 | L1MdTf_III | + | 5171 |
| chr6 | 42432858 | 42433850 | L1MdTf_III | + | 992 |
| chr6 | 42740518 | 42746156 | L1MdTf_II | + | 5638 |
| chr6 | 42746156 | 42747154 | L1MdTf_II | + | 998 |
| chr6 | 42804748 | 42805194 | L1MdTf_III | + | 446 |
| chr6 | 42805194 | 42806187 | L1MdTf_III | + | 993 |
| chr6 | 42830583 | 42835998 | L1MdTf_III | + | 5415 |
| chr6 | 42835998 | 42836990 | L1MdTf_III | + | 992 |
| chr6 | 43602654 | 43608099 | L1MdTf_II | + | 5445 |
| chr6 | 43608099 | 43609091 | L1MdTf_II | + | 992 |
| chr6 | 44129852 | 44130219 | L1MdTf_II | + | 367 |
| chr6 | 44130219 | 44131215 | L1MdTf_II | + | 996 |
| chr6 | 44474019 | 44474317 | L1MdTf_II | + | 298 |
| chr6 | 44474318 | 44475301 | L1MdTf_II | + | 983 |
| chr6 | 46368550 | 46370573 | L1MdTf_I | + | 2023 |
| chr6 | 46370573 | 46371569 | L1MdTf_I | + | 996 |
| chr6 | 46554078 | 46559538 | L1MdTf_II | + | 5460 |
| chr6 | 46559538 | 46560531 | L1MdTf_II | + | 993 |
| chr6 | 48205703 | 48211189 | L1MdTf_III | + | 5486 |
| chr6 | 48211189 | 48212181 | L1MdTf_III | + | 992 |
| chr6 | 53379841 | 53385068 | L1MdTf_I | + | 5227 |
| chr6 | 53385068 | 53386061 | L1MdTf_I | + | 993 |
| chr6 | 57264611 | 57270212 | L1MdTf_II | + | 5601 |
| chr6 | 57270212 | 57271205 | L1MdTf_II | + | 993 |

| | | | | | |
|---|---|---|---|---|---|
| chr6 | 57896493 | 57901822 | L1MdTf_III | + | 5329 |
| chr6 | 57901822 | 57902815 | L1MdTf_III | + | 993 |
| chr6 | 58485370 | 58487669 | L1MdTf_III | + | 2299 |
| chr6 | 58487669 | 58488663 | L1MdTf_III | + | 994 |
| chr6 | 58769374 | 58771493 | L1MdTf_II | + | 2119 |
| chr6 | 58771493 | 58772477 | L1MdTf_II | + | 984 |
| chr6 | 59182131 | 59182785 | L1MdTf_II | + | 654 |
| chr6 | 59182785 | 59183778 | L1MdTf_II | + | 993 |
| chr6 | 59202786 | 59206664 | L1MdTf_III | + | 3878 |
| chr6 | 59206664 | 59207653 | L1MdTf_III | + | 989 |
| chr6 | 59607849 | 59613166 | L1MdTf_II | + | 5317 |
| chr6 | 59613166 | 59614162 | L1MdTf_II | + | 996 |
| chr6 | 59970558 | 59970674 | L1MdTf_III | + | 116 |
| chr6 | 59970675 | 59971668 | L1MdTf_III | + | 993 |
| chr6 | 60068624 | 60068998 | L1MdTf_II | + | 374 |
| chr6 | 60068998 | 60069982 | L1MdTf_II | + | 984 |
| chr6 | 60072021 | 60077312 | L1MdTf_III | + | 5291 |
| chr6 | 60077312 | 60078305 | L1MdTf_III | + | 993 |
| chr6 | 61065386 | 61070636 | L1MdTf_III | + | 5250 |
| chr6 | 61070636 | 61071632 | L1MdTf_III | + | 996 |
| chr6 | 62265678 | 62267385 | L1MdTf_I | + | 1707 |
| chr6 | 62267385 | 62268379 | L1MdTf_I | + | 994 |
| chr6 | 62368094 | 62373327 | L1MdTf_II | + | 5233 |
| chr6 | 62373327 | 62374320 | L1MdTf_II | + | 993 |
| chr6 | 62991960 | 62998875 | L1MdTf_I | + | 6915 |
| chr6 | 62998875 | 62999868 | L1MdTf_I | + | 993 |
| chr6 | 63990332 | 63990611 | L1MdTf_II | + | 279 |
| chr6 | 63990611 | 63991631 | L1MdTf_II | + | 1020 |
| chr6 | 64471472 | 64471618 | L1MdTf_II | + | 146 |
| chr6 | 64471618 | 64472615 | L1MdTf_II | + | 997 |
| chr6 | 65192322 | 65198192 | L1MdTf_II | + | 5870 |
| chr6 | 65198192 | 65199185 | L1MdTf_II | + | 993 |
| chr6 | 65281946 | 65282441 | L1MdTf_III | + | 495 |
| chr6 | 65282441 | 65283434 | L1MdTf_III | + | 993 |
| chr6 | 65295325 | 65300563 | L1MdTf_II | + | 5238 |
| chr6 | 65300563 | 65301557 | L1MdTf_II | + | 994 |
| chr6 | 66015087 | 66016891 | L1MdTf_II | + | 1804 |
| chr6 | 66016891 | 66017884 | L1MdTf_II | + | 993 |
| chr6 | 66294252 | 66299511 | L1MdTf_III | + | 5259 |
| chr6 | 66299511 | 66300505 | L1MdTf_III | + | 994 |
| chr6 | 66743104 | 66748531 | L1MdTf_II | + | 5427 |
| chr6 | 66748532 | 66749524 | L1MdTf_II | + | 992 |
| chr6 | 67471597 | 67477040 | L1MdTf_I | + | 5443 |
| chr6 | 67477040 | 67478033 | L1MdTf_I | + | 993 |
| chr6 | 68994648 | 69000000 | L1MdTf_II | - | 5352 |

| | | | | | |
|---|---|---|---|---|---|
| chr6 | 69000000 | 69001068 | L1MdTf_II | - | 1068 |
| chr6 | 69929266 | 69934536 | L1MdTf_II | + | 5270 |
| chr6 | 69934536 | 69935529 | L1MdTf_II | + | 993 |
| chr6 | 70501170 | 70507028 | L1MdTf_I | + | 5858 |
| chr6 | 70507028 | 70508021 | L1MdTf_I | + | 993 |
| chr6 | 73321377 | 73327505 | L1MdTf_I | + | 6128 |
| chr6 | 73327505 | 73328498 | L1MdTf_I | + | 993 |
| chr6 | 73355878 | 73356042 | L1MdTf_II | - | 164 |
| chr6 | 73356042 | 73356427 | L1MdTf_I | + | 385 |
| chr6 | 73417920 | 73421451 | L1MdTf_II | - | 3531 |
| chr6 | 73421452 | 73424980 | L1MdTf_II | - | 3528 |
| chr6 | 73478430 | 73484056 | L1MdTf_II | + | 5626 |
| chr6 | 73484056 | 73485052 | L1MdTf_II | + | 996 |
| chr6 | 74340399 | 74346304 | L1MdTf_III | + | 5905 |
| chr6 | 74346304 | 74347297 | L1MdTf_III | + | 993 |
| chr6 | 77243719 | 77244227 | L1MdTf_I | - | 508 |
| chr6 | 77244227 | 77245044 | L1MdTf_II | + | 817 |
| chr6 | 78003957 | 78009402 | L1MdTf_II | + | 5445 |
| chr6 | 78009402 | 78010386 | L1MdTf_II | + | 984 |
| chr6 | 78131922 | 78137527 | L1MdTf_I | + | 5605 |
| chr6 | 78137527 | 78138520 | L1MdTf_I | + | 993 |
| chr6 | 78372368 | 78378454 | L1MdTf_II | + | 6086 |
| chr6 | 78378454 | 78379436 | L1MdTf_II | + | 982 |
| chr6 | 79104387 | 79109832 | L1MdTf_II | + | 5445 |
| chr6 | 79109832 | 79110827 | L1MdTf_II | + | 995 |
| chr6 | 79349736 | 79354869 | L1MdTf_II | + | 5133 |
| chr6 | 79354869 | 79355862 | L1MdTf_II | + | 993 |
| chr6 | 80210291 | 80210726 | L1MdTf_I | + | 435 |
| chr6 | 80210726 | 80211719 | L1MdTf_I | + | 993 |
| chr6 | 80355947 | 80361371 | L1MdTf_I | + | 5424 |
| chr6 | 80361371 | 80362364 | L1MdTf_I | + | 993 |
| chr6 | 80366032 | 80366065 | L1MdTf_I | - | 33 |
| chr6 | 80366065 | 80366516 | L1MdTf_I | + | 451 |
| chr6 | 81660643 | 81660930 | L1MdTf_III | - | 287 |
| chr6 | 81660930 | 81661206 | L1MdTf_III | + | 276 |
| chr6 | 81972648 | 81977721 | L1MdTf_II | + | 5073 |
| chr6 | 81977721 | 81978714 | L1MdTf_II | + | 993 |
| chr6 | 82189598 | 82194807 | L1MdTf_III | + | 5209 |
| chr6 | 82194807 | 82195778 | L1MdTf_III | + | 971 |
| chr6 | 82602588 | 82608247 | L1MdTf_II | + | 5659 |
| chr6 | 82608247 | 82609262 | L1MdTf_II | + | 1015 |
| chr6 | 84741309 | 84747208 | L1MdTf_III | + | 5899 |
| chr6 | 84747208 | 84748222 | L1MdTf_III | + | 1014 |
| chr6 | 85686931 | 85687557 | L1MdTf_III | - | 626 |
| chr6 | 85687557 | 85688319 | L1MdTf_III | + | 762 |

| | | | | | |
|---|---|---|---|---|---|
| chr6 | 89730901 | 89731044 | L1MdTf_III | - | 143 |
| chr6 | 89731044 | 89731271 | L1MdTf_III | + | 227 |
| chr6 | 90186478 | 90191818 | L1MdTf_III | + | 5340 |
| chr6 | 90191818 | 90192811 | L1MdTf_III | + | 993 |
| chr6 | 90210647 | 90211368 | L1MdTf_I | - | 721 |
| chr6 | 90211368 | 90212669 | L1MdTf_I | + | 1301 |
| chr6 | 90414967 | 90421039 | L1MdTf_II | + | 6072 |
| chr6 | 90421039 | 90422032 | L1MdTf_II | + | 993 |
| chr6 | 93358940 | 93359594 | L1MdTf_II | + | 654 |
| chr6 | 93359594 | 93360587 | L1MdTf_II | + | 993 |
| chr6 | 95635040 | 95640728 | L1MdTf_II | + | 5688 |
| chr6 | 95640728 | 95641743 | L1MdTf_II | + | 1015 |
| chr6 | 95928898 | 95934157 | L1MdTf_III | + | 5259 |
| chr6 | 95934157 | 95935150 | L1MdTf_III | + | 993 |
| chr6 | 95957688 | 95963109 | L1MdTf_III | + | 5421 |
| chr6 | 95963109 | 95964101 | L1MdTf_III | + | 992 |
| chr6 | 95997042 | 95998397 | L1MdTf_I | - | 1355 |
| chr6 | 95998397 | 95999683 | L1MdTf_I | + | 1286 |
| chr6 | 96572418 | 96578850 | L1MdTf_I | + | 6432 |
| chr6 | 96578850 | 96579843 | L1MdTf_I | + | 993 |
| chr6 | 96764213 | 96769639 | L1MdTf_I | + | 5426 |
| chr6 | 96769639 | 96770632 | L1MdTf_I | + | 993 |
| chr6 | 96898890 | 96904283 | L1MdTf_III | + | 5393 |
| chr6 | 96904283 | 96905275 | L1MdTf_III | + | 992 |
| chr6 | 96946588 | 96952054 | L1MdTf_II | + | 5466 |
| chr6 | 96952054 | 96953047 | L1MdTf_II | + | 993 |
| chr6 | 101620155 | 101626427 | L1MdTf_I | + | 6272 |
| chr6 | 101626428 | 101627420 | L1MdTf_I | + | 992 |
| chr6 | 102643901 | 102649446 | L1MdTf_II | + | 5545 |
| chr6 | 102649446 | 102650439 | L1MdTf_II | + | 993 |
| chr6 | 102661198 | 102666812 | L1MdTf_III | + | 5614 |
| chr6 | 102666812 | 102667835 | L1MdTf_III | + | 1023 |
| chr6 | 103696489 | 103702340 | L1MdTf_III | + | 5851 |
| chr6 | 103702340 | 103703335 | L1MdTf_III | + | 995 |
| chr6 | 104041431 | 104047523 | L1MdTf_III | + | 6092 |
| chr6 | 104047523 | 104048536 | L1MdTf_III | + | 1013 |
| chr6 | 107858666 | 107863884 | L1MdTf_III | + | 5218 |
| chr6 | 107863884 | 107864877 | L1MdTf_III | + | 993 |
| chr6 | 109043986 | 109049528 | L1MdTf_III | + | 5542 |
| chr6 | 109049528 | 109050517 | L1MdTf_III | + | 989 |
| chr6 | 109987859 | 109988984 | L1MdTf_II | + | 1125 |
| chr6 | 109988984 | 109989977 | L1MdTf_II | + | 993 |
| chr6 | 110193439 | 110196113 | L1MdTf_III | + | 2674 |
| chr6 | 110196113 | 110197106 | L1MdTf_III | + | 993 |
| chr6 | 110340367 | 110340565 | L1MdTf_III | + | 198 |

| | | | | | |
|---|---|---|---|---|---|
| chr6 | 110340565 | 110341558 | L1MdTf_III | + | 993 |
| chr6 | 110990504 | 110995738 | L1MdTf_II | + | 5234 |
| chr6 | 110995738 | 110996731 | L1MdTf_II | + | 993 |
| chr6 | 111368311 | 111373444 | L1MdTf_III | + | 5133 |
| chr6 | 111373444 | 111374437 | L1MdTf_III | + | 993 |
| chr6 | 111540214 | 111540473 | L1MdTf_III | - | 259 |
| chr6 | 111540473 | 111541225 | L1MdTf_III | + | 752 |
| chr6 | 111643999 | 111645205 | L1MdTf_II | + | 1206 |
| chr6 | 111645205 | 111646198 | L1MdTf_II | + | 993 |
| chr6 | 112156509 | 112161796 | L1MdTf_III | + | 5287 |
| chr6 | 112161796 | 112162788 | L1MdTf_III | + | 992 |
| chr6 | 112629634 | 112635576 | L1MdTf_III | + | 5942 |
| chr6 | 112635576 | 112636571 | L1MdTf_III | + | 995 |
| chr6 | 116903644 | 116909082 | L1MdTf_II | + | 5438 |
| chr6 | 116909082 | 116910075 | L1MdTf_II | + | 993 |
| chr6 | 117287336 | 117287939 | L1MdTf_II | + | 603 |
| chr6 | 117287939 | 117288935 | L1MdTf_II | + | 996 |
| chr6 | 121977594 | 121978676 | L1MdTf_III | + | 1082 |
| chr6 | 121978676 | 121979669 | L1MdTf_III | + | 993 |
| chr6 | 122146845 | 122147927 | L1MdTf_III | + | 1082 |
| chr6 | 122147927 | 122148920 | L1MdTf_III | + | 993 |
| chr6 | 124188354 | 124194495 | L1MdTf_III | + | 6141 |
| chr6 | 124194495 | 124195490 | L1MdTf_III | + | 995 |
| chr6 | 124591317 | 124593859 | L1MdTf_III | + | 2542 |
| chr6 | 124593859 | 124594852 | L1MdTf_III | + | 993 |
| chr6 | 125934901 | 125935365 | L1MdTf_II | + | 464 |
| chr6 | 125935365 | 125936359 | L1MdTf_II | + | 994 |
| chr6 | 126432839 | 126438715 | L1MdTf_I | + | 5876 |
| chr6 | 126438715 | 126439708 | L1MdTf_I | + | 993 |
| chr6 | 126879317 | 126882215 | L1MdTf_I | + | 2898 |
| chr6 | 126882215 | 126883208 | L1MdTf_I | + | 993 |
| chr6 | 129833754 | 129839817 | L1MdTf_II | + | 6063 |
| chr6 | 129839817 | 129840813 | L1MdTf_II | + | 996 |
| chr6 | 132279570 | 132284838 | L1MdTf_II | + | 5268 |
| chr6 | 132284838 | 132285822 | L1MdTf_II | + | 984 |
| chr6 | 132422912 | 132423637 | L1MdTf_I | - | 725 |
| chr6 | 132423637 | 132424169 | L1MdTf_I | + | 532 |
| chr6 | 133804742 | 133805517 | L1MdTf_II | + | 775 |
| chr6 | 133805517 | 133806536 | L1MdTf_II | + | 1019 |
| chr6 | 134303680 | 134303897 | L1MdTf_III | + | 217 |
| chr6 | 134303897 | 134304914 | L1MdTf_III | + | 1017 |
| chr6 | 135431480 | 135432888 | L1MdTf_III | + | 1408 |
| chr6 | 135432888 | 135433881 | L1MdTf_III | + | 993 |
| chr6 | 142842770 | 142848050 | L1MdTf_III | + | 5280 |
| chr6 | 142848050 | 142849043 | L1MdTf_III | + | 993 |

| | | | | | |
|---|---|---|---|---|---|
| chr6 | 144600831 | 144601459 | L1MdTf_III | + | 628 |
| chr6 | 144601459 | 144602452 | L1MdTf_III | + | 993 |
| chr7 | 3792135 | 3797867 | L1MdTf_III | + | 5732 |
| chr7 | 3797867 | 3798860 | L1MdTf_III | + | 993 |
| chr7 | 3889996 | 3895944 | L1MdTf_III | + | 5948 |
| chr7 | 3895944 | 3896937 | L1MdTf_III | + | 993 |
| chr7 | 5205618 | 5211476 | L1MdTf_II | + | 5858 |
| chr7 | 5211476 | 5212469 | L1MdTf_II | + | 993 |
| chr7 | 5707036 | 5712438 | L1MdTf_III | + | 5402 |
| chr7 | 5712438 | 5713431 | L1MdTf_III | + | 993 |
| chr7 | 5862783 | 5868827 | L1MdTf_I | + | 6044 |
| chr7 | 5868827 | 5869820 | L1MdTf_I | + | 993 |
| chr7 | 9665664 | 9666834 | L1MdTf_II | + | 1170 |
| chr7 | 9666834 | 9667827 | L1MdTf_II | + | 993 |
| chr7 | 12070157 | 12072772 | L1MdTf_III | + | 2615 |
| chr7 | 12072772 | 12073792 | L1MdTf_III | + | 1020 |
| chr7 | 12260458 | 12265898 | L1MdTf_III | + | 5440 |
| chr7 | 12265899 | 12266889 | L1MdTf_III | + | 990 |
| chr7 | 13820252 | 13827385 | L1MdTf_I | + | 7133 |
| chr7 | 13827385 | 13828378 | L1MdTf_I | + | 993 |
| chr7 | 14712744 | 14712784 | L1MdTf_III | + | 40 |
| chr7 | 14712784 | 14713463 | L1MdTf_III | + | 679 |
| chr7 | 14731986 | 14734539 | L1MdTf_III | + | 2553 |
| chr7 | 14734539 | 14735532 | L1MdTf_III | + | 993 |
| chr7 | 15196167 | 15201405 | L1MdTf_I | + | 5238 |
| chr7 | 15201405 | 15202398 | L1MdTf_I | + | 993 |
| chr7 | 15388113 | 15393117 | L1MdTf_III | + | 5004 |
| chr7 | 15393117 | 15394110 | L1MdTf_III | + | 993 |
| chr7 | 15413993 | 15419125 | L1MdTf_III | + | 5132 |
| chr7 | 15419125 | 15420118 | L1MdTf_III | + | 993 |
| chr7 | 16395295 | 16400760 | L1MdTf_I | + | 5465 |
| chr7 | 16400760 | 16401753 | L1MdTf_I | + | 993 |
| chr7 | 17841610 | 17847217 | L1MdTf_III | + | 5607 |
| chr7 | 17847217 | 17848210 | L1MdTf_III | + | 993 |
| chr7 | 17921884 | 17927279 | L1MdTf_III | + | 5395 |
| chr7 | 17927279 | 17928263 | L1MdTf_III | + | 984 |
| chr7 | 17928979 | 17930169 | L1MdTf_I | + | 1190 |
| chr7 | 17930169 | 17931162 | L1MdTf_I | + | 993 |
| chr7 | 18220962 | 18226174 | L1MdTf_II | + | 5212 |
| chr7 | 18226174 | 18227167 | L1MdTf_II | + | 993 |
| chr7 | 18352923 | 18353758 | L1MdTf_III | + | 835 |
| chr7 | 18353759 | 18354751 | L1MdTf_III | + | 992 |
| chr7 | 20002141 | 20002999 | L1MdTf_I | - | 858 |
| chr7 | 20002999 | 20003385 | L1MdTf_I | + | 386 |
| chr7 | 20132869 | 20138517 | L1MdTf_III | + | 5648 |

| | | | | | |
|---|---|---|---|---|---|
| chr7 | 20138517 | 20139509 | L1MdTf_III | + | 992 |
| chr7 | 20954414 | 20959332 | L1MdTf_III | + | 4918 |
| chr7 | 20959332 | 20960325 | L1MdTf_III | + | 993 |
| chr7 | 21410690 | 21415607 | L1MdTf_III | + | 4917 |
| chr7 | 21415608 | 21416600 | L1MdTf_III | + | 992 |
| chr7 | 21963298 | 21964156 | L1MdTf_I | - | 858 |
| chr7 | 21964156 | 21964542 | L1MdTf_I | + | 386 |
| chr7 | 22094004 | 22099652 | L1MdTf_III | + | 5648 |
| chr7 | 22099652 | 22100644 | L1MdTf_III | + | 992 |
| chr7 | 22633889 | 22638806 | L1MdTf_III | + | 4917 |
| chr7 | 22638807 | 22639799 | L1MdTf_III | + | 992 |
| chr7 | 25285736 | 25288757 | L1MdTf_III | + | 3021 |
| chr7 | 25288757 | 25289750 | L1MdTf_III | + | 993 |
| chr7 | 25983200 | 25988817 | L1MdTf_II | + | 5617 |
| chr7 | 25988817 | 25989810 | L1MdTf_II | + | 993 |
| chr7 | 27767326 | 27767639 | L1MdTf_II | - | 313 |
| chr7 | 27767639 | 27767698 | L1MdTf_II | + | 59 |
| chr7 | 31720341 | 31720593 | L1MdTf_III | - | 252 |
| chr7 | 31720593 | 31723039 | L1MdTf_III | - | 2446 |
| chr7 | 31969890 | 31975469 | L1MdTf_I | + | 5579 |
| chr7 | 31975469 | 31976462 | L1MdTf_I | + | 993 |
| chr7 | 32452616 | 32452868 | L1MdTf_III | - | 252 |
| chr7 | 32452868 | 32455542 | L1MdTf_III | - | 2674 |
| chr7 | 32679515 | 32679767 | L1MdTf_III | - | 252 |
| chr7 | 32679767 | 32682441 | L1MdTf_III | - | 2674 |
| chr7 | 40109154 | 40114334 | L1MdTf_III | + | 5180 |
| chr7 | 40114334 | 40115327 | L1MdTf_III | + | 993 |
| chr7 | 40884451 | 40889650 | L1MdTf_III | + | 5199 |
| chr7 | 40889650 | 40890624 | L1MdTf_III | + | 974 |
| chr7 | 41078641 | 41084104 | L1MdTf_II | + | 5463 |
| chr7 | 41084104 | 41085097 | L1MdTf_II | + | 993 |
| chr7 | 43792973 | 43798161 | L1MdTf_III | + | 5188 |
| chr7 | 43798161 | 43799154 | L1MdTf_III | + | 993 |
| chr7 | 48062092 | 48067540 | L1MdTf_III | + | 5448 |
| chr7 | 48067541 | 48068536 | L1MdTf_III | + | 995 |
| chr7 | 50101317 | 50106593 | L1MdTf_II | + | 5276 |
| chr7 | 50106594 | 50107586 | L1MdTf_II | + | 992 |
| chr7 | 50527884 | 50528538 | L1MdTf_II | + | 654 |
| chr7 | 50528538 | 50529531 | L1MdTf_II | + | 993 |
| chr7 | 50767141 | 50773644 | L1MdTf_II | + | 6503 |
| chr7 | 50773644 | 50774639 | L1MdTf_II | + | 995 |
| chr7 | 50924237 | 50928795 | L1MdTf_I | + | 4558 |
| chr7 | 50928795 | 50929788 | L1MdTf_I | + | 993 |
| chr7 | 51451908 | 51456835 | L1MdTf_I | + | 4927 |
| chr7 | 51456835 | 51457828 | L1MdTf_I | + | 993 |

| | | | | | |
|---|---|---|---|---|---|
| chr7 | 52419196 | 52424647 | L1MdTf_II | + | 5451 |
| chr7 | 52424647 | 52425640 | L1MdTf_II | + | 993 |
| chr7 | 52465384 | 52470640 | L1MdTf_II | + | 5256 |
| chr7 | 52470640 | 52471633 | L1MdTf_II | + | 993 |
| chr7 | 52486716 | 52486882 | L1MdTf_III | - | 166 |
| chr7 | 52486882 | 52486986 | L1MdTf_III | + | 104 |
| chr7 | 52577137 | 52577823 | L1MdTf_III | + | 686 |
| chr7 | 52577824 | 52578816 | L1MdTf_III | + | 992 |
| chr7 | 52603428 | 52604294 | L1MdTf_II | - | 866 |
| chr7 | 52604294 | 52605827 | L1MdTf_II | + | 1533 |
| chr7 | 53027849 | 53033148 | L1MdTf_III | + | 5299 |
| chr7 | 53033148 | 53034138 | L1MdTf_III | + | 990 |
| chr7 | 53207308 | 53212937 | L1MdTf_I | + | 5629 |
| chr7 | 53212937 | 53213930 | L1MdTf_I | + | 993 |
| chr7 | 53708567 | 53709211 | L1MdTf_II | + | 644 |
| chr7 | 53709211 | 53710204 | L1MdTf_II | + | 993 |
| chr7 | 54572753 | 54578622 | L1MdTf_II | + | 5869 |
| chr7 | 54578622 | 54579615 | L1MdTf_II | + | 993 |
| chr7 | 54764533 | 54769835 | L1MdTf_II | + | 5302 |
| chr7 | 54769835 | 54770829 | L1MdTf_II | + | 994 |
| chr7 | 55254925 | 55255525 | L1MdTf_III | + | 600 |
| chr7 | 55255525 | 55256518 | L1MdTf_III | + | 993 |
| chr7 | 56166056 | 56166710 | L1MdTf_I | + | 654 |
| chr7 | 56166710 | 56167727 | L1MdTf_II | + | 1017 |
| chr7 | 57720465 | 57725944 | L1MdTf_III | + | 5479 |
| chr7 | 57725944 | 57726937 | L1MdTf_III | + | 993 |
| chr7 | 57750272 | 57755820 | L1MdTf_I | + | 5548 |
| chr7 | 57755820 | 57756813 | L1MdTf_I | + | 993 |
| chr7 | 58071475 | 58077071 | L1MdTf_I | + | 5596 |
| chr7 | 58077071 | 58078063 | L1MdTf_I | + | 992 |
| chr7 | 58774936 | 58780334 | L1MdTf_I | + | 5398 |
| chr7 | 58780334 | 58781327 | L1MdTf_I | + | 993 |
| chr7 | 59043696 | 59049028 | L1MdTf_II | + | 5332 |
| chr7 | 59049028 | 59050021 | L1MdTf_II | + | 993 |
| chr7 | 59050024 | 59052569 | L1MdTf_III | + | 2545 |
| chr7 | 59052569 | 59053587 | L1MdTf_III | + | 1018 |
| chr7 | 59972133 | 59977405 | L1MdTf_I | + | 5272 |
| chr7 | 59977405 | 59978398 | L1MdTf_I | + | 993 |
| chr7 | 59994651 | 60000000 | L1MdTf_III | - | 5349 |
| chr7 | 60000000 | 60001291 | L1MdTf_III | - | 1291 |
| chr7 | 60109424 | 60113115 | L1MdTf_II | + | 3691 |
| chr7 | 60113115 | 60114130 | L1MdTf_II | + | 1015 |
| chr7 | 60425296 | 60431088 | L1MdTf_I | + | 5792 |
| chr7 | 60431088 | 60432105 | L1MdTf_I | + | 1017 |
| chr7 | 60832052 | 60838389 | L1MdTf_III | + | 6337 |

| | | | | | |
|---|---|---|---|---|---|
| chr7 | 60838389 | 60839403 | L1MdTf_III | + | 1014 |
| chr7 | 60995897 | 61000000 | L1MdTf_III | + | 4103 |
| chr7 | 61000000 | 61000863 | L1MdTf_I | + | 863 |
| chr7 | 61001358 | 61005331 | L1MdTf_I | + | 3973 |
| chr7 | 61005331 | 61006324 | L1MdTf_I | + | 993 |
| chr7 | 61012281 | 61016254 | L1MdTf_I | + | 3973 |
| chr7 | 61016254 | 61017273 | L1MdTf_I | + | 1019 |
| chr7 | 61017762 | 61021735 | L1MdTf_I | + | 3973 |
| chr7 | 61021735 | 61022725 | L1MdTf_I | + | 990 |
| chr7 | 61157111 | 61162355 | L1MdTf_III | + | 5244 |
| chr7 | 61162355 | 61163347 | L1MdTf_III | + | 992 |
| chr7 | 62336048 | 62342092 | L1MdTf_I | + | 6044 |
| chr7 | 62342092 | 62343085 | L1MdTf_I | + | 993 |
| chr7 | 62922047 | 62927389 | L1MdTf_III | + | 5342 |
| chr7 | 62927389 | 62928376 | L1MdTf_III | + | 987 |
| chr7 | 64322871 | 64328354 | L1MdTf_III | + | 5483 |
| chr7 | 64328354 | 64329347 | L1MdTf_III | + | 993 |
| chr7 | 66027383 | 66027422 | L1MdTf_II | - | 39 |
| chr7 | 66027422 | 66027767 | L1MdTf_II | + | 345 |
| chr7 | 66923678 | 66929197 | L1MdTf_II | + | 5519 |
| chr7 | 66929197 | 66930190 | L1MdTf_II | + | 993 |
| chr7 | 69842993 | 69848759 | L1MdTf_II | + | 5766 |
| chr7 | 69848759 | 69849752 | L1MdTf_II | + | 993 |
| chr7 | 72577471 | 72583330 | L1MdTf_I | + | 5859 |
| chr7 | 72583330 | 72584323 | L1MdTf_I | + | 993 |
| chr7 | 72657296 | 72662741 | L1MdTf_II | + | 5445 |
| chr7 | 72662741 | 72663734 | L1MdTf_II | + | 993 |
| chr7 | 74449064 | 74454958 | L1MdTf_II | + | 5894 |
| chr7 | 74454958 | 74455951 | L1MdTf_II | + | 993 |
| chr7 | 74836951 | 74837426 | L1MdTf_I | + | 475 |
| chr7 | 74837426 | 74838419 | L1MdTf_I | + | 993 |
| chr7 | 77524781 | 77530237 | L1MdTf_II | + | 5456 |
| chr7 | 77530237 | 77531252 | L1MdTf_II | + | 1015 |
| chr7 | 77604366 | 77607243 | L1MdTf_II | + | 2877 |
| chr7 | 77607243 | 77608263 | L1MdTf_II | + | 1020 |
| chr7 | 78065783 | 78071359 | L1MdTf_I | + | 5576 |
| chr7 | 78071359 | 78072354 | L1MdTf_I | + | 995 |
| chr7 | 81863102 | 81868976 | L1MdTf_II | + | 5874 |
| chr7 | 81868976 | 81869994 | L1MdTf_II | + | 1018 |
| chr7 | 82117270 | 82122450 | L1MdTf_II | + | 5180 |
| chr7 | 82122450 | 82123443 | L1MdTf_II | + | 993 |
| chr7 | 83142216 | 83143547 | L1MdTf_III | - | 1331 |
| chr7 | 83143548 | 83144528 | L1MdTf_III | + | 980 |
| chr7 | 83432531 | 83437679 | L1MdTf_III | + | 5148 |
| chr7 | 83437679 | 83438672 | L1MdTf_III | + | 993 |

| | | | | | |
|---|---|---|---|---|---|
| chr7 | 85036852 | 85042085 | L1MdTf_I | + | 5233 |
| chr7 | 85042085 | 85043078 | L1MdTf_I | + | 993 |
| chr7 | 85176269 | 85177191 | L1MdTf_III | + | 922 |
| chr7 | 85177192 | 85178184 | L1MdTf_III | + | 992 |
| chr7 | 85759515 | 85760007 | L1MdTf_III | - | 492 |
| chr7 | 85760007 | 85761194 | L1MdTf_III | + | 1187 |
| chr7 | 85767886 | 85774067 | L1MdTf_II | + | 6181 |
| chr7 | 85774067 | 85775060 | L1MdTf_II | + | 993 |
| chr7 | 86266983 | 86272411 | L1MdTf_I | + | 5428 |
| chr7 | 86272411 | 86273404 | L1MdTf_I | + | 993 |
| chr7 | 87064467 | 87069886 | L1MdTf_III | + | 5419 |
| chr7 | 87069886 | 87070881 | L1MdTf_III | + | 995 |
| chr7 | 89338696 | 89343900 | L1MdTf_I | + | 5204 |
| chr7 | 89343900 | 89344893 | L1MdTf_I | + | 993 |
| chr7 | 89344926 | 89345526 | L1MdTf_III | + | 600 |
| chr7 | 89345526 | 89346547 | L1MdTf_III | + | 1021 |
| chr7 | 90519425 | 90519879 | L1MdTf_I | + | 454 |
| chr7 | 90519879 | 90520872 | L1MdTf_I | + | 993 |
| chr7 | 91596658 | 91601701 | L1MdTf_III | + | 5043 |
| chr7 | 91601701 | 91602694 | L1MdTf_III | + | 993 |
| chr7 | 91927126 | 91928205 | L1MdTf_II | + | 1079 |
| chr7 | 91928205 | 91929198 | L1MdTf_II | + | 993 |
| chr7 | 92436152 | 92442550 | L1MdTf_III | + | 6398 |
| chr7 | 92442550 | 92443568 | L1MdTf_III | + | 1018 |
| chr7 | 92443568 | 92443867 | L1MdTf_III | + | 299 |
| chr7 | 92443867 | 92444860 | L1MdTf_III | + | 993 |
| chr7 | 92762414 | 92764747 | L1MdTf_I | + | 2333 |
| chr7 | 92764747 | 92765740 | L1MdTf_I | + | 993 |
| chr7 | 93294349 | 93294825 | L1MdTf_II | - | 476 |
| chr7 | 93294825 | 93295572 | L1MdTf_II | + | 747 |
| chr7 | 93393213 | 93398769 | L1MdTf_II | + | 5556 |
| chr7 | 93398769 | 93399764 | L1MdTf_II | + | 995 |
| chr7 | 93851855 | 93852560 | L1MdTf_II | + | 705 |
| chr7 | 93852560 | 93853544 | L1MdTf_II | + | 984 |
| chr7 | 93942155 | 93947580 | L1MdTf_I | + | 5425 |
| chr7 | 93947580 | 93948573 | L1MdTf_I | + | 993 |
| chr7 | 96876529 | 96882108 | L1MdTf_II | + | 5579 |
| chr7 | 96882108 | 96883101 | L1MdTf_II | + | 993 |
| chr7 | 96883146 | 96884306 | L1MdTf_I | + | 1160 |
| chr7 | 96884306 | 96885327 | L1MdTf_I | + | 1021 |
| chr7 | 102443897 | 102449368 | L1MdTf_II | + | 5471 |
| chr7 | 102449368 | 102450384 | L1MdTf_II | + | 1016 |
| chr7 | 102482514 | 102487729 | L1MdTf_I | + | 5215 |
| chr7 | 102487729 | 102488722 | L1MdTf_I | + | 993 |
| chr7 | 104072587 | 104076029 | L1MdTf_I | + | 3442 |

| | | | | | |
|---|---|---|---|---|---|
| chr7 | 104076029 | 104077022 | L1MdTf_I | + | 993 |
| chr7 | 104759140 | 104764058 | L1MdTf_III | + | 4918 |
| chr7 | 104764058 | 104765084 | L1MdTf_III | + | 1026 |
| chr7 | 104999410 | 105000000 | L1MdTf_III | + | 590 |
| chr7 | 105000000 | 105004845 | L1MdTf_III | + | 4845 |
| chr7 | 105008879 | 105009107 | L1MdTf_I | + | 228 |
| chr7 | 105009107 | 105010100 | L1MdTf_I | + | 993 |
| chr7 | 105071243 | 105071307 | L1MdTf_II | - | 64 |
| chr7 | 105071307 | 105071509 | L1MdTf_II | + | 202 |
| chr7 | 106492482 | 106492933 | L1MdTf_III | - | 451 |
| chr7 | 106492933 | 106493671 | L1MdTf_III | + | 738 |
| chr7 | 107453188 | 107459240 | L1MdTf_II | + | 6052 |
| chr7 | 107459240 | 107460232 | L1MdTf_II | + | 992 |
| chr7 | 112205159 | 112210545 | L1MdTf_III | + | 5386 |
| chr7 | 112210545 | 112211538 | L1MdTf_III | + | 993 |
| chr7 | 113719191 | 113724419 | L1MdTf_II | + | 5228 |
| chr7 | 113724419 | 113725416 | L1MdTf_II | + | 997 |
| chr7 | 114164805 | 114165631 | L1MdTf_III | + | 826 |
| chr7 | 114165631 | 114166624 | L1MdTf_III | + | 993 |
| chr7 | 119104042 | 119109275 | L1MdTf_I | + | 5233 |
| chr7 | 119109275 | 119110268 | L1MdTf_I | + | 993 |
| chr7 | 120105238 | 120111288 | L1MdTf_II | + | 6050 |
| chr7 | 120111288 | 120112309 | L1MdTf_II | + | 1021 |
| chr7 | 123498942 | 123499363 | L1MdTf_I | - | 421 |
| chr7 | 123499363 | 123499787 | L1MdTf_I | + | 424 |
| chr7 | 130803499 | 130804244 | L1MdTf_III | - | 745 |
| chr7 | 130804244 | 130804491 | L1MdTf_III | + | 247 |
| chr7 | 131089162 | 131095499 | L1MdTf_III | + | 6337 |
| chr7 | 131095499 | 131096494 | L1MdTf_III | + | 995 |
| chr7 | 140194161 | 140199733 | L1MdTf_III | + | 5572 |
| chr7 | 140199733 | 140200716 | L1MdTf_III | + | 983 |
| chr8 | 5291897 | 5297370 | L1MdTf_III | + | 5473 |
| chr8 | 5297370 | 5298363 | L1MdTf_III | + | 993 |
| chr8 | 6203086 | 6208330 | L1MdTf_III | + | 5244 |
| chr8 | 6208330 | 6209324 | L1MdTf_III | + | 994 |
| chr8 | 6687573 | 6693213 | L1MdTf_II | + | 5640 |
| chr8 | 6693213 | 6694206 | L1MdTf_II | + | 993 |
| chr8 | 7592222 | 7597707 | L1MdTf_I | + | 5485 |
| chr8 | 7597707 | 7598700 | L1MdTf_I | + | 993 |
| chr8 | 9601190 | 9607169 | L1MdTf_I | + | 5979 |
| chr8 | 9607169 | 9608182 | L1MdTf_I | + | 1013 |
| chr8 | 10270811 | 10271057 | L1MdTf_I | - | 246 |
| chr8 | 10271057 | 10271911 | L1MdTf_II | + | 854 |
| chr8 | 15492208 | 15498469 | L1MdTf_III | + | 6261 |
| chr8 | 15498469 | 15499487 | L1MdTf_III | + | 1018 |

| | | | | | |
|---|---|---|---|---|---|
| chr8 | 15702253 | 15707698 | L1MdTf_I | + | 5445 |
| chr8 | 15707698 | 15708691 | L1MdTf_I | + | 993 |
| chr8 | 15763167 | 15768859 | L1MdTf_III | + | 5692 |
| chr8 | 15768859 | 15769853 | L1MdTf_III | + | 994 |
| chr8 | 16266399 | 16271844 | L1MdTf_II | + | 5445 |
| chr8 | 16271844 | 16272828 | L1MdTf_II | + | 984 |
| chr8 | 16860776 | 16861657 | L1MdTf_II | + | 881 |
| chr8 | 16861657 | 16862650 | L1MdTf_II | + | 993 |
| chr8 | 22626991 | 22628629 | L1MdTf_I | + | 1638 |
| chr8 | 22628629 | 22629622 | L1MdTf_I | + | 993 |
| chr8 | 27809256 | 27815143 | L1MdTf_III | + | 5887 |
| chr8 | 27815143 | 27816159 | L1MdTf_III | + | 1016 |
| chr8 | 27997976 | 28000000 | L1MdTf_II | - | 2024 |
| chr8 | 28000000 | 28004625 | L1MdTf_II | - | 4625 |
| chr8 | 28293039 | 28298042 | L1MdTf_III | + | 5003 |
| chr8 | 28298043 | 28299034 | L1MdTf_III | + | 991 |
| chr8 | 28772790 | 28772943 | L1MdTf_II | + | 153 |
| chr8 | 28772943 | 28773936 | L1MdTf_II | + | 993 |
| chr8 | 29021978 | 29025749 | L1MdTf_III | + | 3771 |
| chr8 | 29025749 | 29026741 | L1MdTf_III | + | 992 |
| chr8 | 29684058 | 29689906 | L1MdTf_II | + | 5848 |
| chr8 | 29689906 | 29690899 | L1MdTf_I | + | 993 |
| chr8 | 30288090 | 30293305 | L1MdTf_III | + | 5215 |
| chr8 | 30293306 | 30294298 | L1MdTf_III | + | 992 |
| chr8 | 30525927 | 30531342 | L1MdTf_III | + | 5415 |
| chr8 | 30531343 | 30532336 | L1MdTf_III | + | 993 |
| chr8 | 31359210 | 31364591 | L1MdTf_III | + | 5381 |
| chr8 | 31364592 | 31365585 | L1MdTf_III | + | 993 |
| chr8 | 31377187 | 31378375 | L1MdTf_III | + | 1188 |
| chr8 | 31378375 | 31379367 | L1MdTf_III | + | 992 |
| chr8 | 33204276 | 33209776 | L1MdTf_III | + | 5500 |
| chr8 | 33209776 | 33210769 | L1MdTf_III | + | 993 |
| chr8 | 33713560 | 33718944 | L1MdTf_III | + | 5384 |
| chr8 | 33718944 | 33719937 | L1MdTf_III | + | 993 |
| chr8 | 33744311 | 33744516 | L1MdTf_III | - | 205 |
| chr8 | 33744516 | 33745274 | L1MdTf_III | + | 758 |
| chr8 | 33978596 | 33980304 | L1MdTf_III | + | 1708 |
| chr8 | 33980304 | 33981295 | L1MdTf_III | + | 991 |
| chr8 | 38039442 | 38044813 | L1MdTf_III | + | 5371 |
| chr8 | 38044813 | 38045805 | L1MdTf_III | + | 992 |
| chr8 | 38094418 | 38099894 | L1MdTf_III | + | 5476 |
| chr8 | 38099894 | 38100886 | L1MdTf_III | + | 992 |
| chr8 | 39342182 | 39347620 | L1MdTf_III | + | 5438 |
| chr8 | 39347620 | 39348611 | L1MdTf_III | + | 991 |
| chr8 | 40277139 | 40282487 | L1MdTf_III | + | 5348 |

| | | | | | |
|---|---|---|---|---|---|
| chr8 | 40282487 | 40283480 | L1MdTf_III | + | 993 |
| chr8 | 40501903 | 40502017 | L1MdTf_III | - | 114 |
| chr8 | 40502017 | 40502296 | L1MdTf_III | + | 279 |
| chr8 | 41228937 | 41234381 | L1MdTf_III | + | 5444 |
| chr8 | 41234381 | 41235374 | L1MdTf_III | + | 993 |
| chr8 | 42700992 | 42702150 | L1MdTf_III | + | 1158 |
| chr8 | 42702151 | 42703143 | L1MdTf_III | + | 992 |
| chr8 | 42905046 | 42905369 | L1MdTf_III | + | 323 |
| chr8 | 42905369 | 42906362 | L1MdTf_III | + | 993 |
| chr8 | 43072386 | 43077792 | L1MdTf_III | + | 5406 |
| chr8 | 43077792 | 43078785 | L1MdTf_III | + | 993 |
| chr8 | 44033485 | 44034584 | L1MdTf_III | + | 1099 |
| chr8 | 44034584 | 44035577 | L1MdTf_III | + | 993 |
| chr8 | 44244722 | 44250353 | L1MdTf_III | + | 5631 |
| chr8 | 44250354 | 44251346 | L1MdTf_III | + | 992 |
| chr8 | 44414278 | 44419735 | L1MdTf_II | + | 5457 |
| chr8 | 44419735 | 44420728 | L1MdTf_II | + | 993 |
| chr8 | 44440300 | 44440479 | L1MdTf_I | - | 179 |
| chr8 | 44440479 | 44442580 | L1MdTf_II | + | 2101 |
| chr8 | 44686889 | 44692656 | L1MdTf_II | + | 5767 |
| chr8 | 44692656 | 44693649 | L1MdTf_II | + | 993 |
| chr8 | 44695976 | 44701194 | L1MdTf_II | + | 5218 |
| chr8 | 44701194 | 44702188 | L1MdTf_II | + | 994 |
| chr8 | 50026732 | 50032742 | L1MdTf_I | + | 6010 |
| chr8 | 50032742 | 50033735 | L1MdTf_I | + | 993 |
| chr8 | 50112493 | 50112983 | L1MdTf_II | - | 490 |
| chr8 | 50112983 | 50113074 | L1MdTf_I | + | 91 |
| chr8 | 50696929 | 50702084 | L1MdTf_I | + | 5155 |
| chr8 | 50702084 | 50703077 | L1MdTf_I | + | 993 |
| chr8 | 51799676 | 51805542 | L1MdTf_II | + | 5866 |
| chr8 | 51805542 | 51806539 | L1MdTf_II | + | 997 |
| chr8 | 52235375 | 52235789 | L1MdTf_III | - | 414 |
| chr8 | 52235789 | 52238544 | L1MdTf_III | + | 2755 |
| chr8 | 52756596 | 52766706 | L1MdTf_I | + | 10110 |
| chr8 | 52766706 | 52767697 | L1MdTf_I | + | 991 |
| chr8 | 53356825 | 53362184 | L1MdTf_III | + | 5359 |
| chr8 | 53362184 | 53363177 | L1MdTf_III | + | 993 |
| chr8 | 53566819 | 53572029 | L1MdTf_I | + | 5210 |
| chr8 | 53572029 | 53573022 | L1MdTf_I | + | 993 |
| chr8 | 53793798 | 53794139 | L1MdTf_III | - | 341 |
| chr8 | 53794139 | 53794551 | L1MdTf_III | + | 412 |
| chr8 | 56259178 | 56264392 | L1MdTf_III | + | 5214 |
| chr8 | 56264392 | 56265384 | L1MdTf_III | + | 992 |
| chr8 | 56292619 | 56297865 | L1MdTf_III | + | 5246 |
| chr8 | 56297865 | 56298859 | L1MdTf_III | + | 994 |

| | | | | | |
|---|---|---|---|---|---|
| chr8 | 57319809 | 57323962 | L1MdTf_I | + | 4153 |
| chr8 | 57323962 | 57324954 | L1MdTf_I | + | 992 |
| chr8 | 57395982 | 57401597 | L1MdTf_III | + | 5615 |
| chr8 | 57401597 | 57402590 | L1MdTf_III | + | 993 |
| chr8 | 57635639 | 57641538 | L1MdTf_I | + | 5899 |
| chr8 | 57641538 | 57642531 | L1MdTf_I | + | 993 |
| chr8 | 58841369 | 58847680 | L1MdTf_II | + | 6311 |
| chr8 | 58847680 | 58848664 | L1MdTf_II | + | 984 |
| chr8 | 58938057 | 58939313 | L1MdTf_III | - | 1256 |
| chr8 | 58939313 | 58939919 | L1MdTf_III | + | 606 |
| chr8 | 60102711 | 60108586 | L1MdTf_I | + | 5875 |
| chr8 | 60108586 | 60109579 | L1MdTf_I | + | 993 |
| chr8 | 60130564 | 60136888 | L1MdTf_III | + | 6324 |
| chr8 | 60136888 | 60137881 | L1MdTf_III | + | 993 |
| chr8 | 60308168 | 60308605 | L1MdTf_I | - | 437 |
| chr8 | 60308605 | 60309383 | L1MdTf_I | + | 778 |
| chr8 | 60509345 | 60514752 | L1MdTf_I | + | 5407 |
| chr8 | 60514752 | 60515745 | L1MdTf_I | + | 993 |
| chr8 | 60556129 | 60560369 | L1MdTf_III | + | 4240 |
| chr8 | 60560369 | 60561360 | L1MdTf_III | + | 991 |
| chr8 | 60903279 | 60904792 | L1MdTf_I | - | 1513 |
| chr8 | 60904792 | 60905182 | L1MdTf_I | + | 390 |
| chr8 | 60925983 | 60931290 | L1MdTf_III | + | 5307 |
| chr8 | 60931290 | 60932283 | L1MdTf_III | + | 993 |
| chr8 | 61307330 | 61309323 | L1MdTf_III | + | 1993 |
| chr8 | 61309323 | 61310316 | L1MdTf_III | + | 993 |
| chr8 | 63500489 | 63505713 | L1MdTf_III | + | 5224 |
| chr8 | 63505714 | 63506706 | L1MdTf_III | + | 992 |
| chr8 | 64898862 | 64905109 | L1MdTf_I | + | 6247 |
| chr8 | 64905109 | 64906102 | L1MdTf_I | + | 993 |
| chr8 | 65343199 | 65348428 | L1MdTf_I | + | 5229 |
| chr8 | 65348428 | 65349449 | L1MdTf_I | + | 1021 |
| chr8 | 65637521 | 65642837 | L1MdTf_III | + | 5316 |
| chr8 | 65642837 | 65643830 | L1MdTf_III | + | 993 |
| chr8 | 65694749 | 65699979 | L1MdTf_III | + | 5230 |
| chr8 | 65699979 | 65700972 | L1MdTf_III | + | 993 |
| chr8 | 65856039 | 65861890 | L1MdTf_II | + | 5851 |
| chr8 | 65861890 | 65862883 | L1MdTf_II | + | 993 |
| chr8 | 65906570 | 65912019 | L1MdTf_I | + | 5449 |
| chr8 | 65912019 | 65913012 | L1MdTf_I | + | 993 |
| chr8 | 65913025 | 65915042 | L1MdTf_III | + | 2017 |
| chr8 | 65915042 | 65916035 | L1MdTf_III | + | 993 |
| chr8 | 66043530 | 66049126 | L1MdTf_I | + | 5596 |
| chr8 | 66049126 | 66050119 | L1MdTf_I | + | 993 |
| chr8 | 67488010 | 67494196 | L1MdTf_III | + | 6186 |

| | | | | | |
|---|---|---|---|---|---|
| chr8 | 67494196 | 67495190 | L1MdTf_III | + | 994 |
| chr8 | 67608238 | 67608299 | L1MdTf_II | - | 61 |
| chr8 | 67608299 | 67608965 | L1MdTf_II | + | 666 |
| chr8 | 68673545 | 68679101 | L1MdTf_I | + | 5556 |
| chr8 | 68679101 | 68680094 | L1MdTf_I | + | 993 |
| chr8 | 73694584 | 73700040 | L1MdTf_III | + | 5456 |
| chr8 | 73700041 | 73701055 | L1MdTf_III | + | 1014 |
| chr8 | 74917695 | 74923095 | L1MdTf_II | + | 5400 |
| chr8 | 74923095 | 74924116 | L1MdTf_II | + | 1021 |
| chr8 | 74997330 | 75000000 | L1MdTf_II | + | 2670 |
| chr8 | 75000000 | 75000247 | L1MdTf_II | + | 247 |
| chr8 | 75211506 | 75216772 | L1MdTf_III | + | 5266 |
| chr8 | 75216772 | 75217765 | L1MdTf_III | + | 993 |
| chr8 | 75542039 | 75542352 | L1MdTf_III | + | 313 |
| chr8 | 75542352 | 75543376 | L1MdTf_III | + | 1024 |
| chr8 | 76824417 | 76830274 | L1MdTf_II | + | 5857 |
| chr8 | 76830274 | 76831267 | L1MdTf_II | + | 993 |
| chr8 | 77141027 | 77145897 | L1MdTf_II | + | 4870 |
| chr8 | 77145897 | 77146890 | L1MdTf_II | + | 993 |
| chr8 | 77504804 | 77506362 | L1MdTf_III | + | 1558 |
| chr8 | 77506362 | 77507347 | L1MdTf_III | + | 985 |
| chr8 | 77672425 | 77673182 | L1MdTf_I | - | 757 |
| chr8 | 77673182 | 77674973 | L1MdTf_I | + | 1791 |
| chr8 | 78333997 | 78334031 | L1MdTf_II | + | 34 |
| chr8 | 78334031 | 78334789 | L1MdTf_II | + | 758 |
| chr8 | 78591301 | 78594480 | L1MdTf_II | + | 3179 |
| chr8 | 78594480 | 78595495 | L1MdTf_II | + | 1015 |
| chr8 | 78633748 | 78639635 | L1MdTf_II | + | 5887 |
| chr8 | 78639635 | 78640631 | L1MdTf_II | + | 996 |
| chr8 | 80794944 | 80795598 | L1MdTf_I | - | 654 |
| chr8 | 80795598 | 80796790 | L1MdTf_I | + | 1192 |
| chr8 | 80885296 | 80885530 | L1MdTf_III | - | 234 |
| chr8 | 80885531 | 80886314 | L1MdTf_III | + | 783 |
| chr8 | 81072597 | 81078023 | L1MdTf_I | + | 5426 |
| chr8 | 81078023 | 81079016 | L1MdTf_I | + | 993 |
| chr8 | 81135041 | 81140494 | L1MdTf_III | + | 5453 |
| chr8 | 81140494 | 81141506 | L1MdTf_III | + | 1012 |
| chr8 | 81180787 | 81184918 | L1MdTf_II | + | 4131 |
| chr8 | 81184918 | 81185915 | L1MdTf_II | + | 997 |
| chr8 | 81366713 | 81367739 | L1MdTf_II | + | 1026 |
| chr8 | 81367739 | 81368753 | L1MdTf_II | + | 1014 |
| chr8 | 81799531 | 81805004 | L1MdTf_III | + | 5473 |
| chr8 | 81805004 | 81805997 | L1MdTf_III | + | 993 |
| chr8 | 82215410 | 82215545 | L1MdTf_III | - | 135 |
| chr8 | 82215545 | 82216158 | L1MdTf_III | + | 613 |

| | | | | | |
|---|---|---|---|---|---|
| chr8 | 82447935 | 82453319 | L1MdTf_III | + | 5384 |
| chr8 | 82453319 | 82454312 | L1MdTf_III | + | 993 |
| chr8 | 82777183 | 82777680 | L1MdTf_III | + | 497 |
| chr8 | 82777680 | 82778671 | L1MdTf_III | + | 991 |
| chr8 | 82935416 | 82938448 | L1MdTf_III | + | 3032 |
| chr8 | 82938449 | 82939441 | L1MdTf_III | + | 992 |
| chr8 | 83162031 | 83164914 | L1MdTf_III | - | 2883 |
| chr8 | 83164914 | 83165745 | L1MdTf_III | + | 831 |
| chr8 | 83371276 | 83376774 | L1MdTf_III | + | 5498 |
| chr8 | 83376774 | 83377767 | L1MdTf_III | + | 993 |
| chr8 | 83545327 | 83550964 | L1MdTf_III | + | 5637 |
| chr8 | 83550964 | 83551957 | L1MdTf_III | + | 993 |
| chr8 | 86674991 | 86680642 | L1MdTf_III | + | 5651 |
| chr8 | 86680642 | 86681635 | L1MdTf_III | + | 993 |
| chr8 | 86942623 | 86947836 | L1MdTf_II | + | 5213 |
| chr8 | 86947836 | 86948829 | L1MdTf_II | + | 993 |
| chr8 | 87806653 | 87807184 | L1MdTf_III | + | 531 |
| chr8 | 87807184 | 87808177 | L1MdTf_III | + | 993 |
| chr8 | 94232369 | 94237660 | L1MdTf_III | + | 5291 |
| chr8 | 94237660 | 94238652 | L1MdTf_III | + | 992 |
| chr8 | 98209719 | 98209994 | L1MdTf_III | + | 275 |
| chr8 | 98209994 | 98210987 | L1MdTf_III | + | 993 |
| chr8 | 98794061 | 98794261 | L1MdTf_III | - | 200 |
| chr8 | 98794261 | 98794437 | L1MdTf_III | + | 176 |
| chr8 | 99155758 | 99161105 | L1MdTf_III | + | 5347 |
| chr8 | 99161105 | 99162098 | L1MdTf_III | + | 993 |
| chr8 | 100192908 | 100198989 | L1MdTf_II | + | 6081 |
| chr8 | 100198989 | 100199982 | L1MdTf_II | + | 993 |
| chr8 | 101101729 | 101105486 | L1MdTf_III | + | 3757 |
| chr8 | 101105487 | 101106477 | L1MdTf_III | + | 990 |
| chr8 | 113210221 | 113215900 | L1MdTf_I | + | 5679 |
| chr8 | 113215900 | 113216916 | L1MdTf_I | + | 1016 |
| chr8 | 117542011 | 117542249 | L1MdTf_III | + | 238 |
| chr8 | 117542249 | 117543242 | L1MdTf_III | + | 993 |
| chr8 | 118704797 | 118710480 | L1MdTf_III | + | 5683 |
| chr8 | 118710480 | 118711473 | L1MdTf_III | + | 993 |
| chr8 | 129539797 | 129545213 | L1MdTf_I | + | 5416 |
| chr8 | 129545213 | 129546206 | L1MdTf_I | + | 993 |
| chr9 | 3145749 | 3151010 | L1MdTf_III | + | 5261 |
| chr9 | 3151010 | 3152003 | L1MdTf_III | + | 993 |
| chr9 | 3743533 | 3748762 | L1MdTf_III | + | 5229 |
| chr9 | 3748762 | 3749755 | L1MdTf_III | + | 993 |
| chr9 | 4182506 | 4188013 | L1MdTf_I | + | 5507 |
| chr9 | 4188013 | 4189006 | L1MdTf_I | + | 993 |
| chr9 | 7245661 | 7246698 | L1MdTf_III | + | 1037 |

| | | | | | |
|---|---|---|---|---|---|
| chr9 | 7246698 | 7247693 | L1MdTf_III | + | 995 |
| chr9 | 7485256 | 7487735 | L1MdTf_I | + | 2479 |
| chr9 | 7487735 | 7488731 | L1MdTf_I | + | 996 |
| chr9 | 8286168 | 8291322 | L1MdTf_III | + | 5154 |
| chr9 | 8291322 | 8292315 | L1MdTf_III | + | 993 |
| chr9 | 8466950 | 8473439 | L1MdTf_II | + | 6489 |
| chr9 | 8473439 | 8474432 | L1MdTf_II | + | 993 |
| chr9 | 9468271 | 9473512 | L1MdTf_III | + | 5241 |
| chr9 | 9473512 | 9474505 | L1MdTf_III | + | 993 |
| chr9 | 9495677 | 9496581 | L1MdTf_II | - | 904 |
| chr9 | 9496581 | 9497411 | L1MdTf_II | + | 830 |
| chr9 | 10964177 | 10969647 | L1MdTf_III | + | 5470 |
| chr9 | 10969647 | 10970640 | L1MdTf_III | + | 993 |
| chr9 | 11343733 | 11344250 | L1MdTf_III | + | 517 |
| chr9 | 11344250 | 11345244 | L1MdTf_III | + | 994 |
| chr9 | 11652039 | 11657343 | L1MdTf_III | + | 5304 |
| chr9 | 11657343 | 11658336 | L1MdTf_III | + | 993 |
| chr9 | 11722534 | 11727790 | L1MdTf_II | + | 5256 |
| chr9 | 11727790 | 11728783 | L1MdTf_II | + | 993 |
| chr9 | 12739978 | 12745836 | L1MdTf_II | + | 5858 |
| chr9 | 12745836 | 12746832 | L1MdTf_II | + | 996 |
| chr9 | 13036286 | 13041492 | L1MdTf_III | + | 5206 |
| chr9 | 13041492 | 13042482 | L1MdTf_III | + | 990 |
| chr9 | 13635834 | 13636463 | L1MdTf_III | + | 629 |
| chr9 | 13636463 | 13637456 | L1MdTf_III | + | 993 |
| chr9 | 16364009 | 16369220 | L1MdTf_I | + | 5211 |
| chr9 | 16369220 | 16370213 | L1MdTf_I | + | 993 |
| chr9 | 16517950 | 16523787 | L1MdTf_III | + | 5837 |
| chr9 | 16523787 | 16524777 | L1MdTf_III | + | 990 |
| chr9 | 16744948 | 16750605 | L1MdTf_I | + | 5657 |
| chr9 | 16750605 | 16751598 | L1MdTf_I | + | 993 |
| chr9 | 18039675 | 18039750 | L1MdTf_III | - | 75 |
| chr9 | 18039750 | 18040701 | L1MdTf_III | + | 951 |
| chr9 | 18067290 | 18072505 | L1MdTf_II | + | 5215 |
| chr9 | 18072505 | 18073498 | L1MdTf_II | + | 993 |
| chr9 | 18891696 | 18897759 | L1MdTf_II | + | 6063 |
| chr9 | 18897759 | 18898756 | L1MdTf_II | + | 997 |
| chr9 | 19199746 | 19205103 | L1MdTf_III | + | 5357 |
| chr9 | 19205103 | 19206096 | L1MdTf_III | + | 993 |
| chr9 | 20265932 | 20271398 | L1MdTf_III | + | 5466 |
| chr9 | 20271398 | 20272392 | L1MdTf_III | + | 994 |
| chr9 | 22435735 | 22436220 | L1MdTf_I | - | 485 |
| chr9 | 22436220 | 22437182 | L1MdTf_I | + | 962 |
| chr9 | 23892817 | 23898407 | L1MdTf_III | + | 5590 |
| chr9 | 23898407 | 23899400 | L1MdTf_III | + | 993 |

| | | | | | |
|---|---|---|---|---|---|
| chr9 | 24023904 | 24024742 | L1MdTf_II | - | 838 |
| chr9 | 24024742 | 24025443 | L1MdTf_II | + | 701 |
| chr9 | 24058641 | 24065311 | L1MdTf_III | + | 6670 |
| chr9 | 24065311 | 24066328 | L1MdTf_III | + | 1017 |
| chr9 | 24436844 | 24442060 | L1MdTf_III | + | 5216 |
| chr9 | 24442060 | 24443053 | L1MdTf_III | + | 993 |
| chr9 | 25144744 | 25150185 | L1MdTf_I | + | 5441 |
| chr9 | 25150185 | 25151178 | L1MdTf_I | + | 993 |
| chr9 | 26504018 | 26505133 | L1MdTf_I | + | 1115 |
| chr9 | 26505133 | 26506126 | L1MdTf_I | + | 993 |
| chr9 | 26882030 | 26882137 | L1MdTf_II | + | 107 |
| chr9 | 26882137 | 26883130 | L1MdTf_II | + | 993 |
| chr9 | 27288781 | 27289029 | L1MdTf_II | - | 248 |
| chr9 | 27289029 | 27289314 | L1MdTf_II | + | 285 |
| chr9 | 29939355 | 29940044 | L1MdTf_I | + | 689 |
| chr9 | 29940044 | 29941057 | L1MdTf_I | + | 1013 |
| chr9 | 33304583 | 33309865 | L1MdTf_III | + | 5282 |
| chr9 | 33309866 | 33310859 | L1MdTf_III | + | 993 |
| chr9 | 36146555 | 36152205 | L1MdTf_I | + | 5650 |
| chr9 | 36152205 | 36153218 | L1MdTf_I | + | 1013 |
| chr9 | 38152024 | 38157874 | L1MdTf_I | + | 5850 |
| chr9 | 38157874 | 38158867 | L1MdTf_I | + | 993 |
| chr9 | 39354533 | 39359940 | L1MdTf_II | + | 5407 |
| chr9 | 39359940 | 39360933 | L1MdTf_II | + | 993 |
| chr9 | 40403387 | 40408674 | L1MdTf_III | + | 5287 |
| chr9 | 40408674 | 40409666 | L1MdTf_III | + | 992 |
| chr9 | 42194003 | 42194185 | L1MdTf_III | - | 182 |
| chr9 | 42194186 | 42194526 | L1MdTf_III | + | 340 |
| chr9 | 52311628 | 52311880 | L1MdTf_III | - | 252 |
| chr9 | 52311880 | 52312014 | L1MdTf_III | + | 134 |
| chr9 | 52661029 | 52666445 | L1MdTf_I | + | 5416 |
| chr9 | 52666445 | 52667438 | L1MdTf_I | + | 993 |
| chr9 | 52796777 | 52797205 | L1MdTf_III | + | 428 |
| chr9 | 52797205 | 52798198 | L1MdTf_III | + | 993 |
| chr9 | 52807214 | 52809128 | L1MdTf_III | + | 1914 |
| chr9 | 52809129 | 52810122 | L1MdTf_III | + | 993 |
| chr9 | 52924782 | 52929923 | L1MdTf_II | + | 5141 |
| chr9 | 52929923 | 52930916 | L1MdTf_II | + | 993 |
| chr9 | 55735927 | 55741422 | L1MdTf_I | + | 5495 |
| chr9 | 55741422 | 55742415 | L1MdTf_I | + | 993 |
| chr9 | 57315279 | 57316190 | L1MdTf_III | - | 911 |
| chr9 | 57316190 | 57316671 | L1MdTf_III | + | 481 |
| chr9 | 58593068 | 58593552 | L1MdTf_II | - | 484 |
| chr9 | 58593552 | 58594286 | L1MdTf_II | + | 734 |
| chr9 | 58856930 | 58857486 | L1MdTf_I | - | 556 |

| | | | | | |
|---|---|---|---|---|---|
| chr9 | 58857486 | 58858501 | L1MdTf_I | + | 1015 |
| chr9 | 59952782 | 59958079 | L1MdTf_II | + | 5297 |
| chr9 | 59958079 | 59959075 | L1MdTf_II | + | 996 |
| chr9 | 68066158 | 68066596 | L1MdTf_II | + | 438 |
| chr9 | 68066596 | 68067640 | L1MdTf_II | + | 1044 |
| chr9 | 68401248 | 68406343 | L1MdTf_III | + | 5095 |
| chr9 | 68406343 | 68407336 | L1MdTf_III | + | 993 |
| chr9 | 68986868 | 68987609 | L1MdTf_II | - | 741 |
| chr9 | 68987609 | 68988171 | L1MdTf_II | + | 562 |
| chr9 | 70811016 | 70816452 | L1MdTf_II | + | 5436 |
| chr9 | 70816452 | 70817445 | L1MdTf_II | + | 993 |
| chr9 | 71690042 | 71690342 | L1MdTf_II | + | 300 |
| chr9 | 71690342 | 71691335 | L1MdTf_II | + | 993 |
| chr9 | 73595546 | 73596307 | L1MdTf_I | - | 761 |
| chr9 | 73596307 | 73597030 | L1MdTf_I | + | 723 |
| chr9 | 73842837 | 73848298 | L1MdTf_III | + | 5461 |
| chr9 | 73848298 | 73849289 | L1MdTf_III | + | 991 |
| chr9 | 74675863 | 74681033 | L1MdTf_III | + | 5170 |
| chr9 | 74681033 | 74682024 | L1MdTf_III | + | 991 |
| chr9 | 77285741 | 77285980 | L1MdTf_I | + | 239 |
| chr9 | 77285980 | 77286973 | L1MdTf_I | + | 993 |
| chr9 | 79371058 | 79372145 | L1MdTf_II | + | 1087 |
| chr9 | 79372145 | 79373138 | L1MdTf_II | + | 993 |
| chr9 | 79375444 | 79377077 | L1MdTf_II | - | 1633 |
| chr9 | 79377077 | 79378070 | L1MdTf_II | + | 993 |
| chr9 | 81064707 | 81070194 | L1MdTf_III | + | 5487 |
| chr9 | 81070194 | 81071187 | L1MdTf_III | + | 993 |
| chr9 | 81180394 | 81184055 | L1MdTf_III | + | 3661 |
| chr9 | 81184055 | 81185048 | L1MdTf_III | + | 993 |
| chr9 | 82171267 | 82172081 | L1MdTf_II | - | 814 |
| chr9 | 82172081 | 82173177 | L1MdTf_II | + | 1096 |
| chr9 | 83342648 | 83348057 | L1MdTf_III | + | 5409 |
| chr9 | 83348057 | 83349049 | L1MdTf_III | + | 992 |
| chr9 | 83874521 | 83879962 | L1MdTf_III | + | 5441 |
| chr9 | 83879962 | 83880955 | L1MdTf_III | + | 993 |
| chr9 | 84411833 | 84412091 | L1MdTf_I | + | 258 |
| chr9 | 84412091 | 84413083 | L1MdTf_I | + | 992 |
| chr9 | 84649269 | 84649510 | L1MdTf_II | - | 241 |
| chr9 | 84649510 | 84650630 | L1MdTf_II | + | 1120 |
| chr9 | 85508214 | 85513959 | L1MdTf_III | + | 5745 |
| chr9 | 85513959 | 85514952 | L1MdTf_III | + | 993 |
| chr9 | 85780926 | 85786583 | L1MdTf_II | + | 5657 |
| chr9 | 85786583 | 85787576 | L1MdTf_II | + | 993 |
| chr9 | 86002165 | 86007739 | L1MdTf_III | + | 5574 |
| chr9 | 86007739 | 86008732 | L1MdTf_III | + | 993 |

| | | | | | |
|---|---|---|---|---|---|
| chr9 | 86988097 | 86993413 | L1MdTf_III | + | 5316 |
| chr9 | 86993413 | 86994409 | L1MdTf_III | + | 996 |
| chr9 | 87669954 | 87672288 | L1MdTf_II | + | 2334 |
| chr9 | 87672288 | 87673281 | L1MdTf_II | + | 993 |
| chr9 | 87697229 | 87702659 | L1MdTf_II | + | 5430 |
| chr9 | 87702659 | 87703675 | L1MdTf_II | + | 1016 |
| chr9 | 88922839 | 88928325 | L1MdTf_III | + | 5486 |
| chr9 | 88928325 | 88929316 | L1MdTf_III | + | 991 |
| chr9 | 91143309 | 91149259 | L1MdTf_I | + | 5950 |
| chr9 | 91149259 | 91150252 | L1MdTf_I | + | 993 |
| chr9 | 91500847 | 91506452 | L1MdTf_I | + | 5605 |
| chr9 | 91506452 | 91507445 | L1MdTf_I | + | 993 |
| chr9 | 94381362 | 94385381 | L1MdTf_I | - | 4019 |
| chr9 | 94385381 | 94386000 | L1MdTf_I | + | 619 |
| chr9 | 95174111 | 95180662 | L1MdTf_I | + | 6551 |
| chr9 | 95180662 | 95181655 | L1MdTf_I | + | 993 |
| chr9 | 95963753 | 95965807 | L1MdTf_I | + | 2054 |
| chr9 | 95965807 | 95966800 | L1MdTf_I | + | 993 |
| chr9 | 97212685 | 97218249 | L1MdTf_III | + | 5564 |
| chr9 | 97218250 | 97219241 | L1MdTf_III | + | 991 |
| chr9 | 98132107 | 98138810 | L1MdTf_I | + | 6703 |
| chr9 | 98138810 | 98139803 | L1MdTf_I | + | 993 |
| chr9 | 99955880 | 99961730 | L1MdTf_II | + | 5850 |
| chr9 | 99961730 | 99962723 | L1MdTf_II | + | 993 |
| chr9 | 101304688 | 101310963 | L1MdTf_I | + | 6275 |
| chr9 | 101310963 | 101311956 | L1MdTf_I | + | 993 |
| chr9 | 106980965 | 106982002 | L1MdTf_III | + | 1037 |
| chr9 | 106982002 | 106982989 | L1MdTf_III | + | 987 |
| chr9 | 107087799 | 107088517 | L1MdTf_III | + | 718 |
| chr9 | 107088517 | 107089510 | L1MdTf_III | + | 993 |
| chr9 | 111409285 | 111409987 | L1MdTf_I | - | 702 |
| chr9 | 111409987 | 111410034 | L1MdTf_I | + | 47 |
| chr9 | 112035525 | 112035710 | L1MdTf_II | + | 185 |
| chr9 | 112035710 | 112036699 | L1MdTf_II | + | 989 |
| chr9 | 112490284 | 112495553 | L1MdTf_III | + | 5269 |
| chr9 | 112495553 | 112496546 | L1MdTf_III | + | 993 |
| chr9 | 112787925 | 112793786 | L1MdTf_II | + | 5861 |
| chr9 | 112793786 | 112794779 | L1MdTf_II | + | 993 |
| chr9 | 114112493 | 114117983 | L1MdTf_II | + | 5490 |
| chr9 | 114117983 | 114118976 | L1MdTf_II | + | 993 |
| chr9 | 119449004 | 119450541 | L1MdTf_I | + | 1537 |
| chr9 | 119450541 | 119451534 | L1MdTf_I | + | 993 |
| chrX | 4395515 | 4396206 | L1MdTf_II | + | 691 |
| chrX | 4396206 | 4397202 | L1MdTf_II | + | 996 |
| chrX | 5589719 | 5590507 | L1MdTf_III | - | 788 |

| | | | | | |
|---|---|---|---|---|---|
| chrX | 5590508 | 5591096 | L1MdTf_III | + | 588 |
| chrX | 5739900 | 5741494 | L1MdTf_I | + | 1594 |
| chrX | 5741494 | 5742487 | L1MdTf_I | + | 993 |
| chrX | 6856919 | 6857557 | L1MdTf_III | + | 638 |
| chrX | 6857557 | 6858550 | L1MdTf_III | + | 993 |
| chrX | 7088697 | 7090607 | L1MdTf_II | + | 1910 |
| chrX | 7090607 | 7091628 | L1MdTf_II | + | 1021 |
| chrX | 8622827 | 8628093 | L1MdTf_III | + | 5266 |
| chrX | 8628093 | 8629076 | L1MdTf_III | + | 983 |
| chrX | 8685197 | 8690401 | L1MdTf_III | + | 5204 |
| chrX | 8690401 | 8691394 | L1MdTf_III | + | 993 |
| chrX | 8845580 | 8847058 | L1MdTf_III | + | 1478 |
| chrX | 8847058 | 8848051 | L1MdTf_III | + | 993 |
| chrX | 9832183 | 9837655 | L1MdTf_III | + | 5472 |
| chrX | 9837655 | 9838660 | L1MdTf_III | + | 1005 |
| chrX | 10391031 | 10396525 | L1MdTf_II | + | 5494 |
| chrX | 10396525 | 10397545 | L1MdTf_II | + | 1020 |
| chrX | 10831461 | 10837948 | L1MdTf_II | + | 6487 |
| chrX | 10837948 | 10838945 | L1MdTf_II | + | 997 |
| chrX | 14276838 | 14277363 | L1MdTf_I | + | 525 |
| chrX | 14277363 | 14278356 | L1MdTf_I | + | 993 |
| chrX | 14436507 | 14443679 | L1MdTf_II | + | 7172 |
| chrX | 14443679 | 14444676 | L1MdTf_II | + | 997 |
| chrX | 18371668 | 18377149 | L1MdTf_III | + | 5481 |
| chrX | 18377149 | 18378142 | L1MdTf_III | + | 993 |
| chrX | 18404677 | 18410926 | L1MdTf_I | + | 6249 |
| chrX | 18410926 | 18411919 | L1MdTf_I | + | 993 |
| chrX | 18491544 | 18497142 | L1MdTf_III | + | 5598 |
| chrX | 18497142 | 18498135 | L1MdTf_III | + | 993 |
| chrX | 18663003 | 18668793 | L1MdTf_I | + | 5790 |
| chrX | 18668793 | 18669786 | L1MdTf_I | + | 993 |
| chrX | 19517523 | 19522907 | L1MdTf_I | + | 5384 |
| chrX | 19522907 | 19523919 | L1MdTf_I | + | 1012 |
| chrX | 19697279 | 19703342 | L1MdTf_II | + | 6063 |
| chrX | 19703342 | 19704338 | L1MdTf_II | + | 996 |
| chrX | 21103031 | 21105184 | L1MdTf_III | - | 2153 |
| chrX | 21105184 | 21112305 | L1MdTf_I | - | 7121 |
| chrX | 21332722 | 21338147 | L1MdTf_I | + | 5425 |
| chrX | 21338147 | 21338590 | L1MdTf_I | + | 443 |
| chrX | 21731347 | 21736833 | L1MdTf_II | + | 5486 |
| chrX | 21736833 | 21737825 | L1MdTf_II | + | 992 |
| chrX | 23401051 | 23407132 | L1MdTf_I | + | 6081 |
| chrX | 23407133 | 23408125 | L1MdTf_I | + | 992 |
| chrX | 24105669 | 24105756 | L1MdTf_II | - | 87 |
| chrX | 24105756 | 24106224 | L1MdTf_II | + | 468 |

| | | | | | |
|---|---|---|---|---|---|
| chrX | 24387587 | 24392815 | L1MdTf_I | + | 5228 |
| chrX | 24392815 | 24393808 | L1MdTf_I | + | 993 |
| chrX | 24399125 | 24399196 | L1MdTf_III | + | 71 |
| chrX | 24399196 | 24400190 | L1MdTf_III | + | 994 |
| chrX | 24663903 | 24664409 | L1MdTf_II | + | 506 |
| chrX | 24664409 | 24665406 | L1MdTf_II | + | 997 |
| chrX | 24864715 | 24869351 | L1MdTf_II | - | 4636 |
| chrX | 24869351 | 24869726 | L1MdTf_II | + | 375 |
| chrX | 25888050 | 25893506 | L1MdTf_III | + | 5456 |
| chrX | 25893506 | 25894498 | L1MdTf_III | + | 992 |
| chrX | 26159643 | 26165089 | L1MdTf_I | + | 5446 |
| chrX | 26165089 | 26166101 | L1MdTf_I | + | 1012 |
| chrX | 27338891 | 27339266 | L1MdTf_II | - | 375 |
| chrX | 27339266 | 27343916 | L1MdTf_II | + | 4650 |
| chrX | 27685594 | 27685969 | L1MdTf_II | - | 375 |
| chrX | 27685969 | 27690832 | L1MdTf_II | + | 4863 |
| chrX | 28346570 | 28346945 | L1MdTf_II | - | 375 |
| chrX | 28346945 | 28351807 | L1MdTf_II | + | 4862 |
| chrX | 28693382 | 28693757 | L1MdTf_II | - | 375 |
| chrX | 28693757 | 28698620 | L1MdTf_II | + | 4863 |
| chrX | 29761900 | 29766762 | L1MdTf_II | - | 4862 |
| chrX | 29766762 | 29767137 | L1MdTf_II | + | 375 |
| chrX | 30003224 | 30003274 | L1MdTf_III | + | 50 |
| chrX | 30003275 | 30004268 | L1MdTf_III | + | 993 |
| chrX | 32082368 | 32087977 | L1MdTf_I | + | 5609 |
| chrX | 32087977 | 32088970 | L1MdTf_I | + | 993 |
| chrX | 32991660 | 32997340 | L1MdTf_I | + | 5680 |
| chrX | 32997340 | 32998333 | L1MdTf_I | + | 993 |
| chrX | 33666675 | 33671550 | L1MdTf_I | + | 4875 |
| chrX | 33671550 | 33672543 | L1MdTf_I | + | 993 |
| chrX | 34367570 | 34372433 | L1MdTf_II | - | 4863 |
| chrX | 34372433 | 34372808 | L1MdTf_II | + | 375 |
| chrX | 34592244 | 34592294 | L1MdTf_III | + | 50 |
| chrX | 34592295 | 34593288 | L1MdTf_III | + | 993 |
| chrX | 35086191 | 35092126 | L1MdTf_II | + | 5935 |
| chrX | 35092126 | 35093119 | L1MdTf_II | + | 993 |
| chrX | 35705793 | 35710234 | L1MdTf_III | + | 4441 |
| chrX | 35710234 | 35711227 | L1MdTf_III | + | 993 |
| chrX | 36328464 | 36328873 | L1MdTf_III | + | 409 |
| chrX | 36328873 | 36329832 | L1MdTf_III | + | 959 |
| chrX | 37000000 | 37004240 | L1MdTf_II | + | 4240 |
| chrX | 37004240 | 37005229 | L1MdTf_II | + | 989 |
| chrX | 37877982 | 37878636 | L1MdTf_II | + | 654 |
| chrX | 37878636 | 37879629 | L1MdTf_I | + | 993 |
| chrX | 38294331 | 38300010 | L1MdTf_III | + | 5679 |

| | | | | | |
|---|---|---|---|---|---|
| chrX | 38300010 | 38301003 | L1MdTf_III | + | 993 |
| chrX | 38420730 | 38426581 | L1MdTf_I | + | 5851 |
| chrX | 38426581 | 38427574 | L1MdTf_I | + | 993 |
| chrX | 39117136 | 39117347 | L1MdTf_III | - | 211 |
| chrX | 39117347 | 39117385 | L1MdTf_II | + | 38 |
| chrX | 39922226 | 39928090 | L1MdTf_I | + | 5864 |
| chrX | 39928090 | 39929112 | L1MdTf_I | + | 1022 |
| chrX | 39999875 | 40000000 | L1MdTf_III | + | 125 |
| chrX | 40000000 | 40001828 | L1MdTf_III | + | 1828 |
| chrX | 40320625 | 40326385 | L1MdTf_I | + | 5760 |
| chrX | 40326385 | 40327378 | L1MdTf_I | + | 993 |
| chrX | 40571338 | 40577402 | L1MdTf_II | + | 6064 |
| chrX | 40577402 | 40578418 | L1MdTf_II | + | 1016 |
| chrX | 40833139 | 40838594 | L1MdTf_III | + | 5455 |
| chrX | 40838594 | 40839587 | L1MdTf_III | + | 993 |
| chrX | 41721830 | 41727011 | L1MdTf_II | + | 5181 |
| chrX | 41727011 | 41728005 | L1MdTf_II | + | 994 |
| chrX | 41850018 | 41855461 | L1MdTf_II | + | 5443 |
| chrX | 41855461 | 41856480 | L1MdTf_II | + | 1019 |
| chrX | 42297672 | 42303541 | L1MdTf_I | + | 5869 |
| chrX | 42303541 | 42304534 | L1MdTf_I | + | 993 |
| chrX | 43105927 | 43107072 | L1MdTf_III | + | 1145 |
| chrX | 43107072 | 43108065 | L1MdTf_III | + | 993 |
| chrX | 43132343 | 43137899 | L1MdTf_I | + | 5556 |
| chrX | 43137899 | 43138892 | L1MdTf_I | + | 993 |
| chrX | 43578100 | 43583720 | L1MdTf_I | + | 5620 |
| chrX | 43583720 | 43584713 | L1MdTf_I | + | 993 |
| chrX | 43681763 | 43682141 | L1MdTf_II | + | 378 |
| chrX | 43682141 | 43683135 | L1MdTf_II | + | 994 |
| chrX | 44100047 | 44105684 | L1MdTf_I | + | 5637 |
| chrX | 44105684 | 44106676 | L1MdTf_I | + | 992 |
| chrX | 44190737 | 44194776 | L1MdTf_III | + | 4039 |
| chrX | 44194776 | 44195760 | L1MdTf_III | + | 984 |
| chrX | 44318182 | 44323429 | L1MdTf_III | + | 5247 |
| chrX | 44323429 | 44324415 | L1MdTf_III | + | 986 |
| chrX | 45619887 | 45620059 | L1MdTf_III | - | 172 |
| chrX | 45620059 | 45620565 | L1MdTf_III | + | 506 |
| chrX | 46066383 | 46071738 | L1MdTf_II | + | 5355 |
| chrX | 46071738 | 46072728 | L1MdTf_II | + | 990 |
| chrX | 46444690 | 46449989 | L1MdTf_II | + | 5299 |
| chrX | 46449989 | 46450979 | L1MdTf_II | + | 990 |
| chrX | 46825299 | 46830722 | L1MdTf_II | + | 5423 |
| chrX | 46830722 | 46831715 | L1MdTf_II | + | 993 |
| chrX | 46955252 | 46961540 | L1MdTf_I | + | 6288 |
| chrX | 46961540 | 46962533 | L1MdTf_I | + | 993 |

| | | | | | |
|---|---|---|---|---|---|
| chrX | 47827561 | 47832652 | L1MdTf_III | + | 5091 |
| chrX | 47832652 | 47833645 | L1MdTf_III | + | 993 |
| chrX | 48445271 | 48450821 | L1MdTf_I | + | 5550 |
| chrX | 48450821 | 48451814 | L1MdTf_I | + | 993 |
| chrX | 48451877 | 48453037 | L1MdTf_I | + | 1160 |
| chrX | 48453037 | 48454030 | L1MdTf_I | + | 993 |
| chrX | 48695681 | 48696960 | L1MdTf_II | - | 1279 |
| chrX | 48696960 | 48702907 | L1MdTf_II | - | 5947 |
| chrX | 49086880 | 49092491 | L1MdTf_II | + | 5611 |
| chrX | 49092491 | 49093484 | L1MdTf_II | + | 993 |
| chrX | 49198436 | 49198556 | L1MdTf_II | - | 120 |
| chrX | 49198557 | 49198810 | L1MdTf_II | + | 253 |
| chrX | 49473587 | 49474735 | L1MdTf_III | + | 1148 |
| chrX | 49474735 | 49475727 | L1MdTf_III | + | 992 |
| chrX | 49475731 | 49476891 | L1MdTf_III | + | 1160 |
| chrX | 49476891 | 49477914 | L1MdTf_III | + | 1023 |
| chrX | 50818400 | 50823625 | L1MdTf_II | + | 5225 |
| chrX | 50823625 | 50824618 | L1MdTf_II | + | 993 |
| chrX | 51005012 | 51006714 | L1MdTf_II | + | 1702 |
| chrX | 51006714 | 51007707 | L1MdTf_II | + | 993 |
| chrX | 54472874 | 54474017 | L1MdTf_II | + | 1143 |
| chrX | 54474017 | 54474992 | L1MdTf_II | + | 975 |
| chrX | 55333994 | 55334514 | L1MdTf_II | - | 520 |
| chrX | 55334514 | 55334735 | L1MdTf_II | + | 221 |
| chrX | 55455659 | 55458887 | L1MdTf_I | + | 3228 |
| chrX | 55458887 | 55459880 | L1MdTf_I | + | 993 |
| chrX | 55926183 | 55932022 | L1MdTf_II | + | 5839 |
| chrX | 55932022 | 55933015 | L1MdTf_II | + | 993 |
| chrX | 56688091 | 56693322 | L1MdTf_I | + | 5231 |
| chrX | 56693322 | 56694315 | L1MdTf_I | + | 993 |
| chrX | 57333802 | 57339043 | L1MdTf_III | + | 5241 |
| chrX | 57339043 | 57340036 | L1MdTf_III | + | 993 |
| chrX | 57906982 | 57912624 | L1MdTf_I | + | 5642 |
| chrX | 57912624 | 57913617 | L1MdTf_I | + | 993 |
| chrX | 58840057 | 58841265 | L1MdTf_II | + | 1208 |
| chrX | 58841265 | 58842258 | L1MdTf_II | + | 993 |
| chrX | 58900074 | 58900372 | L1MdTf_III | + | 298 |
| chrX | 58900372 | 58901365 | L1MdTf_III | + | 993 |
| chrX | 59246213 | 59251429 | L1MdTf_III | + | 5216 |
| chrX | 59251430 | 59252411 | L1MdTf_III | + | 981 |
| chrX | 60490826 | 60491743 | L1MdTf_III | + | 917 |
| chrX | 60491743 | 60492736 | L1MdTf_III | + | 993 |
| chrX | 61240934 | 61246866 | L1MdTf_I | + | 5932 |
| chrX | 61246866 | 61247859 | L1MdTf_I | + | 993 |
| chrX | 61660635 | 61665883 | L1MdTf_II | + | 5248 |

| | | | | | |
|---|---|---|---|---|---|
| chrX | 61665883 | 61666876 | L1MdTf_II | + | 993 |
| chrX | 61674713 | 61675557 | L1MdTf_I | + | 844 |
| chrX | 61675557 | 61676550 | L1MdTf_I | + | 993 |
| chrX | 61884240 | 61889845 | L1MdTf_II | + | 5605 |
| chrX | 61889845 | 61890838 | L1MdTf_II | + | 993 |
| chrX | 61989933 | 61993378 | L1MdTf_I | + | 3445 |
| chrX | 61993378 | 61994371 | L1MdTf_I | + | 993 |
| chrX | 62678147 | 62683614 | L1MdTf_III | + | 5467 |
| chrX | 62683614 | 62684607 | L1MdTf_III | + | 993 |
| chrX | 62989559 | 62995179 | L1MdTf_I | + | 5620 |
| chrX | 62995179 | 62996172 | L1MdTf_I | + | 993 |
| chrX | 63169587 | 63174819 | L1MdTf_I | + | 5232 |
| chrX | 63174819 | 63175812 | L1MdTf_I | + | 993 |
| chrX | 63304094 | 63304321 | L1MdTf_I | + | 227 |
| chrX | 63304321 | 63305314 | L1MdTf_I | + | 993 |
| chrX | 64125994 | 64126354 | L1MdTf_III | - | 360 |
| chrX | 64126354 | 64126547 | L1MdTf_III | + | 193 |
| chrX | 64200884 | 64203085 | L1MdTf_III | + | 2201 |
| chrX | 64203085 | 64204077 | L1MdTf_III | + | 992 |
| chrX | 65374925 | 65380369 | L1MdTf_II | + | 5444 |
| chrX | 65380369 | 65381362 | L1MdTf_II | + | 993 |
| chrX | 65495538 | 65500000 | L1MdTf_II | + | 4462 |
| chrX | 65500000 | 65500961 | L1MdTf_I | + | 961 |
| chrX | 65612104 | 65612735 | L1MdTf_I | + | 631 |
| chrX | 65612735 | 65613728 | L1MdTf_I | + | 993 |
| chrX | 66075418 | 66080294 | L1MdTf_II | + | 4876 |
| chrX | 66080294 | 66081278 | L1MdTf_II | + | 984 |
| chrX | 66268336 | 66269428 | L1MdTf_II | - | 1092 |
| chrX | 66269428 | 66269858 | L1MdTf_II | + | 430 |
| chrX | 66495001 | 66496912 | L1MdTf_II | - | 1911 |
| chrX | 66496912 | 66497600 | L1MdTf_II | + | 688 |
| chrX | 67307502 | 67312764 | L1MdTf_III | + | 5262 |
| chrX | 67312765 | 67313758 | L1MdTf_III | + | 993 |
| chrX | 67387331 | 67388545 | L1MdTf_II | + | 1214 |
| chrX | 67388545 | 67389538 | L1MdTf_II | + | 993 |
| chrX | 68290043 | 68295534 | L1MdTf_III | + | 5491 |
| chrX | 68295534 | 68296518 | L1MdTf_III | + | 984 |
| chrX | 69098915 | 69104543 | L1MdTf_I | + | 5628 |
| chrX | 69104544 | 69105537 | L1MdTf_I | + | 993 |
| chrX | 69903898 | 69909037 | L1MdTf_III | + | 5139 |
| chrX | 69909037 | 69909079 | L1MdTf_III | + | 42 |
| chrX | 69909079 | 69909424 | L1MdTf_III | + | 345 |
| chrX | 69909424 | 69910417 | L1MdTf_III | + | 993 |
| chrX | 71187407 | 71192658 | L1MdTf_III | + | 5251 |
| chrX | 71192658 | 71193651 | L1MdTf_III | + | 993 |

| | | | | | |
|---|---|---|---|---|---|
| chrX | 71616459 | 71621594 | L1MdTf_II | + | 5135 |
| chrX | 71621595 | 71627658 | L1MdTf_I | + | 6063 |
| chrX | 71628661 | 71629086 | L1MdTf_II | + | 425 |
| chrX | 71629086 | 71630079 | L1MdTf_II | + | 993 |
| chrX | 73976609 | 73981864 | L1MdTf_II | + | 5255 |
| chrX | 73981864 | 73982858 | L1MdTf_II | + | 994 |
| chrX | 74346486 | 74352744 | L1MdTf_II | + | 6258 |
| chrX | 74352744 | 74353728 | L1MdTf_II | + | 984 |
| chrX | 74633183 | 74634188 | L1MdTf_I | - | 1005 |
| chrX | 74634188 | 74634645 | L1MdTf_I | + | 457 |
| chrX | 74994739 | 75000000 | L1MdTf_III | + | 5261 |
| chrX | 75000000 | 75001013 | L1MdTf_III | + | 1013 |
| chrX | 75280078 | 75285471 | L1MdTf_II | + | 5393 |
| chrX | 75285471 | 75286469 | L1MdTf_II | + | 998 |
| chrX | 75405682 | 75411944 | L1MdTf_I | + | 6262 |
| chrX | 75411944 | 75412937 | L1MdTf_I | + | 993 |
| chrX | 75507743 | 75513128 | L1MdTf_III | + | 5385 |
| chrX | 75513128 | 75514121 | L1MdTf_III | + | 993 |
| chrX | 75994424 | 75995628 | L1MdTf_II | + | 1204 |
| chrX | 75995628 | 75996625 | L1MdTf_II | + | 997 |
| chrX | 76301430 | 76307278 | L1MdTf_I | + | 5848 |
| chrX | 76307278 | 76308271 | L1MdTf_I | + | 993 |
| chrX | 76309229 | 76315076 | L1MdTf_I | + | 5847 |
| chrX | 76315076 | 76316069 | L1MdTf_I | + | 993 |
| chrX | 76316079 | 76316567 | L1MdTf_III | + | 488 |
| chrX | 76316567 | 76317559 | L1MdTf_III | + | 992 |
| chrX | 76358884 | 76364723 | L1MdTf_III | + | 5839 |
| chrX | 76364723 | 76365716 | L1MdTf_III | + | 993 |
| chrX | 76441365 | 76447022 | L1MdTf_I | + | 5657 |
| chrX | 76447022 | 76448044 | L1MdTf_I | + | 1022 |
| chrX | 76727350 | 76729575 | L1MdTf_III | - | 2225 |
| chrX | 76729575 | 76730386 | L1MdTf_I | - | 811 |
| chrX | 76922100 | 76922775 | L1MdTf_III | + | 675 |
| chrX | 76922776 | 76923769 | L1MdTf_III | + | 993 |
| chrX | 77012257 | 77018084 | L1MdTf_II | + | 5827 |
| chrX | 77018084 | 77019077 | L1MdTf_II | + | 993 |
| chrX | 77274464 | 77280170 | L1MdTf_II | + | 5706 |
| chrX | 77280170 | 77281154 | L1MdTf_II | + | 984 |
| chrX | 77356927 | 77357130 | L1MdTf_II | + | 203 |
| chrX | 77357130 | 77358145 | L1MdTf_II | + | 1015 |
| chrX | 77379489 | 77380653 | L1MdTf_II | + | 1164 |
| chrX | 77380653 | 77381637 | L1MdTf_II | + | 984 |
| chrX | 77467259 | 77467597 | L1MdTf_I | + | 338 |
| chrX | 77467598 | 77468615 | L1MdTf_I | + | 1017 |
| chrX | 78863169 | 78868844 | L1MdTf_II | + | 5675 |

| | | | | | |
|---|---|---|---|---|---|
| chrX | 78868844 | 78869859 | L1MdTf_II | + | 1015 |
| chrX | 79933461 | 79938888 | L1MdTf_I | + | 5427 |
| chrX | 79938888 | 79939881 | L1MdTf_I | + | 993 |
| chrX | 80286747 | 80292029 | L1MdTf_I | + | 5282 |
| chrX | 80292029 | 80293023 | L1MdTf_I | + | 994 |
| chrX | 81698333 | 81703882 | L1MdTf_II | + | 5549 |
| chrX | 81703882 | 81704879 | L1MdTf_II | + | 997 |
| chrX | 81978267 | 81983924 | L1MdTf_III | + | 5657 |
| chrX | 81983924 | 81984917 | L1MdTf_III | + | 993 |
| chrX | 83384890 | 83386356 | L1MdTf_III | + | 1466 |
| chrX | 83386356 | 83387347 | L1MdTf_III | + | 991 |
| chrX | 83851751 | 83856930 | L1MdTf_III | + | 5179 |
| chrX | 83856930 | 83857924 | L1MdTf_III | + | 994 |
| chrX | 83894667 | 83895355 | L1MdTf_II | - | 688 |
| chrX | 83895355 | 83896451 | L1MdTf_II | + | 1096 |
| chrX | 83897878 | 83897926 | L1MdTf_I | - | 48 |
| chrX | 83897926 | 83898565 | L1MdTf_I | + | 639 |
| chrX | 86100289 | 86100498 | L1MdTf_I | + | 209 |
| chrX | 86100498 | 86101527 | L1MdTf_I | + | 1029 |
| chrX | 86882117 | 86887840 | L1MdTf_II | + | 5723 |
| chrX | 86887840 | 86888833 | L1MdTf_II | + | 993 |
| chrX | 88247225 | 88248754 | L1MdTf_III | + | 1529 |
| chrX | 88248754 | 88249747 | L1MdTf_III | + | 993 |
| chrX | 88531690 | 88537322 | L1MdTf_I | + | 5632 |
| chrX | 88537322 | 88538315 | L1MdTf_I | + | 993 |
| chrX | 88538498 | 88542125 | L1MdTf_III | + | 3627 |
| chrX | 88542126 | 88543118 | L1MdTf_III | + | 992 |
| chrX | 89044649 | 89044994 | L1MdTf_II | - | 345 |
| chrX | 89044994 | 89045227 | L1MdTf_II | + | 233 |
| chrX | 89373561 | 89379223 | L1MdTf_I | + | 5662 |
| chrX | 89379223 | 89380216 | L1MdTf_I | + | 993 |
| chrX | 90056193 | 90056631 | L1MdTf_III | + | 438 |
| chrX | 90056631 | 90057624 | L1MdTf_III | + | 993 |
| chrX | 90652332 | 90657564 | L1MdTf_II | + | 5232 |
| chrX | 90657565 | 90658558 | L1MdTf_II | + | 993 |
| chrX | 91611862 | 91614416 | L1MdTf_II | + | 2554 |
| chrX | 91614416 | 91615409 | L1MdTf_II | + | 993 |
| chrX | 91658445 | 91663849 | L1MdTf_III | + | 5404 |
| chrX | 91663850 | 91664842 | L1MdTf_III | + | 992 |
| chrX | 91755849 | 91761227 | L1MdTf_III | + | 5378 |
| chrX | 91761228 | 91762220 | L1MdTf_III | + | 992 |
| chrX | 91844243 | 91849872 | L1MdTf_I | + | 5629 |
| chrX | 91849872 | 91850865 | L1MdTf_I | + | 993 |
| chrX | 91987027 | 91992718 | L1MdTf_II | + | 5691 |
| chrX | 91992718 | 91993711 | L1MdTf_II | + | 993 |

| | | | | | |
|---|---|---|---|---|---|
| chrX | 95422881 | 95428178 | L1MdTf_II | + | 5297 |
| chrX | 95428178 | 95429171 | L1MdTf_II | + | 993 |
| chrX | 95436079 | 95436799 | L1MdTf_II | + | 720 |
| chrX | 95436799 | 95442024 | L1MdTf_II | + | 5225 |
| chrX | 95443023 | 95447518 | L1MdTf_II | + | 4495 |
| chrX | 95447518 | 95448511 | L1MdTf_II | + | 993 |
| chrX | 95672553 | 95673633 | L1MdTf_III | + | 1080 |
| chrX | 95673633 | 95674625 | L1MdTf_III | + | 992 |
| chrX | 96212726 | 96213585 | L1MdTf_III | - | 859 |
| chrX | 96213585 | 96214456 | L1MdTf_III | + | 871 |
| chrX | 96771512 | 96777353 | L1MdTf_II | + | 5841 |
| chrX | 96777353 | 96778337 | L1MdTf_II | + | 984 |
| chrX | 96877216 | 96882709 | L1MdTf_III | + | 5493 |
| chrX | 96882709 | 96883701 | L1MdTf_III | + | 992 |
| chrX | 97154800 | 97158194 | L1MdTf_II | + | 3394 |
| chrX | 97158194 | 97159189 | L1MdTf_II | + | 995 |
| chrX | 97266179 | 97271784 | L1MdTf_I | + | 5605 |
| chrX | 97271784 | 97272777 | L1MdTf_I | + | 993 |
| chrX | 99338314 | 99344375 | L1MdTf_II | + | 6061 |
| chrX | 99344375 | 99345388 | L1MdTf_II | + | 1013 |
| chrX | 103216082 | 103216736 | L1MdTf_II | + | 654 |
| chrX | 103216736 | 103217729 | L1MdTf_II | + | 993 |
| chrX | 103666250 | 103671640 | L1MdTf_II | + | 5390 |
| chrX | 103671640 | 103672637 | L1MdTf_II | + | 997 |
| chrX | 103764968 | 103770414 | L1MdTf_III | + | 5446 |
| chrX | 103770414 | 103771416 | L1MdTf_III | + | 1002 |
| chrX | 103841940 | 103847238 | L1MdTf_I | + | 5298 |
| chrX | 103847238 | 103848231 | L1MdTf_I | + | 993 |
| chrX | 103905186 | 103910996 | L1MdTf_I | + | 5810 |
| chrX | 103910996 | 103911989 | L1MdTf_I | + | 993 |
| chrX | 103911991 | 103912335 | L1MdTf_II | + | 344 |
| chrX | 103912335 | 103913328 | L1MdTf_II | + | 993 |
| chrX | 104003244 | 104008883 | L1MdTf_II | + | 5639 |
| chrX | 104008883 | 104009876 | L1MdTf_II | + | 993 |
| chrX | 104658517 | 104663944 | L1MdTf_I | + | 5427 |
| chrX | 104663944 | 104664937 | L1MdTf_I | + | 993 |
| chrX | 104780233 | 104785484 | L1MdTf_I | + | 5251 |
| chrX | 104785484 | 104786496 | L1MdTf_I | + | 1012 |
| chrX | 104786496 | 104789174 | L1MdTf_I | + | 2678 |
| chrX | 104789174 | 104790186 | L1MdTf_I | + | 1012 |
| chrX | 104790186 | 104792864 | L1MdTf_II | + | 2678 |
| chrX | 104792864 | 104793876 | L1MdTf_II | + | 1012 |
| chrX | 106374024 | 106379249 | L1MdTf_II | + | 5225 |
| chrX | 106379249 | 106380242 | L1MdTf_II | + | 993 |
| chrX | 106645597 | 106651290 | L1MdTf_II | + | 5693 |

| | | | | | |
|---|---|---|---|---|---|
| chrX | 106651290 | 106652283 | L1MdTf_II | + | 993 |
| chrX | 108479468 | 108479669 | L1MdTf_II | + | 201 |
| chrX | 108479670 | 108480665 | L1MdTf_II | + | 995 |
| chrX | 108747624 | 108752863 | L1MdTf_III | + | 5239 |
| chrX | 108752863 | 108753856 | L1MdTf_III | + | 993 |
| chrX | 109460424 | 109466466 | L1MdTf_I | + | 6042 |
| chrX | 109466467 | 109467460 | L1MdTf_I | + | 993 |
| chrX | 109489071 | 109494319 | L1MdTf_III | + | 5248 |
| chrX | 109494319 | 109495303 | L1MdTf_III | + | 984 |
| chrX | 109988877 | 109994509 | L1MdTf_II | + | 5632 |
| chrX | 109994509 | 109995502 | L1MdTf_II | + | 993 |
| chrX | 110159094 | 110159563 | L1MdTf_II | + | 469 |
| chrX | 110159563 | 110160556 | L1MdTf_II | + | 993 |
| chrX | 110388635 | 110392183 | L1MdTf_I | + | 3548 |
| chrX | 110392183 | 110393201 | L1MdTf_I | + | 1018 |
| chrX | 111389625 | 111395281 | L1MdTf_III | + | 5656 |
| chrX | 111395281 | 111396274 | L1MdTf_III | + | 993 |
| chrX | 111749816 | 111755207 | L1MdTf_II | + | 5391 |
| chrX | 111755207 | 111756203 | L1MdTf_II | + | 996 |
| chrX | 112000899 | 112001691 | L1MdTf_II | + | 792 |
| chrX | 112001691 | 112002675 | L1MdTf_II | + | 984 |
| chrX | 112261563 | 112266970 | L1MdTf_I | + | 5407 |
| chrX | 112266970 | 112267963 | L1MdTf_I | + | 993 |
| chrX | 112765778 | 112766099 | L1MdTf_III | - | 321 |
| chrX | 112766099 | 112766588 | L1MdTf_III | + | 489 |
| chrX | 113049902 | 113055345 | L1MdTf_II | + | 5443 |
| chrX | 113055345 | 113056362 | L1MdTf_II | + | 1017 |
| chrX | 113782387 | 113786839 | L1MdTf_III | + | 4452 |
| chrX | 113786839 | 113787830 | L1MdTf_III | + | 991 |
| chrX | 114093202 | 114095007 | L1MdTf_I | + | 1805 |
| chrX | 114095007 | 114096000 | L1MdTf_I | + | 993 |
| chrX | 114324218 | 114329723 | L1MdTf_III | + | 5505 |
| chrX | 114329723 | 114330716 | L1MdTf_III | + | 993 |
| chrX | 114404798 | 114407051 | L1MdTf_II | + | 2253 |
| chrX | 114407051 | 114408043 | L1MdTf_II | + | 992 |
| chrX | 114570219 | 114575627 | L1MdTf_III | + | 5408 |
| chrX | 114575627 | 114576619 | L1MdTf_III | + | 992 |
| chrX | 116831767 | 116832427 | L1MdTf_II | - | 660 |
| chrX | 116832427 | 116832981 | L1MdTf_II | + | 554 |
| chrX | 117109707 | 117115134 | L1MdTf_II | + | 5427 |
| chrX | 117115134 | 117116127 | L1MdTf_II | + | 993 |
| chrX | 117851722 | 117857060 | L1MdTf_III | + | 5338 |
| chrX | 117857060 | 117858052 | L1MdTf_III | + | 992 |
| chrX | 118161149 | 118166578 | L1MdTf_III | + | 5429 |
| chrX | 118166578 | 118167571 | L1MdTf_III | + | 993 |

| | | | | | |
|---|---|---|---|---|---|
| chrX | 118467810 | 118468504 | L1MdTf_I | + | 694 |
| chrX | 118468504 | 118469516 | L1MdTf_I | + | 1012 |
| chrX | 118478263 | 118483665 | L1MdTf_III | + | 5402 |
| chrX | 118483665 | 118484661 | L1MdTf_III | + | 996 |
| chrX | 119295202 | 119300694 | L1MdTf_II | + | 5492 |
| chrX | 119300694 | 119301687 | L1MdTf_II | + | 993 |
| chrX | 120494906 | 120500000 | L1MdTf_II | + | 5094 |
| chrX | 120500000 | 120500552 | L1MdTf_I | + | 552 |
| chrX | 121089061 | 121094450 | L1MdTf_III | + | 5389 |
| chrX | 121094450 | 121095443 | L1MdTf_III | + | 993 |
| chrX | 127262366 | 127263691 | L1MdTf_II | + | 1325 |
| chrX | 127263691 | 127264684 | L1MdTf_II | + | 993 |
| chrX | 127496770 | 127500000 | L1MdTf_I | - | 3230 |
| chrX | 127500000 | 127503173 | L1MdTf_II | - | 3173 |
| chrX | 127695532 | 127700793 | L1MdTf_III | + | 5261 |
| chrX | 127700793 | 127701784 | L1MdTf_III | + | 991 |
| chrX | 128999037 | 129000000 | L1MdTf_I | - | 963 |
| chrX | 129000000 | 129000736 | L1MdTf_I | - | 736 |
| chrX | 129286798 | 129287451 | L1MdTf_I | - | 653 |
| chrX | 129287451 | 129288312 | L1MdTf_I | + | 861 |
| chrX | 130398334 | 130401234 | L1MdTf_III | + | 2900 |
| chrX | 130401234 | 130402227 | L1MdTf_III | + | 993 |
| chrX | 130894341 | 130899571 | L1MdTf_II | + | 5230 |
| chrX | 130899571 | 130900550 | L1MdTf_II | + | 979 |
| chrX | 130958616 | 130959060 | L1MdTf_I | - | 444 |
| chrX | 130959060 | 130959436 | L1MdTf_II | + | 376 |
| chrX | 131175178 | 131175296 | L1MdTf_II | + | 118 |
| chrX | 131175296 | 131176289 | L1MdTf_II | + | 993 |
| chrX | 131435771 | 131436661 | L1MdTf_II | + | 890 |
| chrX | 131436661 | 131437654 | L1MdTf_II | + | 993 |
| chrX | 132699070 | 132700221 | L1MdTf_III | + | 1151 |
| chrX | 132700221 | 132701234 | L1MdTf_III | + | 1013 |
| chrX | 134294821 | 134300503 | L1MdTf_III | + | 5682 |
| chrX | 134300503 | 134301496 | L1MdTf_III | + | 993 |
| chrX | 134355420 | 134361102 | L1MdTf_III | + | 5682 |
| chrX | 134361102 | 134362095 | L1MdTf_III | + | 993 |
| chrX | 135465231 | 135466540 | L1MdTf_I | + | 1309 |
| chrX | 135466540 | 135467533 | L1MdTf_I | + | 993 |
| chrX | 136199063 | 136200423 | L1MdTf_II | - | 1360 |
| chrX | 136200423 | 136201441 | L1MdTf_II | - | 1018 |
| chrX | 136331998 | 136333335 | L1MdTf_II | + | 1337 |
| chrX | 136333335 | 136334328 | L1MdTf_II | + | 993 |
| chrX | 138283703 | 138289102 | L1MdTf_I | + | 5399 |
| chrX | 138289102 | 138290095 | L1MdTf_I | + | 993 |
| chrX | 141645861 | 141651084 | L1MdTf_I | + | 5223 |

| | | | | | |
|---|---|---|---|---|---|
| chrX | 141651084 | 141652080 | L1MdTf_I | + | 996 |
| chrX | 142012358 | 142018317 | L1MdTf_III | + | 5959 |
| chrX | 142018317 | 142019312 | L1MdTf_III | + | 995 |
| chrX | 142037408 | 142037645 | L1MdTf_III | + | 237 |
| chrX | 142037645 | 142038642 | L1MdTf_III | + | 997 |
| chrX | 143372198 | 143377408 | L1MdTf_III | + | 5210 |
| chrX | 143377408 | 143378401 | L1MdTf_III | + | 993 |
| chrX | 143831998 | 143833875 | L1MdTf_III | + | 1877 |
| chrX | 143833875 | 143834868 | L1MdTf_III | + | 993 |
| chrX | 144097716 | 144102936 | L1MdTf_III | + | 5220 |
| chrX | 144102936 | 144103929 | L1MdTf_III | + | 993 |
| chrX | 145020217 | 145025418 | L1MdTf_III | + | 5201 |
| chrX | 145025418 | 145026402 | L1MdTf_III | + | 984 |
| chrX | 145136457 | 145141462 | L1MdTf_I | + | 5005 |
| chrX | 145141462 | 145142483 | L1MdTf_I | + | 1021 |
| chrX | 145541063 | 145543861 | L1MdTf_I | + | 2798 |
| chrX | 145543861 | 145544854 | L1MdTf_I | + | 993 |
| chrX | 145612925 | 145613482 | L1MdTf_I | + | 557 |
| chrX | 145613482 | 145614475 | L1MdTf_I | + | 993 |
| chrX | 145678361 | 145679447 | L1MdTf_III | + | 1086 |
| chrX | 145679448 | 145680436 | L1MdTf_III | + | 988 |
| chrX | 145683482 | 145686742 | L1MdTf_III | + | 3260 |
| chrX | 145686743 | 145687731 | L1MdTf_III | + | 988 |
| chrX | 145893110 | 145898317 | L1MdTf_II | + | 5207 |
| chrX | 145898317 | 145899310 | L1MdTf_II | + | 993 |
| chrX | 149105029 | 149108937 | L1MdTf_II | + | 3908 |
| chrX | 149108937 | 149109928 | L1MdTf_II | + | 991 |
| chrX | 149312982 | 149318289 | L1MdTf_I | + | 5307 |
| chrX | 149318289 | 149319282 | L1MdTf_I | + | 993 |
| chrX | 149724345 | 149724600 | L1MdTf_III | + | 255 |
| chrX | 149724600 | 149725616 | L1MdTf_III | + | 1016 |
| chrX | 150021298 | 150026879 | L1MdTf_I | + | 5581 |
| chrX | 150026879 | 150027872 | L1MdTf_I | + | 993 |
| chrX | 150525806 | 150532000 | L1MdTf_III | + | 6194 |
| chrX | 150532000 | 150532995 | L1MdTf_III | + | 995 |
| chrX | 151306607 | 151312468 | L1MdTf_II | + | 5861 |
| chrX | 151312468 | 151313461 | L1MdTf_II | + | 993 |
| chrX | 151660213 | 151660852 | L1MdTf_III | + | 639 |
| chrX | 151660852 | 151661845 | L1MdTf_III | + | 993 |
| chrX | 153792865 | 153795230 | L1MdTf_II | + | 2365 |
| chrX | 153795230 | 153796223 | L1MdTf_II | + | 993 |
| chrX | 154196249 | 154196843 | L1MdTf_II | - | 594 |
| chrX | 154196843 | 154197240 | L1MdTf_II | + | 397 |
| chrX | 154242222 | 154248069 | L1MdTf_I | + | 5847 |
| chrX | 154248069 | 154249092 | L1MdTf_I | + | 1023 |

| | | | | | |
|---|---|---|---|---|---|
| chrX | 154847002 | 154853115 | L1MdTf_I | + | 6113 |
| chrX | 154853115 | 154854108 | L1MdTf_I | + | 993 |
| chrX | 155159878 | 155161251 | L1MdTf_II | + | 1373 |
| chrX | 155161251 | 155162244 | L1MdTf_II | + | 993 |
| chrX | 155287120 | 155289005 | L1MdTf_III | + | 1885 |
| chrX | 155289005 | 155289997 | L1MdTf_III | + | 992 |
| chrX | 155714736 | 155720404 | L1MdTf_I | + | 5668 |
| chrX | 155720404 | 155721393 | L1MdTf_I | + | 989 |
| chrX | 156065971 | 156071660 | L1MdTf_III | + | 5689 |
| chrX | 156071660 | 156072653 | L1MdTf_III | + | 993 |
| chrX | 156895383 | 156901068 | L1MdTf_III | + | 5685 |
| chrX | 156901069 | 156902061 | L1MdTf_III | + | 992 |
| chrX | 157387331 | 157389274 | L1MdTf_II | + | 1943 |
| chrX | 157389274 | 157390267 | L1MdTf_II | + | 993 |
| chrX | 157652837 | 157658476 | L1MdTf_I | + | 5639 |
| chrX | 157658476 | 157659499 | L1MdTf_I | + | 1023 |
| chrX | 157684317 | 157689541 | L1MdTf_I | + | 5224 |
| chrX | 157689541 | 157690549 | L1MdTf_I | + | 1008 |
| chrX | 157788814 | 157789738 | L1MdTf_II | - | 924 |
| chrX | 157789739 | 157790073 | L1MdTf_II | + | 334 |
| chrX | 160661855 | 160662635 | L1MdTf_III | - | 780 |
| chrX | 160662635 | 160662729 | L1MdTf_III | + | 94 |
| chrX | 160731312 | 160731717 | L1MdTf_II | - | 405 |
| chrX | 160731717 | 160732084 | L1MdTf_II | + | 367 |
| chrX | 163397958 | 163404235 | L1MdTf_II | + | 6277 |
| chrX | 163404235 | 163405228 | L1MdTf_II | + | 993 |
| chrX | 163736579 | 163737243 | L1MdTf_II | - | 664 |
| chrX | 163737243 | 163737714 | L1MdTf_II | + | 471 |
| chrX | 163871740 | 163877193 | L1MdTf_III | + | 5453 |
| chrX | 163877193 | 163878186 | L1MdTf_III | + | 993 |
| chrX | 164249239 | 164255080 | L1MdTf_I | + | 5841 |
| chrX | 164255080 | 164256073 | L1MdTf_I | + | 993 |
| chrX | 164417327 | 164422816 | L1MdTf_III | + | 5489 |
| chrX | 164422816 | 164423809 | L1MdTf_III | + | 993 |
| chrX | 164484193 | 164489656 | L1MdTf_III | + | 5463 |
| chrX | 164489656 | 164490649 | L1MdTf_III | + | 993 |
| chrX | 165996329 | 166000000 | L1MdTf_II | + | 3671 |
| chrX | 166000000 | 166002210 | L1MdTf_I | + | 2210 |
| chrX | 166029090 | 166034260 | L1MdTf_III | + | 5170 |
| chrX | 166034260 | 166035253 | L1MdTf_III | + | 993 |
| chrX | 166532976 | 166533575 | L1MdTf_I | + | 599 |
| chrX | 166533575 | 166534568 | L1MdTf_I | + | 993 |
| chrX | 167942735 | 167948423 | L1MdTf_I | + | 5688 |
| chrX | 167948423 | 167949415 | L1MdTf_I | + | 992 |
| chrX | 168194402 | 168200276 | L1MdTf_I | + | 5874 |

| | | | | | |
|---|---|---|---|---|---|
| chrX | 168200276 | 168201263 | L1MdTf_I | + | 987 |
| chrX | 168523687 | 168528925 | L1MdTf_I | + | 5238 |
| chrX | 168528925 | 168529918 | L1MdTf_I | + | 993 |
| chrY | 123061 | 128316 | L1MdTf_III | + | 5255 |
| chrY | 128316 | 129308 | L1MdTf_III | + | 992 |
| chrY | 248225 | 253480 | L1MdTf_III | + | 5255 |
| chrY | 253480 | 254472 | L1MdTf_III | + | 992 |
| chrY | 998534 | 1000000 | L1MdTf_III | + | 1466 |
| chrY | 1000000 | 1004338 | L1MdTf_III | + | 4338 |
| chrY | 3914605 | 3920360 | L1MdTf_II | + | 5755 |
| chrY | 3920360 | 3921357 | L1MdTf_II | + | 997 |
| chrY | 5156967 | 5162241 | L1MdTf_III | + | 5274 |
| chrY | 5162241 | 5163256 | L1MdTf_III | + | 1015 |
| chrY | 8088926 | 8089412 | L1MdTf_I | - | 486 |
| chrY | 8089412 | 8090370 | L1MdTf_I | + | 958 |
| chrY | 9981167 | 9981376 | L1MdTf_III | - | 209 |
| chrY | 9981376 | 9981594 | L1MdTf_III | + | 218 |
| chrY | 14385406 | 14390839 | L1MdTf_III | + | 5433 |
| chrY | 14390839 | 14391853 | L1MdTf_III | + | 1014 |
| chrY | 14989518 | 14993622 | L1MdTf_III | + | 4104 |
| chrY | 14993622 | 14994613 | L1MdTf_III | + | 991 |
| chrY | 19097704 | 19103137 | L1MdTf_III | + | 5433 |
| chrY | 19103137 | 19104150 | L1MdTf_III | + | 1013 |
| chrY | 19702133 | 19706237 | L1MdTf_III | + | 4104 |
| chrY | 19706237 | 19707228 | L1MdTf_III | + | 991 |
| chrY | 29384795 | 29390638 | L1MdTf_III | + | 5843 |
| chrY | 29390638 | 29391658 | L1MdTf_III | + | 1020 |
| chrY | 34255935 | 34262025 | L1MdTf_III | + | 6090 |
| chrY | 34262025 | 34263018 | L1MdTf_III | + | 993 |
| chrY | 43442664 | 43444002 | L1MdTf_III | + | 1338 |
| chrY | 43444002 | 43444993 | L1MdTf_III | + | 991 |
| chrY | 45495506 | 45500000 | L1MdTf_III | - | 4494 |
| chrY | 45500001 | 45501961 | L1MdTf_III | - | 1960 |
| chrY | 47177716 | 47183123 | L1MdTf_III | + | 5407 |
| chrY | 47183123 | 47184116 | L1MdTf_III | + | 993 |
| chrY | 48111245 | 48116707 | L1MdTf_III | + | 5462 |
| chrY | 48116707 | 48117700 | L1MdTf_III | + | 993 |
| chrY | 60905550 | 60911012 | L1MdTf_III | + | 5462 |
| chrY | 60911012 | 60912005 | L1MdTf_III | + | 993 |
| chrY | 67431221 | 67436674 | L1MdTf_III | + | 5453 |
| chrY | 67436674 | 67437667 | L1MdTf_III | + | 993 |
| chrY | 78126998 | 78132451 | L1MdTf_III | + | 5453 |
| chrY | 78132451 | 78133444 | L1MdTf_III | + | 993 |
| chrY | 80047939 | 80053392 | L1MdTf_III | + | 5453 |
| chrY | 80053392 | 80054411 | L1MdTf_III | + | 1019 |

| | | | | | |
|---|---|---|---|---|---|
| chrY | 81285297 | 81290912 | L1MdTf_III | + | 5615 |
| chrY | 81290912 | 81291904 | L1MdTf_III | + | 992 |
| chrY | 89979813 | 89983793 | L1MdTf_III | + | 3980 |
| chrY | 89983793 | 89984785 | L1MdTf_III | + | 992 |
| chrY | 90207271 | 90212733 | L1MdTf_III | + | 5462 |
| chrY | 90212733 | 90213726 | L1MdTf_III | + | 993 |
| chrY | 90515767 | 90521312 | L1MdTf_III | + | 5545 |
| chrY | 90521312 | 90522325 | L1MdTf_III | + | 1013 |
| chr10 | 4861269 | 4867538 | L1MdTf_II | + | 6269 |
| chr10 | 4867538 | 4868531 | L1MdTf_II | + | 993 |
| chr10 | 5039785 | 5045646 | L1MdTf_II | + | 5861 |
| chr10 | 5045646 | 5046639 | L1MdTf_II | + | 993 |
| chr10 | 7278890 | 7280525 | L1MdTf_III | + | 1635 |
| chr10 | 7280525 | 7281547 | L1MdTf_III | + | 1022 |
| chr10 | 9902480 | 9908128 | L1MdTf_I | + | 5648 |
| chr10 | 9908128 | 9909121 | L1MdTf_I | + | 993 |
| chr10 | 10542862 | 10548391 | L1MdTf_III | + | 5529 |
| chr10 | 10548391 | 10549384 | L1MdTf_III | + | 993 |
| chr10 | 10831535 | 10836936 | L1MdTf_I | + | 5401 |
| chr10 | 10836936 | 10837929 | L1MdTf_I | + | 993 |
| chr10 | 10851116 | 10856559 | L1MdTf_I | + | 5443 |
| chr10 | 10856559 | 10857552 | L1MdTf_I | + | 993 |
| chr10 | 11455815 | 11461017 | L1MdTf_III | + | 5202 |
| chr10 | 11461017 | 11462010 | L1MdTf_III | + | 993 |
| chr10 | 11731984 | 11737215 | L1MdTf_I | + | 5231 |
| chr10 | 11737215 | 11738208 | L1MdTf_I | + | 993 |
| chr10 | 12261628 | 12267479 | L1MdTf_II | + | 5851 |
| chr10 | 12267479 | 12268472 | L1MdTf_II | + | 993 |
| chr10 | 12268484 | 12270777 | L1MdTf_I | + | 2293 |
| chr10 | 12270777 | 12271770 | L1MdTf_I | + | 993 |
| chr10 | 13768781 | 13770562 | L1MdTf_II | + | 1781 |
| chr10 | 13770563 | 13775985 | L1MdTf_II | + | 5422 |
| chr10 | 14741388 | 14741426 | L1MdTf_II | - | 38 |
| chr10 | 14741426 | 14742250 | L1MdTf_II | + | 824 |
| chr10 | 15007141 | 15007212 | L1MdTf_II | + | 71 |
| chr10 | 15007212 | 15008205 | L1MdTf_II | + | 993 |
| chr10 | 15073450 | 15078832 | L1MdTf_III | + | 5382 |
| chr10 | 15078832 | 15079825 | L1MdTf_III | + | 993 |
| chr10 | 15222061 | 15227437 | L1MdTf_III | + | 5376 |
| chr10 | 15227437 | 15228430 | L1MdTf_III | + | 993 |
| chr10 | 15510562 | 15516565 | L1MdTf_II | + | 6003 |
| chr10 | 15516565 | 15517558 | L1MdTf_II | + | 993 |
| chr10 | 15649269 | 15649414 | L1MdTf_I | - | 145 |
| chr10 | 15649414 | 15650119 | L1MdTf_II | + | 705 |
| chr10 | 15774985 | 15775523 | L1MdTf_III | - | 538 |

| | | | | | |
|---|---|---|---|---|---|
| chr10 | 15775523 | 15775717 | L1MdTf_III | - | 194 |
| chr10 | 16066405 | 16066944 | L1MdTf_II | + | 539 |
| chr10 | 16066944 | 16067937 | L1MdTf_II | + | 993 |
| chr10 | 16535980 | 16536785 | L1MdTf_I | - | 805 |
| chr10 | 16536785 | 16537059 | L1MdTf_I | + | 274 |
| chr10 | 16567858 | 16573383 | L1MdTf_II | + | 5525 |
| chr10 | 16573383 | 16574378 | L1MdTf_II | + | 995 |
| chr10 | 16696085 | 16701716 | L1MdTf_II | + | 5631 |
| chr10 | 16701716 | 16702709 | L1MdTf_II | + | 993 |
| chr10 | 17019292 | 17024531 | L1MdTf_II | + | 5239 |
| chr10 | 17024531 | 17025530 | L1MdTf_II | + | 999 |
| chr10 | 19145927 | 19148247 | L1MdTf_III | + | 2320 |
| chr10 | 19148247 | 19149240 | L1MdTf_III | + | 993 |
| chr10 | 22735217 | 22740459 | L1MdTf_II | + | 5242 |
| chr10 | 22740460 | 22741453 | L1MdTf_II | + | 993 |
| chr10 | 23376912 | 23382371 | L1MdTf_III | + | 5459 |
| chr10 | 23382371 | 23383364 | L1MdTf_III | + | 993 |
| chr10 | 23507557 | 23512974 | L1MdTf_I | + | 5417 |
| chr10 | 23512974 | 23513967 | L1MdTf_I | + | 993 |
| chr10 | 27338019 | 27338253 | L1MdTf_II | - | 234 |
| chr10 | 27338253 | 27340159 | L1MdTf_II | + | 1906 |
| chr10 | 28402808 | 28408207 | L1MdTf_II | + | 5399 |
| chr10 | 28408207 | 28409217 | L1MdTf_II | + | 1010 |
| chr10 | 28583235 | 28586745 | L1MdTf_II | + | 3510 |
| chr10 | 28586745 | 28587738 | L1MdTf_II | + | 993 |
| chr10 | 28784916 | 28790035 | L1MdTf_III | + | 5119 |
| chr10 | 28790036 | 28791027 | L1MdTf_III | + | 991 |
| chr10 | 29495631 | 29500000 | L1MdTf_II | + | 4369 |
| chr10 | 29500000 | 29501092 | L1MdTf_I | + | 1092 |
| chr10 | 29636958 | 29639123 | L1MdTf_II | + | 2165 |
| chr10 | 29639123 | 29640116 | L1MdTf_II | + | 993 |
| chr10 | 30085474 | 30087480 | L1MdTf_III | + | 2006 |
| chr10 | 30087480 | 30088472 | L1MdTf_III | + | 992 |
| chr10 | 30092010 | 30098098 | L1MdTf_I | + | 6088 |
| chr10 | 30098098 | 30099091 | L1MdTf_I | + | 993 |
| chr10 | 30344318 | 30345300 | L1MdTf_II | + | 982 |
| chr10 | 30345300 | 30346297 | L1MdTf_II | + | 997 |
| chr10 | 31657686 | 31658345 | L1MdTf_III | - | 659 |
| chr10 | 31658345 | 31659231 | L1MdTf_III | + | 886 |
| chr10 | 31873193 | 31875562 | L1MdTf_II | - | 2369 |
| chr10 | 31875562 | 31876514 | L1MdTf_II | + | 952 |
| chr10 | 32375138 | 32380724 | L1MdTf_I | + | 5586 |
| chr10 | 32380724 | 32381717 | L1MdTf_I | + | 993 |
| chr10 | 32403717 | 32408141 | L1MdTf_III | + | 4424 |
| chr10 | 32408141 | 32409134 | L1MdTf_III | + | 993 |

| | | | | | |
|---|---|---|---|---|---|
| chr10 | 33697512 | 33702947 | L1MdTf_I | + | 5435 |
| chr10 | 33702947 | 33703959 | L1MdTf_I | + | 1012 |
| chr10 | 33743216 | 33744069 | L1MdTf_III | - | 853 |
| chr10 | 33744069 | 33744185 | L1MdTf_III | + | 116 |
| chr10 | 35510458 | 35515746 | L1MdTf_III | + | 5288 |
| chr10 | 35515746 | 35516740 | L1MdTf_III | + | 994 |
| chr10 | 36152036 | 36157489 | L1MdTf_III | + | 5453 |
| chr10 | 36157489 | 36158482 | L1MdTf_III | + | 993 |
| chr10 | 38145579 | 38149050 | L1MdTf_III | - | 3471 |
| chr10 | 38149050 | 38149264 | L1MdTf_II | + | 214 |
| chr10 | 38808631 | 38814711 | L1MdTf_I | + | 6080 |
| chr10 | 38814711 | 38815704 | L1MdTf_I | + | 993 |
| chr10 | 41184822 | 41190682 | L1MdTf_II | + | 5860 |
| chr10 | 41190682 | 41191675 | L1MdTf_II | + | 993 |
| chr10 | 45393040 | 45398458 | L1MdTf_II | + | 5418 |
| chr10 | 45398458 | 45399451 | L1MdTf_II | + | 993 |
| chr10 | 45715053 | 45720506 | L1MdTf_II | + | 5453 |
| chr10 | 45720506 | 45721502 | L1MdTf_II | + | 996 |
| chr10 | 46240553 | 46246056 | L1MdTf_III | + | 5503 |
| chr10 | 46246056 | 46247040 | L1MdTf_III | + | 984 |
| chr10 | 46445677 | 46446264 | L1MdTf_III | + | 587 |
| chr10 | 46446264 | 46447257 | L1MdTf_III | + | 993 |
| chr10 | 46988615 | 46988683 | L1MdTf_II | + | 68 |
| chr10 | 46988683 | 46989676 | L1MdTf_II | + | 993 |
| chr10 | 47206977 | 47212306 | L1MdTf_III | + | 5329 |
| chr10 | 47212306 | 47213299 | L1MdTf_III | + | 993 |
| chr10 | 47933252 | 47933872 | L1MdTf_II | + | 620 |
| chr10 | 47933872 | 47934865 | L1MdTf_II | + | 993 |
| chr10 | 48107794 | 48113244 | L1MdTf_II | + | 5450 |
| chr10 | 48113244 | 48114228 | L1MdTf_II | + | 984 |
| chr10 | 48959067 | 48960613 | L1MdTf_III | + | 1546 |
| chr10 | 48960613 | 48961606 | L1MdTf_III | + | 993 |
| chr10 | 49101133 | 49106375 | L1MdTf_III | + | 5242 |
| chr10 | 49106376 | 49107369 | L1MdTf_III | + | 993 |
| chr10 | 49379901 | 49381262 | L1MdTf_III | - | 1361 |
| chr10 | 49381262 | 49381643 | L1MdTf_III | + | 381 |
| chr10 | 49699592 | 49705891 | L1MdTf_I | + | 6299 |
| chr10 | 49705891 | 49706884 | L1MdTf_I | + | 993 |
| chr10 | 49711997 | 49717249 | L1MdTf_III | + | 5252 |
| chr10 | 49717249 | 49718271 | L1MdTf_III | + | 1022 |
| chr10 | 49985212 | 49990635 | L1MdTf_II | + | 5423 |
| chr10 | 49990635 | 49991631 | L1MdTf_II | + | 996 |
| chr10 | 50413540 | 50419991 | L1MdTf_III | + | 6451 |
| chr10 | 50419991 | 50420983 | L1MdTf_III | + | 992 |
| chr10 | 52949910 | 52950854 | L1MdTf_I | + | 944 |

| | | | | | |
|---|---|---|---|---|---|
| chr10 | 52950854 | 52951847 | L1MdTf_I | + | 993 |
| chr10 | 53542413 | 53547462 | L1MdTf_I | + | 5049 |
| chr10 | 53547462 | 53548455 | L1MdTf_I | + | 993 |
| chr10 | 54354740 | 54354881 | L1MdTf_II | - | 141 |
| chr10 | 54354881 | 54355035 | L1MdTf_II | + | 154 |
| chr10 | 55450932 | 55456198 | L1MdTf_II | + | 5266 |
| chr10 | 55456198 | 55457191 | L1MdTf_II | + | 993 |
| chr10 | 55721332 | 55723045 | L1MdTf_III | + | 1713 |
| chr10 | 55723045 | 55724038 | L1MdTf_III | + | 993 |
| chr10 | 55770630 | 55775851 | L1MdTf_III | + | 5221 |
| chr10 | 55775851 | 55776844 | L1MdTf_III | + | 993 |
| chr10 | 55907182 | 55912622 | L1MdTf_II | + | 5440 |
| chr10 | 55912622 | 55913606 | L1MdTf_II | + | 984 |
| chr10 | 56146023 | 56146677 | L1MdTf_I | + | 654 |
| chr10 | 56146677 | 56147670 | L1MdTf_II | + | 993 |
| chr10 | 56163350 | 56169013 | L1MdTf_II | + | 5663 |
| chr10 | 56169013 | 56170036 | L1MdTf_II | + | 1023 |
| chr10 | 56367161 | 56367449 | L1MdTf_II | - | 288 |
| chr10 | 56367449 | 56367598 | L1MdTf_II | + | 149 |
| chr10 | 56974425 | 56980778 | L1MdTf_III | + | 6353 |
| chr10 | 56980778 | 56981772 | L1MdTf_III | + | 994 |
| chr10 | 57326660 | 57331788 | L1MdTf_III | + | 5128 |
| chr10 | 57331788 | 57332781 | L1MdTf_III | + | 993 |
| chr10 | 57642734 | 57648375 | L1MdTf_I | + | 5641 |
| chr10 | 57648375 | 57649368 | L1MdTf_I | + | 993 |
| chr10 | 63348436 | 63353856 | L1MdTf_I | + | 5420 |
| chr10 | 63353856 | 63354849 | L1MdTf_I | + | 993 |
| chr10 | 63559290 | 63565209 | L1MdTf_I | + | 5919 |
| chr10 | 63565209 | 63566224 | L1MdTf_I | + | 1015 |
| chr10 | 63577428 | 63582742 | L1MdTf_III | + | 5314 |
| chr10 | 63582742 | 63583762 | L1MdTf_III | + | 1020 |
| chr10 | 64656245 | 64657633 | L1MdTf_I | + | 1388 |
| chr10 | 64657633 | 64658626 | L1MdTf_I | + | 993 |
| chr10 | 65908438 | 65914059 | L1MdTf_I | + | 5621 |
| chr10 | 65914059 | 65915080 | L1MdTf_I | + | 1021 |
| chr10 | 66417990 | 66423815 | L1MdTf_I | + | 5825 |
| chr10 | 66423815 | 66424808 | L1MdTf_I | + | 993 |
| chr10 | 69032454 | 69037884 | L1MdTf_I | + | 5430 |
| chr10 | 69037884 | 69038877 | L1MdTf_I | + | 993 |
| chr10 | 69329490 | 69329744 | L1MdTf_III | - | 254 |
| chr10 | 69329745 | 69330390 | L1MdTf_III | + | 645 |
| chr10 | 70718040 | 70721923 | L1MdTf_I | + | 3883 |
| chr10 | 70721924 | 70722916 | L1MdTf_I | + | 992 |
| chr10 | 73273352 | 73279192 | L1MdTf_I | + | 5840 |
| chr10 | 73279192 | 73280185 | L1MdTf_I | + | 993 |

| | | | | | |
|---|---|---|---|---|---|
| chr10 | 73801276 | 73801339 | L1MdTf_I | + | 63 |
| chr10 | 73801339 | 73802332 | L1MdTf_I | + | 993 |
| chr10 | 73972059 | 73977314 | L1MdTf_III | + | 5255 |
| chr10 | 73977314 | 73978307 | L1MdTf_III | + | 993 |
| chr10 | 74178861 | 74184076 | L1MdTf_II | + | 5215 |
| chr10 | 74184076 | 74185069 | L1MdTf_II | + | 993 |
| chr10 | 74693248 | 74695380 | L1MdTf_III | + | 2132 |
| chr10 | 74695380 | 74696373 | L1MdTf_III | + | 993 |
| chr10 | 78569040 | 78570627 | L1MdTf_III | + | 1587 |
| chr10 | 78570627 | 78571620 | L1MdTf_III | + | 993 |
| chr10 | 78730472 | 78735915 | L1MdTf_III | + | 5443 |
| chr10 | 78735915 | 78736910 | L1MdTf_III | + | 995 |
| chr10 | 78776370 | 78781798 | L1MdTf_II | + | 5428 |
| chr10 | 78781798 | 78782796 | L1MdTf_II | + | 998 |
| chr10 | 86069927 | 86076017 | L1MdTf_II | + | 6090 |
| chr10 | 86076017 | 86077010 | L1MdTf_II | + | 993 |
| chr10 | 86442664 | 86444188 | L1MdTf_II | + | 1524 |
| chr10 | 86444189 | 86445184 | L1MdTf_II | + | 995 |
| chr10 | 87468245 | 87470076 | L1MdTf_II | + | 1831 |
| chr10 | 87470076 | 87471069 | L1MdTf_II | + | 993 |
| chr10 | 87637016 | 87637598 | L1MdTf_III | + | 582 |
| chr10 | 87637598 | 87638591 | L1MdTf_III | + | 993 |
| chr10 | 90084816 | 90090252 | L1MdTf_II | + | 5436 |
| chr10 | 90090252 | 90091245 | L1MdTf_II | + | 993 |
| chr10 | 90436666 | 90436893 | L1MdTf_II | - | 227 |
| chr10 | 90436893 | 90437303 | L1MdTf_II | + | 410 |
| chr10 | 97752957 | 97755264 | L1MdTf_III | + | 2307 |
| chr10 | 97755264 | 97756257 | L1MdTf_III | + | 993 |
| chr10 | 97833954 | 97839195 | L1MdTf_III | + | 5241 |
| chr10 | 97839195 | 97840188 | L1MdTf_III | + | 993 |
| chr10 | 97996655 | 98000000 | L1MdTf_III | - | 3345 |
| chr10 | 98000000 | 98002912 | L1MdTf_III | - | 2912 |
| chr10 | 99497854 | 99500000 | L1MdTf_III | - | 2146 |
| chr10 | 99500000 | 99504145 | L1MdTf_III | - | 4145 |
| chr10 | 100048616 | 100053965 | L1MdTf_III | + | 5349 |
| chr10 | 100053965 | 100054958 | L1MdTf_III | + | 993 |
| chr10 | 100482457 | 100487759 | L1MdTf_III | + | 5302 |
| chr10 | 100487759 | 100488752 | L1MdTf_III | + | 993 |
| chr10 | 101876427 | 101876761 | L1MdTf_II | + | 334 |
| chr10 | 101876761 | 101877782 | L1MdTf_II | + | 1021 |
| chr10 | 102159437 | 102164877 | L1MdTf_III | + | 5440 |
| chr10 | 102164877 | 102165868 | L1MdTf_III | + | 991 |
| chr10 | 103703370 | 103708592 | L1MdTf_II | + | 5222 |
| chr10 | 103708592 | 103709585 | L1MdTf_II | + | 993 |
| chr10 | 103829622 | 103832517 | L1MdTf_II | + | 2895 |

| | | | | | |
|---|---|---|---|---|---|
| chr10 | 103832517 | 103833510 | L1MdTf_II | + | 993 |
| chr10 | 104064966 | 104069771 | L1MdTf_I | + | 4805 |
| chr10 | 104069771 | 104070764 | L1MdTf_I | + | 993 |
| chr10 | 104324908 | 104325050 | L1MdTf_III | - | 142 |
| chr10 | 104325050 | 104325104 | L1MdTf_III | + | 54 |
| chr10 | 104600986 | 104606386 | L1MdTf_II | + | 5400 |
| chr10 | 104606386 | 104607375 | L1MdTf_II | + | 989 |
| chr10 | 105454276 | 105455819 | L1MdTf_III | + | 1543 |
| chr10 | 105455820 | 105456812 | L1MdTf_III | + | 992 |
| chr10 | 106042381 | 106048444 | L1MdTf_I | + | 6063 |
| chr10 | 106048444 | 106049437 | L1MdTf_I | + | 993 |
| chr10 | 106226498 | 106232468 | L1MdTf_II | + | 5970 |
| chr10 | 106232468 | 106237890 | L1MdTf_I | + | 5422 |
| chr10 | 110060780 | 110066181 | L1MdTf_III | + | 5401 |
| chr10 | 110066181 | 110067173 | L1MdTf_III | + | 992 |
| chr10 | 110340293 | 110345724 | L1MdTf_II | + | 5431 |
| chr10 | 110345724 | 110346720 | L1MdTf_II | + | 996 |
| chr10 | 111984407 | 111988002 | L1MdTf_III | + | 3595 |
| chr10 | 111988002 | 111988994 | L1MdTf_III | + | 992 |
| chr10 | 112260993 | 112266835 | L1MdTf_II | + | 5842 |
| chr10 | 112266835 | 112267828 | L1MdTf_II | + | 993 |
| chr10 | 112422918 | 112428555 | L1MdTf_II | + | 5637 |
| chr10 | 112428555 | 112429548 | L1MdTf_II | + | 993 |
| chr10 | 112540176 | 112540830 | L1MdTf_II | + | 654 |
| chr10 | 112540830 | 112541823 | L1MdTf_II | + | 993 |
| chr10 | 113891534 | 113896909 | L1MdTf_III | + | 5375 |
| chr10 | 113896909 | 113897903 | L1MdTf_III | + | 994 |
| chr10 | 114371375 | 114377235 | L1MdTf_I | + | 5860 |
| chr10 | 114377235 | 114378247 | L1MdTf_I | + | 1012 |
| chr10 | 114696506 | 114701860 | L1MdTf_III | + | 5354 |
| chr10 | 114701860 | 114702853 | L1MdTf_III | + | 993 |
| chr10 | 114872051 | 114877678 | L1MdTf_II | + | 5627 |
| chr10 | 114877678 | 114878673 | L1MdTf_II | + | 995 |
| chr10 | 115860209 | 115865440 | L1MdTf_I | + | 5231 |
| chr10 | 115865440 | 115866433 | L1MdTf_I | + | 993 |
| chr10 | 115997231 | 115997302 | L1MdTf_II | - | 71 |
| chr10 | 115997302 | 115997360 | L1MdTf_II | + | 58 |
| chr10 | 122070834 | 122076703 | L1MdTf_II | + | 5869 |
| chr10 | 122076703 | 122077696 | L1MdTf_II | + | 993 |
| chr10 | 122227281 | 122233139 | L1MdTf_II | + | 5858 |
| chr10 | 122233139 | 122234132 | L1MdTf_II | + | 993 |
| chr10 | 123159828 | 123160397 | L1MdTf_I | - | 569 |
| chr10 | 123160397 | 123160605 | L1MdTf_I | + | 208 |
| chr10 | 124544517 | 124544581 | L1MdTf_II | - | 64 |
| chr10 | 124544581 | 124545531 | L1MdTf_II | + | 950 |

| | | | | | |
|---|---|---|---|---|---|
| chr10 | 126536954 | 126542425 | L1MdTf_I | + | 5471 |
| chr10 | 126542425 | 126543418 | L1MdTf_I | + | 993 |
| chr10 | 129025348 | 129030927 | L1MdTf_I | + | 5579 |
| chr10 | 129030927 | 129031920 | L1MdTf_I | + | 993 |
| chr10 | 129249139 | 129254468 | L1MdTf_II | + | 5329 |
| chr10 | 129254468 | 129255463 | L1MdTf_II | + | 995 |
| chr11 | 8099359 | 8104999 | L1MdTf_I | + | 5640 |
| chr11 | 8104999 | 8106013 | L1MdTf_I | + | 1014 |
| chr11 | 11237224 | 11243500 | L1MdTf_I | + | 6276 |
| chr11 | 11243500 | 11244493 | L1MdTf_I | + | 993 |
| chr11 | 12079182 | 12084986 | L1MdTf_II | + | 5804 |
| chr11 | 12084986 | 12085979 | L1MdTf_II | + | 993 |
| chr11 | 13329106 | 13334747 | L1MdTf_II | + | 5641 |
| chr11 | 13334747 | 13335740 | L1MdTf_II | + | 993 |
| chr11 | 13565249 | 13571726 | L1MdTf_I | + | 6477 |
| chr11 | 13571726 | 13572746 | L1MdTf_I | + | 1020 |
| chr11 | 13772944 | 13773618 | L1MdTf_III | + | 674 |
| chr11 | 13773618 | 13774602 | L1MdTf_III | + | 984 |
| chr11 | 14230933 | 14231489 | L1MdTf_III | + | 556 |
| chr11 | 14231489 | 14232480 | L1MdTf_III | + | 991 |
| chr11 | 14865873 | 14871098 | L1MdTf_II | + | 5225 |
| chr11 | 14871098 | 14872091 | L1MdTf_II | + | 993 |
| chr11 | 16138557 | 16141408 | L1MdTf_III | + | 2851 |
| chr11 | 16141408 | 16142400 | L1MdTf_III | + | 992 |
| chr11 | 16335704 | 16341343 | L1MdTf_II | + | 5639 |
| chr11 | 16341343 | 16342336 | L1MdTf_II | + | 993 |
| chr11 | 17996001 | 18000000 | L1MdTf_III | - | 3999 |
| chr11 | 18000000 | 18002722 | L1MdTf_III | - | 2722 |
| chr11 | 18586168 | 18591614 | L1MdTf_III | + | 5446 |
| chr11 | 18591614 | 18592606 | L1MdTf_III | + | 992 |
| chr11 | 19032668 | 19038044 | L1MdTf_III | + | 5376 |
| chr11 | 19038044 | 19039037 | L1MdTf_III | + | 993 |
| chr11 | 20545860 | 20546677 | L1MdTf_III | - | 817 |
| chr11 | 20546677 | 20548128 | L1MdTf_III | + | 1451 |
| chr11 | 21324596 | 21324744 | L1MdTf_II | - | 148 |
| chr11 | 21324744 | 21325114 | L1MdTf_II | + | 370 |
| chr11 | 21856987 | 21862429 | L1MdTf_I | + | 5442 |
| chr11 | 21862429 | 21863422 | L1MdTf_I | + | 993 |
| chr11 | 24906871 | 24907922 | L1MdTf_III | + | 1051 |
| chr11 | 24907922 | 24908915 | L1MdTf_III | + | 993 |
| chr11 | 25920275 | 25925498 | L1MdTf_III | + | 5223 |
| chr11 | 25925498 | 25926491 | L1MdTf_III | + | 993 |
| chr11 | 26121474 | 26125229 | L1MdTf_I | + | 3755 |
| chr11 | 26125229 | 26126222 | L1MdTf_I | + | 993 |
| chr11 | 26376530 | 26382362 | L1MdTf_I | + | 5832 |

| | | | | | |
|---|---|---|---|---|---|
| chr11 | 26382362 | 26383355 | L1MdTf_I | + | 993 |
| chr11 | 26619333 | 26624859 | L1MdTf_III | + | 5526 |
| chr11 | 26624859 | 26625852 | L1MdTf_III | + | 993 |
| chr11 | 26682852 | 26688276 | L1MdTf_II | + | 5424 |
| chr11 | 26688276 | 26689261 | L1MdTf_II | + | 985 |
| chr11 | 27485073 | 27490308 | L1MdTf_III | + | 5235 |
| chr11 | 27490309 | 27491301 | L1MdTf_III | + | 992 |
| chr11 | 29388716 | 29394141 | L1MdTf_I | + | 5425 |
| chr11 | 29394141 | 29395137 | L1MdTf_I | + | 996 |
| chr11 | 30985685 | 30991384 | L1MdTf_III | + | 5699 |
| chr11 | 30991384 | 30992377 | L1MdTf_III | + | 993 |
| chr11 | 33963888 | 33969156 | L1MdTf_III | + | 5268 |
| chr11 | 33969156 | 33970162 | L1MdTf_III | + | 1006 |
| chr11 | 34740768 | 34746001 | L1MdTf_II | + | 5233 |
| chr11 | 34746001 | 34746995 | L1MdTf_II | + | 994 |
| chr11 | 36233266 | 36238676 | L1MdTf_III | + | 5410 |
| chr11 | 36238676 | 36239669 | L1MdTf_III | + | 993 |
| chr11 | 37643268 | 37649361 | L1MdTf_II | + | 6093 |
| chr11 | 37649361 | 37650354 | L1MdTf_II | + | 993 |
| chr11 | 37904107 | 37909514 | L1MdTf_II | + | 5407 |
| chr11 | 37909514 | 37910507 | L1MdTf_II | + | 993 |
| chr11 | 38870796 | 38876047 | L1MdTf_III | + | 5251 |
| chr11 | 38876048 | 38877040 | L1MdTf_III | + | 992 |
| chr11 | 39933561 | 39934215 | L1MdTf_III | - | 654 |
| chr11 | 39934215 | 39934658 | L1MdTf_III | + | 443 |
| chr11 | 40524722 | 40530162 | L1MdTf_II | + | 5440 |
| chr11 | 40530162 | 40531155 | L1MdTf_II | + | 993 |
| chr11 | 40907093 | 40912672 | L1MdTf_I | + | 5579 |
| chr11 | 40912672 | 40913663 | L1MdTf_I | + | 991 |
| chr11 | 40996104 | 41000000 | L1MdTf_II | - | 3896 |
| chr11 | 41000000 | 41002736 | L1MdTf_II | - | 2736 |
| chr11 | 41449473 | 41454693 | L1MdTf_III | + | 5220 |
| chr11 | 41454693 | 41455683 | L1MdTf_III | + | 990 |
| chr11 | 42562190 | 42562376 | L1MdTf_III | - | 186 |
| chr11 | 42562376 | 42562641 | L1MdTf_III | - | 265 |
| chr11 | 42577222 | 42579916 | L1MdTf_I | + | 2694 |
| chr11 | 42579916 | 42580909 | L1MdTf_I | + | 993 |
| chr11 | 42670038 | 42675272 | L1MdTf_I | + | 5234 |
| chr11 | 42675272 | 42676265 | L1MdTf_I | + | 993 |
| chr11 | 43194530 | 43200166 | L1MdTf_III | + | 5636 |
| chr11 | 43200166 | 43201159 | L1MdTf_III | + | 993 |
| chr11 | 44196905 | 44202355 | L1MdTf_III | + | 5450 |
| chr11 | 44202355 | 44203348 | L1MdTf_III | + | 993 |
| chr11 | 44474769 | 44474969 | L1MdTf_I | + | 200 |
| chr11 | 44474969 | 44475962 | L1MdTf_I | + | 993 |

| | | | | | |
|---|---|---|---|---|---|
| chr11 | 45258612 | 45264463 | L1MdTf_I | + | 5851 |
| chr11 | 45264463 | 45265456 | L1MdTf_I | + | 993 |
| chr11 | 45615264 | 45622109 | L1MdTf_I | + | 6845 |
| chr11 | 45622109 | 45623102 | L1MdTf_I | + | 993 |
| chr11 | 45660374 | 45666861 | L1MdTf_I | + | 6487 |
| chr11 | 45666861 | 45667854 | L1MdTf_I | + | 993 |
| chr11 | 47741223 | 47746902 | L1MdTf_I | + | 5679 |
| chr11 | 47746902 | 47747895 | L1MdTf_I | + | 993 |
| chr11 | 48342679 | 48348159 | L1MdTf_III | + | 5480 |
| chr11 | 48348159 | 48349152 | L1MdTf_III | + | 993 |
| chr11 | 56623158 | 56628614 | L1MdTf_III | + | 5456 |
| chr11 | 56628614 | 56629607 | L1MdTf_III | + | 993 |
| chr11 | 63537739 | 63543396 | L1MdTf_I | + | 5657 |
| chr11 | 63543396 | 63544389 | L1MdTf_I | + | 993 |
| chr11 | 64189972 | 64195813 | L1MdTf_I | + | 5841 |
| chr11 | 64195813 | 64196806 | L1MdTf_I | + | 993 |
| chr11 | 66351550 | 66357005 | L1MdTf_III | + | 5455 |
| chr11 | 66357006 | 66357998 | L1MdTf_III | + | 992 |
| chr11 | 73360486 | 73366140 | L1MdTf_II | + | 5654 |
| chr11 | 73366141 | 73367133 | L1MdTf_II | + | 992 |
| chr11 | 74192689 | 74203093 | L1MdTf_I | + | 10404 |
| chr11 | 74203093 | 74204086 | L1MdTf_I | + | 993 |
| chr11 | 89851650 | 89856872 | L1MdTf_III | + | 5222 |
| chr11 | 89856872 | 89857865 | L1MdTf_III | + | 993 |
| chr11 | 90911318 | 90916746 | L1MdTf_II | + | 5428 |
| chr11 | 90916746 | 90917743 | L1MdTf_II | + | 997 |
| chr11 | 91629212 | 91629844 | L1MdTf_I | - | 632 |
| chr11 | 91629844 | 91630642 | L1MdTf_I | + | 798 |
| chr11 | 92189690 | 92189849 | L1MdTf_III | + | 159 |
| chr11 | 92189849 | 92190842 | L1MdTf_III | + | 993 |
| chr11 | 99721332 | 99722549 | L1MdTf_II | + | 1217 |
| chr11 | 99722549 | 99723562 | L1MdTf_II | + | 1013 |
| chr11 | 107996687 | 108000000 | L1MdTf_III | + | 3313 |
| chr11 | 108000000 | 108003293 | L1MdTf_III | + | 3293 |
| chr11 | 112269092 | 112269665 | L1MdTf_I | + | 573 |
| chr11 | 112269665 | 112270658 | L1MdTf_I | + | 993 |
| chr11 | 120986535 | 120986612 | L1MdTf_II | + | 77 |
| chr11 | 120986613 | 120986851 | L1MdTf_II | + | 238 |
| chr12 | 6573679 | 6575436 | L1MdTf_III | + | 1757 |
| chr12 | 6575436 | 6576429 | L1MdTf_III | + | 993 |
| chr12 | 7851859 | 7857516 | L1MdTf_II | + | 5657 |
| chr12 | 7857516 | 7858513 | L1MdTf_II | + | 997 |
| chr12 | 8173892 | 8179497 | L1MdTf_II | + | 5605 |
| chr12 | 8179497 | 8180520 | L1MdTf_II | + | 1023 |
| chr12 | 9676730 | 9682197 | L1MdTf_II | + | 5467 |

| | | | | | |
|---|---|---|---|---|---|
| chr12 | 9682197 | 9683187 | L1MdTf_II | + | 990 |
| chr12 | 10747925 | 10753308 | L1MdTf_III | + | 5383 |
| chr12 | 10753308 | 10754301 | L1MdTf_III | + | 993 |
| chr12 | 10962099 | 10967289 | L1MdTf_II | + | 5190 |
| chr12 | 10967289 | 10968285 | L1MdTf_II | + | 996 |
| chr12 | 10968289 | 10970130 | L1MdTf_II | + | 1841 |
| chr12 | 10970130 | 10971126 | L1MdTf_II | + | 996 |
| chr12 | 10971130 | 10972971 | L1MdTf_II | + | 1841 |
| chr12 | 10972971 | 10973967 | L1MdTf_II | + | 996 |
| chr12 | 11038632 | 11043880 | L1MdTf_III | + | 5248 |
| chr12 | 11043880 | 11044873 | L1MdTf_III | + | 993 |
| chr12 | 11237536 | 11243141 | L1MdTf_II | + | 5605 |
| chr12 | 11243141 | 11244153 | L1MdTf_II | + | 1012 |
| chr12 | 12142515 | 12148002 | L1MdTf_III | + | 5487 |
| chr12 | 12148002 | 12148995 | L1MdTf_III | + | 993 |
| chr12 | 13270848 | 13271831 | L1MdTf_III | + | 983 |
| chr12 | 13271831 | 13272824 | L1MdTf_III | + | 993 |
| chr12 | 14030974 | 14031683 | L1MdTf_II | - | 709 |
| chr12 | 14031684 | 14031729 | L1MdTf_II | - | 45 |
| chr12 | 14499716 | 14500000 | L1MdTf_II | - | 284 |
| chr12 | 14500000 | 14505957 | L1MdTf_I | - | 5957 |
| chr12 | 14808503 | 14808992 | L1MdTf_III | - | 489 |
| chr12 | 14808992 | 14809779 | L1MdTf_III | + | 787 |
| chr12 | 15473423 | 15478886 | L1MdTf_II | + | 5463 |
| chr12 | 15478886 | 15479903 | L1MdTf_II | + | 1017 |
| chr12 | 16088892 | 16089330 | L1MdTf_III | - | 438 |
| chr12 | 16089330 | 16090062 | L1MdTf_III | + | 732 |
| chr12 | 19410702 | 19416625 | L1MdTf_II | + | 5923 |
| chr12 | 19416625 | 19417618 | L1MdTf_II | + | 993 |
| chr12 | 19676887 | 19682720 | L1MdTf_II | + | 5833 |
| chr12 | 19682720 | 19683713 | L1MdTf_II | + | 993 |
| chr12 | 20679262 | 20679648 | L1MdTf_II | - | 386 |
| chr12 | 20679648 | 20680692 | L1MdTf_I | + | 1044 |
| chr12 | 22979357 | 22979425 | L1MdTf_III | + | 68 |
| chr12 | 22979425 | 22980418 | L1MdTf_III | + | 993 |
| chr12 | 26098251 | 26104740 | L1MdTf_III | + | 6489 |
| chr12 | 26104740 | 26105733 | L1MdTf_III | + | 993 |
| chr12 | 28211279 | 28216732 | L1MdTf_III | + | 5453 |
| chr12 | 28216732 | 28217725 | L1MdTf_III | + | 993 |
| chr12 | 29101562 | 29101824 | L1MdTf_III | + | 262 |
| chr12 | 29101824 | 29102817 | L1MdTf_III | + | 993 |
| chr12 | 30237324 | 30242589 | L1MdTf_III | + | 5265 |
| chr12 | 30242589 | 30243582 | L1MdTf_III | + | 993 |
| chr12 | 31041425 | 31047083 | L1MdTf_II | + | 5658 |
| chr12 | 31047083 | 31048067 | L1MdTf_II | + | 984 |

| | | | | | |
|---|---|---|---|---|---|
| chr12 | 32420101 | 32422793 | L1MdTf_II | + | 2692 |
| chr12 | 32422793 | 32423786 | L1MdTf_II | + | 993 |
| chr12 | 33714376 | 33719802 | L1MdTf_I | + | 5426 |
| chr12 | 33719802 | 33720795 | L1MdTf_I | + | 993 |
| chr12 | 35904168 | 35910658 | L1MdTf_II | + | 6490 |
| chr12 | 35910658 | 35911651 | L1MdTf_II | + | 993 |
| chr12 | 36679085 | 36679258 | L1MdTf_III | + | 173 |
| chr12 | 36679258 | 36680250 | L1MdTf_III | + | 992 |
| chr12 | 37001699 | 37007143 | L1MdTf_III | + | 5444 |
| chr12 | 37007143 | 37008136 | L1MdTf_III | + | 993 |
| chr12 | 38097610 | 38103886 | L1MdTf_I | + | 6276 |
| chr12 | 38103886 | 38104879 | L1MdTf_I | + | 993 |
| chr12 | 38124668 | 38130094 | L1MdTf_I | + | 5426 |
| chr12 | 38130094 | 38131117 | L1MdTf_I | + | 1023 |
| chr12 | 38577380 | 38582835 | L1MdTf_III | + | 5455 |
| chr12 | 38582835 | 38583828 | L1MdTf_III | + | 993 |
| chr12 | 39039084 | 39041885 | L1MdTf_II | + | 2801 |
| chr12 | 39041885 | 39042878 | L1MdTf_II | + | 993 |
| chr12 | 41429235 | 41434501 | L1MdTf_III | + | 5266 |
| chr12 | 41434502 | 41435495 | L1MdTf_III | + | 993 |
| chr12 | 41552555 | 41557507 | L1MdTf_II | + | 4952 |
| chr12 | 41557507 | 41558500 | L1MdTf_II | + | 993 |
| chr12 | 41906709 | 41907693 | L1MdTf_II | + | 984 |
| chr12 | 41907693 | 41908712 | L1MdTf_II | + | 1019 |
| chr12 | 41915604 | 41916073 | L1MdTf_II | + | 469 |
| chr12 | 41916073 | 41917066 | L1MdTf_II | + | 993 |
| chr12 | 42378936 | 42384193 | L1MdTf_I | + | 5257 |
| chr12 | 42384193 | 42385186 | L1MdTf_I | + | 993 |
| chr12 | 42713567 | 42718993 | L1MdTf_II | + | 5426 |
| chr12 | 42718993 | 42719986 | L1MdTf_II | + | 993 |
| chr12 | 43083423 | 43089690 | L1MdTf_I | + | 6267 |
| chr12 | 43089690 | 43090683 | L1MdTf_I | + | 993 |
| chr12 | 43222055 | 43227488 | L1MdTf_I | + | 5433 |
| chr12 | 43227488 | 43228504 | L1MdTf_I | + | 1016 |
| chr12 | 43523023 | 43523292 | L1MdTf_I | + | 269 |
| chr12 | 43523292 | 43524284 | L1MdTf_II | + | 992 |
| chr12 | 43795545 | 43801081 | L1MdTf_I | + | 5536 |
| chr12 | 43801081 | 43802074 | L1MdTf_I | + | 993 |
| chr12 | 43933226 | 43938475 | L1MdTf_III | + | 5249 |
| chr12 | 43938475 | 43939481 | L1MdTf_III | + | 1006 |
| chr12 | 43975642 | 43981071 | L1MdTf_III | + | 5429 |
| chr12 | 43981071 | 43982064 | L1MdTf_III | + | 993 |
| chr12 | 44529869 | 44535992 | L1MdTf_I | + | 6123 |
| chr12 | 44535992 | 44536985 | L1MdTf_I | + | 993 |
| chr12 | 44536987 | 44537940 | L1MdTf_I | + | 953 |

| | | | | | |
|---|---|---|---|---|---|
| chr12 | 44537940 | 44538933 | L1MdTf_I | + | 993 |
| chr12 | 47326816 | 47332467 | L1MdTf_I | + | 5651 |
| chr12 | 47332467 | 47333238 | L1MdTf_I | + | 771 |
| chr12 | 48387139 | 48392739 | L1MdTf_III | + | 5600 |
| chr12 | 48392739 | 48393207 | L1MdTf_III | + | 468 |
| chr12 | 48603114 | 48605450 | L1MdTf_III | + | 2336 |
| chr12 | 48605450 | 48606443 | L1MdTf_III | + | 993 |
| chr12 | 49123093 | 49128517 | L1MdTf_III | + | 5424 |
| chr12 | 49128517 | 49129510 | L1MdTf_III | + | 993 |
| chr12 | 49178060 | 49178254 | L1MdTf_III | + | 194 |
| chr12 | 49178254 | 49179245 | L1MdTf_III | + | 991 |
| chr12 | 49652894 | 49658337 | L1MdTf_III | + | 5443 |
| chr12 | 49658337 | 49659328 | L1MdTf_III | + | 991 |
| chr12 | 49748617 | 49754279 | L1MdTf_I | + | 5662 |
| chr12 | 49754279 | 49755272 | L1MdTf_I | + | 993 |
| chr12 | 51214852 | 51220189 | L1MdTf_II | + | 5337 |
| chr12 | 51220189 | 51221182 | L1MdTf_II | + | 993 |
| chr12 | 55679400 | 55685472 | L1MdTf_I | + | 6072 |
| chr12 | 55685472 | 55686465 | L1MdTf_I | + | 993 |
| chr12 | 55826765 | 55827071 | L1MdTf_II | - | 306 |
| chr12 | 55827071 | 55827547 | L1MdTf_II | + | 476 |
| chr12 | 58816594 | 58822034 | L1MdTf_I | + | 5440 |
| chr12 | 58822034 | 58823045 | L1MdTf_I | + | 1011 |
| chr12 | 59414128 | 59415315 | L1MdTf_III | + | 1187 |
| chr12 | 59415315 | 59416308 | L1MdTf_III | + | 993 |
| chr12 | 59545291 | 59550648 | L1MdTf_III | + | 5357 |
| chr12 | 59550648 | 59551641 | L1MdTf_III | + | 993 |
| chr12 | 59608878 | 59609252 | L1MdTf_III | - | 374 |
| chr12 | 59609252 | 59609351 | L1MdTf_III | + | 99 |
| chr12 | 60450378 | 60451708 | L1MdTf_III | + | 1330 |
| chr12 | 60451709 | 60452703 | L1MdTf_III | + | 994 |
| chr12 | 61201269 | 61206685 | L1MdTf_III | + | 5416 |
| chr12 | 61206685 | 61207678 | L1MdTf_III | + | 993 |
| chr12 | 61679798 | 61685433 | L1MdTf_I | + | 5635 |
| chr12 | 61685433 | 61686426 | L1MdTf_I | + | 993 |
| chr12 | 61861466 | 61862377 | L1MdTf_I | - | 911 |
| chr12 | 61862378 | 61862476 | L1MdTf_I | + | 98 |
| chr12 | 62304653 | 62307048 | L1MdTf_I | + | 2395 |
| chr12 | 62307048 | 62308041 | L1MdTf_I | + | 993 |
| chr12 | 63006595 | 63012787 | L1MdTf_I | + | 6192 |
| chr12 | 63012787 | 63013806 | L1MdTf_I | + | 1019 |
| chr12 | 63031809 | 63037209 | L1MdTf_II | + | 5400 |
| chr12 | 63037210 | 63038202 | L1MdTf_II | + | 992 |
| chr12 | 63440584 | 63446229 | L1MdTf_II | + | 5645 |
| chr12 | 63446229 | 63447213 | L1MdTf_II | + | 984 |

| | | | | | |
|---|---|---|---|---|---|
| chr12 | 63616936 | 63622162 | L1MdTf_II | + | 5226 |
| chr12 | 63622162 | 63623155 | L1MdTf_II | + | 993 |
| chr12 | 63738339 | 63744001 | L1MdTf_I | + | 5662 |
| chr12 | 63744001 | 63744994 | L1MdTf_I | + | 993 |
| chr12 | 64772759 | 64778765 | L1MdTf_I | + | 6006 |
| chr12 | 64778765 | 64779758 | L1MdTf_I | + | 993 |
| chr12 | 65961994 | 65967843 | L1MdTf_I | + | 5849 |
| chr12 | 65967843 | 65968836 | L1MdTf_I | + | 993 |
| chr12 | 66991299 | 66992140 | L1MdTf_II | - | 841 |
| chr12 | 66992140 | 66993608 | L1MdTf_II | + | 1468 |
| chr12 | 67923221 | 67923817 | L1MdTf_III | - | 596 |
| chr12 | 67923817 | 67924286 | L1MdTf_III | + | 469 |
| chr12 | 68986447 | 68990418 | L1MdTf_III | + | 3971 |
| chr12 | 68990418 | 68991410 | L1MdTf_III | + | 992 |
| chr12 | 69005631 | 69010699 | L1MdTf_III | + | 5068 |
| chr12 | 69010699 | 69011682 | L1MdTf_III | + | 983 |
| chr12 | 69011690 | 69012850 | L1MdTf_III | + | 1160 |
| chr12 | 69012850 | 69013842 | L1MdTf_III | + | 992 |
| chr12 | 71253660 | 71259235 | L1MdTf_III | + | 5575 |
| chr12 | 71259235 | 71260228 | L1MdTf_III | + | 993 |
| chr12 | 72249759 | 72255083 | L1MdTf_I | + | 5324 |
| chr12 | 72255083 | 72256074 | L1MdTf_I | + | 991 |
| chr12 | 74624807 | 74625382 | L1MdTf_III | - | 575 |
| chr12 | 74625382 | 74626033 | L1MdTf_III | + | 651 |
| chr12 | 74667525 | 74669894 | L1MdTf_I | + | 2369 |
| chr12 | 74669894 | 74670887 | L1MdTf_I | + | 993 |
| chr12 | 74973304 | 74979215 | L1MdTf_II | + | 5911 |
| chr12 | 74979215 | 74980208 | L1MdTf_II | + | 993 |
| chr12 | 75426190 | 75432079 | L1MdTf_I | + | 5889 |
| chr12 | 75432079 | 75433072 | L1MdTf_I | + | 993 |
| chr12 | 77925691 | 77930899 | L1MdTf_III | + | 5208 |
| chr12 | 77930899 | 77931894 | L1MdTf_III | + | 995 |
| chr12 | 77931930 | 77933088 | L1MdTf_III | + | 1158 |
| chr12 | 77933089 | 77934073 | L1MdTf_III | + | 984 |
| chr12 | 78335626 | 78336024 | L1MdTf_I | - | 398 |
| chr12 | 78336024 | 78337004 | L1MdTf_I | + | 980 |
| chr12 | 87736259 | 87737332 | L1MdTf_I | + | 1073 |
| chr12 | 87737332 | 87738352 | L1MdTf_I | + | 1020 |
| chr12 | 87884039 | 87884625 | L1MdTf_II | + | 586 |
| chr12 | 87884625 | 87885618 | L1MdTf_II | + | 993 |
| chr12 | 88196306 | 88201535 | L1MdTf_II | + | 5229 |
| chr12 | 88201535 | 88202528 | L1MdTf_II | + | 993 |
| chr12 | 90404177 | 90405306 | L1MdTf_III | + | 1129 |
| chr12 | 90405306 | 90406299 | L1MdTf_III | + | 993 |
| chr12 | 90628380 | 90629516 | L1MdTf_III | + | 1136 |

| | | | | | |
|---|---|---|---|---|---|
| chr12 | 90629516 | 90630500 | L1MdTf_III | + | 984 |
| chr12 | 91441769 | 91442237 | L1MdTf_III | - | 468 |
| chr12 | 91442237 | 91442709 | L1MdTf_III | + | 472 |
| chr12 | 92724930 | 92726857 | L1MdTf_I | + | 1927 |
| chr12 | 92726857 | 92727850 | L1MdTf_I | + | 993 |
| chr12 | 94404185 | 94409248 | L1MdTf_III | + | 5063 |
| chr12 | 94409248 | 94410242 | L1MdTf_III | + | 994 |
| chr12 | 94479631 | 94480543 | L1MdTf_I | + | 912 |
| chr12 | 94480543 | 94481536 | L1MdTf_I | + | 993 |
| chr12 | 95215191 | 95221285 | L1MdTf_III | + | 6094 |
| chr12 | 95221285 | 95222280 | L1MdTf_III | + | 995 |
| chr12 | 95307312 | 95312792 | L1MdTf_I | + | 5480 |
| chr12 | 95312792 | 95313785 | L1MdTf_I | + | 993 |
| chr12 | 95788819 | 95795102 | L1MdTf_II | + | 6283 |
| chr12 | 95795102 | 95796095 | L1MdTf_I | + | 993 |
| chr12 | 96477522 | 96483526 | L1MdTf_I | + | 6004 |
| chr12 | 96483526 | 96484519 | L1MdTf_I | + | 993 |
| chr12 | 97004007 | 97010078 | L1MdTf_I | + | 6071 |
| chr12 | 97010078 | 97011071 | L1MdTf_I | + | 993 |
| chr12 | 98449683 | 98454590 | L1MdTf_III | + | 4907 |
| chr12 | 98454590 | 98455583 | L1MdTf_III | + | 993 |
| chr12 | 101413144 | 101418813 | L1MdTf_III | + | 5669 |
| chr12 | 101418813 | 101419808 | L1MdTf_III | + | 995 |
| chr12 | 101668501 | 101669430 | L1MdTf_II | - | 929 |
| chr12 | 101669430 | 101669650 | L1MdTf_II | + | 220 |
| chr12 | 104167880 | 104173580 | L1MdTf_III | + | 5700 |
| chr12 | 104173580 | 104174573 | L1MdTf_III | + | 993 |
| chr12 | 115689333 | 115694577 | L1MdTf_III | + | 5244 |
| chr12 | 115694577 | 115695570 | L1MdTf_III | + | 993 |
| chr12 | 118647150 | 118652780 | L1MdTf_II | + | 5630 |
| chr12 | 118652780 | 118653777 | L1MdTf_II | + | 997 |
| chr12 | 119309942 | 119315225 | L1MdTf_III | + | 5283 |
| chr12 | 119315225 | 119316217 | L1MdTf_III | + | 992 |
| chr13 | 3159379 | 3165903 | L1MdTf_II | + | 6524 |
| chr13 | 3165903 | 3166889 | L1MdTf_II | + | 986 |
| chr13 | 5031926 | 5038854 | L1MdTf_II | + | 6928 |
| chr13 | 5038854 | 5039874 | L1MdTf_II | + | 1020 |
| chr13 | 6485771 | 6491466 | L1MdTf_III | + | 5695 |
| chr13 | 6491466 | 6492482 | L1MdTf_III | + | 1016 |
| chr13 | 6539457 | 6541417 | L1MdTf_III | + | 1960 |
| chr13 | 6541417 | 6542398 | L1MdTf_III | + | 981 |
| chr13 | 6812115 | 6813532 | L1MdTf_I | + | 1417 |
| chr13 | 6813532 | 6814525 | L1MdTf_I | + | 993 |
| chr13 | 6828440 | 6834300 | L1MdTf_II | + | 5860 |
| chr13 | 6834300 | 6835293 | L1MdTf_II | + | 993 |

| | | | | | |
|---|---|---|---|---|---|
| chr13 | 8217053 | 8222263 | L1MdTf_II | + | 5210 |
| chr13 | 8222263 | 8223256 | L1MdTf_II | + | 993 |
| chr13 | 8422693 | 8424130 | L1MdTf_II | + | 1437 |
| chr13 | 8424130 | 8425123 | L1MdTf_II | + | 993 |
| chr13 | 16582931 | 16588141 | L1MdTf_III | + | 5210 |
| chr13 | 16588142 | 16589134 | L1MdTf_III | + | 992 |
| chr13 | 16674401 | 16680465 | L1MdTf_I | + | 6064 |
| chr13 | 16680465 | 16681458 | L1MdTf_I | + | 993 |
| chr13 | 17114578 | 17119834 | L1MdTf_II | + | 5256 |
| chr13 | 17119834 | 17120846 | L1MdTf_II | + | 1012 |
| chr13 | 18861941 | 18862927 | L1MdTf_III | - | 986 |
| chr13 | 18862927 | 18863131 | L1MdTf_III | + | 204 |
| chr13 | 19627910 | 19627936 | L1MdTf_III | + | 26 |
| chr13 | 19627936 | 19628383 | L1MdTf_I | - | 447 |
| chr13 | 19628383 | 19629108 | L1MdTf_III | + | 725 |
| chr13 | 19629109 | 19630125 | L1MdTf_III | + | 1016 |
| chr13 | 21044863 | 21050318 | L1MdTf_II | + | 5455 |
| chr13 | 21050318 | 21051313 | L1MdTf_II | + | 995 |
| chr13 | 21055333 | 21057077 | L1MdTf_II | + | 1744 |
| chr13 | 21057077 | 21058075 | L1MdTf_II | + | 998 |
| chr13 | 21513956 | 21519278 | L1MdTf_III | + | 5322 |
| chr13 | 21519278 | 21520271 | L1MdTf_III | + | 993 |
| chr13 | 22607137 | 22613035 | L1MdTf_I | + | 5898 |
| chr13 | 22613035 | 22614028 | L1MdTf_I | + | 993 |
| chr13 | 24212313 | 24217905 | L1MdTf_I | + | 5592 |
| chr13 | 24217905 | 24218898 | L1MdTf_I | + | 993 |
| chr13 | 26624581 | 26631225 | L1MdTf_I | + | 6644 |
| chr13 | 26631225 | 26632218 | L1MdTf_I | + | 993 |
| chr13 | 26705861 | 26711094 | L1MdTf_II | + | 5233 |
| chr13 | 26711094 | 26712078 | L1MdTf_II | + | 984 |
| chr13 | 26889093 | 26889168 | L1MdTf_II | - | 75 |
| chr13 | 26889168 | 26889232 | L1MdTf_II | + | 64 |
| chr13 | 29888864 | 29889943 | L1MdTf_I | + | 1079 |
| chr13 | 29889943 | 29890936 | L1MdTf_I | + | 993 |
| chr13 | 33498931 | 33500000 | L1MdTf_III | - | 1069 |
| chr13 | 33500000 | 33505024 | L1MdTf_III | - | 5024 |
| chr13 | 33557148 | 33563409 | L1MdTf_II | + | 6261 |
| chr13 | 33563409 | 33564402 | L1MdTf_II | + | 993 |
| chr13 | 33656723 | 33661955 | L1MdTf_III | + | 5232 |
| chr13 | 33661955 | 33662948 | L1MdTf_III | + | 993 |
| chr13 | 35751138 | 35756961 | L1MdTf_I | + | 5823 |
| chr13 | 35756961 | 35757954 | L1MdTf_I | + | 993 |
| chr13 | 36847647 | 36852910 | L1MdTf_III | + | 5263 |
| chr13 | 36852910 | 36853903 | L1MdTf_III | + | 993 |
| chr13 | 36980138 | 36980378 | L1MdTf_I | + | 240 |

| | | | | | |
|---|---|---|---|---|---|
| chr13 | 36980378 | 36981371 | L1MdTf_I | + | 993 |
| chr13 | 37446969 | 37452425 | L1MdTf_III | + | 5456 |
| chr13 | 37452425 | 37453417 | L1MdTf_III | + | 992 |
| chr13 | 39870724 | 39871742 | L1MdTf_I | + | 1018 |
| chr13 | 39871742 | 39872739 | L1MdTf_I | + | 997 |
| chr13 | 40093059 | 40098729 | L1MdTf_I | + | 5670 |
| chr13 | 40098729 | 40099726 | L1MdTf_I | + | 997 |
| chr13 | 40741123 | 40746580 | L1MdTf_II | + | 5457 |
| chr13 | 40746580 | 40747573 | L1MdTf_II | + | 993 |
| chr13 | 48450556 | 48456386 | L1MdTf_I | + | 5830 |
| chr13 | 48456386 | 48457379 | L1MdTf_I | + | 993 |
| chr13 | 50072047 | 50073241 | L1MdTf_III | + | 1194 |
| chr13 | 50073241 | 50074234 | L1MdTf_III | + | 993 |
| chr13 | 50191111 | 50196774 | L1MdTf_III | + | 5663 |
| chr13 | 50196774 | 50196928 | L1MdTf_III | + | 154 |
| chr13 | 50273364 | 50279458 | L1MdTf_II | + | 6094 |
| chr13 | 50279458 | 50280453 | L1MdTf_II | + | 995 |
| chr13 | 50447180 | 50452580 | L1MdTf_II | + | 5400 |
| chr13 | 50452580 | 50453573 | L1MdTf_II | + | 993 |
| chr13 | 61003655 | 61009319 | L1MdTf_II | + | 5664 |
| chr13 | 61009319 | 61010315 | L1MdTf_II | + | 996 |
| chr13 | 61545126 | 61550533 | L1MdTf_II | + | 5407 |
| chr13 | 61550533 | 61551529 | L1MdTf_II | + | 996 |
| chr13 | 61557498 | 61558015 | L1MdTf_III | - | 517 |
| chr13 | 61558015 | 61558655 | L1MdTf_III | + | 640 |
| chr13 | 62566659 | 62573144 | L1MdTf_II | + | 6485 |
| chr13 | 62573144 | 62574159 | L1MdTf_II | + | 1015 |
| chr13 | 62599361 | 62599955 | L1MdTf_II | - | 594 |
| chr13 | 62599955 | 62600189 | L1MdTf_II | + | 234 |
| chr13 | 63585456 | 63590685 | L1MdTf_I | + | 5229 |
| chr13 | 63590685 | 63591678 | L1MdTf_I | + | 993 |
| chr13 | 66296051 | 66301305 | L1MdTf_III | + | 5254 |
| chr13 | 66301305 | 66302298 | L1MdTf_III | + | 993 |
| chr13 | 66802485 | 66807727 | L1MdTf_III | + | 5242 |
| chr13 | 66807727 | 66808720 | L1MdTf_III | + | 993 |
| chr13 | 66982967 | 66988429 | L1MdTf_III | + | 5462 |
| chr13 | 66988429 | 66989422 | L1MdTf_III | + | 993 |
| chr13 | 67439825 | 67440246 | L1MdTf_III | - | 421 |
| chr13 | 67440246 | 67440657 | L1MdTf_III | + | 411 |
| chr13 | 68307346 | 68308130 | L1MdTf_III | - | 784 |
| chr13 | 68308130 | 68308874 | L1MdTf_III | + | 744 |
| chr13 | 68763450 | 68763904 | L1MdTf_III | - | 454 |
| chr13 | 68763904 | 68766864 | L1MdTf_III | + | 2960 |
| chr13 | 71072193 | 71076777 | L1MdTf_III | + | 4584 |
| chr13 | 71076777 | 71077774 | L1MdTf_III | + | 997 |

| | | | | | |
|---|---|---|---|---|---|
| chr13 | 71573202 | 71574173 | L1MdTf_II | + | 971 |
| chr13 | 71574174 | 71575166 | L1MdTf_II | + | 992 |
| chr13 | 71576378 | 71582032 | L1MdTf_II | + | 5654 |
| chr13 | 71582032 | 71583046 | L1MdTf_II | + | 1014 |
| chr13 | 71639764 | 71645236 | L1MdTf_I | + | 5472 |
| chr13 | 71645236 | 71646229 | L1MdTf_I | + | 993 |
| chr13 | 72027631 | 72033053 | L1MdTf_III | + | 5422 |
| chr13 | 72033053 | 72034044 | L1MdTf_III | + | 991 |
| chr13 | 72352427 | 72354250 | L1MdTf_III | + | 1823 |
| chr13 | 72354250 | 72355243 | L1MdTf_III | + | 993 |
| chr13 | 74853714 | 74859337 | L1MdTf_II | + | 5623 |
| chr13 | 74859337 | 74860321 | L1MdTf_II | + | 984 |
| chr13 | 75328386 | 75329109 | L1MdTf_III | + | 723 |
| chr13 | 75329109 | 75330101 | L1MdTf_III | + | 992 |
| chr13 | 77577002 | 77582349 | L1MdTf_III | + | 5347 |
| chr13 | 77582349 | 77583327 | L1MdTf_III | + | 978 |
| chr13 | 77685948 | 77691607 | L1MdTf_I | + | 5659 |
| chr13 | 77691607 | 77692620 | L1MdTf_I | + | 1013 |
| chr13 | 77721332 | 77723263 | L1MdTf_I | + | 1931 |
| chr13 | 77723263 | 77724256 | L1MdTf_I | + | 993 |
| chr13 | 78409319 | 78414541 | L1MdTf_III | + | 5222 |
| chr13 | 78414542 | 78415532 | L1MdTf_III | + | 990 |
| chr13 | 78419460 | 78419800 | L1MdTf_II | - | 340 |
| chr13 | 78419800 | 78420948 | L1MdTf_II | + | 1148 |
| chr13 | 79093815 | 79099504 | L1MdTf_III | + | 5689 |
| chr13 | 79099505 | 79100498 | L1MdTf_III | + | 993 |
| chr13 | 79569950 | 79575111 | L1MdTf_I | + | 5161 |
| chr13 | 79575111 | 79576104 | L1MdTf_I | + | 993 |
| chr13 | 79686801 | 79686985 | L1MdTf_III | + | 184 |
| chr13 | 79686985 | 79687994 | L1MdTf_III | + | 1009 |
| chr13 | 80172142 | 80178030 | L1MdTf_II | + | 5888 |
| chr13 | 80178030 | 80179025 | L1MdTf_II | + | 995 |
| chr13 | 81343062 | 81348297 | L1MdTf_I | + | 5235 |
| chr13 | 81348297 | 81349290 | L1MdTf_I | + | 993 |
| chr13 | 81349292 | 81351166 | L1MdTf_III | + | 1874 |
| chr13 | 81351166 | 81352157 | L1MdTf_III | + | 991 |
| chr13 | 81601030 | 81606267 | L1MdTf_III | + | 5237 |
| chr13 | 81606267 | 81607261 | L1MdTf_III | + | 994 |
| chr13 | 82488811 | 82489464 | L1MdTf_III | + | 653 |
| chr13 | 82489464 | 82490485 | L1MdTf_III | + | 1021 |
| chr13 | 82697674 | 82698038 | L1MdTf_III | + | 364 |
| chr13 | 82698038 | 82699031 | L1MdTf_III | + | 993 |
| chr13 | 83960874 | 83966079 | L1MdTf_II | + | 5205 |
| chr13 | 83966079 | 83967091 | L1MdTf_II | + | 1012 |
| chr13 | 84175151 | 84176988 | L1MdTf_III | + | 1837 |

| | | | | | |
|---|---|---|---|---|---|
| chr13 | 84176988 | 84177981 | L1MdTf_III | + | 993 |
| chr13 | 84191027 | 84191429 | L1MdTf_II | - | 402 |
| chr13 | 84191429 | 84191685 | L1MdTf_II | + | 256 |
| chr13 | 84261272 | 84262744 | L1MdTf_I | + | 1472 |
| chr13 | 84262744 | 84263737 | L1MdTf_I | + | 993 |
| chr13 | 84589101 | 84594723 | L1MdTf_II | + | 5622 |
| chr13 | 84594723 | 84595716 | L1MdTf_II | + | 993 |
| chr13 | 85187669 | 85192912 | L1MdTf_III | + | 5243 |
| chr13 | 85192912 | 85193905 | L1MdTf_III | + | 993 |
| chr13 | 86436179 | 86441681 | L1MdTf_III | + | 5502 |
| chr13 | 86441682 | 86442665 | L1MdTf_III | + | 983 |
| chr13 | 86655125 | 86657103 | L1MdTf_II | + | 1978 |
| chr13 | 86657103 | 86658096 | L1MdTf_II | + | 993 |
| chr13 | 87717575 | 87723656 | L1MdTf_I | + | 6081 |
| chr13 | 87723656 | 87724649 | L1MdTf_I | + | 993 |
| chr13 | 89589357 | 89589511 | L1MdTf_III | + | 154 |
| chr13 | 89589511 | 89590504 | L1MdTf_III | + | 993 |
| chr13 | 90010108 | 90015488 | L1MdTf_II | + | 5380 |
| chr13 | 90015488 | 90016484 | L1MdTf_II | + | 996 |
| chr13 | 90191012 | 90196665 | L1MdTf_I | + | 5653 |
| chr13 | 90196665 | 90197658 | L1MdTf_I | + | 993 |
| chr13 | 90741583 | 90747033 | L1MdTf_II | + | 5450 |
| chr13 | 90747033 | 90748026 | L1MdTf_II | + | 993 |
| chr13 | 91963299 | 91968726 | L1MdTf_II | + | 5427 |
| chr13 | 91968726 | 91969719 | L1MdTf_II | + | 993 |
| chr13 | 92094450 | 92099102 | L1MdTf_II | + | 4652 |
| chr13 | 92099102 | 92100095 | L1MdTf_I | + | 993 |
| chr13 | 101117889 | 101122831 | L1MdTf_I | + | 4942 |
| chr13 | 101122831 | 101123823 | L1MdTf_I | + | 992 |
| chr13 | 102557836 | 102563470 | L1MdTf_I | + | 5634 |
| chr13 | 102563470 | 102564491 | L1MdTf_I | + | 1021 |
| chr13 | 103718681 | 103723997 | L1MdTf_III | + | 5316 |
| chr13 | 103723997 | 103724991 | L1MdTf_III | + | 994 |
| chr13 | 104621064 | 104624972 | L1MdTf_III | + | 3908 |
| chr13 | 104624972 | 104625965 | L1MdTf_III | + | 993 |
| chr13 | 105929639 | 105935096 | L1MdTf_III | + | 5457 |
| chr13 | 105935096 | 105936089 | L1MdTf_III | + | 993 |
| chr13 | 106258382 | 106258733 | L1MdTf_III | + | 351 |
| chr13 | 106258733 | 106259726 | L1MdTf_III | + | 993 |
| chr13 | 106433549 | 106435906 | L1MdTf_III | + | 2357 |
| chr13 | 106435906 | 106436899 | L1MdTf_III | + | 993 |
| chr13 | 106786733 | 106792329 | L1MdTf_I | + | 5596 |
| chr13 | 106792329 | 106793322 | L1MdTf_I | + | 993 |
| chr13 | 106816143 | 106821606 | L1MdTf_III | + | 5463 |
| chr13 | 106821606 | 106822599 | L1MdTf_III | + | 993 |

| | | | | | |
|---|---|---|---|---|---|
| chr13 | 107656705 | 107656951 | L1MdTf_I | + | 246 |
| chr13 | 107656951 | 107657944 | L1MdTf_I | + | 993 |
| chr13 | 108704088 | 108704710 | L1MdTf_II | - | 622 |
| chr13 | 108704710 | 108705597 | L1MdTf_II | + | 887 |
| chr13 | 109546504 | 109551923 | L1MdTf_II | + | 5419 |
| chr13 | 109551923 | 109552916 | L1MdTf_II | + | 993 |
| chr13 | 110446037 | 110451716 | L1MdTf_III | + | 5679 |
| chr13 | 110451716 | 110452709 | L1MdTf_III | + | 993 |
| chr13 | 111425686 | 111425831 | L1MdTf_I | + | 145 |
| chr13 | 111425831 | 111426854 | L1MdTf_I | + | 1023 |
| chr13 | 114726414 | 114728340 | L1MdTf_I | + | 1926 |
| chr13 | 114728340 | 114729333 | L1MdTf_I | + | 993 |
| chr13 | 116601915 | 116604398 | L1MdTf_I | + | 2483 |
| chr13 | 116604398 | 116605391 | L1MdTf_I | + | 993 |
| chr13 | 116637392 | 116637524 | L1MdTf_II | + | 132 |
| chr13 | 116637524 | 116638517 | L1MdTf_II | + | 993 |
| chr13 | 118481757 | 118481854 | L1MdTf_III | + | 97 |
| chr13 | 118481854 | 118482847 | L1MdTf_III | + | 993 |
| chr13 | 120751829 | 120757456 | L1MdTf_II | + | 5627 |
| chr13 | 120757456 | 120758449 | L1MdTf_II | + | 993 |
| chr14 | 3451932 | 3452213 | L1MdTf_III | - | 281 |
| chr14 | 3452213 | 3452266 | L1MdTf_III | + | 53 |
| chr14 | 4922763 | 4923430 | L1MdTf_II | + | 667 |
| chr14 | 4923430 | 4924422 | L1MdTf_II | + | 992 |
| chr14 | 6959737 | 6965132 | L1MdTf_III | + | 5395 |
| chr14 | 6965132 | 6966125 | L1MdTf_III | + | 993 |
| chr14 | 7357304 | 7357743 | L1MdTf_II | + | 439 |
| chr14 | 7357743 | 7358739 | L1MdTf_II | + | 996 |
| chr14 | 7375603 | 7380609 | L1MdTf_I | + | 5006 |
| chr14 | 7380609 | 7381602 | L1MdTf_I | + | 993 |
| chr14 | 8986296 | 8986758 | L1MdTf_III | - | 462 |
| chr14 | 8986758 | 8987811 | L1MdTf_III | + | 1053 |
| chr14 | 10211756 | 10214015 | L1MdTf_I | + | 2259 |
| chr14 | 10214015 | 10215008 | L1MdTf_I | + | 993 |
| chr14 | 11972497 | 11979022 | L1MdTf_III | + | 6525 |
| chr14 | 11979022 | 11980017 | L1MdTf_III | + | 995 |
| chr14 | 12361879 | 12367325 | L1MdTf_II | + | 5446 |
| chr14 | 12367325 | 12368318 | L1MdTf_II | + | 993 |
| chr14 | 12463716 | 12464038 | L1MdTf_III | - | 322 |
| chr14 | 12464038 | 12464068 | L1MdTf_III | + | 30 |
| chr14 | 12609308 | 12614742 | L1MdTf_II | + | 5434 |
| chr14 | 12614742 | 12615752 | L1MdTf_II | + | 1010 |
| chr14 | 13298276 | 13301599 | L1MdTf_I | + | 3323 |
| chr14 | 13301599 | 13302592 | L1MdTf_I | + | 993 |
| chr14 | 15119426 | 15120065 | L1MdTf_III | - | 639 |

| | | | | | |
|---|---|---|---|---|---|
| chr14 | 15120065 | 15121112 | L1MdTf_III | + | 1047 |
| chr14 | 15544818 | 15545458 | L1MdTf_III | - | 640 |
| chr14 | 15545458 | 15546500 | L1MdTf_III | + | 1042 |
| chr14 | 15677304 | 15677944 | L1MdTf_III | - | 640 |
| chr14 | 15677944 | 15678985 | L1MdTf_III | + | 1041 |
| chr14 | 15839596 | 15845023 | L1MdTf_I | + | 5427 |
| chr14 | 15845023 | 15846015 | L1MdTf_I | + | 992 |
| chr14 | 16325981 | 16326622 | L1MdTf_III | - | 641 |
| chr14 | 16326622 | 16327664 | L1MdTf_III | + | 1042 |
| chr14 | 17898001 | 17902429 | L1MdTf_I | + | 4428 |
| chr14 | 17902429 | 17903422 | L1MdTf_I | + | 993 |
| chr14 | 18168770 | 18169578 | L1MdTf_III | - | 808 |
| chr14 | 18169578 | 18170218 | L1MdTf_III | + | 640 |
| chr14 | 18849731 | 18855043 | L1MdTf_III | + | 5312 |
| chr14 | 18855043 | 18856036 | L1MdTf_III | + | 993 |
| chr14 | 19089140 | 19094452 | L1MdTf_III | + | 5312 |
| chr14 | 19094452 | 19095445 | L1MdTf_III | + | 993 |
| chr14 | 19292530 | 19297742 | L1MdTf_I | + | 5212 |
| chr14 | 19297742 | 19298735 | L1MdTf_I | + | 993 |
| chr14 | 19302313 | 19307625 | L1MdTf_III | + | 5312 |
| chr14 | 19307625 | 19308618 | L1MdTf_III | + | 993 |
| chr14 | 19353314 | 19358818 | L1MdTf_III | + | 5504 |
| chr14 | 19358818 | 19359811 | L1MdTf_III | + | 993 |
| chr14 | 19361348 | 19366660 | L1MdTf_III | + | 5312 |
| chr14 | 19366660 | 19367653 | L1MdTf_III | + | 993 |
| chr14 | 19522119 | 19527562 | L1MdTf_I | + | 5443 |
| chr14 | 19527562 | 19528555 | L1MdTf_I | + | 993 |
| chr14 | 33879942 | 33880116 | L1MdTf_III | - | 174 |
| chr14 | 33880116 | 33881305 | L1MdTf_III | + | 1189 |
| chr14 | 34997198 | 35000000 | L1MdTf_II | + | 2802 |
| chr14 | 35000000 | 35002448 | L1MdTf_I | + | 2448 |
| chr14 | 36159088 | 36160445 | L1MdTf_III | - | 1357 |
| chr14 | 36160445 | 36161314 | L1MdTf_III | + | 869 |
| chr14 | 37808725 | 37808923 | L1MdTf_III | + | 198 |
| chr14 | 37808923 | 37809916 | L1MdTf_III | + | 993 |
| chr14 | 39818981 | 39825077 | L1MdTf_II | + | 6096 |
| chr14 | 39825077 | 39826070 | L1MdTf_II | + | 993 |
| chr14 | 39905823 | 39911261 | L1MdTf_II | + | 5438 |
| chr14 | 39911261 | 39912245 | L1MdTf_II | + | 984 |
| chr14 | 40268559 | 40275100 | L1MdTf_II | + | 6541 |
| chr14 | 40275100 | 40276093 | L1MdTf_II | + | 993 |
| chr14 | 40969750 | 40972835 | L1MdTf_I | + | 3085 |
| chr14 | 40972835 | 40973828 | L1MdTf_I | + | 993 |
| chr14 | 43075759 | 43081172 | L1MdTf_I | + | 5413 |
| chr14 | 43081172 | 43082164 | L1MdTf_I | + | 992 |

| | | | | | |
|---|---|---|---|---|---|
| chr14 | 43823287 | 43829174 | L1MdTf_I | + | 5887 |
| chr14 | 43829174 | 43830167 | L1MdTf_I | + | 993 |
| chr14 | 43983377 | 43983613 | L1MdTf_II | + | 236 |
| chr14 | 43983613 | 43984608 | L1MdTf_II | + | 995 |
| chr14 | 44470766 | 44475772 | L1MdTf_II | + | 5006 |
| chr14 | 44475772 | 44476776 | L1MdTf_II | + | 1004 |
| chr14 | 44536421 | 44542571 | L1MdTf_II | + | 6150 |
| chr14 | 44542571 | 44543565 | L1MdTf_II | + | 994 |
| chr14 | 46983239 | 46988662 | L1MdTf_III | + | 5423 |
| chr14 | 46988662 | 46989655 | L1MdTf_III | + | 993 |
| chr14 | 49865975 | 49871506 | L1MdTf_III | + | 5531 |
| chr14 | 49871506 | 49872499 | L1MdTf_III | + | 993 |
| chr14 | 50034666 | 50035419 | L1MdTf_III | + | 753 |
| chr14 | 50035419 | 50036412 | L1MdTf_III | + | 993 |
| chr14 | 50726575 | 50728714 | L1MdTf_III | + | 2139 |
| chr14 | 50728714 | 50729707 | L1MdTf_III | + | 993 |
| chr14 | 50776665 | 50778072 | L1MdTf_III | + | 1407 |
| chr14 | 50778072 | 50779065 | L1MdTf_III | + | 993 |
| chr14 | 50874138 | 50875485 | L1MdTf_III | + | 1347 |
| chr14 | 50875485 | 50876478 | L1MdTf_III | + | 993 |
| chr14 | 51725364 | 51730798 | L1MdTf_III | + | 5434 |
| chr14 | 51730798 | 51731793 | L1MdTf_III | + | 995 |
| chr14 | 51940839 | 51945980 | L1MdTf_III | + | 5141 |
| chr14 | 51945980 | 51946973 | L1MdTf_III | + | 993 |
| chr14 | 56432863 | 56438714 | L1MdTf_I | + | 5851 |
| chr14 | 56438714 | 56439707 | L1MdTf_I | + | 993 |
| chr14 | 58563119 | 58568111 | L1MdTf_II | + | 4992 |
| chr14 | 58568111 | 58569104 | L1MdTf_II | + | 993 |
| chr14 | 58824815 | 58830303 | L1MdTf_III | + | 5488 |
| chr14 | 58830303 | 58831296 | L1MdTf_III | + | 993 |
| chr14 | 59402243 | 59407747 | L1MdTf_III | + | 5504 |
| chr14 | 59407747 | 59408738 | L1MdTf_III | + | 991 |
| chr14 | 60588331 | 60593737 | L1MdTf_I | + | 5406 |
| chr14 | 60593737 | 60594730 | L1MdTf_I | + | 993 |
| chr14 | 61112919 | 61118129 | L1MdTf_I | + | 5210 |
| chr14 | 61118129 | 61119148 | L1MdTf_I | + | 1019 |
| chr14 | 73185456 | 73191096 | L1MdTf_II | + | 5640 |
| chr14 | 73191096 | 73192089 | L1MdTf_II | + | 993 |
| chr14 | 73192096 | 73196652 | L1MdTf_II | + | 4556 |
| chr14 | 73196652 | 73197645 | L1MdTf_II | + | 993 |
| chr14 | 77536826 | 77542249 | L1MdTf_III | + | 5423 |
| chr14 | 77542249 | 77543242 | L1MdTf_III | + | 993 |
| chr14 | 78445723 | 78451193 | L1MdTf_III | + | 5470 |
| chr14 | 78451193 | 78452186 | L1MdTf_III | + | 993 |
| chr14 | 79234416 | 79239783 | L1MdTf_I | + | 5367 |

| | | | | | |
|---|---|---|---|---|---|
| chr14 | 79239783 | 79240776 | L1MdTf_I | + | 993 |
| chr14 | 80331998 | 80333145 | L1MdTf_I | + | 1147 |
| chr14 | 80333145 | 80334138 | L1MdTf_I | + | 993 |
| chr14 | 80423068 | 80423704 | L1MdTf_I | - | 636 |
| chr14 | 80423704 | 80424478 | L1MdTf_II | + | 774 |
| chr14 | 81887847 | 81888532 | L1MdTf_III | - | 685 |
| chr14 | 81888533 | 81888740 | L1MdTf_III | + | 207 |
| chr14 | 82779960 | 82785410 | L1MdTf_III | + | 5450 |
| chr14 | 82785410 | 82786394 | L1MdTf_III | + | 984 |
| chr14 | 83195133 | 83200563 | L1MdTf_II | + | 5430 |
| chr14 | 83200563 | 83201555 | L1MdTf_II | + | 992 |
| chr14 | 83434918 | 83440044 | L1MdTf_I | + | 5126 |
| chr14 | 83440044 | 83441037 | L1MdTf_I | + | 993 |
| chr14 | 83792244 | 83793651 | L1MdTf_I | - | 1407 |
| chr14 | 83793651 | 83794631 | L1MdTf_I | + | 980 |
| chr14 | 83934333 | 83938049 | L1MdTf_II | + | 3716 |
| chr14 | 83938049 | 83939042 | L1MdTf_II | + | 993 |
| chr14 | 84605591 | 84611156 | L1MdTf_III | + | 5565 |
| chr14 | 84611156 | 84612149 | L1MdTf_III | + | 993 |
| chr14 | 85658311 | 85658876 | L1MdTf_II | - | 565 |
| chr14 | 85658876 | 85659155 | L1MdTf_II | + | 279 |
| chr14 | 86190733 | 86196022 | L1MdTf_III | + | 5289 |
| chr14 | 86196023 | 86197014 | L1MdTf_III | + | 991 |
| chr14 | 86312113 | 86312463 | L1MdTf_I | + | 350 |
| chr14 | 86312463 | 86313456 | L1MdTf_I | + | 993 |
| chr14 | 86582014 | 86582959 | L1MdTf_III | - | 945 |
| chr14 | 86582959 | 86583297 | L1MdTf_III | + | 338 |
| chr14 | 87177862 | 87183331 | L1MdTf_III | + | 5469 |
| chr14 | 87183331 | 87184324 | L1MdTf_III | + | 993 |
| chr14 | 88505246 | 88506320 | L1MdTf_III | - | 1074 |
| chr14 | 88506321 | 88506971 | L1MdTf_III | + | 650 |
| chr14 | 88675242 | 88675986 | L1MdTf_III | - | 744 |
| chr14 | 88675986 | 88676484 | L1MdTf_III | + | 498 |
| chr14 | 88850786 | 88856391 | L1MdTf_III | + | 5605 |
| chr14 | 88856391 | 88857382 | L1MdTf_III | + | 991 |
| chr14 | 91363250 | 91368634 | L1MdTf_III | + | 5384 |
| chr14 | 91368634 | 91369627 | L1MdTf_III | + | 993 |
| chr14 | 91993565 | 91996440 | L1MdTf_III | - | 2875 |
| chr14 | 91996440 | 91997160 | L1MdTf_III | + | 720 |
| chr14 | 92520421 | 92521300 | L1MdTf_III | + | 879 |
| chr14 | 92521300 | 92522293 | L1MdTf_III | + | 993 |
| chr14 | 92571975 | 92572116 | L1MdTf_III | + | 141 |
| chr14 | 92572116 | 92573107 | L1MdTf_III | + | 991 |
| chr14 | 92895979 | 92896810 | L1MdTf_III | + | 831 |
| chr14 | 92896810 | 92897803 | L1MdTf_III | + | 993 |

| | | | | | |
|---|---|---|---|---|---|
| chr14 | 93026488 | 93026540 | L1MdTf_III | + | 52 |
| chr14 | 93026540 | 93027551 | L1MdTf_III | + | 1011 |
| chr14 | 93725759 | 93726373 | L1MdTf_III | + | 614 |
| chr14 | 93726373 | 93727357 | L1MdTf_III | + | 984 |
| chr14 | 95970455 | 95971042 | L1MdTf_III | - | 587 |
| chr14 | 95971042 | 95971606 | L1MdTf_III | + | 564 |
| chr14 | 96131471 | 96131893 | L1MdTf_III | + | 422 |
| chr14 | 96131893 | 96132885 | L1MdTf_III | + | 992 |
| chr14 | 98788248 | 98791782 | L1MdTf_III | + | 3534 |
| chr14 | 98791782 | 98792775 | L1MdTf_III | + | 993 |
| chr14 | 99444526 | 99444650 | L1MdTf_III | + | 124 |
| chr14 | 99444650 | 99445667 | L1MdTf_III | + | 1017 |
| chr14 | 100616539 | 100622894 | L1MdTf_I | - | 6355 |
| chr14 | 100622895 | 100627235 | L1MdTf_II | - | 4340 |
| chr14 | 102796727 | 102801909 | L1MdTf_III | + | 5182 |
| chr14 | 102801909 | 102802903 | L1MdTf_III | + | 994 |
| chr14 | 104359151 | 104364573 | L1MdTf_I | + | 5422 |
| chr14 | 104364573 | 104365566 | L1MdTf_I | + | 993 |
| chr14 | 106256297 | 106257199 | L1MdTf_II | - | 902 |
| chr14 | 106257199 | 106257863 | L1MdTf_II | + | 664 |
| chr14 | 107094183 | 107094639 | L1MdTf_II | + | 456 |
| chr14 | 107094639 | 107095632 | L1MdTf_II | + | 993 |
| chr14 | 107201387 | 107207729 | L1MdTf_I | + | 6342 |
| chr14 | 107207729 | 107208722 | L1MdTf_I | + | 993 |
| chr14 | 107444485 | 107449835 | L1MdTf_III | + | 5350 |
| chr14 | 107449835 | 107450828 | L1MdTf_III | + | 993 |
| chr14 | 107994076 | 108000000 | L1MdTf_II | + | 5924 |
| chr14 | 108000000 | 108000725 | L1MdTf_I | + | 725 |
| chr14 | 109818484 | 109819559 | L1MdTf_III | + | 1075 |
| chr14 | 109819560 | 109820549 | L1MdTf_III | + | 989 |
| chr14 | 111141793 | 111147049 | L1MdTf_III | + | 5256 |
| chr14 | 111147049 | 111148042 | L1MdTf_III | + | 993 |
| chr14 | 111461536 | 111462575 | L1MdTf_II | + | 1039 |
| chr14 | 111462575 | 111463559 | L1MdTf_II | + | 984 |
| chr14 | 112140620 | 112146249 | L1MdTf_I | + | 5629 |
| chr14 | 112146249 | 112147242 | L1MdTf_I | + | 993 |
| chr14 | 112960767 | 112961749 | L1MdTf_II | - | 982 |
| chr14 | 112961749 | 112962205 | L1MdTf_II | + | 456 |
| chr14 | 113522837 | 113523989 | L1MdTf_III | - | 1152 |
| chr14 | 113523989 | 113524981 | L1MdTf_III | + | 992 |
| chr14 | 114212313 | 114217685 | L1MdTf_II | + | 5372 |
| chr14 | 114217685 | 114218681 | L1MdTf_II | + | 996 |
| chr14 | 115171280 | 115171841 | L1MdTf_I | - | 561 |
| chr14 | 115171841 | 115171973 | L1MdTf_II | + | 132 |
| chr14 | 116562876 | 116568174 | L1MdTf_III | + | 5298 |

| | | | | | |
|---|---|---|---|---|---|
| chr14 | 116568174 | 116569167 | L1MdTf_III | + | 993 |
| chr14 | 117025847 | 117031504 | L1MdTf_II | + | 5657 |
| chr14 | 117031504 | 117032500 | L1MdTf_II | + | 996 |
| chr14 | 117495634 | 117500000 | L1MdTf_I | + | 4366 |
| chr14 | 117500000 | 117501062 | L1MdTf_II | + | 1062 |
| chr14 | 120299330 | 120304581 | L1MdTf_II | + | 5251 |
| chr14 | 120304581 | 120305565 | L1MdTf_II | + | 984 |
| chr14 | 123651116 | 123651944 | L1MdTf_I | + | 828 |
| chr14 | 123651944 | 123652937 | L1MdTf_I | + | 993 |
| chr14 | 123846747 | 123852060 | L1MdTf_I | + | 5313 |
| chr14 | 123852060 | 123853053 | L1MdTf_I | + | 993 |
| chr14 | 123967701 | 123971919 | L1MdTf_I | + | 4218 |
| chr14 | 123971919 | 123972934 | L1MdTf_I | + | 1015 |
| chr14 | 124058509 | 124063802 | L1MdTf_III | + | 5293 |
| chr14 | 124063802 | 124064786 | L1MdTf_III | + | 984 |
| chr14 | 124728761 | 124734147 | L1MdTf_II | + | 5386 |
| chr14 | 124734147 | 124735147 | L1MdTf_II | + | 1000 |
| chr14 | 124891643 | 124897696 | L1MdTf_I | + | 6053 |
| chr14 | 124897696 | 124898689 | L1MdTf_I | + | 993 |
| chr14 | 125030351 | 125034693 | L1MdTf_II | + | 4342 |
| chr14 | 125034693 | 125035686 | L1MdTf_II | + | 993 |
| chr15 | 3507419 | 3510271 | L1MdTf_II | + | 2852 |
| chr15 | 3510271 | 3511287 | L1MdTf_II | + | 1016 |
| chr15 | 3949771 | 3955639 | L1MdTf_I | + | 5868 |
| chr15 | 3955639 | 3956651 | L1MdTf_I | + | 1012 |
| chr15 | 5255032 | 5255586 | L1MdTf_III | + | 554 |
| chr15 | 5255586 | 5256579 | L1MdTf_III | + | 993 |
| chr15 | 5296339 | 5302167 | L1MdTf_II | + | 5828 |
| chr15 | 5302167 | 5303160 | L1MdTf_II | + | 993 |
| chr15 | 7878846 | 7884076 | L1MdTf_II | + | 5230 |
| chr15 | 7884076 | 7885069 | L1MdTf_II | + | 993 |
| chr15 | 9394621 | 9399847 | L1MdTf_III | + | 5226 |
| chr15 | 9399847 | 9400839 | L1MdTf_III | + | 992 |
| chr15 | 9813859 | 9814425 | L1MdTf_III | - | 566 |
| chr15 | 9814425 | 9814756 | L1MdTf_III | + | 331 |
| chr15 | 13687793 | 13690919 | L1MdTf_II | + | 3126 |
| chr15 | 13690919 | 13691914 | L1MdTf_II | + | 995 |
| chr15 | 14169201 | 14171892 | L1MdTf_II | + | 2691 |
| chr15 | 14171892 | 14172838 | L1MdTf_I | + | 946 |
| chr15 | 15004031 | 15005397 | L1MdTf_II | - | 1366 |
| chr15 | 15005397 | 15005928 | L1MdTf_II | + | 531 |
| chr15 | 15265459 | 15270710 | L1MdTf_III | + | 5251 |
| chr15 | 15270710 | 15271703 | L1MdTf_III | + | 993 |
| chr15 | 15914617 | 15919827 | L1MdTf_II | + | 5210 |
| chr15 | 15919827 | 15920811 | L1MdTf_II | + | 984 |

| | | | | | |
|---|---|---|---|---|---|
| chr15 | 16082212 | 16083542 | L1MdTf_III | + | 1330 |
| chr15 | 16083542 | 16084535 | L1MdTf_III | + | 993 |
| chr15 | 17266495 | 17272748 | L1MdTf_II | + | 6253 |
| chr15 | 17272748 | 17273745 | L1MdTf_II | + | 997 |
| chr15 | 17941999 | 17942360 | L1MdTf_III | - | 361 |
| chr15 | 17942360 | 17943060 | L1MdTf_III | + | 700 |
| chr15 | 18238343 | 18238584 | L1MdTf_III | + | 241 |
| chr15 | 18238585 | 18239561 | L1MdTf_III | + | 976 |
| chr15 | 18314610 | 18320062 | L1MdTf_I | + | 5452 |
| chr15 | 18320062 | 18321055 | L1MdTf_I | + | 993 |
| chr15 | 19843695 | 19848936 | L1MdTf_III | + | 5241 |
| chr15 | 19848936 | 19849929 | L1MdTf_III | + | 993 |
| chr15 | 21262133 | 21267344 | L1MdTf_II | + | 5211 |
| chr15 | 21267344 | 21268337 | L1MdTf_II | + | 993 |
| chr15 | 21450141 | 21453410 | L1MdTf_I | + | 3269 |
| chr15 | 21453410 | 21454403 | L1MdTf_I | + | 993 |
| chr15 | 21905887 | 21906471 | L1MdTf_III | - | 584 |
| chr15 | 21906471 | 21907172 | L1MdTf_III | + | 701 |
| chr15 | 22487952 | 22493441 | L1MdTf_III | + | 5489 |
| chr15 | 22493441 | 22494434 | L1MdTf_III | + | 993 |
| chr15 | 22962433 | 22963137 | L1MdTf_I | - | 704 |
| chr15 | 22963137 | 22963682 | L1MdTf_I | + | 545 |
| chr15 | 24834908 | 24836351 | L1MdTf_II | - | 1443 |
| chr15 | 24836351 | 24837139 | L1MdTf_II | + | 788 |
| chr15 | 26112717 | 26118356 | L1MdTf_II | + | 5639 |
| chr15 | 26118356 | 26119346 | L1MdTf_II | + | 990 |
| chr15 | 26280340 | 26285997 | L1MdTf_III | + | 5657 |
| chr15 | 26285997 | 26286990 | L1MdTf_III | + | 993 |
| chr15 | 27053125 | 27054741 | L1MdTf_II | + | 1616 |
| chr15 | 27054741 | 27055725 | L1MdTf_II | + | 984 |
| chr15 | 27638245 | 27638726 | L1MdTf_II | + | 481 |
| chr15 | 27638727 | 27639741 | L1MdTf_II | + | 1014 |
| chr15 | 29648329 | 29653538 | L1MdTf_I | + | 5209 |
| chr15 | 29653538 | 29654531 | L1MdTf_I | + | 993 |
| chr15 | 30251624 | 30256882 | L1MdTf_III | + | 5258 |
| chr15 | 30256882 | 30257875 | L1MdTf_III | + | 993 |
| chr15 | 32833125 | 32838764 | L1MdTf_II | + | 5639 |
| chr15 | 32838764 | 32839757 | L1MdTf_II | + | 993 |
| chr15 | 32856888 | 32862181 | L1MdTf_III | + | 5293 |
| chr15 | 32862181 | 32863174 | L1MdTf_III | + | 993 |
| chr15 | 33710360 | 33710542 | L1MdTf_I | + | 182 |
| chr15 | 33710542 | 33711535 | L1MdTf_I | + | 993 |
| chr15 | 34763245 | 34768659 | L1MdTf_III | + | 5414 |
| chr15 | 34768659 | 34769635 | L1MdTf_III | + | 976 |
| chr15 | 35395476 | 35396305 | L1MdTf_III | - | 829 |

| | | | | | |
|---|---|---|---|---|---|
| chr15 | 35396305 | 35396933 | L1MdTf_III | + | 628 |
| chr15 | 35783918 | 35785850 | L1MdTf_II | + | 1932 |
| chr15 | 35785850 | 35786834 | L1MdTf_II | + | 984 |
| chr15 | 36171305 | 36171650 | L1MdTf_II | + | 345 |
| chr15 | 36171650 | 36172671 | L1MdTf_II | + | 1021 |
| chr15 | 37860632 | 37866043 | L1MdTf_III | + | 5411 |
| chr15 | 37866043 | 37867036 | L1MdTf_III | + | 993 |
| chr15 | 39359379 | 39361791 | L1MdTf_III | + | 2412 |
| chr15 | 39361791 | 39362782 | L1MdTf_III | + | 991 |
| chr15 | 41107716 | 41113211 | L1MdTf_III | + | 5495 |
| chr15 | 41113211 | 41114204 | L1MdTf_III | + | 993 |
| chr15 | 43004788 | 43010469 | L1MdTf_I | + | 5681 |
| chr15 | 43010469 | 43011462 | L1MdTf_I | + | 993 |
| chr15 | 43456489 | 43461731 | L1MdTf_II | + | 5242 |
| chr15 | 43461731 | 43462724 | L1MdTf_II | + | 993 |
| chr15 | 44087671 | 44087737 | L1MdTf_I | - | 66 |
| chr15 | 44087737 | 44087812 | L1MdTf_II | + | 75 |
| chr15 | 44180946 | 44181773 | L1MdTf_II | + | 827 |
| chr15 | 44181773 | 44182766 | L1MdTf_II | + | 993 |
| chr15 | 45440274 | 45442290 | L1MdTf_III | + | 2016 |
| chr15 | 45442290 | 45443283 | L1MdTf_III | + | 993 |
| chr15 | 45793078 | 45794902 | L1MdTf_I | + | 1824 |
| chr15 | 45794902 | 45795895 | L1MdTf_I | + | 993 |
| chr15 | 46226893 | 46232118 | L1MdTf_III | + | 5225 |
| chr15 | 46232118 | 46233104 | L1MdTf_III | + | 986 |
| chr15 | 46337636 | 46337967 | L1MdTf_III | + | 331 |
| chr15 | 46337967 | 46338960 | L1MdTf_III | + | 993 |
| chr15 | 47043993 | 47049396 | L1MdTf_II | + | 5403 |
| chr15 | 47049396 | 47050389 | L1MdTf_II | + | 993 |
| chr15 | 47113124 | 47118457 | L1MdTf_II | + | 5333 |
| chr15 | 47118457 | 47119451 | L1MdTf_II | + | 994 |
| chr15 | 47842158 | 47842605 | L1MdTf_II | - | 447 |
| chr15 | 47842605 | 47843060 | L1MdTf_II | + | 455 |
| chr15 | 48261168 | 48266807 | L1MdTf_I | + | 5639 |
| chr15 | 48266807 | 48267794 | L1MdTf_I | + | 987 |
| chr15 | 48376974 | 48382095 | L1MdTf_III | + | 5121 |
| chr15 | 48382095 | 48383065 | L1MdTf_III | + | 970 |
| chr15 | 49992623 | 49998227 | L1MdTf_I | + | 5604 |
| chr15 | 49998227 | 49999220 | L1MdTf_I | + | 993 |
| chr15 | 50227858 | 50233000 | L1MdTf_I | + | 5142 |
| chr15 | 50233000 | 50234014 | L1MdTf_I | + | 1014 |
| chr15 | 53437549 | 53442987 | L1MdTf_I | + | 5438 |
| chr15 | 53442987 | 53443981 | L1MdTf_I | + | 994 |
| chr15 | 54208655 | 54208921 | L1MdTf_III | - | 266 |
| chr15 | 54208921 | 54209401 | L1MdTf_III | + | 480 |

| | | | | | |
|---|---|---|---|---|---|
| chr15 | 55022547 | 55023960 | L1MdTf_I | - | 1413 |
| chr15 | 55023960 | 55024627 | L1MdTf_I | + | 667 |
| chr15 | 55259511 | 55265485 | L1MdTf_III | + | 5974 |
| chr15 | 55265485 | 55266480 | L1MdTf_III | + | 995 |
| chr15 | 56387331 | 56389293 | L1MdTf_III | + | 1962 |
| chr15 | 56389293 | 56390286 | L1MdTf_III | + | 993 |
| chr15 | 60200670 | 60201681 | L1MdTf_I | + | 1011 |
| chr15 | 60201681 | 60202674 | L1MdTf_I | + | 993 |
| chr15 | 60269944 | 60275384 | L1MdTf_II | + | 5440 |
| chr15 | 60275384 | 60276406 | L1MdTf_II | + | 1022 |
| chr15 | 60310373 | 60314645 | L1MdTf_III | + | 4272 |
| chr15 | 60314645 | 60315636 | L1MdTf_III | + | 991 |
| chr15 | 60845236 | 60850912 | L1MdTf_III | + | 5676 |
| chr15 | 60850913 | 60851906 | L1MdTf_III | + | 993 |
| chr15 | 61534290 | 61535846 | L1MdTf_III | + | 1556 |
| chr15 | 61535847 | 61536871 | L1MdTf_III | + | 1024 |
| chr15 | 65144547 | 65150411 | L1MdTf_II | + | 5864 |
| chr15 | 65150411 | 65151404 | L1MdTf_II | + | 993 |
| chr15 | 65382648 | 65387838 | L1MdTf_III | + | 5190 |
| chr15 | 65387838 | 65388825 | L1MdTf_III | + | 987 |
| chr15 | 65444891 | 65448829 | L1MdTf_III | + | 3938 |
| chr15 | 65448829 | 65449842 | L1MdTf_III | + | 1013 |
| chr15 | 65478147 | 65478578 | L1MdTf_I | - | 431 |
| chr15 | 65478579 | 65479270 | L1MdTf_I | + | 691 |
| chr15 | 65609412 | 65615258 | L1MdTf_I | + | 5846 |
| chr15 | 65615258 | 65616251 | L1MdTf_I | + | 993 |
| chr15 | 65701983 | 65704195 | L1MdTf_III | + | 2212 |
| chr15 | 65704195 | 65705188 | L1MdTf_III | + | 993 |
| chr15 | 67113243 | 67114228 | L1MdTf_III | - | 985 |
| chr15 | 67114228 | 67114798 | L1MdTf_III | + | 570 |
| chr15 | 67130626 | 67131521 | L1MdTf_II | + | 895 |
| chr15 | 67131521 | 67132514 | L1MdTf_II | + | 993 |
| chr15 | 67255480 | 67256690 | L1MdTf_II | - | 1210 |
| chr15 | 67256690 | 67257586 | L1MdTf_II | + | 896 |
| chr15 | 67566854 | 67567349 | L1MdTf_II | - | 495 |
| chr15 | 67567349 | 67568363 | L1MdTf_II | + | 1014 |
| chr15 | 67789625 | 67789969 | L1MdTf_I | + | 344 |
| chr15 | 67789969 | 67790962 | L1MdTf_I | + | 993 |
| chr15 | 67904071 | 67909448 | L1MdTf_III | + | 5377 |
| chr15 | 67909449 | 67910440 | L1MdTf_III | + | 991 |
| chr15 | 68240030 | 68245484 | L1MdTf_III | + | 5454 |
| chr15 | 68245484 | 68246477 | L1MdTf_III | + | 993 |
| chr15 | 68757734 | 68763159 | L1MdTf_II | + | 5425 |
| chr15 | 68763159 | 68764152 | L1MdTf_II | + | 993 |
| chr15 | 70117638 | 70118949 | L1MdTf_I | + | 1311 |

| | | | | | |
|---|---|---|---|---|---|
| chr15 | 70118949 | 70119942 | L1MdTf_I | + | 993 |
| chr15 | 70376845 | 70377140 | L1MdTf_III | + | 295 |
| chr15 | 70377140 | 70378138 | L1MdTf_III | + | 998 |
| chr15 | 71521247 | 71527960 | L1MdTf_I | + | 6713 |
| chr15 | 71527960 | 71528976 | L1MdTf_I | + | 1016 |
| chr15 | 77433762 | 77439124 | L1MdTf_II | + | 5362 |
| chr15 | 77439124 | 77440113 | L1MdTf_II | + | 989 |
| chr15 | 90076879 | 90082539 | L1MdTf_II | + | 5660 |
| chr15 | 90082539 | 90083531 | L1MdTf_II | + | 992 |
| chr15 | 92711433 | 92717031 | L1MdTf_III | + | 5598 |
| chr15 | 92717031 | 92718024 | L1MdTf_III | + | 993 |
| chr15 | 93067618 | 93073085 | L1MdTf_II | + | 5467 |
| chr15 | 93073085 | 93074078 | L1MdTf_II | + | 993 |
| chr15 | 98253087 | 98255025 | L1MdTf_II | + | 1938 |
| chr15 | 98255025 | 98256018 | L1MdTf_II | + | 993 |
| chr15 | 101564088 | 101569559 | L1MdTf_I | + | 5471 |
| chr15 | 101569559 | 101570552 | L1MdTf_I | + | 993 |
| chr15 | 103319052 | 103325971 | L1MdTf_I | + | 6919 |
| chr15 | 103325971 | 103326964 | L1MdTf_I | + | 993 |
| chr16 | 3202141 | 3208576 | L1MdTf_II | + | 6435 |
| chr16 | 3208576 | 3209566 | L1MdTf_II | + | 990 |
| chr16 | 7114103 | 7114626 | L1MdTf_III | - | 523 |
| chr16 | 7114626 | 7116001 | L1MdTf_III | + | 1375 |
| chr16 | 7770153 | 7775365 | L1MdTf_III | + | 5212 |
| chr16 | 7775366 | 7776358 | L1MdTf_III | + | 992 |
| chr16 | 7967860 | 7973193 | L1MdTf_I | + | 5333 |
| chr16 | 7973193 | 7974209 | L1MdTf_I | + | 1016 |
| chr16 | 11677504 | 11683345 | L1MdTf_II | + | 5841 |
| chr16 | 11683345 | 11684341 | L1MdTf_II | + | 996 |
| chr16 | 12280260 | 12286313 | L1MdTf_II | + | 6053 |
| chr16 | 12286313 | 12287306 | L1MdTf_II | + | 993 |
| chr16 | 13140399 | 13140716 | L1MdTf_III | - | 317 |
| chr16 | 13140716 | 13141121 | L1MdTf_III | + | 405 |
| chr16 | 14491737 | 14493652 | L1MdTf_III | + | 1915 |
| chr16 | 14493653 | 14495753 | L1MdTf_II | + | 2100 |
| chr16 | 14504068 | 14507203 | L1MdTf_II | + | 3135 |
| chr16 | 14507203 | 14508196 | L1MdTf_II | + | 993 |
| chr16 | 15133256 | 15138905 | L1MdTf_III | + | 5649 |
| chr16 | 15138905 | 15139898 | L1MdTf_III | + | 993 |
| chr16 | 16324925 | 16331140 | L1MdTf_II | + | 6215 |
| chr16 | 16331141 | 16332133 | L1MdTf_II | + | 992 |
| chr16 | 16486071 | 16486385 | L1MdTf_III | + | 314 |
| chr16 | 16486385 | 16487378 | L1MdTf_III | + | 993 |
| chr16 | 19231819 | 19237448 | L1MdTf_I | + | 5629 |
| chr16 | 19237448 | 19238441 | L1MdTf_I | + | 993 |

| | | | | | |
|---|---|---|---|---|---|
| chr16 | 19552675 | 19558127 | L1MdTf_III | + | 5452 |
| chr16 | 19558127 | 19559109 | L1MdTf_III | + | 982 |
| chr16 | 25009629 | 25014871 | L1MdTf_III | + | 5242 |
| chr16 | 25014871 | 25015864 | L1MdTf_III | + | 993 |
| chr16 | 25384411 | 25390068 | L1MdTf_II | + | 5657 |
| chr16 | 25390068 | 25391061 | L1MdTf_II | + | 993 |
| chr16 | 26039554 | 26041436 | L1MdTf_II | + | 1882 |
| chr16 | 26041436 | 26042429 | L1MdTf_II | + | 993 |
| chr16 | 26136090 | 26141410 | L1MdTf_III | + | 5320 |
| chr16 | 26141410 | 26142403 | L1MdTf_III | + | 993 |
| chr16 | 26685197 | 26691320 | L1MdTf_I | + | 6123 |
| chr16 | 26691320 | 26692313 | L1MdTf_I | + | 993 |
| chr16 | 28009891 | 28015339 | L1MdTf_II | + | 5448 |
| chr16 | 28015339 | 28016332 | L1MdTf_II | + | 993 |
| chr16 | 28471875 | 28472999 | L1MdTf_II | - | 1124 |
| chr16 | 28472999 | 28473629 | L1MdTf_II | + | 630 |
| chr16 | 35262729 | 35268099 | L1MdTf_II | + | 5370 |
| chr16 | 35268099 | 35269116 | L1MdTf_II | + | 1017 |
| chr16 | 36938649 | 36941041 | L1MdTf_II | + | 2392 |
| chr16 | 36941041 | 36942034 | L1MdTf_II | + | 993 |
| chr16 | 39255652 | 39260896 | L1MdTf_III | + | 5244 |
| chr16 | 39260896 | 39261889 | L1MdTf_III | + | 993 |
| chr16 | 39356067 | 39357332 | L1MdTf_III | - | 1265 |
| chr16 | 39357332 | 39358092 | L1MdTf_III | + | 760 |
| chr16 | 40348956 | 40351368 | L1MdTf_II | + | 2412 |
| chr16 | 40351368 | 40352361 | L1MdTf_II | + | 993 |
| chr16 | 40634586 | 40639840 | L1MdTf_III | + | 5254 |
| chr16 | 40639840 | 40640820 | L1MdTf_III | + | 980 |
| chr16 | 41395412 | 41401911 | L1MdTf_I | + | 6499 |
| chr16 | 41401911 | 41402928 | L1MdTf_I | + | 1017 |
| chr16 | 42137608 | 42137995 | L1MdTf_III | + | 387 |
| chr16 | 42137995 | 42138988 | L1MdTf_III | + | 993 |
| chr16 | 42633738 | 42638999 | L1MdTf_III | + | 5261 |
| chr16 | 42638999 | 42639991 | L1MdTf_III | + | 992 |
| chr16 | 42906651 | 42908065 | L1MdTf_I | + | 1414 |
| chr16 | 42908065 | 42909057 | L1MdTf_I | + | 992 |
| chr16 | 44714378 | 44719624 | L1MdTf_III | + | 5246 |
| chr16 | 44719624 | 44720615 | L1MdTf_III | + | 991 |
| chr16 | 44743008 | 44744258 | L1MdTf_I | - | 1250 |
| chr16 | 44744258 | 44744754 | L1MdTf_I | + | 496 |
| chr16 | 46095712 | 46099931 | L1MdTf_I | + | 4219 |
| chr16 | 46099931 | 46100924 | L1MdTf_I | + | 993 |
| chr16 | 46462939 | 46469646 | L1MdTf_I | + | 6707 |
| chr16 | 46469646 | 46470639 | L1MdTf_I | + | 993 |
| chr16 | 46577593 | 46583039 | L1MdTf_III | + | 5446 |

| | | | | | |
|---|---|---|---|---|---|
| chr16 | 46583039 | 46584048 | L1MdTf_III | + | 1009 |
| chr16 | 47125321 | 47131433 | L1MdTf_II | + | 6112 |
| chr16 | 47131433 | 47132427 | L1MdTf_II | + | 994 |
| chr16 | 47196096 | 47201945 | L1MdTf_I | + | 5849 |
| chr16 | 47201945 | 47202966 | L1MdTf_I | + | 1021 |
| chr16 | 47841958 | 47847640 | L1MdTf_III | + | 5682 |
| chr16 | 47847640 | 47848633 | L1MdTf_III | + | 993 |
| chr16 | 47963741 | 47968928 | L1MdTf_III | + | 5187 |
| chr16 | 47968929 | 47969921 | L1MdTf_III | + | 992 |
| chr16 | 48676076 | 48681755 | L1MdTf_II | + | 5679 |
| chr16 | 48681755 | 48682749 | L1MdTf_II | + | 994 |
| chr16 | 48783198 | 48788828 | L1MdTf_I | + | 5630 |
| chr16 | 48788828 | 48789821 | L1MdTf_I | + | 993 |
| chr16 | 49077235 | 49079617 | L1MdTf_I | + | 2382 |
| chr16 | 49079617 | 49080632 | L1MdTf_I | + | 1015 |
| chr16 | 49286767 | 49292324 | L1MdTf_III | + | 5557 |
| chr16 | 49292324 | 49293317 | L1MdTf_III | + | 993 |
| chr16 | 50649154 | 50649217 | L1MdTf_III | + | 63 |
| chr16 | 50649217 | 50650210 | L1MdTf_III | + | 993 |
| chr16 | 50949706 | 50955642 | L1MdTf_I | + | 5936 |
| chr16 | 50955642 | 50956654 | L1MdTf_I | + | 1012 |
| chr16 | 51549901 | 51555586 | L1MdTf_III | + | 5685 |
| chr16 | 51555586 | 51556579 | L1MdTf_III | + | 993 |
| chr16 | 51757314 | 51757399 | L1MdTf_II | - | 85 |
| chr16 | 51757399 | 51757655 | L1MdTf_I | + | 256 |
| chr16 | 52358304 | 52358868 | L1MdTf_II | - | 564 |
| chr16 | 52358868 | 52359683 | L1MdTf_II | + | 815 |
| chr16 | 53645289 | 53651658 | L1MdTf_I | + | 6369 |
| chr16 | 53651658 | 53652651 | L1MdTf_I | + | 993 |
| chr16 | 54990085 | 54990657 | L1MdTf_II | - | 572 |
| chr16 | 54990657 | 54991596 | L1MdTf_II | + | 939 |
| chr16 | 58069789 | 58075058 | L1MdTf_I | + | 5269 |
| chr16 | 58075058 | 58076051 | L1MdTf_I | + | 993 |
| chr16 | 58139716 | 58145155 | L1MdTf_III | + | 5439 |
| chr16 | 58145155 | 58146142 | L1MdTf_III | + | 987 |
| chr16 | 58238042 | 58239265 | L1MdTf_II | - | 1223 |
| chr16 | 58239265 | 58239895 | L1MdTf_II | + | 630 |
| chr16 | 58682843 | 58688111 | L1MdTf_III | + | 5268 |
| chr16 | 58688111 | 58689134 | L1MdTf_III | + | 1023 |
| chr16 | 58797709 | 58803388 | L1MdTf_II | + | 5679 |
| chr16 | 58803388 | 58804381 | L1MdTf_II | + | 993 |
| chr16 | 59192869 | 59198196 | L1MdTf_III | + | 5327 |
| chr16 | 59198196 | 59199191 | L1MdTf_III | + | 995 |
| chr16 | 60256614 | 60256900 | L1MdTf_III | + | 286 |
| chr16 | 60256901 | 60262174 | L1MdTf_III | + | 5273 |

| | | | | | |
|---|---|---|---|---|---|
| chr16 | 60263169 | 60264282 | L1MdTf_III | + | 1113 |
| chr16 | 60264283 | 60265274 | L1MdTf_III | + | 991 |
| chr16 | 60265283 | 60266396 | L1MdTf_III | + | 1113 |
| chr16 | 60266397 | 60267388 | L1MdTf_III | + | 991 |
| chr16 | 60276702 | 60277192 | L1MdTf_I | + | 490 |
| chr16 | 60277193 | 60278185 | L1MdTf_I | + | 992 |
| chr16 | 61166655 | 61167297 | L1MdTf_I | - | 642 |
| chr16 | 61167297 | 61167723 | L1MdTf_I | + | 426 |
| chr16 | 61567982 | 61568150 | L1MdTf_I | - | 168 |
| chr16 | 61568150 | 61568985 | L1MdTf_I | + | 835 |
| chr16 | 61673594 | 61679249 | L1MdTf_I | + | 5655 |
| chr16 | 61679249 | 61680242 | L1MdTf_I | + | 993 |
| chr16 | 62165603 | 62171063 | L1MdTf_III | + | 5460 |
| chr16 | 62171063 | 62172056 | L1MdTf_III | + | 993 |
| chr16 | 63259538 | 63264968 | L1MdTf_II | + | 5430 |
| chr16 | 63264968 | 63265961 | L1MdTf_II | + | 993 |
| chr16 | 63451036 | 63451555 | L1MdTf_I | + | 519 |
| chr16 | 63451555 | 63452548 | L1MdTf_I | + | 993 |
| chr16 | 63788885 | 63794408 | L1MdTf_III | + | 5523 |
| chr16 | 63794408 | 63795403 | L1MdTf_III | + | 995 |
| chr16 | 64230790 | 64236047 | L1MdTf_III | + | 5257 |
| chr16 | 64236047 | 64237040 | L1MdTf_III | + | 993 |
| chr16 | 64270631 | 64276036 | L1MdTf_II | + | 5405 |
| chr16 | 64276036 | 64277033 | L1MdTf_II | + | 997 |
| chr16 | 65045777 | 65046112 | L1MdTf_II | + | 335 |
| chr16 | 65046112 | 65047136 | L1MdTf_II | + | 1024 |
| chr16 | 66026093 | 66031520 | L1MdTf_II | + | 5427 |
| chr16 | 66031520 | 66032517 | L1MdTf_II | + | 997 |
| chr16 | 66067261 | 66067882 | L1MdTf_III | + | 621 |
| chr16 | 66067882 | 66068876 | L1MdTf_III | + | 994 |
| chr16 | 67276731 | 67277291 | L1MdTf_II | - | 560 |
| chr16 | 67277291 | 67277496 | L1MdTf_II | + | 205 |
| chr16 | 67607391 | 67608226 | L1MdTf_II | - | 835 |
| chr16 | 67608226 | 67608788 | L1MdTf_II | + | 562 |
| chr16 | 68663139 | 68663418 | L1MdTf_II | - | 279 |
| chr16 | 68663418 | 68663788 | L1MdTf_II | + | 370 |
| chr16 | 69005561 | 69011442 | L1MdTf_II | + | 5881 |
| chr16 | 69011442 | 69012438 | L1MdTf_II | + | 996 |
| chr16 | 69496875 | 69500000 | L1MdTf_III | - | 3125 |
| chr16 | 69500000 | 69503283 | L1MdTf_III | - | 3283 |
| chr16 | 69689810 | 69695491 | L1MdTf_II | + | 5681 |
| chr16 | 69695491 | 69696484 | L1MdTf_II | + | 993 |
| chr16 | 71073319 | 71079048 | L1MdTf_I | + | 5729 |
| chr16 | 71079048 | 71080064 | L1MdTf_I | + | 1016 |
| chr16 | 71154976 | 71156130 | L1MdTf_III | + | 1154 |

| | | | | | |
|---|---|---|---|---|---|
| chr16 | 71156130 | 71157123 | L1MdTf_III | + | 993 |
| chr16 | 71925916 | 71926242 | L1MdTf_I | - | 326 |
| chr16 | 71926242 | 71926905 | L1MdTf_I | + | 663 |
| chr16 | 72393400 | 72398658 | L1MdTf_III | + | 5258 |
| chr16 | 72398658 | 72399649 | L1MdTf_III | + | 991 |
| chr16 | 72420551 | 72426412 | L1MdTf_II | + | 5861 |
| chr16 | 72426412 | 72427405 | L1MdTf_II | + | 993 |
| chr16 | 73151264 | 73156584 | L1MdTf_III | + | 5320 |
| chr16 | 73156584 | 73157577 | L1MdTf_III | + | 993 |
| chr16 | 73453604 | 73459131 | L1MdTf_III | + | 5527 |
| chr16 | 73459131 | 73460123 | L1MdTf_III | + | 992 |
| chr16 | 73478234 | 73484085 | L1MdTf_II | + | 5851 |
| chr16 | 73484085 | 73485101 | L1MdTf_II | + | 1016 |
| chr16 | 74002645 | 74008165 | L1MdTf_II | + | 5520 |
| chr16 | 74008165 | 74009158 | L1MdTf_II | + | 993 |
| chr16 | 74768429 | 74773675 | L1MdTf_II | + | 5246 |
| chr16 | 74773675 | 74774668 | L1MdTf_II | + | 993 |
| chr16 | 75749274 | 75754449 | L1MdTf_III | + | 5175 |
| chr16 | 75754449 | 75755442 | L1MdTf_III | + | 993 |
| chr16 | 76585927 | 76591213 | L1MdTf_II | + | 5286 |
| chr16 | 76591213 | 76592206 | L1MdTf_II | + | 993 |
| chr16 | 76952148 | 76957591 | L1MdTf_I | + | 5443 |
| chr16 | 76957591 | 76958605 | L1MdTf_I | + | 1014 |
| chr16 | 78147084 | 78153178 | L1MdTf_III | + | 6094 |
| chr16 | 78153178 | 78154173 | L1MdTf_III | + | 995 |
| chr16 | 78785011 | 78790236 | L1MdTf_II | + | 5225 |
| chr16 | 78790236 | 78791228 | L1MdTf_II | + | 992 |
| chr16 | 79480823 | 79481123 | L1MdTf_III | - | 300 |
| chr16 | 79481123 | 79481638 | L1MdTf_III | + | 515 |
| chr16 | 79578585 | 79583834 | L1MdTf_III | + | 5249 |
| chr16 | 79583834 | 79584827 | L1MdTf_III | + | 993 |
| chr16 | 82162795 | 82163190 | L1MdTf_III | - | 395 |
| chr16 | 82163190 | 82163312 | L1MdTf_III | + | 122 |
| chr16 | 82304512 | 82310047 | L1MdTf_II | + | 5535 |
| chr16 | 82310047 | 82311039 | L1MdTf_II | + | 992 |
| chr16 | 82340233 | 82341995 | L1MdTf_III | + | 1762 |
| chr16 | 82341995 | 82342987 | L1MdTf_III | + | 992 |
| chr16 | 83058871 | 83064416 | L1MdTf_I | + | 5545 |
| chr16 | 83064416 | 83065409 | L1MdTf_I | + | 993 |
| chr16 | 84078435 | 84079299 | L1MdTf_III | + | 864 |
| chr16 | 84079299 | 84080291 | L1MdTf_III | + | 992 |
| chr16 | 84452268 | 84452377 | L1MdTf_II | + | 109 |
| chr16 | 84452377 | 84453370 | L1MdTf_II | + | 993 |
| chr16 | 85037476 | 85042694 | L1MdTf_III | + | 5218 |
| chr16 | 85042694 | 85043684 | L1MdTf_III | + | 990 |

| | | | | | |
|---|---|---|---|---|---|
| chr16 | 85927390 | 85928347 | L1MdTf_II | + | 957 |
| chr16 | 85928347 | 85929340 | L1MdTf_II | + | 993 |
| chr16 | 86148621 | 86150212 | L1MdTf_I | + | 1591 |
| chr16 | 86150212 | 86151205 | L1MdTf_I | + | 993 |
| chr16 | 86331642 | 86332115 | L1MdTf_III | - | 473 |
| chr16 | 86332115 | 86332539 | L1MdTf_III | + | 424 |
| chr16 | 86522160 | 86527544 | L1MdTf_III | + | 5384 |
| chr16 | 86527545 | 86528537 | L1MdTf_III | + | 992 |
| chr16 | 86747736 | 86753953 | L1MdTf_III | + | 6217 |
| chr16 | 86753953 | 86754948 | L1MdTf_III | + | 995 |
| chr16 | 86909008 | 86914448 | L1MdTf_I | + | 5440 |
| chr16 | 86914448 | 86915441 | L1MdTf_I | + | 993 |
| chr16 | 86946134 | 86951934 | L1MdTf_II | + | 5800 |
| chr16 | 86951934 | 86952927 | L1MdTf_II | + | 993 |
| chr16 | 86990109 | 86995563 | L1MdTf_III | + | 5454 |
| chr16 | 86995563 | 86996584 | L1MdTf_III | + | 1021 |
| chr16 | 88324309 | 88329932 | L1MdTf_I | + | 5623 |
| chr16 | 88329932 | 88330953 | L1MdTf_I | + | 1021 |
| chr16 | 89432874 | 89438894 | L1MdTf_III | + | 6020 |
| chr16 | 89438894 | 89439913 | L1MdTf_III | + | 1019 |
| chr16 | 95583696 | 95589123 | L1MdTf_II | + | 5427 |
| chr16 | 95589123 | 95590139 | L1MdTf_II | + | 1016 |
| chr17 | 12091123 | 12091565 | L1MdTf_II | - | 442 |
| chr17 | 12091565 | 12092148 | L1MdTf_II | + | 583 |
| chr17 | 14253547 | 14254544 | L1MdTf_I | - | 997 |
| chr17 | 14254544 | 14255459 | L1MdTf_I | + | 915 |
| chr17 | 16357275 | 16362728 | L1MdTf_III | + | 5453 |
| chr17 | 16362728 | 16363721 | L1MdTf_III | + | 993 |
| chr17 | 17774373 | 17780517 | L1MdTf_II | + | 6144 |
| chr17 | 17780517 | 17781510 | L1MdTf_II | + | 993 |
| chr17 | 19070128 | 19072608 | L1MdTf_III | + | 2480 |
| chr17 | 19072608 | 19073601 | L1MdTf_III | + | 993 |
| chr17 | 19324052 | 19329896 | L1MdTf_I | + | 5844 |
| chr17 | 19329896 | 19330889 | L1MdTf_I | + | 993 |
| chr17 | 19817514 | 19819599 | L1MdTf_I | - | 2085 |
| chr17 | 19819599 | 19820224 | L1MdTf_I | + | 625 |
| chr17 | 20859279 | 20864889 | L1MdTf_II | + | 5610 |
| chr17 | 20864889 | 20865882 | L1MdTf_II | + | 993 |
| chr17 | 21424412 | 21429769 | L1MdTf_III | + | 5357 |
| chr17 | 21429769 | 21430762 | L1MdTf_III | + | 993 |
| chr17 | 22687884 | 22693470 | L1MdTf_III | + | 5586 |
| chr17 | 22693471 | 22694464 | L1MdTf_III | + | 993 |
| chr17 | 22867431 | 22872670 | L1MdTf_III | + | 5239 |
| chr17 | 22872670 | 22873690 | L1MdTf_III | + | 1020 |
| chr17 | 22977301 | 22978221 | L1MdTf_I | - | 920 |

| | | | | | |
|---|---|---|---|---|---|
| chr17 | 22978221 | 22979028 | L1MdTf_I | + | 807 |
| chr17 | 33497613 | 33500000 | L1MdTf_II | + | 2387 |
| chr17 | 33500000 | 33503479 | L1MdTf_II | + | 3479 |
| chr17 | 33633310 | 33636508 | L1MdTf_III | + | 3198 |
| chr17 | 33636508 | 33636966 | L1MdTf_III | - | 458 |
| chr17 | 33636966 | 33639487 | L1MdTf_III | + | 2521 |
| chr17 | 33639487 | 33640480 | L1MdTf_III | + | 993 |
| chr17 | 36902401 | 36903187 | L1MdTf_III | + | 786 |
| chr17 | 36903188 | 36904185 | L1MdTf_III | + | 997 |
| chr17 | 37018216 | 37024097 | L1MdTf_II | + | 5881 |
| chr17 | 37024097 | 37025090 | L1MdTf_II | + | 993 |
| chr17 | 37025126 | 37027013 | L1MdTf_II | + | 1887 |
| chr17 | 37027013 | 37028006 | L1MdTf_II | + | 993 |
| chr17 | 38122034 | 38127564 | L1MdTf_II | + | 5530 |
| chr17 | 38127564 | 38128560 | L1MdTf_II | + | 996 |
| chr17 | 39201612 | 39207292 | L1MdTf_II | + | 5680 |
| chr17 | 39207292 | 39208290 | L1MdTf_II | + | 998 |
| chr17 | 39208549 | 39213788 | L1MdTf_II | + | 5239 |
| chr17 | 39213788 | 39214785 | L1MdTf_II | + | 997 |
| chr17 | 39939276 | 39939754 | L1MdTf_II | - | 478 |
| chr17 | 39939754 | 39940697 | L1MdTf_II | + | 943 |
| chr17 | 40303275 | 40308925 | L1MdTf_II | + | 5650 |
| chr17 | 40308925 | 40309918 | L1MdTf_II | + | 993 |
| chr17 | 41065668 | 41070886 | L1MdTf_I | + | 5218 |
| chr17 | 41070886 | 41071897 | L1MdTf_I | + | 1011 |
| chr17 | 41110666 | 41112061 | L1MdTf_II | + | 1395 |
| chr17 | 41112061 | 41113054 | L1MdTf_II | + | 993 |
| chr17 | 41276240 | 41281879 | L1MdTf_I | + | 5639 |
| chr17 | 41281879 | 41282872 | L1MdTf_I | + | 993 |
| chr17 | 51233859 | 51239226 | L1MdTf_II | + | 5367 |
| chr17 | 51239226 | 51240219 | L1MdTf_II | + | 993 |
| chr17 | 53252193 | 53255769 | L1MdTf_II | + | 3576 |
| chr17 | 53255769 | 53256762 | L1MdTf_II | + | 993 |
| chr17 | 53423958 | 53429640 | L1MdTf_III | + | 5682 |
| chr17 | 53429640 | 53430631 | L1MdTf_III | + | 991 |
| chr17 | 53490095 | 53495454 | L1MdTf_III | + | 5359 |
| chr17 | 53495455 | 53496447 | L1MdTf_III | + | 992 |
| chr17 | 55882143 | 55887263 | L1MdTf_III | + | 5120 |
| chr17 | 55887263 | 55888256 | L1MdTf_III | + | 993 |
| chr17 | 57683140 | 57684120 | L1MdTf_II | - | 980 |
| chr17 | 57684121 | 57684796 | L1MdTf_II | - | 675 |
| chr17 | 58006044 | 58011701 | L1MdTf_II | + | 5657 |
| chr17 | 58011701 | 58012726 | L1MdTf_II | + | 1025 |
| chr17 | 58478097 | 58483564 | L1MdTf_I | + | 5467 |
| chr17 | 58483564 | 58484557 | L1MdTf_I | + | 993 |

| | | | | | |
|---|---|---|---|---|---|
| chr17 | 58858469 | 58863896 | L1MdTf_II | + | 5427 |
| chr17 | 58863896 | 58864889 | L1MdTf_II | + | 993 |
| chr17 | 59598746 | 59601521 | L1MdTf_II | + | 2775 |
| chr17 | 59601521 | 59602514 | L1MdTf_II | + | 993 |
| chr17 | 60784554 | 60784619 | L1MdTf_I | + | 65 |
| chr17 | 60784619 | 60785622 | L1MdTf_I | + | 1003 |
| chr17 | 61090287 | 61096331 | L1MdTf_II | + | 6044 |
| chr17 | 61096331 | 61097324 | L1MdTf_II | + | 993 |
| chr17 | 61689190 | 61689835 | L1MdTf_III | + | 645 |
| chr17 | 61689835 | 61690828 | L1MdTf_III | + | 993 |
| chr17 | 63918757 | 63924200 | L1MdTf_III | + | 5443 |
| chr17 | 63924200 | 63925183 | L1MdTf_III | + | 983 |
| chr17 | 65233737 | 65233966 | L1MdTf_III | + | 229 |
| chr17 | 65233966 | 65234959 | L1MdTf_III | + | 993 |
| chr17 | 65638093 | 65643733 | L1MdTf_II | + | 5640 |
| chr17 | 65643733 | 65644726 | L1MdTf_II | + | 993 |
| chr17 | 66998059 | 66998328 | L1MdTf_II | + | 269 |
| chr17 | 66998328 | 66999321 | L1MdTf_II | + | 993 |
| chr17 | 69194413 | 69196073 | L1MdTf_III | - | 1660 |
| chr17 | 69196073 | 69196744 | L1MdTf_III | + | 671 |
| chr17 | 70205072 | 70209283 | L1MdTf_III | - | 4211 |
| chr17 | 70209283 | 70209656 | L1MdTf_III | - | 373 |
| chr17 | 70456962 | 70462823 | L1MdTf_II | + | 5861 |
| chr17 | 70462823 | 70463816 | L1MdTf_II | + | 993 |
| chr17 | 70536317 | 70542636 | L1MdTf_I | + | 6319 |
| chr17 | 70542636 | 70543629 | L1MdTf_I | + | 993 |
| chr17 | 70630652 | 70636071 | L1MdTf_II | + | 5419 |
| chr17 | 70636071 | 70637064 | L1MdTf_II | + | 993 |
| chr17 | 74416963 | 74422369 | L1MdTf_III | + | 5406 |
| chr17 | 74422369 | 74423362 | L1MdTf_III | + | 993 |
| chr17 | 75706251 | 75711618 | L1MdTf_II | + | 5367 |
| chr17 | 75711618 | 75712611 | L1MdTf_II | + | 993 |
| chr17 | 75731134 | 75736824 | L1MdTf_II | + | 5690 |
| chr17 | 75736824 | 75737844 | L1MdTf_II | + | 1020 |
| chr17 | 77539415 | 77545071 | L1MdTf_II | + | 5656 |
| chr17 | 77545071 | 77546092 | L1MdTf_II | + | 1021 |
| chr17 | 78136479 | 78142022 | L1MdTf_I | + | 5543 |
| chr17 | 78142022 | 78143015 | L1MdTf_I | + | 993 |
| chr17 | 78441750 | 78441848 | L1MdTf_II | - | 98 |
| chr17 | 78441848 | 78441909 | L1MdTf_II | + | 61 |
| chr17 | 81991276 | 81996915 | L1MdTf_I | + | 5639 |
| chr17 | 81996915 | 81997908 | L1MdTf_I | + | 993 |
| chr17 | 81999842 | 82000000 | L1MdTf_III | + | 158 |
| chr17 | 82000000 | 82000785 | L1MdTf_III | + | 785 |
| chr17 | 82023845 | 82029246 | L1MdTf_I | + | 5401 |

| | | | | | |
|---|---|---|---|---|---|
| chr17 | 82029246 | 82030239 | L1MdTf_I | + | 993 |
| chr17 | 82118049 | 82118772 | L1MdTf_I | + | 723 |
| chr17 | 82118772 | 82119765 | L1MdTf_I | + | 993 |
| chr17 | 82196366 | 82196483 | L1MdTf_II | - | 117 |
| chr17 | 82196483 | 82196514 | L1MdTf_II | + | 31 |
| chr17 | 82587251 | 82587501 | L1MdTf_III | + | 250 |
| chr17 | 82587501 | 82588487 | L1MdTf_III | + | 986 |
| chr17 | 82771531 | 82777183 | L1MdTf_I | + | 5652 |
| chr17 | 82777183 | 82778176 | L1MdTf_I | + | 993 |
| chr17 | 82789079 | 82790420 | L1MdTf_I | + | 1341 |
| chr17 | 82790420 | 82791413 | L1MdTf_I | + | 993 |
| chr17 | 83296289 | 83297563 | L1MdTf_I | - | 1274 |
| chr17 | 83297563 | 83299026 | L1MdTf_I | + | 1463 |
| chr17 | 85571480 | 85572173 | L1MdTf_II | + | 693 |
| chr17 | 85572173 | 85573166 | L1MdTf_II | + | 993 |
| chr17 | 89748008 | 89753415 | L1MdTf_II | + | 5407 |
| chr17 | 89753415 | 89754408 | L1MdTf_II | + | 993 |
| chr17 | 90331383 | 90335959 | L1MdTf_I | + | 4576 |
| chr17 | 90335959 | 90336952 | L1MdTf_I | + | 993 |
| chr17 | 94843329 | 94848866 | L1MdTf_III | + | 5537 |
| chr17 | 94848866 | 94849859 | L1MdTf_III | + | 993 |
| chr17 | 95004062 | 95004658 | L1MdTf_II | + | 596 |
| chr17 | 95004658 | 95005651 | L1MdTf_II | + | 993 |
| chr18 | 3031031 | 3036712 | L1MdTf_II | + | 5681 |
| chr18 | 3036712 | 3037705 | L1MdTf_II | + | 993 |
| chr18 | 4021882 | 4028364 | L1MdTf_II | + | 6482 |
| chr18 | 4028364 | 4029393 | L1MdTf_II | + | 1029 |
| chr18 | 5751051 | 5757356 | L1MdTf_II | + | 6305 |
| chr18 | 5757356 | 5758346 | L1MdTf_II | + | 990 |
| chr18 | 7494796 | 7495664 | L1MdTf_I | - | 868 |
| chr18 | 7495664 | 7497869 | L1MdTf_I | + | 2205 |
| chr18 | 9202462 | 9207713 | L1MdTf_III | + | 5251 |
| chr18 | 9207713 | 9208706 | L1MdTf_III | + | 993 |
| chr18 | 11185874 | 11191102 | L1MdTf_I | + | 5228 |
| chr18 | 11191102 | 11192095 | L1MdTf_I | + | 993 |
| chr18 | 14865883 | 14871132 | L1MdTf_II | + | 5249 |
| chr18 | 14871132 | 14872125 | L1MdTf_II | + | 993 |
| chr18 | 16114078 | 16120343 | L1MdTf_I | + | 6265 |
| chr18 | 16120344 | 16121337 | L1MdTf_I | + | 993 |
| chr18 | 16854039 | 16854856 | L1MdTf_I | + | 817 |
| chr18 | 16854856 | 16855849 | L1MdTf_I | + | 993 |
| chr18 | 17094964 | 17100398 | L1MdTf_II | + | 5434 |
| chr18 | 17100398 | 17101415 | L1MdTf_II | + | 1017 |
| chr18 | 17351180 | 17356401 | L1MdTf_III | + | 5221 |
| chr18 | 17356401 | 17357394 | L1MdTf_III | + | 993 |

| | | | | | |
|---|---|---|---|---|---|
| chr18 | 17558154 | 17563336 | L1MdTf_III | + | 5182 |
| chr18 | 17563336 | 17563406 | L1MdTf_III | + | 70 |
| chr18 | 17587201 | 17592497 | L1MdTf_III | + | 5296 |
| chr18 | 17592497 | 17593490 | L1MdTf_III | + | 993 |
| chr18 | 18797550 | 18802995 | L1MdTf_I | + | 5445 |
| chr18 | 18802995 | 18804011 | L1MdTf_I | + | 1016 |
| chr18 | 19687057 | 19692303 | L1MdTf_III | + | 5246 |
| chr18 | 19692303 | 19693296 | L1MdTf_III | + | 993 |
| chr18 | 19995128 | 20000000 | L1MdTf_II | + | 4872 |
| chr18 | 20000000 | 20000781 | L1MdTf_I | + | 781 |
| chr18 | 21669406 | 21674620 | L1MdTf_II | + | 5214 |
| chr18 | 21674620 | 21675617 | L1MdTf_II | + | 997 |
| chr18 | 22246948 | 22247605 | L1MdTf_III | - | 657 |
| chr18 | 22247605 | 22249058 | L1MdTf_III | + | 1453 |
| chr18 | 22430426 | 22436150 | L1MdTf_I | + | 5724 |
| chr18 | 22436150 | 22437143 | L1MdTf_I | + | 993 |
| chr18 | 23010663 | 23016302 | L1MdTf_I | + | 5639 |
| chr18 | 23016302 | 23017283 | L1MdTf_I | + | 981 |
| chr18 | 23122052 | 23128539 | L1MdTf_I | + | 6487 |
| chr18 | 23128539 | 23129532 | L1MdTf_I | + | 993 |
| chr18 | 25913944 | 25919405 | L1MdTf_I | + | 5461 |
| chr18 | 25919405 | 25920398 | L1MdTf_I | + | 993 |
| chr18 | 26357033 | 26357354 | L1MdTf_II | + | 321 |
| chr18 | 26357354 | 26358347 | L1MdTf_II | + | 993 |
| chr18 | 26693479 | 26694150 | L1MdTf_II | - | 671 |
| chr18 | 26694150 | 26695582 | L1MdTf_II | + | 1432 |
| chr18 | 26829515 | 26834981 | L1MdTf_III | + | 5466 |
| chr18 | 26834981 | 26835974 | L1MdTf_III | + | 993 |
| chr18 | 27366784 | 27372884 | L1MdTf_III | + | 6100 |
| chr18 | 27372884 | 27373906 | L1MdTf_III | + | 1022 |
| chr18 | 28284276 | 28285797 | L1MdTf_II | - | 1521 |
| chr18 | 28285797 | 28285921 | L1MdTf_II | + | 124 |
| chr18 | 29331998 | 29333759 | L1MdTf_I | + | 1761 |
| chr18 | 29333759 | 29334752 | L1MdTf_I | + | 993 |
| chr18 | 32173433 | 32173528 | L1MdTf_II | - | 95 |
| chr18 | 32173528 | 32174108 | L1MdTf_II | + | 580 |
| chr18 | 32815830 | 32821267 | L1MdTf_III | + | 5437 |
| chr18 | 32821267 | 32822258 | L1MdTf_III | + | 991 |
| chr18 | 33177132 | 33182585 | L1MdTf_I | + | 5453 |
| chr18 | 33182585 | 33183577 | L1MdTf_I | + | 992 |
| chr18 | 33527977 | 33533630 | L1MdTf_II | + | 5653 |
| chr18 | 33533630 | 33534643 | L1MdTf_II | + | 1013 |
| chr18 | 37763267 | 37769115 | L1MdTf_II | + | 5848 |
| chr18 | 37769115 | 37770140 | L1MdTf_II | + | 1025 |
| chr18 | 40793971 | 40796080 | L1MdTf_III | - | 2109 |

| | | | | | |
|---|---|---|---|---|---|
| chr18 | 40796080 | 40796575 | L1MdTf_III | + | 495 |
| chr18 | 40895611 | 40898274 | L1MdTf_III | + | 2663 |
| chr18 | 40898274 | 40899266 | L1MdTf_III | + | 992 |
| chr18 | 41684223 | 41689624 | L1MdTf_I | + | 5401 |
| chr18 | 41689624 | 41690617 | L1MdTf_I | + | 993 |
| chr18 | 42943629 | 42944101 | L1MdTf_I | + | 472 |
| chr18 | 42944101 | 42945116 | L1MdTf_I | + | 1015 |
| chr18 | 44375186 | 44380421 | L1MdTf_III | + | 5235 |
| chr18 | 44380421 | 44381414 | L1MdTf_III | + | 993 |
| chr18 | 45996092 | 46000000 | L1MdTf_III | + | 3908 |
| chr18 | 46000000 | 46002083 | L1MdTf_III | + | 2083 |
| chr18 | 48165237 | 48165897 | L1MdTf_II | + | 660 |
| chr18 | 48165897 | 48166909 | L1MdTf_II | + | 1012 |
| chr18 | 48891430 | 48896682 | L1MdTf_III | + | 5252 |
| chr18 | 48896682 | 48897676 | L1MdTf_III | + | 994 |
| chr18 | 50915943 | 50921605 | L1MdTf_II | + | 5662 |
| chr18 | 50921605 | 50922602 | L1MdTf_II | + | 997 |
| chr18 | 52676349 | 52681714 | L1MdTf_III | + | 5365 |
| chr18 | 52681714 | 52682707 | L1MdTf_III | + | 993 |
| chr18 | 53009356 | 53015123 | L1MdTf_II | + | 5767 |
| chr18 | 53015123 | 53016141 | L1MdTf_II | + | 1018 |
| chr18 | 53410450 | 53411680 | L1MdTf_III | + | 1230 |
| chr18 | 53411680 | 53412705 | L1MdTf_III | + | 1025 |
| chr18 | 58320459 | 58320498 | L1MdTf_II | - | 39 |
| chr18 | 58320498 | 58320542 | L1MdTf_II | + | 44 |
| chr18 | 58940524 | 58942103 | L1MdTf_III | + | 1579 |
| chr18 | 58942103 | 58943114 | L1MdTf_III | + | 1011 |
| chr18 | 59248441 | 59253996 | L1MdTf_II | + | 5555 |
| chr18 | 59253996 | 59254989 | L1MdTf_II | + | 993 |
| chr18 | 59863054 | 59869753 | L1MdTf_I | + | 6699 |
| chr18 | 59869753 | 59870746 | L1MdTf_I | + | 993 |
| chr18 | 63167354 | 63172563 | L1MdTf_II | + | 5209 |
| chr18 | 63172563 | 63173556 | L1MdTf_II | + | 993 |
| chr18 | 71272113 | 71277438 | L1MdTf_III | + | 5325 |
| chr18 | 71277439 | 71278432 | L1MdTf_III | + | 993 |
| chr18 | 71378383 | 71378562 | L1MdTf_III | + | 179 |
| chr18 | 71378562 | 71379555 | L1MdTf_III | + | 993 |
| chr18 | 71763998 | 71769659 | L1MdTf_II | + | 5661 |
| chr18 | 71769659 | 71770677 | L1MdTf_II | + | 1018 |
| chr18 | 72639496 | 72645151 | L1MdTf_I | + | 5655 |
| chr18 | 72645151 | 72646144 | L1MdTf_I | + | 993 |
| chr18 | 72709896 | 72715726 | L1MdTf_I | + | 5830 |
| chr18 | 72715726 | 72716719 | L1MdTf_I | + | 993 |
| chr18 | 73243551 | 73248769 | L1MdTf_II | + | 5218 |
| chr18 | 73248769 | 73249762 | L1MdTf_II | + | 993 |

| | | | | | |
|---|---|---|---|---|---|
| chr18 | 73364075 | 73369271 | L1MdTf_III | + | 5196 |
| chr18 | 73369271 | 73370264 | L1MdTf_III | + | 993 |
| chr18 | 76874161 | 76879333 | L1MdTf_III | + | 5172 |
| chr18 | 76879333 | 76880324 | L1MdTf_III | + | 991 |
| chr18 | 78456618 | 78461859 | L1MdTf_III | + | 5241 |
| chr18 | 78461859 | 78462852 | L1MdTf_III | + | 993 |
| chr18 | 79555333 | 79556457 | L1MdTf_I | + | 1124 |
| chr18 | 79556457 | 79557450 | L1MdTf_I | + | 993 |
| chr18 | 83278103 | 83284542 | L1MdTf_II | - | 6439 |
| chr18 | 83284542 | 83289442 | L1MdTf_II | - | 4900 |
| chr18 | 83517248 | 83517327 | L1MdTf_I | - | 79 |
| chr18 | 83517327 | 83517974 | L1MdTf_I | + | 647 |
| chr18 | 85801711 | 85807174 | L1MdTf_I | + | 5463 |
| chr18 | 85807174 | 85808167 | L1MdTf_I | + | 993 |
| chr18 | 86578684 | 86584538 | L1MdTf_II | + | 5854 |
| chr18 | 86584538 | 86585534 | L1MdTf_II | + | 996 |
| chr18 | 86917808 | 86919298 | L1MdTf_II | + | 1490 |
| chr18 | 86919298 | 86920291 | L1MdTf_II | + | 993 |
| chr18 | 87550108 | 87550893 | L1MdTf_II | - | 785 |
| chr18 | 87550893 | 87551717 | L1MdTf_II | + | 824 |
| chr18 | 88411784 | 88413809 | L1MdTf_I | + | 2025 |
| chr18 | 88413809 | 88414802 | L1MdTf_I | + | 993 |
| chr18 | 89183710 | 89184376 | L1MdTf_II | + | 666 |
| chr18 | 89184376 | 89185369 | L1MdTf_II | + | 993 |
| chr18 | 90391857 | 90397706 | L1MdTf_I | + | 5849 |
| chr18 | 90397706 | 90398699 | L1MdTf_I | + | 993 |
| chr19 | 3223193 | 3227013 | L1MdTf_III | + | 3820 |
| chr19 | 3227014 | 3228006 | L1MdTf_III | + | 992 |
| chr19 | 11064675 | 11065814 | L1MdTf_I | + | 1139 |
| chr19 | 11065814 | 11066832 | L1MdTf_I | + | 1018 |
| chr19 | 12480460 | 12485641 | L1MdTf_I | + | 5181 |
| chr19 | 12485641 | 12486634 | L1MdTf_I | + | 993 |
| chr19 | 13747903 | 13753542 | L1MdTf_I | + | 5639 |
| chr19 | 13753542 | 13754535 | L1MdTf_I | + | 993 |
| chr19 | 13876836 | 13882684 | L1MdTf_II | + | 5848 |
| chr19 | 13882684 | 13883677 | L1MdTf_II | + | 993 |
| chr19 | 15291903 | 15293061 | L1MdTf_III | + | 1158 |
| chr19 | 15293061 | 15294085 | L1MdTf_III | + | 1024 |
| chr19 | 20552527 | 20552656 | L1MdTf_II | + | 129 |
| chr19 | 20552656 | 20553649 | L1MdTf_II | + | 993 |
| chr19 | 21195545 | 21201174 | L1MdTf_I | + | 5629 |
| chr19 | 21201174 | 21202167 | L1MdTf_I | + | 993 |
| chr19 | 21425915 | 21426493 | L1MdTf_III | - | 578 |
| chr19 | 21426493 | 21427062 | L1MdTf_III | + | 569 |
| chr19 | 27728773 | 27734444 | L1MdTf_II | + | 5671 |

| | | | | | |
|---|---|---|---|---|---|
| chr19 | 27734444 | 27735437 | L1MdTf_II | + | 993 |
| chr19 | 27948438 | 27949362 | L1MdTf_III | + | 924 |
| chr19 | 27949362 | 27949572 | L1MdTf_III | + | 210 |
| chr19 | 28456908 | 28457008 | L1MdTf_II | + | 100 |
| chr19 | 28457009 | 28458001 | L1MdTf_II | + | 992 |
| chr19 | 30605871 | 30606497 | L1MdTf_II | + | 626 |
| chr19 | 30606497 | 30607490 | L1MdTf_II | + | 993 |
| chr19 | 31700163 | 31706438 | L1MdTf_I | + | 6275 |
| chr19 | 31706438 | 31707431 | L1MdTf_I | + | 993 |
| chr19 | 31712019 | 31712221 | L1MdTf_II | + | 202 |
| chr19 | 31712221 | 31713214 | L1MdTf_II | + | 993 |
| chr19 | 31914548 | 31920200 | L1MdTf_I | + | 5652 |
| chr19 | 31920200 | 31921212 | L1MdTf_I | + | 1012 |
| chr19 | 33263685 | 33269148 | L1MdTf_II | + | 5463 |
| chr19 | 33269148 | 33270143 | L1MdTf_II | + | 995 |
| chr19 | 33572250 | 33577920 | L1MdTf_I | + | 5670 |
| chr19 | 33577920 | 33578941 | L1MdTf_I | + | 1021 |
| chr19 | 33739808 | 33745238 | L1MdTf_II | + | 5430 |
| chr19 | 33745238 | 33746231 | L1MdTf_II | + | 993 |
| chr19 | 33931741 | 33934738 | L1MdTf_III | + | 2997 |
| chr19 | 33934738 | 33935735 | L1MdTf_III | + | 997 |
| chr19 | 34002803 | 34008057 | L1MdTf_III | + | 5254 |
| chr19 | 34008057 | 34009050 | L1MdTf_III | + | 993 |
| chr19 | 35211988 | 35217370 | L1MdTf_III | + | 5382 |
| chr19 | 35217370 | 35218363 | L1MdTf_III | + | 993 |
| chr19 | 35881401 | 35887616 | L1MdTf_II | + | 6215 |
| chr19 | 35887616 | 35888609 | L1MdTf_II | + | 993 |
| chr19 | 40135340 | 40140768 | L1MdTf_III | + | 5428 |
| chr19 | 40140768 | 40141761 | L1MdTf_III | + | 993 |
| chr19 | 47793279 | 47798620 | L1MdTf_III | + | 5341 |
| chr19 | 47798620 | 47799065 | L1MdTf_III | + | 445 |
| chr19 | 48920796 | 48926439 | L1MdTf_I | + | 5643 |
| chr19 | 48926439 | 48927432 | L1MdTf_I | + | 993 |
| chr19 | 49328740 | 49334928 | L1MdTf_I | + | 6188 |
| chr19 | 49334928 | 49335922 | L1MdTf_I | + | 994 |
| chr19 | 49511726 | 49517172 | L1MdTf_II | + | 5446 |
| chr19 | 49517172 | 49518190 | L1MdTf_II | + | 1018 |
| chr19 | 50414709 | 50420560 | L1MdTf_II | + | 5851 |
| chr19 | 50420560 | 50421553 | L1MdTf_II | + | 993 |
| chr19 | 51888141 | 51893770 | L1MdTf_I | + | 5629 |
| chr19 | 51893770 | 51894763 | L1MdTf_I | + | 993 |
| chr19 | 52829147 | 52834533 | L1MdTf_III | + | 5386 |
| chr19 | 52834534 | 52835527 | L1MdTf_III | + | 993 |
| chr19 | 54565413 | 54566184 | L1MdTf_II | + | 771 |
| chr19 | 54566184 | 54567177 | L1MdTf_II | + | 993 |

| | | | | | |
|---|---|---|---|---|---|
| chr1_GL4562 | 62551 | 62622 | L1MdTf_III | - | 71 |
| chr1_GL4562 | 62622 | 62939 | L1MdTf_III | - | 317 |
| chr1_GL4562 | 43940 | 50277 | L1MdTf_III | + | 6337 |
| chr1_GL4562 | 50277 | 51293 | L1MdTf_III | + | 1016 |
| chr5_JH5842 | 13873 | 14007 | L1MdTf_II | + | 134 |
| chr5_JH5842 | 14008 | 15000 | L1MdTf_II | + | 992 |
| chr7_GL4562 | 47633 | 53050 | L1MdTf_II | + | 5417 |
| chr7_GL4562 | 53050 | 54043 | L1MdTf_II | + | 993 |

**Table S4. TipseqHunter predicted non-reference insertions and manual curation**

| chr | start | end | label |
|---|---|---|---|
| chr1 | 20203193 | 20203497 | true insertions |
| chr1 | 62362101 | 62362407 | poor mapping quality |
| chr1 | 73943772 | 73944075 | mis-prime |
| chr1 | 99513888 | 99514192 | poor mapping quality |
| chr1 | 132231364 | 132231665 | mis-prime |
| chr1 | 138720350 | 138720652 | between L1s |
| chr1 | 178742415 | 178742716 | mis-prime |
| chr10 | 31236257 | 31236563 | true insertions |
| chr10 | 35898330 | 35898633 | true insertions |
| chr10 | 58313086 | 58313391 | mis-prime |
| chr10 | 61279396 | 61279701 | mis-prime |
| chr10 | 102657417 | 102657718 | mis-prime |
| chr11 | 14180217 | 14180518 | mis-prime |
| chr11 | 93470453 | 93470754 | mis-prime |
| chr11 | 99720511 | 99720814 | between L1s |
| chr12 | 5461550 | 5461851 | mis-prime |
| chr12 | 55366510 | 55366813 | mis-prime |
| chr12 | 88052321 | 88052627 | poor mapping quality |
| chr12 | 88297753 | 88298056 | poor mapping quality |
| chr12 | 95796324 | 95796626 | exisit L1 |
| chr12 | 113253280 | 113253585 | mis-prime |
| chr13 | 19188672 | 19188973 | mis-prime |
| chr13 | 21058217 | 21058527 | exisit L1 |
| chr13 | 64323492 | 64323794 | mis-prime |
| chr13 | 69735710 | 69736012 | mis-prime |
| chr13 | 77720438 | 77720739 | between L1s |
| chr13 | 82186217 | 82186527 | mis-prime |
| chr13 | 118414353 | 118414654 | poor mapping quality |
| chr14 | 14862644 | 14862951 | mis-prime |
| chr14 | 15413449 | 15413755 | poor mapping quality |
| chr14 | 15760314 | 15760615 | poor mapping quality |
| chr14 | 16920379 | 16920681 | true insertions |
| chr14 | 17273011 | 17273312 | poor mapping quality |
| chr14 | 18610623 | 18610924 | poor mapping quality |
| chr14 | 18768059 | 18768360 | poor mapping quality |
| chr14 | 18977121 | 18977423 | poor mapping quality |
| chr14 | 19007461 | 19007764 | poor mapping quality |
| chr14 | 36788509 | 36788814 | mis-prime |
| chr14 | 41479101 | 41479406 | poor mapping quality |
| chr14 | 42858202 | 42858507 | poor mapping quality |
| chr14 | 80328937 | 80329238 | exisit L1 |
| chr14 | 88546524 | 88546828 | poor mapping quality |
| chr15 | 88520502 | 88520807 | mis-prime |
| chr16 | 9569421 | 9569722 | mis-prime |
| chr16 | 67111043 | 67111344 | poor mapping quality |
| chr16 | 78831155 | 78831462 | true insertions |
| chr17 | 23019577 | 23019880 | poor mapping quality |
| chr17 | 24374911 | 24375215 | mis-prime |
| chr17 | 40155769 | 40156071 | mis-prime |
| chr17 | 41110025 | 41110330 | exisit L1 |
| chr18 | 29330901 | 29331203 | exisit L1 |
| chr18 | 79553508 | 79553815 | exisit L1 |
| chr2 | 6809826 | 6810127 | mis-prime |
| chr2 | 9220189 | 9220497 | exisit L1 |
| chr2 | 21868821 | 21869129 | mis-prime |
| chr2 | 75496553 | 75496855 | mis-prime |
| chr2 | 114330641 | 114330942 | exisit L1 |
| chr2 | 148788101 | 148788411 | true insertions |
| chr2 | 155483536 | 155483837 | mis-prime |
| chr3 | 56781247 | 56781555 | exisit L1 |
| chr3 | 139599308 | 139599613 | true insertions |
| chr3 | 146643889 | 146644191 | mis-prime |
| chr4 | 12234776 | 12235080 | mis-prime |
| chr4 | 17219027 | 17219333 | exisit L1 |
| chr4 | 106303136 | 106303437 | mis-prime |
| chr5 | 11040242 | 11040548 | poor mapping quality |
| chr5 | 11595091 | 11595398 | poor mapping quality |

| summary | |
|---|---|
| 17 | true insertions |
| 37 | mispriming |
| 26 | between close L1 or exist L1 |
| 36 | poor mapping quality |

| | | | |
|---|---|---|---|
| chr5 | 15104109 | 15104411 | poor mapping quality |
| chr5 | 15643423 | 15643731 | poor mapping quality |
| chr5 | 15886057 | 15886359 | poor mapping quality |
| chr5 | 34238268 | 34238569 | exisit L1 |
| chr5 | 42209869 | 42210171 | mis-prime |
| chr5 | 44638182 | 44638483 | mis-prime |
| chr5 | 58886063 | 58886364 | exisit L1 |
| chr5 | 77572550 | 77572851 | exisit L1 |
| chr5 | 82267080 | 82267382 | true insertions |
| chr5 | 126455014 | 126455323 | exisit L1 |
| chr5 | 145538514 | 145538817 | true insertions |
| chr6 | 56033929 | 56034230 | mis-prime |
| chr6 | 93512584 | 93512885 | mis-prime |
| chr6 | 111642744 | 111643048 | poor mapping quality |
| chr7 | 21080863 | 21081168 | true insertions |
| chr7 | 23880287 | 23880593 | mis-prime |
| chr7 | 69161596 | 69161897 | true insertions |
| chr7 | 120653943 | 120654246 | mis-prime |
| chr8 | 46292840 | 46293141 | mis-prime |
| chr8 | 50111895 | 50112198 | exisit L1 |
| chr8 | 57563653 | 57563955 | true insertions |
| chr9 | 16229368 | 16229672 | mis-prime |
| chr9 | 30342950 | 30343253 | poor mapping quality |
| chr9 | 66539847 | 66540150 | exisit L1 |
| chr9 | 79165010 | 79165311 | exisit L1 |
| chr9 | 91010627 | 91010929 | poor mapping quality |
| chr9 | 94733030 | 94733331 | poor mapping quality |
| chr9 | 97594457 | 97594758 | mis-prime |
| chrM | 971 | 1272 | poor mapping quality |
| chrX | 14623828 | 14624129 | poor mapping quality |
| chrX | 25236085 | 25236386 | true insertions |
| chrX | 26434584 | 26434888 | true insertions |
| chrX | 65565720 | 65566023 | true insertions |
| chrX | 67385687 | 67385988 | exisit L1 |
| chrX | 84995940 | 84996241 | poor mapping quality |
| chrX | 125601713 | 125602019 | poor mapping quality |
| chrX | 125657568 | 125657873 | poor mapping quality |
| chrX | 125664675 | 125664979 | poor mapping quality |
| chrX | 125672045 | 125672348 | poor mapping quality |
| chrX | 125676200 | 125676506 | poor mapping quality |
| chrX | 125685575 | 125685882 | poor mapping quality |
| chrX | 136331299 | 136331604 | exisit L1 |
| chrX | 145472097 | 145472402 | poor mapping quality |
| chrX | 157385521 | 157385822 | exisit L1 |
| chrX | 159525174 | 159525477 | true insertions |
| chrY | 21293618 | 21293924 | true insertions |

**Table S5. Coverage of annotation L1s identified by nanoTIPseq**

| | support reads | | | | | | |
|---|---|---|---|---|---|---|---|
| sample | >0 | >5 | >10 | >20 | >50 | total reads (10^3) | total base (Million base) |
| enzyme_B6_230112 | 3206 | 2917 | 2638 | 2100 | 929 | 200 | 129 |
| 4226_MDA_230321 | 3148 | 2873 | 2641 | 2183 | 1208 | 200 | 166 |
| 4226_PTA1 230327 | 3238 | 3125 | 3006 | 2579 | 1138 | 200 | 114 |
| 4226_PTA2_230327 | 3230 | 3118 | 2958 | 2569 | 1163 | 200 | 114 |
| 4226_PTA3_230327 | 3233 | 3151 | 3012 | 2648 | 1189 | 200 | 114 |
| 4226_PTA4_230327 | 3232 | 3066 | 2873 | 2383 | 1109 | 200 | 115 |
| 4226_PTA5_230327 | 3233 | 3114 | 2911 | 2446 | 992 | 200 | 111 |
| Sheared_B6_230217 | 3252 | 3207 | 3151 | 2953 | 1498 | 200 | 115 |
| B6+FVB_230218 | 3250 | 3199 | 3128 | 2833 | 848 | 200 | 126 |
| FVB_230218 | 2554 | 2142 | 2070 | 1857 | 620 | 200 | 121 |

| | support reads | | | | | | |
|---|---|---|---|---|---|---|---|
| sample | >0 | >5 | >10 | >20 | >50 | total reads (10^3) | total base (Million base) |
| enzyme_B6_230112 | 98.2% | 89.3% | 80.8% | 64.3% | 28.4% | 200 | 129 |
| 4226_MDA_230321 | 96.4% | 88.0% | 80.9% | 66.8% | 37.0% | 200 | 166 |
| 4226_PTA1 230327 | 99.1% | 95.7% | 92.0% | 79.0% | 34.8% | 200 | 114 |
| 4226_PTA2_230327 | 98.9% | 95.5% | 90.6% | 78.7% | 35.6% | 200 | 114 |
| 4226_PTA3_230327 | 99.0% | 96.5% | 92.2% | 81.1% | 36.4% | 200 | 114 |
| 4226_PTA4_230327 | 99.0% | 93.9% | 88.0% | 73.0% | 34.0% | 200 | 115 |
| 4226_PTA5_230327 | 99.0% | 95.3% | 89.1% | 74.9% | 30.4% | 200 | 111 |
| Sheared_B6_230217 | 99.6% | 98.2% | 96.5% | 90.4% | 45.9% | 200 | 115 |
| B6+FVB_230218 | 99.5% | 97.9% | 95.8% | 86.7% | 26.0% | 200 | 126 |
| FVB_230218 | 78.2% | 65.6% | 63.4% | 56.9% | 19.0% | 200 | 121 |

**Table S6. Predicted non-reference insertions from enzymatically digested gDNA/nanoTIPseq**

| | | |
|---|---|---|
| chr1 | 22443981 | 22444311 |
| chr1 | 164797542 | 164797730 |
| chr1 | 174993497 | 174993667 |
| chr10 | 35898398 | 35898637 |
| chr12 | 37250143 | 37250314 |
| chr12 | 41914804 | 41915096 |
| chr12 | 62017960 | 62018160 |
| chr14 | 101031794 | 101032227 |
| chr15 | 6728303 | 6728533 |
| chr19 | 50374116 | 50374315 |
| chr2 | 98497575 | 98497804 |
| chr2 | 144553351 | 144553515 |
| chr3 | 122695185 | 122695347 |
| chr4 | 52144534 | 52144844 |
| chr4 | 119674709 | 119675273 |
| chr5 | 15057713 | 15057907 |
| chr5 | 15643426 | 15643598 |
| chr5 | 15714371 | 15714561 |
| chr5 | 15781217 | 15781398 |
| chr5 | 15863028 | 15863201 |
| chr5 | 15886057 | 15886243 |
| chr5 | 18554902 | 18555062 |
| chr5 | 70361178 | 70361418 |
| chr5 | 77571396 | 77571566 |
| chr5 | 95118009 | 95118227 |
| chr5 | 126454831 | 126455363 |
| chr5 | 145538657 | 145538819 |
| chr6 | 6739319 | 6739501 |
| chr6 | 93309020 | 93309203 |
| chr7 | 20680640 | 20680825 |
| chr7 | 21080971 | 21081170 |
| chr7 | 51248681 | 51248840 |
| chr7 | 52586684 | 52586859 |
| chr7 | 56935796 | 56935966 |
| chr7 | 72271476 | 72271733 |
| chr7 | 87638054 | 87638292 |
| chr7 | 94960100 | 94960753 |
| chr8 | 20487141 | 20487303 |
| chr8 | 21166503 | 21166695 |
| chr8 | 21192770 | 21192956 |
| chr8 | 21228242 | 21228443 |
| chr8 | 96696448 | 96696649 |

| | | |
|---|---|---|
| false positive | 11 | |
| non-reference insertion | 43 | 81% |

| | | |
|---|---|---|
| chr9 | 3000598 | 3000781 |
| chrX | 3312957 | 3313139 |
| chrX | 4442610 | 4442919 |
| chrX | 5091050 | 5091230 |
| chrX | 30841485 | 30841655 |
| chrX | 31975219 | 31975423 |
| chrX | 33365163 | 33365322 |
| chrX | 33738049 | 33738230 |
| chrX | 33900273 | 33900443 |
| chrX | 80200864 | 80201022 |
| chrX | 81997786 | 81998005 |
| chrX | 125685714 | 125685875 |

**Table S7. Predicted non-reference insertions from sheared gDNA/nanoTIPseq**

| | | |
|---|---|---|
| chr1 | 109806349 | 109806744 |
| chr1 | 164797529 | 164797728 |
| chr1 | 182771638 | 182771823 |
| chr10 | 3542416 | 3542599 |
| chr10 | 6759135 | 6759318 |
| chr10 | 6936895 | 6937090 |
| chr10 | 10203482 | 10203653 |
| chr10 | 10285631 | 10285807 |
| chr10 | 14067789 | 14068006 |
| chr10 | 14078431 | 14078605 |
| chr10 | 14456638 | 14456849 |
| chr10 | 14935174 | 14935359 |
| chr10 | 15645404 | 15645583 |
| chr10 | 35898425 | 35898637 |
| chr12 | 37250143 | 37250335 |
| chr12 | 41914804 | 41915015 |
| chr12 | 49540974 | 49541160 |
| chr12 | 62017979 | 62018163 |
| chr12 | 65029363 | 65029533 |
| chr15 | 6728336 | 6728527 |
| chr15 | 45435394 | 45435564 |
| chr16 | 58052689 | 58052859 |
| chr18 | 57244576 | 57244762 |
| chr18 | 87271809 | 87271971 |
| chr19 | 50374143 | 50374315 |
| chr2 | 3050442 | 3050604 |
| chr2 | 98492738 | 98492959 |
| chr2 | 98496770 | 98497130 |
| chr2 | 101181322 | 101181496 |
| chr3 | 24880806 | 24881003 |
| chr4 | 3275451 | 3275621 |
| chr4 | 3706128 | 3706330 |
| chr4 | 6520166 | 6520336 |
| chr4 | 7144170 | 7144358 |
| chr4 | 7538019 | 7538192 |
| chr4 | 10056028 | 10056217 |
| chr4 | 10605187 | 10605373 |
| chr4 | 13342390 | 13342576 |
| chr4 | 14994207 | 14994411 |
| chr4 | 15324322 | 15324507 |
| chr4 | 15688061 | 15688268 |
| chr4 | 16708713 | 16709271 |
| chr4 | 17928110 | 17928280 |

| | | |
|---|---|---|
| chr4 | 18321958 | 18322153 |
| chr4 | 18671075 | 18671280 |
| chr4 | 19199005 | 19199175 |
| chr4 | 19961915 | 19962098 |
| chr4 | 24704072 | 24704250 |
| chr4 | 25456896 | 25457066 |
| chr4 | 26377548 | 26377727 |
| chr4 | 26495242 | 26495452 |
| chr4 | 27594017 | 27594167 |
| chr4 | 29269070 | 29269293 |
| chr4 | 90015795 | 90015961 |
| chr5 | 15083973 | 15084153 |
| chr5 | 15643426 | 15643606 |
| chr5 | 15654375 | 15654551 |
| chr5 | 15682498 | 15682668 |
| chr5 | 15714379 | 15714555 |
| chr5 | 15781166 | 15781396 |
| chr5 | 15886040 | 15886233 |
| chr5 | 82267279 | 82267448 |
| chr5 | 95118040 | 95118215 |
| chr5 | 109279414 | 109279584 |
| chr5 | 145538591 | 145538825 |
| chr7 | 20680642 | 20680821 |
| chr7 | 51248664 | 51248841 |
| chr7 | 56935796 | 56935966 |
| chr7 | 60995257 | 60995429 |
| chr7 | 61997075 | 61997258 |
| chr8 | 20487110 | 20487301 |
| chr8 | 21166460 | 21166691 |
| chr8 | 21192770 | 21192957 |
| chr8 | 65589348 | 65589549 |
| chr8 | 69110032 | 69110216 |
| chr9 | 3000604 | 3001159 |
| chr9 | 3001392 | 3001838 |
| chr9 | 105692517 | 105692704 |
| chr9 | 107023673 | 107023879 |
| chr9 | 107040277 | 107040469 |
| chr9 | 112685758 | 112685954 |
| chrX | 3312956 | 3313127 |
| chrX | 4442610 | 4442783 |
| chrX | 5091050 | 5091220 |
| chrX | 31975242 | 31975404 |
| chrX | 33311863 | 33312051 |
| chrX | 33365170 | 33365324 |
| chrX | 33738024 | 33738232 |

| | | |
|---|---|---|
| chrX | 33900273 | 33900443 |
| chrX | 81997828 | 81998004 |
| chrX | 125674936 | 125675105 |
| chrX | 125679077 | 125679274 |
| chrX | 125685683 | 125685881 |

**Table S8. Confirmation of selected non-reference insertions**

| CHROMOSOME | VALIDATION PRIMER | VALIDATION PRIMER | PCR CONFIRMED |
|---|---|---|---|
| **1**(164,797.576-7; minus) | TuJH922 | TuJH1083 | yes |
| 5(145,538,591-825; minus) | TuJH922 | TuJH1067 | yes |
| **7**(51,248,689-90; minus) | TuJH922 | TuJH1071 | yes |
| **7**(56,935,951-2; plus) | TuJH922 | TuJH1076 | yes |
| **8**(20,487,110-301; minus) | TuJH922 | TuJH1143 | yes |
| **10**(35,898,425-637; minus) | TuJH922 | TuJH1055 | yes |
| **12**(37,250,143-335; plus) | TuJH922 | TuJH1132 | no |
| **12**(41,914,804-5,015; plus) | TuJH922 | TuJH1060 | yes |
| **12**(62,017,979-8,163; minus) | TuJH922 | TuJH1087 | yes |
| **19**(50,374,143-315; minus) | TuJH922 | TuJH1063 | yes |
| **X**(3,312,993-4; minus) | TuJH922 | TuJH1095 | yes |
| **X**(4,442,610-783; plus) | TuJH922 | TuJH1108 | yes |
| **X**(5,091,199-200; plus) | TuJH922 | TuJH1112 | yes |
| **X**(31,975,242-404; minus) | TuJH922 | TuJH1099 | yes |
| **X**(33,311,863; minua) | TuJH922 | TuJH1103 | no |
| **X**(33,900,422-3; plus) | TuJH922 | TuJH1120 | yes |
| **X**(81,997,828-8,004; minus) | TuJH922 | TuJH1079 | yes |
| X(125,685,683-881; minus) | TuJH922 | TuJH1091 | yes |

confirmation by whole genome sequencing (different mouse)

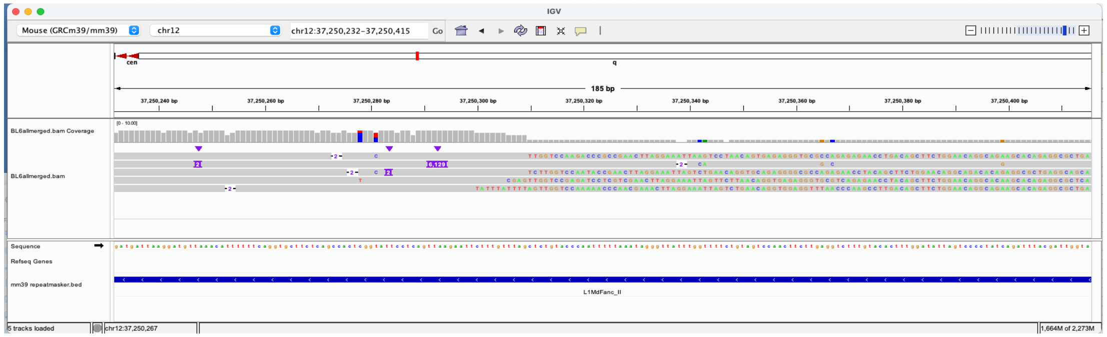


confirmation by whole genome sequencing (different mouse)

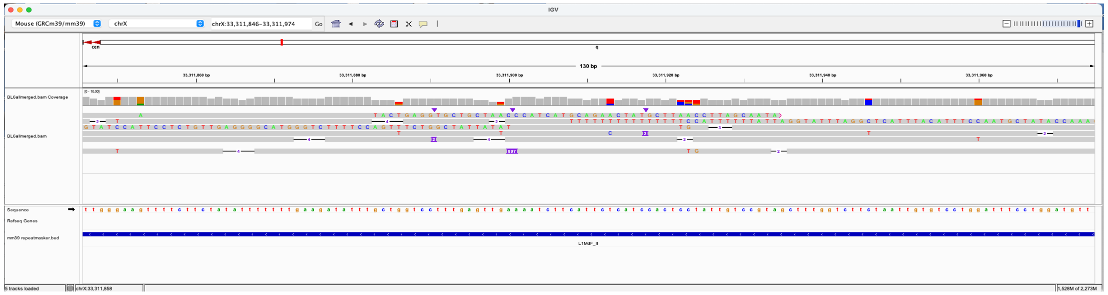

**Table S9. Quality control of whole genome amplification by chromosome PCRs**

| Sample | Chr 1 | Chr 2 | Chr 3 | Chr 4 | Chr 5 | Chr 6 | Chr 7 | Chr 8 | Chr 9 | Chr 10 | Chr 11 | Chr 12 | Chr 13 | Chr 14 | Chr 15 | Chr 16 | Chr 17 | Chr 18 | Chr 19 | Chr X |
|---|---|---|---|---|---|---|---|---|---|---|---|---|---|---|---|---|---|---|---|---|
| PTA-1 | y | y | y | y | y | y | y | y | y | y | y | y | y | y | y | y | y | y | y | y |
| PTA-2 | y | y | y | y | y | y | y | y | y | y | y | y | y | y | y | y | y | y | y | y |
| PTA-3 | y | y | y | y | y | y | y | y | y | y | y | y | y | y | y | y | y | y | y | y |
| PTA-4 | y | y | y | y | y | y | y | y | y | y | y | y | y | y | y | y | y | y | y | y |
| PTA-5 | y | y | y | y | y | y | y | y | y | y | y | y | y | y | y | y | y | y | y | y |
| MDA-1 | y | y | y | y | y | y | y | y | y | y | y | y | y | y | y | y | y | y | y | y |
| MDA-2 | y | y | y | y | y | y | y | y | y | y | y | y | y | y | y | y | y | y | y | y |

**Table S10. Unique non-reference L1s identified in 4226 cells.**

| | | |
|---|---|---|
| chr1 | 21986360 | 21986599 |
| chr1 | 96648322 | 96648492 |
| chr1 | 110747550 | 110747751 |
| chr1 | 111907832 | 111908198 |
| chr1 | 112065835 | 112066017 |
| chr1 | 115305634 | 115305803 |
| chr1 | 116220581 | 116220807 |
| chr1 | 116430762 | 116431263 |
| chr1 | 117729582 | 117729771 |
| chr1 | 122388723 | 122388893 |
| chr1 | 122775713 | 122775887 |
| chr1 | 122905585 | 122905756 |
| chr1 | 125518879 | 125519066 |
| chr1 | 140637368 | 140637580 |
| chr1 | 145296966 | 145297142 |
| chr1 | 145693024 | 145693222 |
| chr1 | 147133371 | 147133552 |
| chr1 | 147149216 | 147149413 |
| chr1 | 147516061 | 147516256 |
| chr1 | 147770792 | 147770965 |
| chr1 | 147772040 | 147772221 |
| chr1 | 148130479 | 148130727 |
| chr1 | 148688721 | 148688911 |
| chr1 | 151143261 | 151143440 |
| chr1 | 151153628 | 151153941 |
| chr1 | 152167213 | 152167398 |
| chr11 | 103512941 | 103513139 |
| chr11 | 103853216 | 103853425 |
| chr13 | 109111811 | 109111992 |
| chr16 | 10035833 | 10036001 |
| chr16 | 27541445 | 27541615 |
| chr16 | 27665347 | 27665570 |
| chr17 | 51372871 | 51373068 |
| chr17 | 52376208 | 52376446 |
| chr19 | 60530342 | 60530536 |
| chr3 | 39087748 | 39087946 |
| chr3 | 39101373 | 39101694 |
| chr3 | 39133897 | 39134099 |
| chr3 | 41939522 | 41939748 |
| chr3 | 142829271 | 142829453 |
| chr4 | 74038222 | 74038400 |
| chr4 | 84968817 | 84969001 |
| chr4 | 85680908 | 85681119 |

| | | |
|---|---|---|
| chr4 | 85958397 | 85958584 |
| chr4 | 86890867 | 86891037 |
| chr4 | 87139575 | 87139771 |
| chr4 | 88245639 | 88245841 |
| chr4 | 94324736 | 94324947 |
| chr4 | 94329395 | 94329589 |
| chr4 | 94677081 | 94677429 |
| chr4 | 96831649 | 96831853 |
| chr5 | 110139490 | 110139677 |
| chr6 | 23947540 | 23947787 |
| chr6 | 106762350 | 106762523 |
| chr9 | 101364249 | 101364420 |
| chr9 | 101528567 | 101528797 |
| chr9 | 101790357 | 101790558 |
| chr9 | 101837332 | 101837531 |
